# Secreted SARS-CoV-2 ORF8 modulates the cytokine expression profile of human macrophages

**DOI:** 10.1101/2021.08.13.456266

**Authors:** Nisha Kriplani, Sara Clohisey, Sonia Fonseca, Sarah Fletcher, Hui-Min Lee, Jordan Ashworth, Dominic Kurian, Samantha J. Lycett, Christine Tait-Burkard, J. Kenneth Baillie, Mark E. J. Woolhouse, Simon R. Carding, James P. Stewart, Paul Digard

**Affiliations:** The Roslin Institute, University of Edinburgh, Easter Bush, Midlothian, EH25 9RG, UK; Usher Institute, Ashworth Laboratories, Kings Buildings, Charlotte Auerbach Road, University of Edinburgh, Edinburgh, EH9 3FL, UK; Quadram Institute Bioscience, Rosalind Franklin Road, Norwich Research Park, Norwich, NR4 7UQ, UK; Norwich Medical School, University of East Anglia, Norwich Research Park, Norwich, NR4 7TJ, UK; Department of Infection Biology and Microbiomes, University of Liverpool, LSP IC2, Brownlow Hill, Liverpool, L3 5RF, UK

**Keywords:** ORF8, virokine, monocyte, accessory protein, ER stress, L84S

## Abstract

Severe acute respiratory syndrome coronavirus 2 (SARS-CoV-2) is still adapting to its new human host. Attention has focussed on the viral spike protein, but substantial variation has been seen in the *ORF8* gene. Here, we show that SARS-CoV-2 ORF8 protein undergoes signal peptide-mediated processing through the endoplasmic reticulum and is secreted as a glycosylated, disulphide-linked dimer. The secreted protein from the prototype SARS-CoV-2 virus had no major effect on viability of a variety of cell types, or on IFN or NF-κB signalling. However, it modulated cytokine expression from primary CSF1-derived human macrophages, most notably by decreasing IL-6 and IL-8 secretion. Furthermore, a sequence polymorphism L84S that appeared early in the pandemic associated with the Clade S lineage of virus, showed a markedly different effect, of increasing IL-6 production. We conclude that *ORF8* sequence polymorphisms can potentially affect SARS-CoV-2 virulence and should therefore be monitored in sequencing-based surveillance.

## Introduction

Severe Acute Respiratory Syndrome coronavirus-2 (SARS-CoV-2), the causative agent of the COVID-19 pandemic, is a positive sense RNA virus belonging to the *Coronaviridae* family [1]. Its RNA genome of around 30 kb contains 12 major open reading frames (ORFs), which with polyprotein processing and various alternative translation events encode upwards of 30 proteins [2]. Based on their function, these can be variously classified as replication proteins (*e.g.* the many products of the Orf1a/b polyprotein), structural proteins (including S, M, E and N) and accessory proteins that include the products of ORF3a, ORF6, ORF7a/b, ORF8, ORF9b/c and ORF10 [3]. Coronavirus accessory proteins are not necessarily essential for replication of the virus *in vitro* and tend to be less well conserved between different viral species [3–8], Nonetheless, accessory proteins are often involved in immune modulation and can play essential roles in virulence, pathogenesis, and host-adaptation. In line with this, SARS-CoV-1 acquired a 29 bp deletion disrupting *ORF8* early in the epidemic, which was associated with host adaptation [9–12]. Similarly, in the early stages of the COVID-19 pandemic, the SARS-CoV-2 *ORF8* gene was an early hotspot of mutational change, with several studies identifying a coding change (L84S) as being under significant selection pressure [13–16]. Large gene deletions affecting SARS-CoV-2 *ORF8* have also been seen, and as with SARS-CoV-1, these, as well as the L84S polymorphism, have been linked to changes in disease severity [8, 17–19]. Understanding the function of SARS-CoV-2 ORF8 polypeptide is therefore likely to provide insights into viral adaptation and pathogenesis. An intact *ORF8* gene is non-essential for SARS-CoV-1 replication in tissue culture, while the intact ORF8 polypeptide was shown to be glycosylated and partially resident in the endoplasmic reticulum (ER), as well as being associated with inhibition of innate immune signalling [20–22]. Bioinformatics analyses of SARS-CoV-2 ORF8 similarly indicated the presence of a cleavable signal peptide (SP) [23, 24] and consistent with this, a subsequent study detected SP-dependent secretion of ORF8 in transfected cell culture supernatants as well as in the serum of patients infected with SARS-CoV-2 [25]. SARS-CoV-2 ORF8 has also been proposed to inhibit various steps of the interferon signalling pathway [26–29], although not all studies have confirmed this [30]. Two studies have also linked ORF8 expression to induction of ER stress and the unfolded protein response [26, 31]. ORF8 has also been suggested to facilitate immune evasion by directly interacting with and downregulating MHC-I molecules in various cell types [32]. Conversely, it has been suggested to contribute to “cytokine storms” through an interaction with the IL-17 receptor [33]. Here, we confirm SP-dependent secretion of SARS-CoV-2 ORF8 and show that the secreted form is predominantly a disulphide-linked dimer glycosylated at position 78. Furthermore, we show that secreted ORF8 is capable of modulating the cytokine response of human blood monocytes-derived macrophages (MDMs) in a manner specifically affected by the L84S polymorphism.

## Materials and Methods

### Cell culture and transfections

A549 (Human adenocarcinoma alveolar basal epithelial cell line), X293T (Human Embryonic Kidney cell line) and HEK-Blue IFN α/β cells (InvivoGen) were cultured in Dulbecco’s modified eagle’s medium (DMEM, Sigma Aldrich) with 10% (v/v) Foetal Bovine Serum (FBS) (Life Technologies), 2 mM L-Glutamine (Life Technologies), 100 U/mL penicillin (Gibco) and 100 μg/mL streptomycin (Gibco). For HEK-Blue IFNα/β cells, 30 μg/mL blasticidin (InvivoGen) and 100 μg/mL zeocin (InivoGen) were also added to complete medium. Similarly, the human monocyte cell line THP1-Blue expressing an NF-κB-inducible secreted embryonic alkaline phosphatase (SEAP) reporter gene (InvivoGen) were cultured in RPMI-1640 (Sigma Aldrich) with 10% (v/v) FBS (Biosera), 2 mM L- Glutamine, 100 U/mL penicillin, 100 μg/ml streptomycin, 30 μg/mL blasticidin and 100 μg/mL normocin. To maintain selection pressure, 10 μg/ml blasticidin was added every other passage. Primary human CD14^+^ monocytes were isolated from fresh blood from volunteer donors under ethical approval from the Lothian Research Ethics Committee (11/AL/0168) as described before [34]. Primary monocytes were seeded in 96-well or 48-well cell culture plates in RPMI supplemented with 10% (v/v) FBS, 2 mM L- Glutamine, 100 U/mL penicillin, 100 μg/ml streptomycin and 100 ng/ml rhCSF-1 for 7 days to facilitate differentiation into macrophages. Stimulation-based experiments were performed on 8^th^ day. Plasmid and Poly I:C (InvivoGen) transfections were done using Lipofectamine-2000 (Life Technologies) transfection reagent according to the manufacturer’s instructions. Cell viability was quantified by measuring ATP content using CellTitre-Glo reagent (Promega) according to manufacturer’s instructions.

### Plasmid constructs

*ORF8* untagged and FLAG-tagged synthetic cDNA sequences (from MN994468.1; Wuhan seafood market pneumonia virus isolate 2019-nCoV/USA-CA2/2020) were ordered from Integrated DNA Technologies Inc. Synthetic cDNAs for two naturally occurring polymorphisms: L84S and V62L/L84S; and two functional mutants: cleavage signal (CS) dead and signal peptide (SP) dead (Figure 1A), were ordered in FLAG epitope-tagged versions. Both untagged and tagged versions of *ORF8* cDNAs were subcloned into a strong expression vector pVR1255 [35] by replacing a Firefly luciferase gene between NotI and BamHI restriction sites. An N-linked glycosylation site mutation (N78Q) was introduced by site directed mutagenesis using the primer pair, CATCGATATCGGTCAATATACAGTTTCCTG and CAGGAAACTGTATATTGACCGATATCGATG. An expression construct for human short palate, lung and nasal epithelium clone 1 (SPLUNC 1) (also called BPI fold containing family A, member 1 (BPIFA1) was constructed as follows. The hSPLUNC1 coding sequence with a 3’ sequence encoding the FLAG tag was amplified from cDNA using KOD polymerase (Sigma) and the primers (Forward) 5’ ATGCGGCCGCCGCCGCCACCATGTTTCAAACTGGAGGCCTC 3’ and (Reverse) 5’ CAGATCTTTACTTGTCATCGTCGTCCTTGTAGTCGACCTTGATGACAAACTGTAGTC 3’. The Forward primer incorporated a NotI site and consensus translational start site and the Reverse, a BglII site and the FLAG-encoding sequence. The amplified sequence was cut with NotI and BglII, inserted between the NotI and BamHI sites of pVR1255 and verified by sequencing.

**Figure 1:**
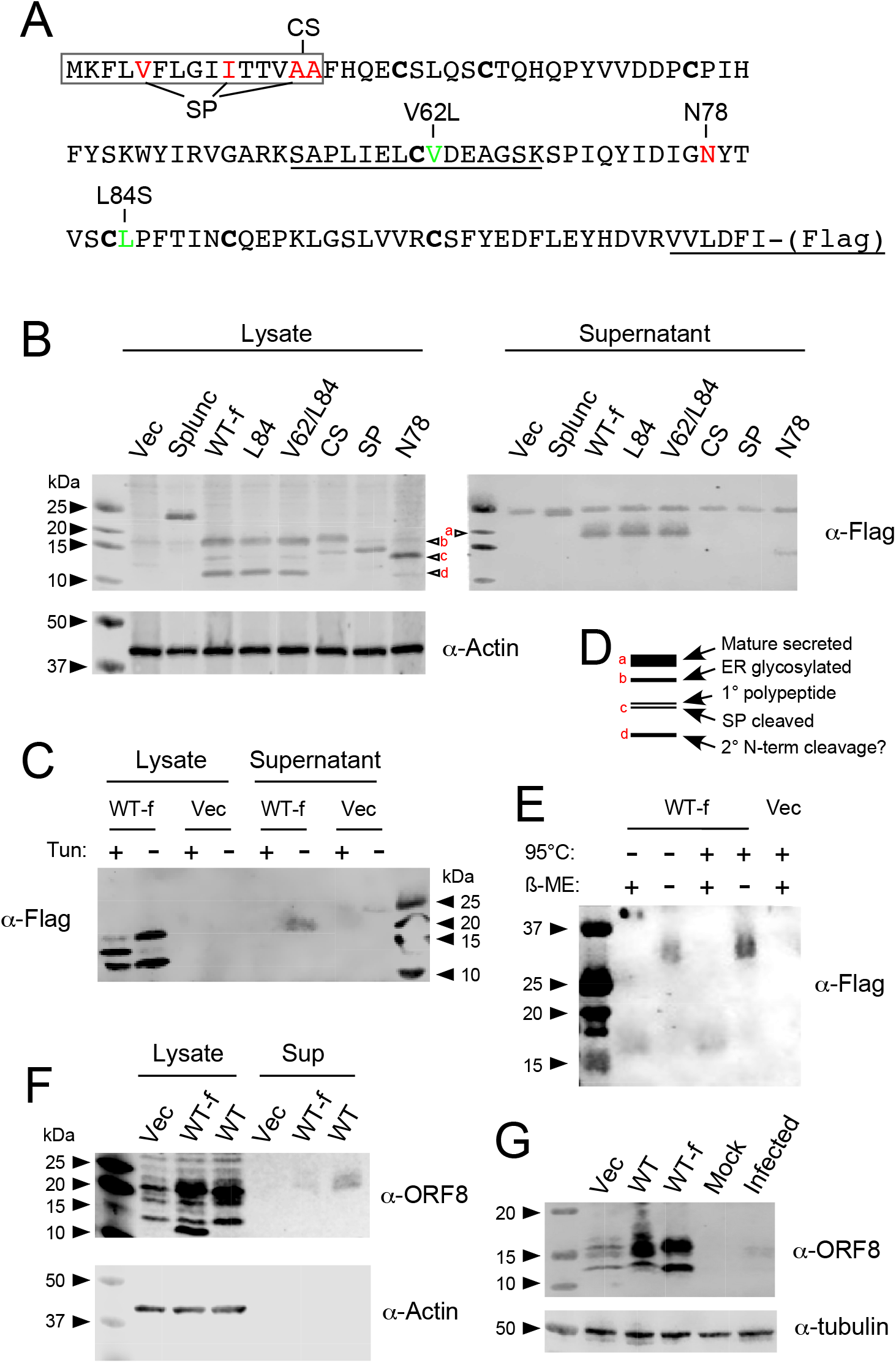
Biochemical analysis of SARS-CoV-2 ORF8. (A) Amino acid sequence of ORF8. Green colour represents naturally occurring variants and red colour represent artificial functional mutants of ORF8. (FLAG) indicates a FLAG epitope tag (DYKDDDDK) present in some constructs. Peptides identified by mass spectrometry of secreted WT-f polypeptide are underlined. (B) X293T cells were transfected with the indicated plasmids and 48h post-transfection cells lysates and supernatants were examined by SDS-PAGE and western blotting. (C) Vector control or ORF8-transfected X293T cells were treated with Tunicamycin (5 μg/ml) for 24h hours. Cell lysates and supernatants were examined by western blotting. (D) Schematic explanation of different ORF8 species observed by western blotting. (E) Cell lysate and supernatants from ORF8 WT-f-expressing cells were examined by western blotting with or without β-mercaptoethanol treatment and/or heating the samples at 95°C. (F) Untagged and FLAG-tagged ORF8 cell lysates and supernatants were immunoblotted using ORF8 antibody. (G) Vero cells were infected with SARS-CoV2 at MOI of 1 and 24h post-infection cell lysates were examined by western blotting. WT denotes untagged ORF8, WT-f denotes wildtype ORF8 with a FLAG tag; all ORF8 and Splunc clones were FLAG-tagged. Red lettering on (B) and (D) indicates the forms of ORF8 produced by the WT-f construct. Blot images were modified during figure preparation by separation of individual colour channels, changing colour mode to greyscale followed by inversion and linear adjustment of brightness and contrast settings, as well as cropping to remove unwanted parts of the image.

### Production and testing of ORF8 supernatants

To produce cell supernatants containing secreted ORF8 polypeptides, 10 cm dishes of X293Ts were transfected with 5 μg empty vector or ORF8 expression plasmids. 6h later media was replaced with fresh complete medium. 48 h post transfection supernatants were harvested, clarified by centrifugation at 2100 x g for 5 min, aliquoted and stored at − 80°C. For treatment of X293T or A549 cells, supernatants were diluted 1:2 with fresh medium before adding to cells, for MDMs, supernatants were diluted 1:3. For simultaneous treatment with IFNβ, human type I IFN (R&D systems, 11415-1) was typically added at a final concentration of 150 U/ml. For treatment with poly I:C (Invivogen), cells were transfected with 2.5 μg (A549 cells) or 15 μg (MDMs) using Lipofectamine (Life Technologies) according to standard protocols. For double treatment with ORF8 and poly I:C, ORF8-containing supernatants were applied to the cells first, followed immediately by the transfection mixes.

### SARS-CoV2 infections

Virus isolate EDB-2 (Scotland/EDB1827/2020, with an intact S1/S2 furin cleavage site was UK isolated, propagated and titred in Vero E6 (ATCC CRL-1586) cells [36]. Vero E6 cells were cultured in DMEM (Lonza), 10% HI FBS (Gibco), 1X ultraglutamine (Lonza), and 1X non-essential amino acids solution (Invitrogen) and infected (at confluence) with passage 2 SARS-CoV-2 EDB-2 at an MOI of 1. At 24 hours post infection (hpi), the cells were lysed in 1x final concentration Laemmli sample buffer (Biorad) containing 100 mM DTT.

### Immuno-precipitation and western blotting

Cells were transfected with control vector or ORF8 expressing vectors. 48h post-transfection, supernatants were collected and clarified by centrifugation (3000 rpm for 5 minutes). Cells were washed with PBS and lysed in Laemmli buffer containing β-mercaptoethanol (unless otherwise indicated). A fraction of clarified supernatant was mixed with 4X Laemmli buffer in 3:1 ratio for western blotting. The remaining supernatants were incubated with magnetic FLAG beads (Sigma) overnight at 4° C. Beads were harvested and washed with TBS (50 mM Tris-Cl, 150 mM NaCl, pH 7.5) prior to mixing with Laemmli buffer. Cell lysates, supernatants, and magnetic beads (in Laemmli buffer) were then heated at 95° C for 10 minutes before resolving them in 10-20% SDS-PAGE gels (Novex, Life Technologies) and then transferred onto nitrocellulose membranes (VWR) using the Trans-Blot® Turbo™ Transfer System (Bio-Rad). Membranes were blocked using 1% BSA in PBS-T at room temperature for 1 hour and then incubated with primary antibodies, anti-FLAG (Cell Signalling Technologies, 2908S), anti-SARS-CoV-2-ORF8 (MRCPPU, University of Dundee, DA088, [37]) and anti-Tubulin (BioRad, MCA77G). Donkey anti-rabbit IRDye 800 and goat anti-rat IRDye 680 (Li-cor Biosciences) secondary antibodies were used to detect bound IgG, followed by imaging on a Licor Odyssey instrument.

### Mass spectrometry

LC-MS analysis was performed using a RSLCnano system (Thermo Fisher Scientific) coupled to a micrOTOF QII mass spectrometer (Bruker) on in-gel trypsin digested protein bands excised from Coomassie Blue-stained SDS-PAGE gels [38].

### Immunofluorescence

A549 cells seeded on glass coverslips were washed with PBS and then fixed with 4% formaldehyde in PBS (Sigma) for 20 min prior to permeabilisation with 1% Triton X-100 in PBS for 5 min at room temperature. Primary antibodies, anti-FLAG (Cell Signalling, 2368S) and anti-Calreticulin (Abcam, ab2907), were used at 1:800 dilution, Alexa Fluor-conjugated secondary antibodies (Donkey α-Mouse Alexa-Fluor 488 for Flag and Donkey α-Mouse Alexa-Fluor 594 for Calreticulin, Life Technologies) were used at 1:2000 and nuclei were stained using Hoechst 33342 at 1:5000. Staining steps were performed sequentially at room temperature in PBS/1% FBS with PBS-T wash steps in between, followed by mounting in ProLong Gold antifade mount (Thermo Fisher Scientific). Cells were imaged using a Zeiss LSM 710 confocal microscope under a 63X objective. All images shown are single optical slices.

### Cytokine and immune signalling assays

To measure type I interferons or NF-κB activation, 20 μl of cell supernatants were incubated with HEK-Blue™ IFNα/β cells or THP1-Blue™ NF-κB cells respectively. For antagonist experiments IFNβ or lipopolysaccharide (LPS; Sigma Aldrich, L2630) were added at indicated concentrations. 24 hours later, 20 μl of supernatants from the HEK-Blue™ or THP1-Blue™ cells were mixed with 180 μl of QuantiBlue reagent (InvivoGen) and optical density (O.D.) was measured at 650nm using a plate reader (CLARIOstar, BMG Labtech). To measure IL-6, ELISAs were performed on clarified supernatants (undiluted) using Human IL-6 ELISA kit (Life Technology) according to manufacturer’s instructions. Statistical tests (one-way ANOVA using repeated measures between samples from the same experiment or donor as appropriate, or mixed-effects analysis in cases where sample loss had occurred, followed by Dunnett’s multiple comparison tests) were performed using Graphpad Prism 8.4.3.

### Cytokine profiler assay

A subset of clarified (by centrifugation at 400 x g for 10 minutes) supernatants from MDMs of three donors were pooled together. Cytokine profiling of these pooled supernatants was performed using a human cytokine array kit (human cytokine array kit, panel A; R&D Systems) and near-infrared fluorescence detection using IRDye 800CW Streptavidin (Li-Cor Biosciences).

## Results

### The ORF8 polypeptide undergoes complex processing in the secretory pathway

Inspection of the ORF8 protein sequence indicates a small protein of 121 aa (molecular weight ~11kDa) with a predicted cleavable signal peptide at the N-terminus, no further transmembrane domain and an N-glycosylation motif at amino acid 78, as well as an abundance of cysteine residues (Figure 1A). Taken together, these suggested the hypothesis that ORF8 is a secreted viral chemokine, or “virokine”. Analysis of naturally occurring ORF8 polymorphisms early in the COVID-19 pandemic using sequences available on GISAID [39] shows that the first recorded L84S mutation was detected in one of the 24 sequences now available from Wuhan, China in December 2019 (Strain Wuhan_IME-WH01, Genbank accession MT291826). In January, February and March 2020, the L84S mutation was present in 35%, 18% and 11% of the available GISAID sequences (Table S1, Figure S1), corresponding to Nextstrain clade 19B, GISAID Clade S (98% of L84S sequences) or several Pango A lineages (20% A[basal], 37% A.1, 18% A.2 of L84S sequences). Subsequently, the first recorded V62L mutations were detected in sequences from China and Hong Kong in January 2020. The majority (87%) of the V62L mutations co-occur with L84S in Clade S or Pango lineage A[basal] (70%) and A.3 (28%) for the period December 2019-March 2020 (Figure S1), and these sequences with both L84S and V62L were mostly (86%) detected in North America (Figure S2). The early appearance and spread of L84S possibly along with V62L is suggestive of host adaption. As a first test of these hypotheses, we constructed a series of C-terminally FLAG-tagged ORF8 constructs and expressed them in X293T cells, along with (as a positive control for a small cellular secreted protein [40]) a similarly FLAG-tagged version of the human secreted protein Splunc1. Western blots of intracellular material showed a single species of the expected size for Splunc1 and only background reactivity from cells transfected with empty plasmid vector (Figure 1B, left hand panel). The wild type ORF8 FLAG construct (WT-f) as well as the L84 and V62/L84 constructs produced three polypeptides ranging in apparent molecular weight between ~ 11kDa and 16kDa. Mutation of the signal peptide (SP) cleavage signal (CS) led to the loss of the smallest 11kD band and a slight upwards size shift of the two larger species, whereas mutation of the SP led to the disappearance of both top 16kDa and bottom 11kDa bands as well as more prominent accumulation of the middle species. Mutation of the glycosylation motif (N78Q) also led to a marked reduction in the accumulation of the top and bottom polypeptides and a more prominent central band, but without a mobility shift. Analysis of FLAG tag immunoprecipitated supernatants from the cells confirmed the secretion of Splunc1 and all three natural variants of ORF8, with the latter all running as heterogeneous products at around 20 kDa, slightly larger than the presumed intracellular glycosylated form. Mass spectrometric analysis of the secreted WT-f product showed the presence of two peptides from ORF8 (underlined, Figure 1A). Secretion of ORF8 was blocked by the CS and SP mutations and reduced by alteration of the glycosylation motif (Figure 1B, right hand panel). Treatment with tunicamycin, an inhibitor of glycosylation and inducer of ER stress [41, 42] blocked secretion of the WT-f ORF8 polypeptide and shifted the balance of the two larger intracellular isoforms towards the middle species (Figure 1C). Overall, this suggests that the primary ORF8 polypeptide is processed by removal of the SP and an initial round of glycosylation in the ER, followed by further glycosylation and secretion of this mature form (Figure 1D). A final intracellular species apparently results from further N-terminal cleavage in the secretory pathway.

The crystal structure of ORF8 produced from *Escherichia coli* and refolded *in vitro* indicates a disulphide-linked dimer [43]. To test if this occurred for the fully glycosylated form synthesised in mammalian cells, we analysed the mature secreted form by SDS-PAGE with and without reducing agent in the sample buffer. In the absence of β-mercaptoethanol, a prominent band migrating at the size expected for a dimer was seen, as well a small amount of the monomer form (Figure 1E), indicating that ORF8 is primarily secreted from mammalian cells as a disulphide-linked dimer.

To test if the addition of the FLAG tag affected biosynthesis of ORF8, we examined cell lysates and supernatants from cells transfected with either empty vector, the WT-f construct or one containing an untagged ORF8 gene by western blotting with an anti-ORF8 polyclonal antibody [37]. Similarly sized polypeptides were detected from both tagged and untagged constructs in the cell lysate and supernatants (Figure 1F), suggesting that the FLAG tag had no major effect on ORF8 secretion. Finally, we examined lysates from cells transfected with the ORF8 constructs or infected with SARS-CoV-2, which revealed that authentic virus infection produced a similarly-sized polypeptide to the exogenously-expressed protein (Figure 1G), again validating the synthetic constructs and overall, consistent with the virokine hypothesis.

To test if the subcellular location of ORF8 was consistent with the inferences drawn from sequence and mutational analyses, A549 cells transfected with WT-f ORF8 or empty vector were fixed, stained with anti-FLAG and anti-calreticulin (as a marker of the ER) and examined by confocal microscopy. Anti-FLAG staining gave a reticular pattern in the cytoplasm as well as outlining the nuclear rim, showing partial colocalization with the calreticulin staining (Figure 2), consistent with the biochemical analyses. Examination of cells transfected with the L84S, V62L/L84S, CS and N78Q mutants gave similar results (Figure S3). In contrast, the SP mutant gave a more punctate cytoplasmic distribution which did not colocalise with the ER.

**Figure 2:**
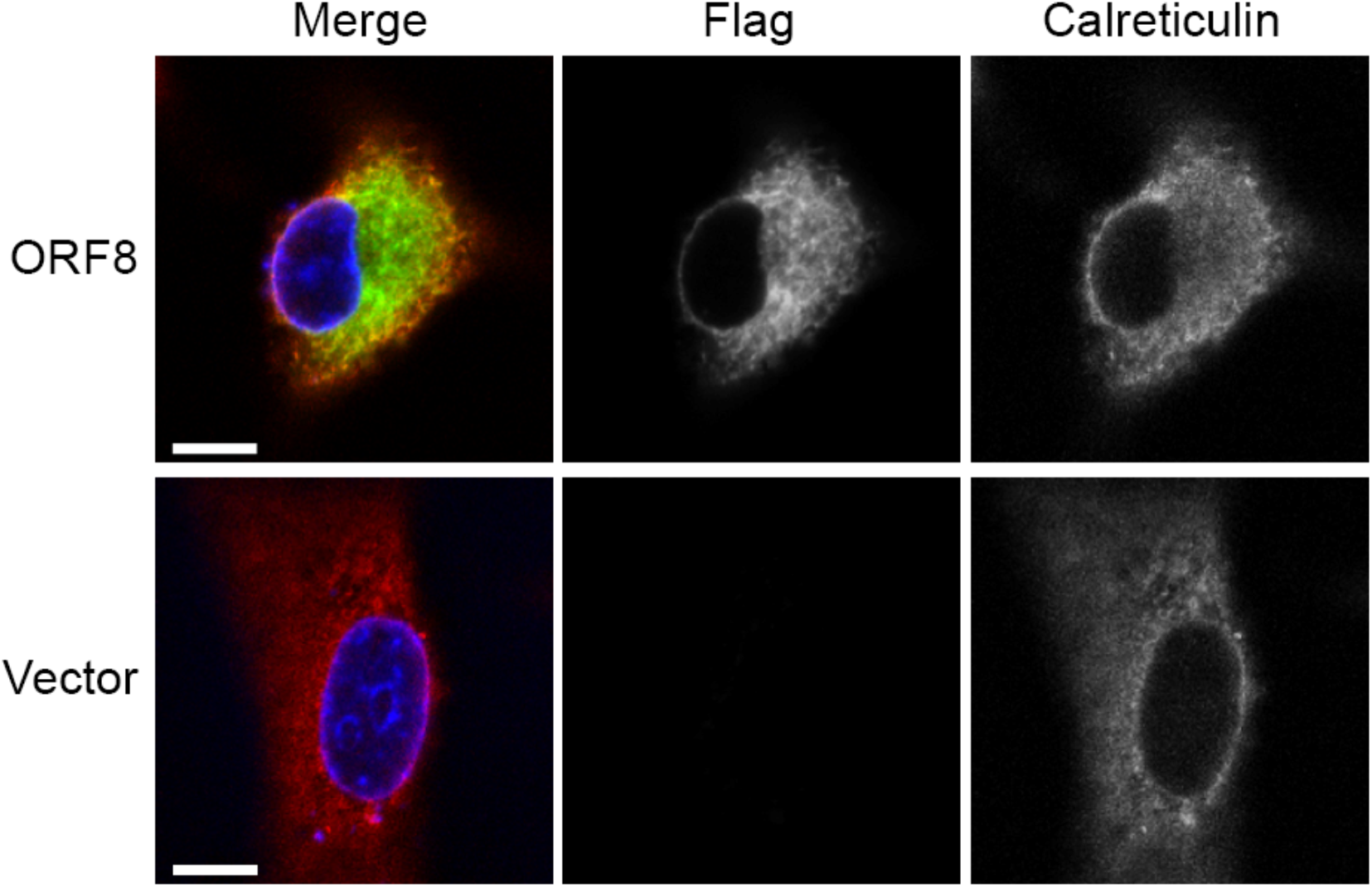
Subcellular localisation of ORF8. A549 cells were transfected with vector control or ORF8 WT-f expressing construct. 48h post-transfection cells were fixed and stained with anti-FLAG (green) and ER marker Calreticulin (red), as well as for DNA with Hoechst (blue). Cells were imaged using a confocal microscope. Scale bar = 10μm. Colour channels were adjusted individually using Adobe Photoshop during figure preparation to ensure visibility and the images were cropped to increase visibility of the chosen cells. All adjustments were linear and applied to all images equally. See also Fig S3.

### ORF8 has little effect on IFN and NF-κBsignalling

ORF8 homologues from SARS-CoV-1 i.e., ORF8ab, ORF8a and ORF8b, have been reported to variously induce apoptosis, modulate cellular DNA synthesis, induce protein degradation (by ubiquitination) and interfere with the host immune response [5, 8, 44]. Less functional information is available on SARS-CoV-2 ORF8 but here too it has been implicated in immune evasion [27, 45]. Our virokine hypothesis predicted that secreted ORF8 would act as a paracrine modulator of cell function; either by inducing a response and/or by reducing the response of cells to another stimulus. To test this, supernatants from transfected X293T cells were harvested and assayed for effects when added to the medium over fresh cells. Unlike SARS-CoV-1 ORF8 polypeptides [44, 46], ORF8 of SARS-CoV-2 either in expressed or secreted form did not notably affect cell viability of X293T or A549 cells (Figure S4), although A549 cells appeared sensitive to apoptosis induced by expression of the SP mutant (Figure S3). Next, we tested the role of secreted ORF8 in interferon (IFN) signalling. When added to A549 cells, the ORF8 polypeptides did not induce detectable type I IFN secretion as assessed by bio-assay using HEK-Blue cells, and nor did they inhibit type I IFN secretion when the cells were stimulated by poly I:C transfection (Figure 3A). Moreover, when an IFNβ dose response curve was performed in the presence of secreted ORF8 (either WT or the L84S variant), no significant effects were observed (Figure 3B). The effects of secreted ORF8 on NF-κB signalling were also assessed, using THP1-Blue-NF-κB cells; a reporter cell line that contains a SEAP reporter gene under an NF-κB-inducible promoter [47]. Supernatants from ORF8 expressing cells or control cells were added to the reporter cells with or without the NF-κB signalling activator LPS. However, the ORF8-containing supernatants did not induce NF-κB signalling on their own and nor did they exhibit any antagonistic effect against LPS-induced responses (Figure 3C). Thus, secreted ORF8 had no obvious effect on IFN signalling in epithelial cell models or NF-κB signalling in a monocyte model.

**Figure 3:**
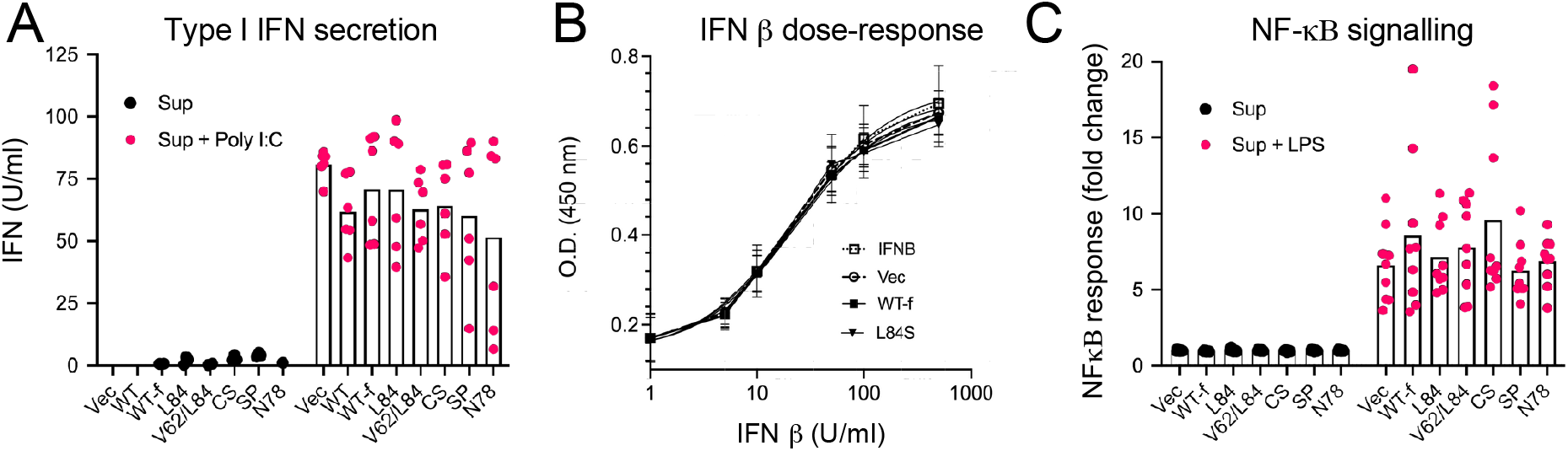
Secreted ORF8 does not affect NF-κB or IFN signalling. (A) ORF8 containing supernatants were added onto A549 cells, either alone or in combination with Poly I:C (2.5 μg/ml). 24 h later supernatants from A549s were harvested and assayed for type I Interferon using HEK -Blue IFNα/β cells. Type I IFN amounts were extrapolated from a standard curve and values below baseline are not plotted. Data points are from two independent experiments each done in triplicate. Bar indicates the mean. (B) An IFN β dose-response curve analysis was performed on HEK-Blue IFN α/β cells in the absence (IFN) or presence of control (Vec) or ORF8 (WT-f or L84S)-containing supernatants. Data are the mean ± SD of three independent experiments each performed in triplicate and have been fitted with a three-parameter agonist-response curve in Graphpad Prism. All R^2^ values were > 0.93 and the estimated EC_50_ values were not significantly different (p > 0.05). (C) Control or various ORF8-containing supernatants were added to THP-1 Blue cells in the presence or absence of LPS (10 ng/ml) and NF-κB response was measured using QuantiBlue. Data points from three independent experiments done in triplicate. Bars indicate the mean.

### Secreted ORF8 alters the cytokine expression profile of primary human macrophages

Macrophages are an indispensable component of the innate immune system and upon viral infections, alter their cytokine profile to mount a greater anti-viral response [48]. A dysregulated monocytic response is thought to be key to severe pathology in SARS-CoV-2 infections [49]. Accordingly, we next investigated the effect of ORF8 on the cytokine/chemokine secretion profile of primary human macrophages. Monocyte-derived macrophages (MDMs) were incubated with supernatants containing the panel of ORF8 polypeptides in the presence or absence of poly I:C or IFNβ. Individual samples showed variability, perhaps reflecting differences in primary cells from donors, but overall, none of the secreted ORF8 or control cell supernatants had any significant effect on MDM viability, whilst transfection with poly I:C or treatment with IFNβ reduced viability by around 50% (Figure 4A). As with A549 cells, the addition of ORF8-containing supernatants to MDMs did not induce type I IFN production by human macrophages and nor did it significantly alter type I IFN production after they were stimulated with poly I:C (Figure 4B).

**Figure 4:**
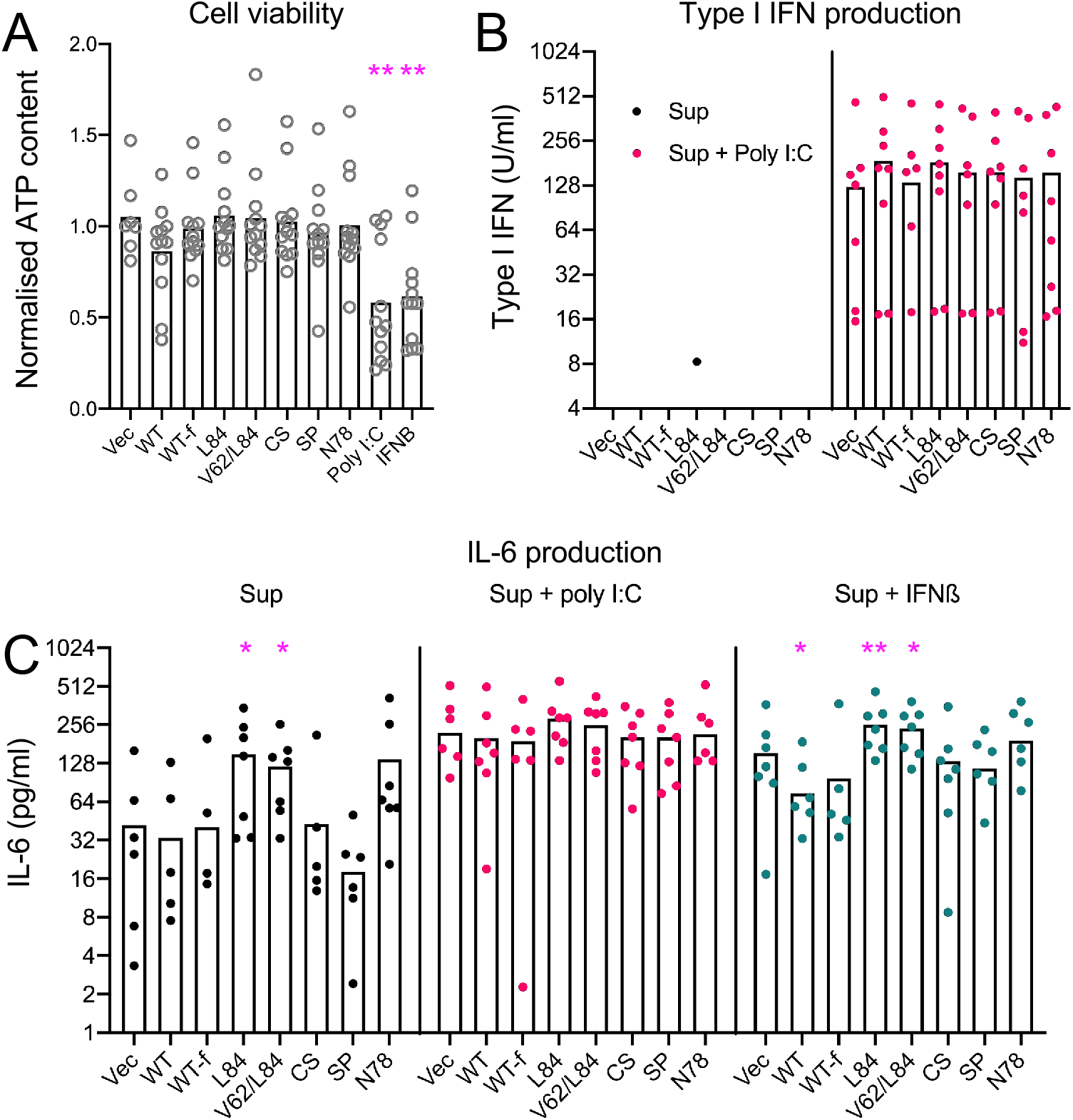
Differential regulation by ORF8 variants of IL-6 production from human peripheral blood monocyte-derived macrophages. (A) Cell viability of MDMs was assayed after treatment for 24 h with ORF8 supernatants, IFNβ (150 U/mL) or Poly I:C (15 μg/mL). Data are from cells from seven donors. ** = p < 0.01 following one-way ANOVA followed by Dunnett’s multiple comparison tests against the mean of the Vec samples. (B) Supernatants from MDMs treated with ORF8 supernatants with or without concomitant Poly I:C stimulation were assayed for type I IFN using HEK-Blue IFN α/β cells. Data points are from the same seven donors as (A). Values below the assay limit of detection (4 U/ml) are not plotted. No significant differences were seen within groups (one-way ANOVA followed by Dunnett’s multiple comparison tests). (C) Secreted IL-6 levels from MDMs in response to ORF8 supernatants in the presence or absence of Poly I:C or IFNβ were assessed by IL-6 ELISA. Data are the mean of seven donors * = p < 0.05; ** = p < 0.01 (one -way ANOVA for IL-6 only, mixed-effects analysis for poly I:C and IFN β samples followed by Dunnett’s multiple comparisons test against the matching Vec samples).

The severity of COVID-19 disease has been linked to magnitude of the IL-6 response [50, 51], so we next examined the effect of secreted ORF8 on production of this cytokine. Cells treated with supernatant from vector-transfected cells produced on average around 50 pg/ml IL-6 and this was not significantly altered by either tagged or untagged WT ORF8, or supernatants from cells transfected with the non-secreted CS or SP mutant forms of the protein (Figure 4C, left hand panel). In contrast, the L84 and V62/L84 mutants significantly increased IL-6 production by around 3-fold. A similar trend (although not statistically significant) was seen for the non-glycosylated N78 mutant. Stimulation of MDMs with poly I:C increased IL-6 production to around 250 pg/ml but against this background, none of the ORF8 supernatants had any significant effect (Figure 4C, middle panel). Stimulation of the cells with IFNβ also increased IL-6 secretion by the cells (to around 150 pg/ml) in the absence of ORF8. The two WT forms of ORF8 modestly decreased IL-6 secretion by around two-fold (Figure 4C, right hand panel), but both the L84 and V62/L84 ORF8 variants significantly increased IL-6 production to around 250 pg/ml, while the other mutant forms of the protein had no significant effect. Thus, ORF8 shows sequence-specific modulation of IL-6 production by human primary MDMs.

These effects of ORF8 on IL-6 production by MDMs led us to investigate the effects on a wider range of cytokine/chemokines. For this, MDM supernatants from individual donors stimulated with control or ORF8-containing supernatants with or without additional IFNβ treatment were pooled into three groups and the levels of various secreted cytokines and chemokines were quantified using an antibody capture array. Plotting the averaged data as a heat map showed that treatment of MDMs with both WT and L84 versions of secreted ORF8 in the absence of IFN was moderately suppressive to cytokine secretion, with most being reduced and none enhanced more than 2-fold (Figure 5A). Five cytokines showed greater than 2-fold reductions: IL-8, CXCL1 and TNFa by the WT-f supernatant, and CXCL10 and MIF by L84S ORF8. IFN treatment of the MDMs in the absence of ORF8 had a greater effect on cytokine production, with MIP-1, CXCL11, CCL5 and CXCL10 being upregulated more than 5-fold and several down regulated more than 2-fold (Figure 5A). Both ORF8 polypeptides generally flattened the response to IFN treatment, with greater suppression of IL-2 by both ORF8 variants, IL-8 by WT but not L84S ORF8, conversely IL-27 production by L84S but not WT ORF8 and, as previously noted, differential effects on IL-6 release (Figure 5B). Thus, secreted ORF8 modulates cytokine production by primary human macrophages in a manner that varies according to naturally-occurring sequence polymorphisms.

**Figure 5:**
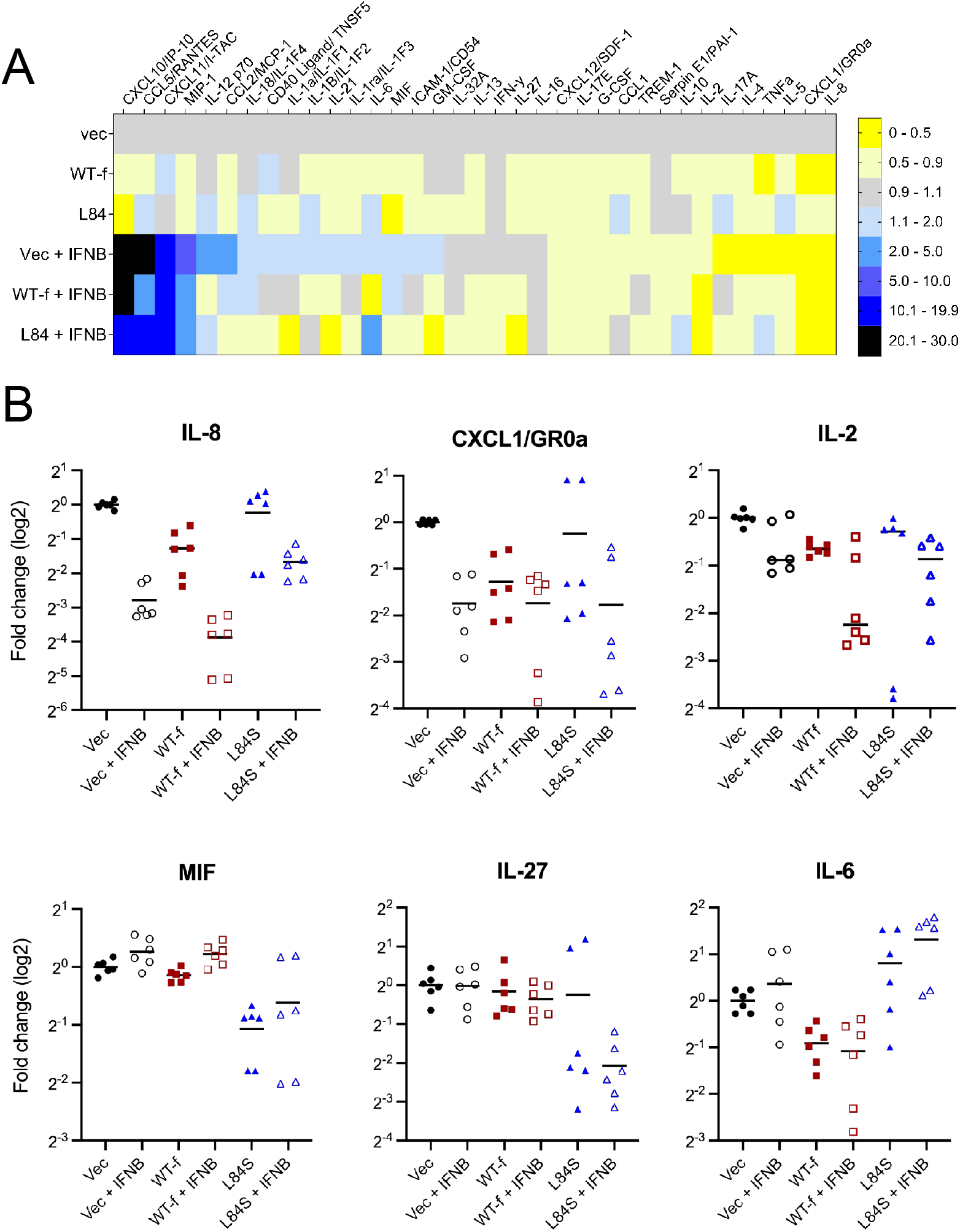
Secreted ORF8 alters the cytokine profile of monocyte derived macrophages. Cell supernatants from MDMs treated with FLAG-tagged WT or L84S ORF8-containing or control supernatants with or without concomitant IFN stimulation were assayed for the indicated cytokines using antibody capture arrays imaged on a Licor Odyssey instrument, after pooling into three batches, each from 2-3 individual donors (see Figure 4). (A) Heatmap of data normalised to cells treated with control (Vec) supernatants without IFN treatment. Cytokines are ordered according to the magnitude of the response seen with IFN treatment in the absence of ORF8. (B) Individual data points are shown for selected cytokines. Data are the technical replicates from the three individual pools. Bars indicate the mean.

## Discussion

Here, we confirm that SARS-CoV-2 ORF8 is secreted via a cleavable SP-dependent mechanism, further showing that this is largely in the form of a disulphide-linked dimer glycosylated at position 78. We also provide evidence that secreted ORF8 acts as an immunomodulator, altering the cytokine profile of human primary MDMs in a way that depends on the ORF8 L84S sequence polymorphism.

Consistent with a previous study of SARS-CoV-2 ORF8 [25], we found SP-dependent secretion of WT and L84S ORF8 and could trace a biosynthetic pathway up to a mature glycosylated and secreted disulphide-linked dimer. We also detected an additional smaller species (migrating at ~ 11 kDa) of unknown origin, that retained the C-terminal FLAG tag, thus most likely following an N-terminal cleavage event. This product was less abundant following mutation of either the SP, the SP cleavage site or the N-linked glycosylation signal, suggesting it is produced in the ER. Like others [26, 31], we find that ORF8 over-expression induces ER stress (data not shown); whether formation of this isoform of ORF8 is involved in that process (or possibly other functions) remains to be determined.

Some, but not all studies have linked SARS-CoV-2 ORF8 to interference with innate immune signalling [26, 27, 29, 32]. Here, we found no major effect (positive or negative) of secreted ORF8 on IFN secretion, response, or NF-κB signalling in A549, X293T and THP-1 cells respectively. In this respect, our data are in broad agreement with a study that compared the response of human primary nasal epithelial cells to infection with WT or a naturally occurring ORF8 deletion (Δ382 variant) and found no substantial difference in the IFN response [52]. The reason(s) behind these discrepant results remain to be determined; cell type and assay method seem likely contributors. Cell type did play a role in the major effect we found for secreted ORF8 - modulating IL-6 production by human primary MDMs, as we did not recapitulate this effect in A549 cells, despite their being competent for IL-6 secretion (data not shown). Nevertheless, our finding that secreted ORF8 can alter cytokine secretion by MDMs supported our original hypothesis (also proposed by others [14]) that it has an immunomodulatory “virokine” function. Furthermore, our data indicating that the two common ORF8 polymorphisms present early in the pandemic had differential effects is consistent with work associating the L84S variant with reduced clinical severity of disease [19]; although perhaps counterintuitive because L84S increased IL-6 production in our system and higher IL-6 levels are accepted as prognostic of poorer clinical outcomes [50, 51].

The mechanism by which ORF8 modulates cytokine expression remains to be determined. We have not tested purified ORF8 polypeptides so cannot exclude the possibility that ORF8 synthesis in cells induced the release of factors that alter MDM function (though this would be analogous to natural infection where infected epithelial cells will be the source of secreted ORF8), but the simplest hypothesis is that ORF8 acts directly on one or more MDM receptors. Based on structural predictions and supported by interactome analyses, ORF8 has been suggested to interact with integrins [14, 28]. It has also been shown to interact with the IL-17 receptor [33]. Neither observation provides an immediate explanation of how IL-6 and other cytokine release by MDMs is perturbed, so further analysis is required. It is also not clear how the L84S polymorphism affects ORF8 function. We saw no obvious differences in intracellular processing or secretion and consistent with this, biochemical analyses indicate no substantial effect on protein stability other than decreased sensitivity to acidic (~ pH 6) conditions [53]. The L84S variant is largely associated with the early clade S lineage of SARS-CoV-2 and the available surveillance sequencing suggests it is currently (summer 2021) restricted geographically to parts of Africa, Central America and northern regions of South America (Nextstrain.org). However, with the subsequent rise of the Variant of Concern (VOC) Alpha (B.1.1.7) in autumn 2020, viruses with truncated *ORF8* (mutation Q27*) have been circulating. During Spring 2021, the VOC Delta (B.1.617.2 and others) strains which have mostly L84 and V62 have risen to dominance, and by June 2021 less than 0.1% of available sequences have L84S in an intact ORF8 polypeptide. This implies that L84S does not provide a major fitness advantage to SARS-CoV-2. Nevertheless, our work revealing that ORF8 and its variants can modulate cytokine expression by a cell type that is highly relevant to viral pathogenesis suggests that further work into the protein so that sequence data from viral variant surveillance can be more fully interpreted is worthwhile.

## Acknowledgments

PD, CTB, SL and JKB acknowledge Strategic Programme Grant support from the Biotechnology and Biological Sciences Research Council (BBSRC) (no.

BB/P013740/1). MEJW and SL received funding from the European Union’s Horizon 2020 research and innovation programme under grant agreement No. 874735 (VEO) (https://www.veo-europe.eu/). SRC and SF acknowledge the support of BBSRC Institute Strategic Programme Grant Gut Microbes and Health BB/R012490/1 and its constituent project(s) BBS/E/F/000PR10353 and BBS/E/F/000PR10355. JPS acknowledges support from the MRC (MR/W005611/1) and BBSRC (BB/R00904X/1; BB/R018863/1).

We also thank the anonymous donors who provided blood for experiments, as well as authors from the Originating laboratories responsible for obtaining the specimens and the Submitting laboratories where genetic sequence data were generated and shared via the GISAID Initiative, on which this research is based (supplementary tables S2 and S3).

## Author contributions

Conceptualisation, PD, JPS, MW, NK; Investigation, NK, SRC, both SFs, H-ML, JA, DK, SL, CT-B, PD; Writing – Original Draft, NK, PD; Writing – Review & Editing, NK, SRC, both SFs, H-ML, DK, CT-B, SC, JPS, PD; Funding Acquisition, PD, CT-B, JKB, SL, MEJW, SRC, JPS; Supervision, PD, CT-B, JKB, SRC, MEJW.

