## Supplementary Figs 1-4 and Table S1 for "Secreted SARS-CoV-2 ORF8 modulates the cytokine expression profile of human macrophages"

### Supplementary Material

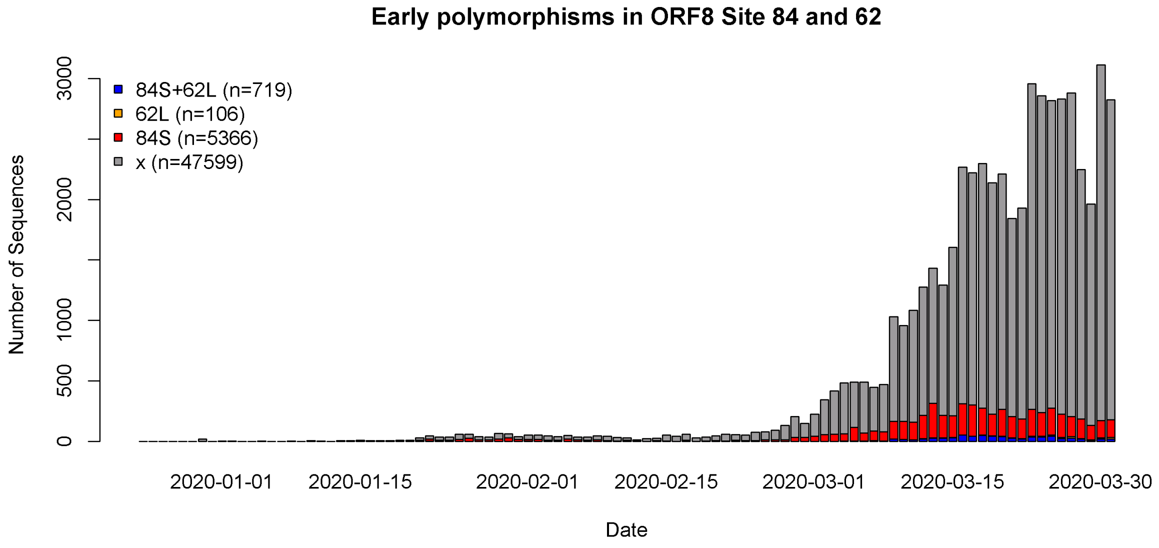


**Figure S1: Number of ORF8 sequences per day from Dec 2019 – March 2020, with L84S (red), V62L (orange), L84S and V62L (blue), or neither mutation (grey).** Sequences not labelled as human are excluded, and the minority of sequences with only month information are randomly distributed across the month.

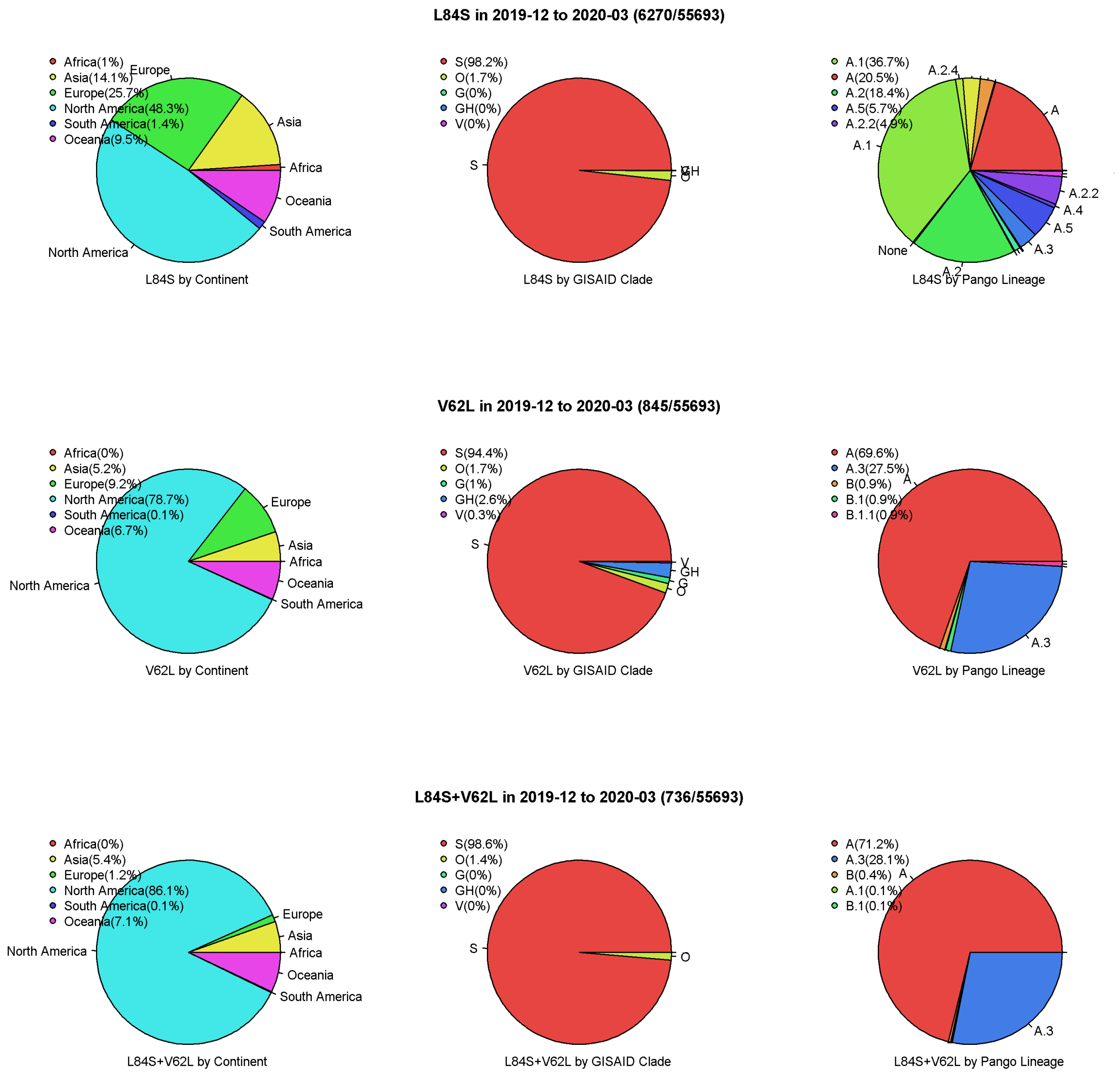


**Figure S2: Distribution of sequences with L84S, V62L and L84S and V62L from Dec 2010-March 2020 by continent, GISAID clade and Pango Lineage.** The majority of sequences with these mutations are from North America, in GISAID clade S and Pango A lineages. Most of the V62L mutations co-occur with L84S.

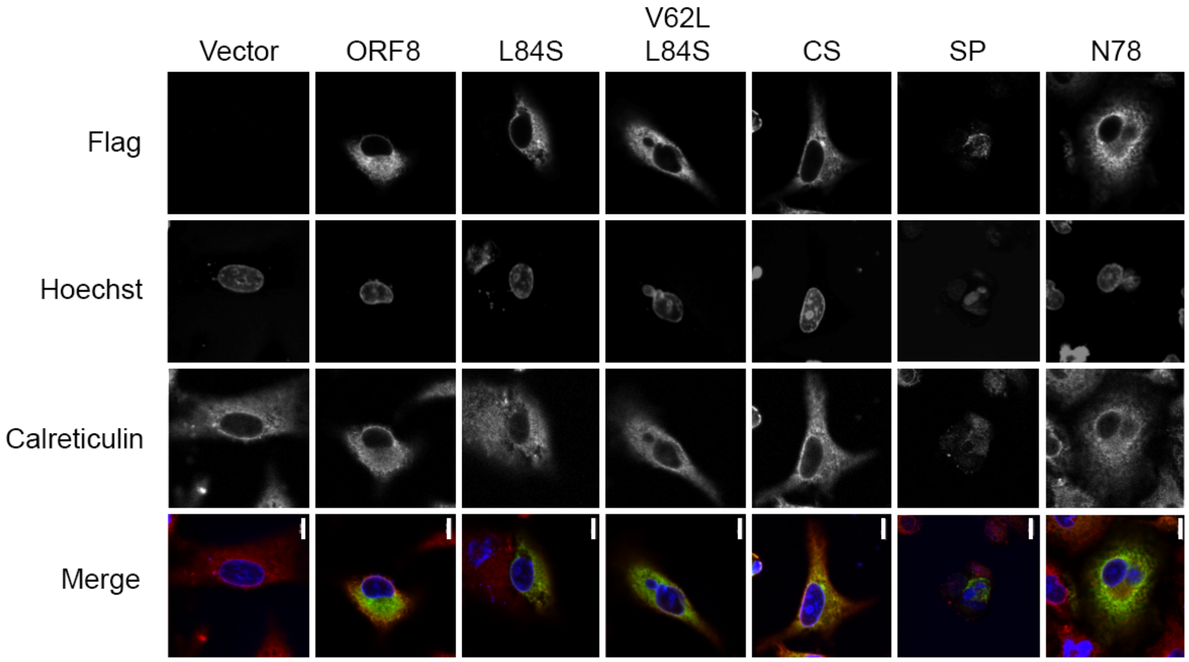


**Figure S3: Subcellular localisation of ORF8 WT-f and mutant proteins.** A549 cells were transfected with vector control or ORF8 WT-f expressing constructs as labelled. 48h post-transfection cells were fixed and stained with anti-FLAG (green) and ER marker Calreticulin (red), as well as for DNA with Hoechst (blue). Cells were imaged using a confocal microscope and a 63x objective. Images are single optical slices. Scale bars = 10µm. Note that Vec and ORF8 images are repeated from Figure S1 (without cropping), and that expression of the SP mutant typically induced cytotoxicity, as seen by apparently apoptotic nuclei. Colour channels were adjusted individually using Adobe Photoshop during figure preparation to ensure visibility. All adjustments were linear and applied to all images equally.

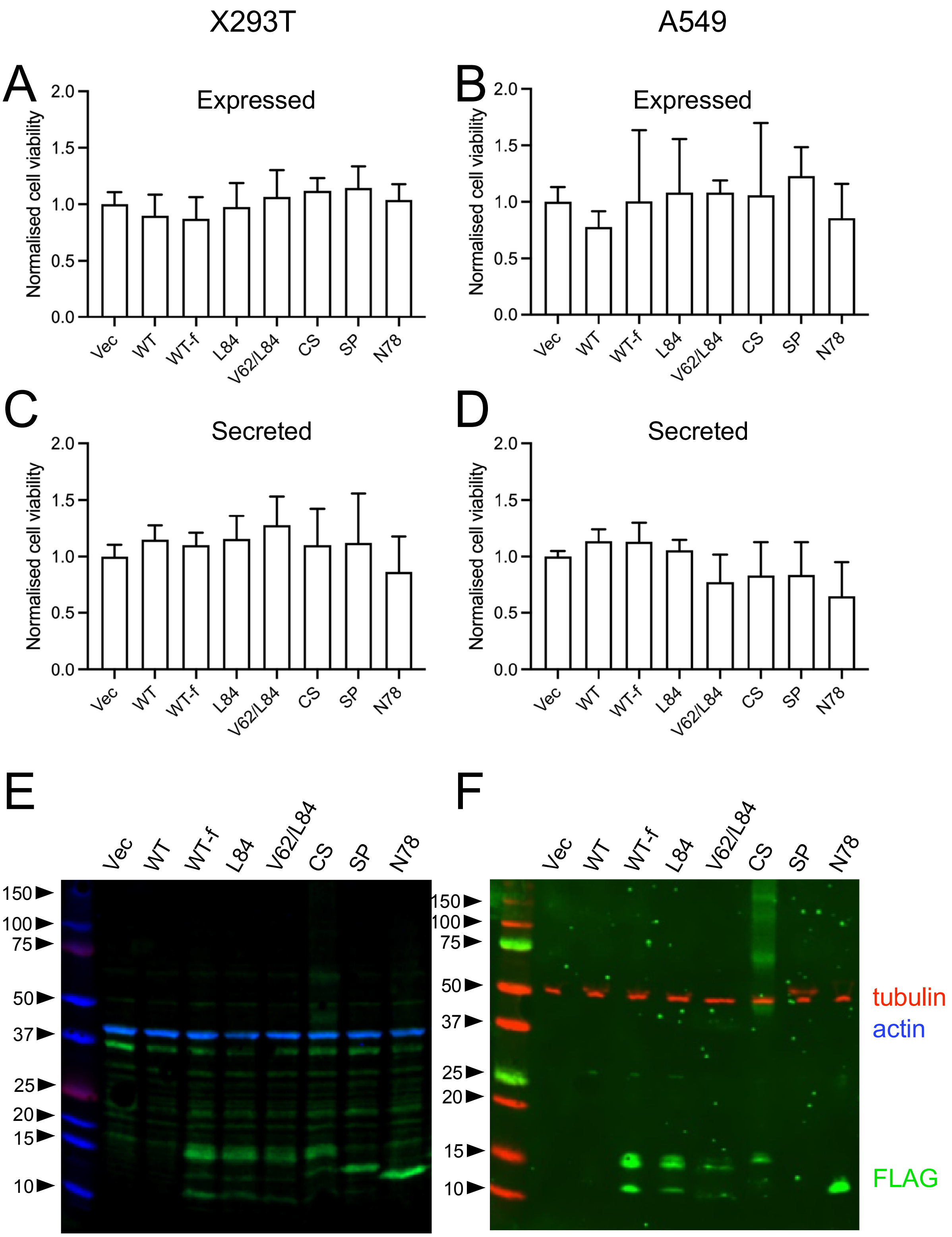


**Figure S4: ORF8 is not overtly cytotoxic in cultured epithelial cells**. X293T or A549 cells were transfected with the indicated plasmids and 24h later, (A, B) intracellular ATP content measured to estimate cell viability. Data are the mean ± SD of 2 independent experiments each with 3 technical replicates. (C, D) In parallel, cell supernatants were harvested and used to treat further dishes of cells for 24h, after which cell viability was measured as above. Data are the mean ± SD of three independent experiments each with three technical replicates. (E,F) cell lysates from the transfected cells were also analysed by western blotting with anti-FLAG (green) to confirm ORF8 expression and anti-actin (blue) or anti-tubulin (red) as loading controls. Note that WT (untagged) ORF8 will not be detected by this method. In addition, the mutant SP mutant expressed very poorly in A549 cells; from the appearance of the few transfected cells, because of the induction of cell death (see Fig S1). The higher molecular weight products visible in the CS mutant were not consistently seen in all experiments (data not shown).

**Tables**

| Year-Month | L84S | V62L | Q27* |
|---|---|---|---|
| **2019-12** | 4.2 | 0.0 | 0.0 |
| **2020-01** | 34.8 | 2.4 | 0.2 |
| **2020-02** | 17.6 | 1.1 | 0.1 |
| **2020-03** | 10.9 | 1.6 | 0.0 |
| **2020-04** | 4.0 | 1.1 | 0.1 |
| **2020-05** | 2.9 | 0.5 | 0.1 |
| **2020-06** | 1.0 | 0.4 | 0.1 |
| **2020-07** | 0.5 | 0.6 | 0.1 |
| **2020-08** | 0.3 | 0.4 | 0.1 |
| **2020-09** | 0.2 | 0.4 | 0.1 |
| **2020-10** | 0.1 | 0.6 | 0.2 |
| **2020-11** | 0.1 | 0.6 | 2.5 |
| **2020-12** | 0.4 | 0.6 | 14.6 |
| **2021-01** | 0.6 | 0.6 | 32.8 |
| **2021-02** | 0.8 | 0.7 | 49.8 |
| **2021-03** | 0.8 | 0.4 | 67.7 |
| **2021-04** | 0.6 | 0.3 | 73.4 |
| **2021-05** | 0.4 | 0.3 | 69.9 |
| **2021-06** | 0.1 | 0.1 | 26.3 |

**Table S1: Percentage of sequences with mutations from wild-type to main alterative amino acid (or stop codon) at sites 84, 62 and 27.** Sequences containing X or – (gap) are excluded from the percentage calculation, hence 84S and 62L especially from December 2020 onwards are being compared to the remaining full length ORF8 sequences.

All ORF8 protein sequences were downloaded from the GISAID database (https://www.gisaid.org) in August 2021 (1st August 2021 data set) and summarised with custom R scripts. We gratefully acknowledge the Authors from Originating and Submitting laboratories of sequence data on which the analysis is based.

**Table S2: GISAID Acknowledgement table for strains from Wuhan, China isolated in Dec 2019.** These are also published on Genbank with accession numbers: MN90894; MT019529; MT291826; MT291827; MT291828; MT291829; MT291830; MT019530; MT019531; MT019532; MN996527; MN996528; MN996529; MN996530; MN996531. Strain hCoV-19/Wuhan/IME-WH01/2019 (EPI_ISL_529213, MT291826, Wuhan_IME-WH01) has the L84S substitution.

**Table S3: GISAID Acknowledgement table for all GISAID Clade S strains from Dec 2019-March 2020.**
