## Supplementary Table 2 for "Secreted SARS-CoV-2 ORF8 modulates the cytokine expression profile of human macrophages"

We gratefully acknowledge the following Authors from the Originating laboratories responsible for obtaining the specimens, as well as the Submitting laboratories where the genome data were generated and shared via GISAID, on which this research is based.

All Submitters of data may be contacted directly via www.gisaid.org

Authors are sorted alphabetically.

| Accession ID | Originating Laboratory | Submitting Laboratory | Authors |
|---|---|---|---|
| EPI_ISL_529213, EPI_ISL_529214, EPI_ISL_529215, EPI_ISL_529216, EPI_ISL_529217 | Beijing Institute of Microbiology and Epidemiology | Beijing Institute of Microbiology and Epidemiology | Cui, Y.; Fan; Guo, Y.; Hang; Hou, J.; Li, B.; Mi, Z.; Mu, J.; Qin, E.; Song; Teng; Wu, Y.; Xu, Z.; Yajun.; Yang, R.; Yong, Y.; Yue; Zhang, X. |
| EPI_ISL_406798, EPI_ISL_406799 | General Hospital of Central Theater Command of People's Liberation Army of China | BGI & Institute of Microbiology, Chinese Academy of Sciences & Shandong First Medical University & Shandong Academy of Medical Sciences & General Hospital of Central Theater Command of People's Liberation Army of China | Weifeng Shi and Zhenhong Hu; Weijun Chen; Yuhai Bi |
| EPI_ISL_402123, EPI_ISL_403929, EPI_ISL_403930, EPI_ISL_403931 | Institute of Pathogen Biology, Chinese Academy of Medical Sciences & Peking Union Medical College | Institute of Pathogen Biology, Chinese Academy of Medical Sciences & Peking Union Medical College | Chao Wu; Jianwei Wang; Lili Ren; Qi Jin; Yiwei Liu; Zhiqiang Wu; Zichun Xiang |
| EPI_ISL_402125 | National Institute for Communicable Disease Control and Prevention (ICDC) Chinese Center for Disease Control and Prevention (China CDC) | National Institute for Communicable Disease Control and Prevention (ICDC) Chinese Center for Disease Control and Prevention (China CDC) | Chen; Dai; F.-H.; Hu, Y.; J.-H.; J.-J.; J.-L. and Zhu; Liu, Y.; Pei; Q.-M.; She; Song; T.-Y.; Tao; Tian; Wang; Wang, W.; Wu, F.; Xu, L.; Y.-L.; Y.-M.; Y.-Y.; Y.-Z.; Yu, B.; Z.-G.; Z.-W.; Zhang; Zhao, S.; Zheng |
| EPI_ISL_402119, EPI_ISL_402121 | National Institute for Viral Disease Control and Prevention, China CDC | National Institute for Viral Disease Control and Prevention, China CDC | Faxian Zhan , Weifeng Shi , Baoying Huang , Jun Liu , Li Zhao , Yao Meng , Fei Ye , Na Zhu; Ji Wang; Weimin Zhou; Weimin Zhou , Peihua Niu , Peipei Liu , Faxian Zhan , Weifeng Shi , Baoying Huang , Jun Liu , Li Zhao , Yao Meng , Xiaozhou He , Fei Ye , Na Zhu , Yang Li , Jing Chen , Wenbo Xu , George F. Gao , Guizhen Wu; Wenjie Tan , Xiang Zhao , Wenling Wang , Xuejun Ma , Yongzhong Jiang , Roujian Lu; Wenjie Tan , Xuejun Ma , Xiang Zhao , Wenling Wang , Yongzhong Jiang , Roujian Lu , Ji Wang , Peihua Niu; Xiaozhou He , Peipei Liu; Yang Li , Jing Chen , Wenbo Xu , George F. Gao , Guizhen Wu |
| EPI_ISL_434534 | National Institute for Viral Disease Control and Prevention, China CDC | National Institute for Viral Disease Control and Prevention, China CDC, Yunnan Provincial CDC | Baoying Huang; Fei Ye; Guizhen Wu; Huijuan Wang; Li Zhao; Peihua Niu; Roujian Lu; Wenjie Tan; Wenling Wang |
| EPI_ISL_402132, EPI_ISL_412898, EPI_ISL_412899, EPI_ISL_412900 | Wuhan Jinyintan Hospital | Hubei Provincial Center for Disease Control and Prevention | Bin Fang; Bo Yang; Bo Yu; Faxian Zhan; Guojun Ye; Jing Li; Junqiang Xu; Kun Cai; Linlin Liu; Xiang Li; Xiao Yu; Xixiang Huo; Yongzhong Jiang. |
| EPI_ISL_402124, EPI_ISL_402127, EPI_ISL_402128, EPI_ISL_402129, EPI_ISL_402130 | Wuhan Jinyintan Hospital | Wuhan Institute of Virology, Chinese Academy of Sciences | Ding-Yu Zhang; Hao-Rui Si; Lei Zhang; Peng Zhou; Xing-Lou Yang; Yan Zhu; Zhengli Shi |
