## Supplementary Table 3 for "Secreted SARS-CoV-2 ORF8 modulates the cytokine expression profile of human macrophages"

We gratefully acknowledge the following Authors from the Originating laboratories responsible for obtaining the specimens, as well as the Submitting laboratories where the genome data were generated and shared via GISAID, on which this research is based.

All Submitters of data may be contacted directly via www.gisaid.org

Authors are sorted alphabetically.

| Accession ID | Originating Laboratory | Submitting Laboratory | Authors |
|---|---|---|---|
| EPI_ISL_455042 | ACT Pathology | NSW Health Pathology - Institute of Clinical Pathology and Medical Research; Westmead Hospital; University of Sydney | CIDM-PH et al. |
| EPI_ISL_498469, EPI_ISL_498473, EPI_ISL_498475, EPI_ISL_498476, EPI_ISL_498480, EPI_ISL_498483, EPI_ISL_498486, EPI_ISL_498487, EPI_ISL_498490, EPI_ISL_498496, EPI_ISL_498497, EPI_ISL_498499, EPI_ISL_498500, EPI_ISL_498502, EPI_ISL_498505, EPI_ISL_498506, EPI_ISL_498507, EPI_ISL_498509, EPI_ISL_498511, EPI_ISL_498512, EPI_ISL_498517, EPI_ISL_498525, EPI_ISL_498540 | | | |
| see above | ACT Pathology | Schwessinger Lab | Ashley Jones; Benjamin Schwessinger; Craig Kennedy; Karina Kennedy; Kevin Murray; Megan McDonald; Ming-Dao Chia; Robert Lanfear; Robyn N Hall |
| EPI_ISL_451590, EPI_ISL_451592, EPI_ISL_451593 | ACT pathology | NSW Health Pathology - Institute of Clinical Pathology and Medical Research; Westmead Hospital; University of Sydney | CIDM-PH et al. |
| EPI_ISL_424848, EPI_ISL_424849 | AZ SPHL, Arizona Department of Health Services | Pathogen Discovery, Respiratory Viruses Branch, Division of Viral Diseases, Centers for Disease Control and Prevention | Alison S. Laufer Halpin; Anna Uehara; Christopher A. Elkins; Clinton R. Paden; Haibin Wang; Jing Zhang; Krista Queen; Mary S. Keckler; Rachel Marine; Suxiang Tong; Yan Li; Ying Tao |
| EPI_ISL_426483, EPI_ISL_426512, EPI_ISL_426513, EPI_ISL_426517, EPI_ISL_426537, EPI_ISL_426540 | AZ SPHL, Arizona Department of Health Services | TGen North | Darrin Lemmer; Dave Engelthaler; Jolene Bowers; Megan Folkerts |
| EPI_ISL_805714, EPI_ISL_805897, EPI_ISL_806157, EPI_ISL_806168, EPI_ISL_806248, EPI_ISL_806288, EPI_ISL_806416 | | | |
| see above | Alberta Precision Labs (APL) | Alberta Precision Labs (APL) | Berenger B; Bernier F; Chui L; Croxen M; Gordon P; Kellner J; Lam LG; Li V; Ma R; Melin A; Pabbaraju K; Tipples G; Wong A; Zelyas N |
| EPI_ISL_514647 | Allina Health Laboratory | Minnesota Department of Health, Public Health Laboratory | Jacob Garfin; Matt Plumb; and Xiong Wang |
| EPI_ISL_456656 | American Type Culture Collection Inc. (ATCC) | American Type Culture Collection Inc. (ATCC) | A.M.; Benton, B.; Chu; D.K.W.; Flores, B.; Frank; J.S.M.; King; M. and Rashid, S.; M.A.; N.P.; Parker; Peiris; Puthuveetil; Riojas; S.P. |
| EPI_ISL_500601, EPI_ISL_500602, EPI_ISL_500603, EPI_ISL_500605, EPI_ISL_500606, EPI_ISL_500613, EPI_ISL_500615, EPI_ISL_500616, EPI_ISL_500620, EPI_ISL_500626, EPI_ISL_500629, EPI_ISL_500635, EPI_ISL_500638, EPI_ISL_500641, EPI_ISL_500643, EPI_ISL_500648, EPI_ISL_500649, EPI_ISL_500650, EPI_ISL_500651, EPI_ISL_500653, EPI_ISL_500654, EPI_ISL_500655, EPI_ISL_500656, EPI_ISL_500662, EPI_ISL_500663, EPI_ISL_500664, EPI_ISL_500667, EPI_ISL_500668, EPI_ISL_500669, EPI_ISL_500671, EPI_ISL_500672, EPI_ISL_500673, EPI_ISL_500675, EPI_ISL_500676, EPI_ISL_500685, EPI_ISL_500686, EPI_ISL_500689, EPI_ISL_500694, EPI_ISL_500697, EPI_ISL_500698, EPI_ISL_500699, EPI_ISL_500705, EPI_ISL_509494, EPI_ISL_509495, EPI_ISL_509496, EPI_ISL_509497, EPI_ISL_509501, EPI_ISL_509502, EPI_ISL_509504, EPI_ISL_509506, EPI_ISL_509509, EPI_ISL_509511, EPI_ISL_509515, EPI_ISL_509516, EPI_ISL_509517, EPI_ISL_509518, EPI_ISL_509520, EPI_ISL_527035 | | | |
| see above | Area of Virology, Serology and Virology Division (SAViD), New South Wales Health Pathology Randwick | Area of Virology, Serology and Virology Division (SAViD), New South Wales Health Pathology Randwick | Rawlinson, W. |
| EPI_ISL_678293, EPI_ISL_678304, EPI_ISL_678333, EPI_ISL_678338, EPI_ISL_678382, EPI_ISL_707892, EPI_ISL_2404979, EPI_ISL_2534567, EPI_ISL_2534568, EPI_ISL_2534580, EPI_ISL_2534581 | | | |
| see above | Area of Virology, Serology and Virology Division (SAViD), New South Wales Health Pathology Randwick | Virology Research Laboratory; Area of Virology, Serology and Virology Division (SAViD), New South Wales Health Pathology Randwick | Au, J.; Bull, R.; Deveson, I.; Foster, C.; Rawlinson, W.; Ruiz Silva, M.; Van Hal, S. |
| EPI_ISL_406223 | Arizona Department of Health Services | Pathogen Discovery, Respiratory Viruses Branch, Division of Viral Diseases, Centers for Disease Control and Prevention | Anna Uehara; Brett L. Whitaker; Brian Lynch; Clinton R. Paden; Janna' R. Murray; Jing Zhang; Krista Queen; Lijuan Wang; Senthil Kumar K. Sakthivel; Shifaq Kamili; Stephen Lindstrom; Susan I. Gerber; Suxiang Tong; Xiaoyan Lu; Yan Li; Ying Tao |
| EPI_ISL_2135982, EPI_ISL_2135983, EPI_ISL_2135984, EPI_ISL_2227495 | Arizona State Public Health Laboratory | Arizona State Public Health Laboratory | Jessica Escobar; Katherine Fullerton; Linda Getsinger; Nobuko Fukushima; Stacy White; Trung Huynh; Victor Waddell |
| EPI_ISL_424668 | Arizona State University Health Services | Arizona State University | Arvind Varsani; Bereket Estifanos; Brenda G. Hogue; Efrem S. Lim; Emily A. Kaelin; Jason Steel; LaRinda A. Holland; Lily I. Wu; Matthew Scotch; Nicholas J. Mellor; Rabia Maqsood; Rolf U. Halden |
| EPI_ISL_415584, EPI_ISL_415586, EPI_ISL_415588, EPI_ISL_415589, EPI_ISL_418816, EPI_ISL_418817, EPI_ISL_418818, EPI_ISL_418819, EPI_ISL_418820, EPI_ISL_418821, EPI_ISL_418822, EPI_ISL_418823, EPI_ISL_418825, EPI_ISL_418826, EPI_ISL_418829, EPI_ISL_418830, EPI_ISL_418840, EPI_ISL_418841, EPI_ISL_418843, EPI_ISL_418847, EPI_ISL_418848, EPI_ISL_418850, EPI_ISL_418851, EPI_ISL_418852, EPI_ISL_418853, EPI_ISL_418854, EPI_ISL_418855 | | | |
| see above | BCCDC Public Health Laboratory | BCCDC Public Health Laboratory | Choi; Gilmour; Harrigan; Hoang; Kamelian; Krajden; Lapointe; Lee; Levett; Li; Loman; Prystajecky; Quick; Sekirov; Snutch; Tyson |
| EPI_ISL_445082, EPI_ISL_445084, EPI_ISL_501167 | Baylor College of Medicine | Baylor College of Medicine: HGSC | David Henke; Donna Muzny; Erin Nicholson; George Weissenberger; Ginger Metcalf; Harsha Doddapaneni; Hsu Chao; Hua Shen; Joseph F. Petrosino; Kavya Kottapalli; Kristi L. Hoffman; Matthew C. Ross; Matthew Wong; Pedro Piedra; Qingchang Meng; Richard Sucgang; Sara J.J. Cregeen; Sejal Salvi; Tulin Ayvaz; Vasanthi Avadhanula; Vipin Menon; Yimiti Meiheerguli; Zeineen Momin |
| EPI_ISL_424355, EPI_ISL_424356, EPI_ISL_424357, EPI_ISL_424358, EPI_ISL_424360, EPI_ISL_529213 | Beijing Institute of Microbiology and Epidemiology | Beijing Institute of Microbiology and Epidemiology | B. and Cui, Y.; Cui, Y.; Fan; Fan, H.; Guo, Y.; Hang; Hou, J.; Li; Li, B.; Mi, Z.; Mu, J.; Qin, E.; Song; Song, Y.; Teng; Teng, Y.; Wu, Y.; Xu, Z.; Yajun.; Yang, R.; Yong, Y.; Yue; Zhang, X. |
| EPI_ISL_429998 | Biolab Diagnostic Laboratories | Andersen lab at Scripps Research | Ahmad Tibi; Amid Abdelnour with SEARCH Alliance San Diego; Issa Abu-Dayyeh; Lama Hussein; Lina Mohammad; Zein Naber |
| EPI_ISL_414521 | Bundeswehr Institute of Microbiology | Bundeswehr Institute of Microbiology | Markus H Antwerpen and Roman Wölfel; Mathias C Walter |
| EPI_ISL_424982, EPI_ISL_424990 | Bureau of Laboratories, Michigan Department of Health and Human Services | Michigan Department of Health and Human Services | Blankenship HM; Riner D; Soehnlen MK |
| EPI_ISL_445266, EPI_ISL_445267 | CENTRO ONCOLOGICO DEL NORTE | Instituto de Salud Publica de Chile | Alejandra Acevedo; Andrés E Castillo; Bárbara Parra; Carolina Tambley; Gabriel Leal; Jaime Lagos; Jorge Fernandez; Loredana Arata; Patricia Bustos; Paz Tapia; Rodrigo Fasce; Winston Andrade |
| EPI_ISL_453816, EPI_ISL_511723 | CH Leiria | Instituto Nacional de Saude (INSA) | Borges et al |
| EPI_ISL_454011, EPI_ISL_454012, EPI_ISL_454013, EPI_ISL_511476 | CH Porto - H Sto Antonio | Instituto Nacional de Saude (INSA) | Borges et al |
| EPI_ISL_454225, EPI_ISL_511521, EPI_ISL_511522 | CHLO | Instituto Nacional de Saude (INSA) | Borges et al |
| EPI_ISL_418027 | CHTMAD | Instituto Nacional de Saude (INSA) | Guiomar et al |
| EPI_ISL_660360 | CHU Clermont-Ferrand | CNR Virus des Infections Respiratoires - France SUD | Amélie Brebion; Antonin Bal; Audrey Mirand; Bruno Lina; Christel Regagnon; Christine Archimbaud; Cécile Henquell; Gregory Destras; Gwendolyne Burfin; Hadrien Règue; Hélène Chabrolles; Laurence Josset; Martine Chambon; Martine Valette; Maxime Bisseux; Patricia Combes; Quentin Semanas |
| EPI_ISL_644691, EPI_ISL_660674 | CHU Montpellier | CNR Virus des Infections Respiratoires - France SUD | Antonin Bal; Bruno Lina; Gregory Destras; Gwendolyne Burfin; Hadrien Règue; Laurence Josset; Martine Valette; Michel Segondy; Quentin Semanas; Vincent Foulongne |
| EPI_ISL_663216 | CHU de Limoges | CNR Virus des Infections Respiratoires - France SUD | Antonin Bal; Bruno Lina; Gregory Destras; Gwendolyne Burfin; Hadrien Règue; Laurence Josset; Martine Valette; Quentin Semanas; Sylvie Rogez |
| EPI_ISL_641527 | CHU de Nice - Hôpital Archet 9 | CNR Virus des Infections Respiratoires - France SUD | Antonin Bal; Bruno Lina; Gregory Destras; Gwendolyne Burfin; Géraldine Gonfrier; Hadrien Règue; Laurence Josset; Martine Valette; Quentin Semanas; Valérie Giordanengo |
| EPI_ISL_641543, EPI_ISL_660334, EPI_ISL_660335, EPI_ISL_660336, EPI_ISL_660337, EPI_ISL_660339 | CHU de Saint-Étienne Hôpital Nord | CNR Virus des Infections Respiratoires - France SUD | Antonin Bal; Bruno Lina; Bruno Pozzetto; Gregory Destras; Gwendolyne Burfin; Hadrien Règue; Issam Bechri; Laurence Josset; Manon Vogrig; Marine Delorme; Martine Valette; Quentin Semanas; Sylvie Gonzalo; Sylvie Pillet; Thomas Bourlet |
| EPI_ISL_511058, EPI_ISL_511061, EPI_ISL_511063 | CHULC - H Curry Cabral | Instituto Nacional de Saude (INSA) | Borges et al |
| EPI_ISL_511727 | CHULN | Instituto Nacional de Saude (INSA) | Borges et al |
| EPI_ISL_535729, EPI_ISL_535814, EPI_ISL_535827 | CHUM - Microbiologie - Hôpital Saint-Luc | Laboratoire de santé publique du Québec | Guillaume Bourque; Ioannis Ragoussis; Jesse Shapiro; Mark Lathrop and Michel Roger; Mark Lathrop and Michel Roger on behalf of the CoVSeQ research group; Sandrine Moreira |
| EPI_ISL_445245, | CLINICA ALEMANA DE SANTIAGO S.A. | Instituto de Salud Publica de Chile | Alejandra Acevedo; Andrés E Castillo; Bárbara Parra; Carolina Tambley; Gabriel Leal; Jaime Lagos; Jorge Fernandez; Loredana Arata; Patricia Bustos; Paz Tapia; Rodrigo Fasce; Winston Andrade |

| | | | |
|---|---|---|---|
| EPI_ISL_445258, EPI_ISL_445260 | | | |
| EPI_ISL_445259 | CLINICA LAS CONDES S.A. | Instituto de Salud Publica de Chile | Alejandra Acevedo; Andrés E Castillo; Bárbara Parra; Carolina Tambley; Gabriel Leal; Jaime Lagos; Jorge Fernandez; Loredana Arata; Patricia Bustos; Paz Tapia; Rodrigo Fasce; Winston Andrade |
| EPI_ISL_445257 | CLINICA TABANCURA | Instituto de Salud Publica de Chile | Alejandra Acevedo; Andrés E Castillo; Bárbara Parra; Carolina Tambley; Gabriel Leal; Jaime Lagos; Jorge Fernandez; Loredana Arata; Patricia Bustos; Paz Tapia; Rodrigo Fasce; Winston Andrade |
| EPI_ISL_445306 | CLINICA UC SAN CARLOS DE APOQUINDO | Instituto de Salud Publica de Chile | Alejandra Acevedo; Andrés E Castillo; Bárbara Parra; Carolina Tambley; Gabriel Leal; Jaime Lagos; Jorge Fernandez; Loredana Arata; Patricia Bustos; Paz Tapia; Rodrigo Fasce; Winston Andrade |
| EPI_ISL_416994, EPI_ISL_418248, EPI_ISL_418249 | COMPLEJO ASISTENCIAL UNIVERSITARIO DE BURGOS | Instituto de Salud Carlos III | A. Monzón; F. Casas; G. Hospital: ------; I. Jiménez; I. Megias-Lobon G.; Iglesias-Caballero; M. Camarero; M. Camarero S. Pozo F. Casas I. Jiménez P. Jiménez M. Zaballos A. Monzón; M. Cuesta; M. González-Esguevillas; M. Molinero Calamita; M. Zaballos; P. Jiménez; S. Juliá; S. Juliá M. Cuesta I. Megias Lobón; S. Pozo; S. Varona |
| EPI_ISL_426416 | CT-Dr. Katherine A. Kelley State Public Health Lab | Pathogen Discovery, Respiratory Viruses Branch, Division of Viral Diseases, Centers for Disease Control and Prevention | Alison S. Laufer Halpin; Anna Uehara; Christopher A. Elkins; Clinton R. Paden; Haibin Wang; Jing Zhang; Krista Queen; Mary S. Keckler; Rachel Marine; Suxiang Tong; Yan Li; Ying Tao |
| EPI_ISL_418812, EPI_ISL_429819 | Cadham Provincial Laboratory | National Microbiology Laboratory | Anna Majer; David Alexander; Elsie Grudeski; Gary Van Domselaar; Grace Seo; Jared Bullard; Jennifer Tanner; Kerry Dust; Kristyn Burak; Matthew Gilmour; Morag Graham; Natalie Knox; Nathalie Bastien; Paul Van Caeseele; Philip Mabon; Rhiannon Huzarewich; Russell Mandes; Shari Tyson; Timothy Booth; Yan Li |
| EPI_ISL_582242, EPI_ISL_582244, EPI_ISL_582255, EPI_ISL_582260, EPI_ISL_582261, EPI_ISL_582275, EPI_ISL_582288, EPI_ISL_582292, EPI_ISL_582303, EPI_ISL_582304, EPI_ISL_582306, EPI_ISL_582328, EPI_ISL_582335, EPI_ISL_582338, EPI_ISL_582349, EPI_ISL_582351, EPI_ISL_582354, EPI_ISL_582400 | | | |
| see above | Cadham Provincial Laboratory | National Microbiology Laboratory (NML) | Anna Majer; Anneliese Landgraff; CanCOGeN's metadata curation team; Darian Hole; David Alexander; Elsie Grudeski; Gary Van Domselaar; Grace Seo; Jared Bullard; Jennifer Tanner; Kerry Dust; Madison Chapel; Morag Graham; Natalie Knox; Nathalie Bastien; Paul Van Caeseele; Philip Mabon; Public Health Agency of Canada CanCOGeN team; Rhiannon Huzarewich; Russell Mandes; Shari Tyson; Timothy Booth; Yan Li |
| EPI_ISL_632912, EPI_ISL_632915, EPI_ISL_632919, EPI_ISL_632921, EPI_ISL_632931 | Cadham Provincial laboratory | Cadham Provincial laboratory | Anna Majer; Anneliese Landgraff; CanCOGeN's metadata curation team; Darian Hole; David Alexander; Elsie Grudeski; Gary Van Domselaar; Grace Seo; Jared Bullard; Jennifer Tanner; Kerry Dust; Madison Chapel; Morag Graham; Natalie Knox; Nathalie Bastien; Paul Van Caeseele; Philip Mabon; Public Health Agency of Canada CanCOGeN team; Rhiannon Huzarewich; Russell Mandes; Shari Tyson; Timothy Booth; Yan Li |
| EPI_ISL_979276, EPI_ISL_979281, EPI_ISL_979286, EPI_ISL_979290 | Cadham Provincial laboratory | National Microbiology Laboratory (NML) | Anna Majer; Anneliese Landgraff; CanCOGeN's metadata curation team; Darian Hole; David Alexander; Elsie Grudeski; Gary Van Domselaar; Grace Seo; Jared Bullard; Jennifer Tanner; Kerry Dust; Kirsten Biggar; Madison Chapel; Morag Graham; Natalie Knox; Nathalie Bastien; Paul Van Caeseele; Philip Mabon; Public Health Agency of Canada CanCOGeN team; Rhiannon Huzarewich; Russell Mandes; Shari Tyson; Timothy Booth; Yan Li |
| EPI_ISL_515894, EPI_ISL_515895, EPI_ISL_515898, EPI_ISL_515901, EPI_ISL_515902, EPI_ISL_515911, EPI_ISL_754984, EPI_ISL_754991, EPI_ISL_754996, EPI_ISL_755022 | | | |
| see above | California Department of Public Health | California Department of Public Health | CDPH IDLB COVIDNet |
| EPI_ISL_413930 | California Department of Public Health | Chiu Laboratory UCSF-Abbott Viral Diagnostics and Discovery Center University of California, San Francisco | Alicia Sotomayor-Gonzalez; Allan Gopez; Chao-Yang Pan; Debra A. Wadford; Guixia Yu; Hugo Guevara; Scot Federman; Steve Miller; Wei Gu; Xianding Deng; and Charles Y. Chiu |
| EPI_ISL_413557, EPI_ISL_413922, EPI_ISL_413924, EPI_ISL_413925, EPI_ISL_413926, EPI_ISL_413928, EPI_ISL_413931, EPI_ISL_417323, EPI_ISL_417324, EPI_ISL_417326, EPI_ISL_417327, EPI_ISL_429875, EPI_ISL_429876 | | | |
| see above | California Department of Public Health | Chiu Laboratory, University of California, San Francisco | Chao-Yang Pan; Debra A. Wadford; Hugo Guevara; Scot Federman; Wei Gu; Xianding Deng; and Charles Y. Chiu |
| EPI_ISL_406034, EPI_ISL_411954 | California Department of Public Health | Pathogen Discovery, Respiratory Viruses Branch, Division of Viral Diseases, Centers for Dieases Control and Prevention | Anna Uehara; Brett L. Whitaker; Brian Lynch; Clinton R. Paden; Haibin Wang; Janna' R. Murray; Jing Zhang; Krista Queen; Lijuan Wang; Senthil Kumar K. Sakthivel; Shifaq Kamili; Stephen Lindstrom; Susan I. Gerber; Suxiang Tong; Xiaoyan Lu; Yan Li; Ying Tao |
| EPI_ISL_418865 | California Department of Public Health | University of California, San Francisco | Chao-Yang Pan; Debra A. Wadford; Hugo Guevara; Scot Federman; Wei Gu; Xianding Deng; and Charles Y. Chiu |
| EPI_ISL_579061, EPI_ISL_579067, EPI_ISL_579068, EPI_ISL_579069, EPI_ISL_579077, EPI_ISL_579439, EPI_ISL_579440, EPI_ISL_579451, EPI_ISL_579458, EPI_ISL_579476, EPI_ISL_622788, EPI_ISL_622790, EPI_ISL_622794, EPI_ISL_622797, EPI_ISL_622798 | | | |
| see above | Canterbury Health Laboratories | Institute of Environmental Science and Research (ESR) | Anja Werno; Antje van der Linden; Arlo Upton; Chris Mansell; David Hammer; Dragana Drinkovic; Erasmus Smit; Gary McAuliffe; Hana Sofia Andersson; Hermes Perez; James Ussher; Jill Sherwood; Jing Wang; Joep de Ligt; Josh Freeman; Julia Howard; Juliet Elvy; Lauren Jelly; Mary DeAlmeida; Matt Blakiston; Matt Storey; Matthew Rogers; Max Bloomfield; Michael Addidle; Michelle Balm; Muhammad Faisal; Nikki Freed; Olin Silander; Sally Roberts; Sarah Jefferies; Sharmini Muttaiyah; Susan Morpeth; Susan Taylor; Timothy Blackmore; Vani Sathyendran; Veronica Playle; Virginia Hope; Xiaoyun Ren |
| EPI_ISL_475586, EPI_ISL_475607, EPI_ISL_475612, EPI_ISL_475613, EPI_ISL_475630, EPI_ISL_475708 | Cedars-Sinai Medical Center, Department of Pathology & Laboratory Medicine, Molecular Pathology Laboratory | Cedars-Sinai Medical Center, Molecular Pathology Laboratory of Department of Pathology & Laboratory Medicine and Genomic Core | Brian Davis; Eric Vail; Jasmine T Plummer; Jean Lopategui; Jianbo Song; John Paul Govindavari; Jong Taek Kim; Stephanie Chen; Wenjuan Zhang |
| EPI_ISL_459858, EPI_ISL_459863 | Center for Genome Regulation (CRG) | Center for Mathematical Modeling and Center for Genome Regulation. Santiago, Chile | Allende ML; Gaete A; González M.; Maass A; Palma R; Travisany D; Urra C; Varas M |
| EPI_ISL_2835622, EPI_ISL_2835623 | Center for Medical and Sanitary Research (CERMES) | CDC Atlanta | Adamou Lagare; Dhwani Batra; Justin Lee |
| EPI_ISL_493137 | Center for Research and Innovation, Faculty of Medical Technology, Mahidol University | Center for Research and Innovation, Faculty of Medical Technology, Mahidol University | Anek Mungaomklang; Hatairat Lerdsamran; Jarunee Prasertsopon; Kamolthip Atsawawaranunt; Kantima Sangsiriwut; Nattakan Thinpan; Pilaipan Puthavathana; Prabda Praphasiri; Somrak Sirikhetkon; Tipsuda Chanmanee |
| EPI_ISL_438048, EPI_ISL_438065, EPI_ISL_438070, EPI_ISL_438092, EPI_ISL_438106, EPI_ISL_475775, EPI_ISL_475787, EPI_ISL_475789, EPI_ISL_475790, EPI_ISL_583566 | | | |
| see above | Center for Virology, Medical University of Vienna | Bergthaler laboratory, CeMM Research Center for Molecular Medicine of the Austrian Academy of Sciences | Adi Steinrigl; Alexander Lercher; Alexandra Popa; Andreas Bergthaler; Benedikt Agerer; Christian Paar; Christoph Bock; Daniela Schmid; Dorothee von Laer; Elisabeth Puchhammer-Stoeckl; Franz Allerberger; Gernot Walder; Gregor Hörmann; Guenter Weiss; Gunther Vogl; Henrique Colaco; Jakob-Wendelin Genger; Jan Laine; Judith Aberle; Kinga Rigler-Hohenwarter; Lukas Endler; Manfred Nairz; Mark Smyth; Martin Senekowitsch; Michael Schuster; Peter Hufnagl; Peter Obrist; Rainer Gattringer; Sabine Sussitz-Rack; Stephan Aberle; Thomas Penz; Wegene Borena |
| EPI_ISL_425123 | Center of Medical Microbiology, Virology, and Hospital Hygiene, University of Duesseldorf | Center of Medical Microbiology, Virology, and Hospital Hygiene, University of Duesseldorf | Alexander Dilthey; Andreas Walker; Björn-Erik Jensen; Daniel Strelow; Detlef Kindgen-Milles; Jörg Timm; Klaus Pfeffer; Malte Kohns Vasconcelos; Marcel Andree; Ortwin Adams; Sandra Hauka; Tina Senff; Tobias Wienemann; Torsten Feldt; Torsten Houwaart |
| EPI_ISL_429854, EPI_ISL_429855 | Centers for Disease Control and Prevention of Lishui | Department of Inspection , Centers for Disease Control and Prevention of Lishui | Ji Jiansong; Ji Qiaoying; Wang Xiaoguang; Ye Bifeng; Ye Ling |
| EPI_ISL_978504 | Central Public Health Laboratory - LACEN -Bahia, Salvador, Brazil | Central Public Health Laboratory - LACEN -Bahia, Salvador, Brazil | Arabela Leal; Breno Dominguez; Felicidade Pereira; Jaqueline Gomes; Luciana Oliveira; Luiz Alcantara; Marcela Gómez; Marta Giovanetti; Patrícia Cajado; Stephane Tosta; Vagner Fonseca; Vanessa Nardy |
| EPI_ISL_535843, EPI_ISL_536059, EPI_ISL_536060 | Centre Hospitalier Régional de Lanaudière | Laboratoire de santé publique du Québec | Guillaume Bourque; Ioannis Ragoussis; Jesse Shapiro; Mark Lathrop and Michel Roger; Mark Lathrop and Michel Roger on behalf of the CoVSeQ research group; Sandrine Moreira |
| EPI_ISL_539573, EPI_ISL_539575 | Centre de Recherches Medicales de Lambarene (CERMEL) | Department of Emerging Infectious Diseases, Institute of Tropical Medicine, Nagasaki University | Akim A. Adegnika; Bertrand Lell; Haruka Abe; Jiro Yasuda; Rodrigue Bikangui; Yuri Ushijima |
| EPI_ISL_483926, EPI_ISL_483988, EPI_ISL_484151 | Centre for Clinical Infection and Diagnostics Research and Genomics Innovation Unit, Guy's and St. Thomas' NHS Trust | COVID-19 Genomics UK (COG-UK) Consortium | Ali Raza Awan; Chloe Fisher; Jonathan Edgeworth; Luke Snell; Penny Cliff; Rahul Batra |
| EPI_ISL_407893, EPI_ISL_417030 | Centre for Infectious Diseases and Microbiology Laboratory Services | NSW Health Pathology - Institute of Clinical Pathology and Medical Research; Westmead Hospital; University of Sydney | Carter I; Chen SC; Eden J-S; Holmes EC; Kok J and Dwyer DE for the 2019-nCoV Study Group; Kok J and Dwyer DE for the 2019-nCoV Study Group*; Maddocks S; O’Sullivan MV; Rahman H; Rockett R; Sintchenko V |
| EPI_ISL_417395, EPI_ISL_427667, EPI_ISL_427670, EPI_ISL_427671, EPI_ISL_427672, EPI_ISL_427675, EPI_ISL_427678, EPI_ISL_427681, EPI_ISL_427684, EPI_ISL_427685, EPI_ISL_427691, EPI_ISL_427694, EPI_ISL_427695, EPI_ISL_427699, EPI_ISL_427701, EPI_ISL_427707, EPI_ISL_427709, EPI_ISL_427710, EPI_ISL_427713, EPI_ISL_427714, EPI_ISL_427716, EPI_ISL_427722, EPI_ISL_427726, EPI_ISL_427727, EPI_ISL_427730, EPI_ISL_427732, EPI_ISL_427736, EPI_ISL_427740, EPI_ISL_427745, EPI_ISL_427746, EPI_ISL_427747, EPI_ISL_427752, EPI_ISL_427753, EPI_ISL_427755, EPI_ISL_427762, EPI_ISL_427763, EPI_ISL_427765, EPI_ISL_427766, EPI_ISL_427767, EPI_ISL_427769, EPI_ISL_427774, EPI_ISL_427777, EPI_ISL_427786, EPI_ISL_427792, EPI_ISL_427793, EPI_ISL_427797, EPI_ISL_427798, EPI_ISL_427803, EPI_ISL_427806, EPI_ISL_427807 | | | |
| see above | Centre for Infectious Diseases and Microbiology Public Health | NSW Health Pathology - Institute of Clinical Pathology and Medical Research; Westmead Hospital; University of Sydney | Arnott A; Bachmann N; Basile K; Byun R; Carter I; Chang S; Chen SC; Draper J; Dwyer DE; Dwyer DE for the 2019-nCoV Study Group; Eden J-S; Eden JS; Gall M; Gray K; Holmes EC; Holmes EC and O’Sullivan MV for the 2019-nCoV Study Group; Kok J; Lam C; Maddocks S; O’Sullivan MV; Propenko M; Rahman H; Rockett R; Sadsad R; Sim E; Sintchenko V; Sorrell T; Timms; Timms V; V |
| EPI_ISL_536043 | Centre hospitalier Anna-Laberge | Laboratoire de santé publique du Québec | Guillaume Bourque; Ioannis Ragoussis; Jesse Shapiro; Mark Lathrop and Michel Roger on behalf of the CoVSeQ research group; Sandrine Moreira |
| EPI_ISL_837576 | Centro Nacional de Enfermedades Tropicales (CENETROP) | Laboratory of Respiratory Viruses and Measles, Oswaldo Cruz Institute, FIOCRUZ | Ana Carolina Mendonca; Anna Carolina Paixao; Cinthia Avila; Fernando Motta; Luciana Appolinario; Marilda Siqueira on behalf of the Fiocruz COVID-19 Genomic Surveillance Network; Paola Resende; Roxana Loayza |
| EPI_ISL_468762, EPI_ISL_468763, EPI_ISL_468764, EPI_ISL_500370, EPI_ISL_500377, EPI_ISL_500378, EPI_ISL_500387, EPI_ISL_500388, EPI_ISL_500390, EPI_ISL_500391, EPI_ISL_500392, EPI_ISL_500393, EPI_ISL_500394, EPI_ISL_500395, EPI_ISL_500396, EPI_ISL_500397, EPI_ISL_500398, EPI_ISL_500399, EPI_ISL_500400, EPI_ISL_500401, EPI_ISL_500402, EPI_ISL_500403, EPI_ISL_500404, EPI_ISL_500405, EPI_ISL_500406, EPI_ISL_500408, EPI_ISL_500414, EPI_ISL_500419, EPI_ISL_500420, EPI_ISL_500423, EPI_ISL_500424, EPI_ISL_500426, EPI_ISL_500429, EPI_ISL_500434, EPI_ISL_500435, EPI_ISL_500436, EPI_ISL_500437, EPI_ISL_500438, EPI_ISL_500439, EPI_ISL_500441, EPI_ISL_500442, EPI_ISL_500444, EPI_ISL_500446, EPI_ISL_500447, EPI_ISL_500448, EPI_ISL_500457, EPI_ISL_500458, EPI_ISL_537383, EPI_ISL_537386, EPI_ISL_537387, EPI_ISL_537388, EPI_ISL_537389, EPI_ISL_537391, EPI_ISL_537392, EPI_ISL_537400, EPI_ISL_537401, EPI_ISL_537402, EPI_ISL_537403, EPI_ISL_537404, EPI_ISL_537405, EPI_ISL_537406, EPI_ISL_537407, EPI_ISL_537408, EPI_ISL_537409, EPI_ISL_537410, EPI_ISL_537411, EPI_ISL_537412, EPI_ISL_537413, EPI_ISL_537415, EPI_ISL_537416, EPI_ISL_537417, EPI_ISL_537418, EPI_ISL_537419, EPI_ISL_537420, EPI_ISL_537421, EPI_ISL_537423, EPI_ISL_537424, EPI_ISL_537425, EPI_ISL_537426, EPI_ISL_537430, EPI_ISL_537438, EPI_ISL_537448, EPI_ISL_537449, EPI_ISL_537454, EPI_ISL_537462, EPI_ISL_537811, EPI_ISL_537812, EPI_ISL_537813, EPI_ISL_537814, EPI_ISL_537815, EPI_ISL_537816, EPI_ISL_537817, EPI_ISL_537818, EPI_ISL_537819, EPI_ISL_537820, EPI_ISL_537821, EPI_ISL_537822, EPI_ISL_537823, EPI_ISL_537824, EPI_ISL_537825, EPI_ISL_537826, EPI_ISL_537827, EPI_ISL_537830, EPI_ISL_537831, EPI_ISL_537832, EPI_ISL_537833, EPI_ISL_537835, EPI_ISL_537838, EPI_ISL_537839, EPI_ISL_537843, EPI_ISL_537848, EPI_ISL_537854, EPI_ISL_537860, EPI_ISL_537861, EPI_ISL_537862, EPI_ISL_537867, EPI_ISL_537868, EPI_ISL_537872, EPI_ISL_732761, EPI_ISL_732762 | | | |
| see above | Centro de Investigación Biomédica de La Rioja - Hospital San Pedro Logroño | SeqCOVID-SPAIN consortium/IBV(CSIC) | José Manuel Azcona Gutiérrez; María Pilar Bea Escudero; María de Toro; Miriam Blasco Alberdi and SeqCOVID-SPAIN consortium |
| EPI_ISL_753843 | Charité Universitätsmedizin Berlin, Institut für Virologie/Labor Berlin | Charité Universitätsmedizin Berlin, Institut für Virologie | Barbara Mühlemann; Christian Drosten; Julia Schneider; Jörn Beheim-Schwarzbach; Talitha Veith; Terry Jones; Victor M Corman |
| EPI_ISL_430722, EPI_ISL_430723, EPI_ISL_430724, EPI_ISL_430725, EPI_ISL_430728, EPI_ISL_430730, EPI_ISL_430731, EPI_ISL_430733, EPI_ISL_430737, EPI_ISL_430738, EPI_ISL_430741 | | | |
| see above | Chinese PLA Institute for Disease Control and Prevention | Chinese PLA Institute for Disease Control and Prevention | Lizhong Li; Peng Li , Jinhui Li |
| EPI_ISL_417332 | Chiu Laboratory, University of California, San Francisco | Chiu Laboratory, University of California, San Francisco | Scot Federman; Wei Gu; Xianding Deng; and Charles Y. Chiu |
| EPI_ISL_414579 | Clinica Alemana de Santiago, Chile | Instituto de Salud Publica de Chile | Alejandra Acevedo; Andrés E. Castillo; Bárbara Parra; Carolina Tambley; Gabriel Leal; Gisselle Barra; Jaime Lagos; Javier Tognarelli; Jorge Fernández.; Loredana Arata; Patricia Bustos; Paz Tapia; Rodrigo Fasce; Soledad Ulloa; Winston Andrade |

| | | | |
|---|---|---|---|
| EPI_ISL_437455, EPI_ISL_902978, EPI_ISL_911403, EPI_ISL_911450, EPI_ISL_911470, EPI_ISL_911576, EPI_ISL_911607, EPI_ISL_911612, EPI_ISL_911722 | | | |
| see above | Clinical Diagnostics Laboratory, Diagnostic & Experimental Pathology, Lilly Research Laboratories | Clinical Diagnostics Laboratory, Diagnostic & Experimental Pathology, Lilly Research Laboratories | Andrew Schade; Angie Fulford; Erin Wray; Jeff Fill; Joe Oakley; John Calley; John McElwee; Leslie ONeill Reising; Mayuri Vaidya; Pat Finnegan; Phil Ebert; Rachael Redmond; Sam McNeely; Tim Holzer |
| EPI_ISL_1233898 | Clinical Division, Fred Hutchinson Cancer Research Center | Clinical Division, Fred Hutchinson Cancer Research Center | A.K.; C.-W.; D.E.; Geraghty; Greninger, A.; Ishitani, A.; Jerome, K.; L.P.; Li, R.; M.G.; Mills; Nelson; Pyo; Randhawa; Roychoudhury, P.; W.C.; Wang, R.; Zhao |
| EPI_ISL_877590, EPI_ISL_877591 | Clinical Molecular Microbiology Laboratory, UNC Hospital | Dirk Dittmer | Angelica Juarez; Blossom Damania.; Brent A. Eason; Cameroon Grant; Carolina Caro-Vegas; Cecilia Thompson; Dirk Dittmer; Evelyn Hoffman; Jason Wong; Justin T. Landis; Linda Pluta; Melissa B. Miller; Patricio Cano; Razia Moorad |
| EPI_ISL_862652, EPI_ISL_862653 | Clínica Universidad de Navarra | Instituto de Salud Carlos III | A. Monzón; F. Casas; I. Fernández, M.; I. Jiménez; Iglesias-Caballero; M. Pozo; M. Cuesta; M. González-Esguevillas; M. Zaballos; M.Camarero; P. Jiménez; S. Juliá; S. Molinero Calamita; S. Varona |
| EPI_ISL_452478, EPI_ISL_452481, EPI_ISL_452482, EPI_ISL_452489, EPI_ISL_452493, EPI_ISL_452494, EPI_ISL_452499, EPI_ISL_452500, EPI_ISL_452501, EPI_ISL_452502, EPI_ISL_452504, EPI_ISL_452508, EPI_ISL_452512, EPI_ISL_452515, EPI_ISL_452520, EPI_ISL_452522, EPI_ISL_452523, EPI_ISL_452525, EPI_ISL_452526, EPI_ISL_452529, EPI_ISL_452534, EPI_ISL_452535, EPI_ISL_452536, EPI_ISL_452537, EPI_ISL_452538, EPI_ISL_452539, EPI_ISL_452540, EPI_ISL_452541, EPI_ISL_452542, EPI_ISL_452543, EPI_ISL_538082, EPI_ISL_538083, EPI_ISL_538100 | | | |
| see above | Clínica Universidad de Navarra. Servicio de Enfermedades Infecciosas y Microbiología clínica | SeqCOVID-SPAIN consortium/IBV(CSIC) | Jose Luis del Pozo and SeqCOVID-SPAIN consortium; Mirian Fernández-Alonso |
| EPI_ISL_479662 | College of Veterinary Medicine, Chungnam National University | College of Veterinary Medicine, Chungnam National University | Seo, S. |
| EPI_ISL_677241 | Colorado Department of Public Health and Environment | Colorado Department of Public Health & Environment | Emily A. Travanty; Laura Bankers; Molly Hetherington-Rauth; Sarah Elizabeth Totten; Shannon Ely; Shannon R. Matzinger |
| EPI_ISL_2549577, EPI_ISL_2549578, EPI_ISL_2549579, EPI_ISL_2549580, EPI_ISL_2549581, EPI_ISL_2549582, EPI_ISL_2549583, EPI_ISL_2549584, EPI_ISL_2549585, EPI_ISL_2549586, EPI_ISL_2549587, EPI_ISL_2549588, EPI_ISL_2549593, EPI_ISL_2549594, EPI_ISL_2549595, EPI_ISL_2549596, EPI_ISL_2549597, EPI_ISL_2549598, EPI_ISL_2549599, EPI_ISL_2549600, EPI_ISL_2549601, EPI_ISL_2549608, EPI_ISL_2549609, EPI_ISL_2549610, EPI_ISL_2549611, EPI_ISL_2549612, EPI_ISL_2549613, EPI_ISL_2549614, EPI_ISL_2549615, EPI_ISL_2549616, EPI_ISL_2549617, EPI_ISL_2549618, EPI_ISL_2549619, EPI_ISL_2549620, EPI_ISL_2549632, EPI_ISL_2549633, EPI_ISL_2549636, EPI_ISL_2549640, EPI_ISL_2549642, EPI_ISL_2549643, EPI_ISL_2549644, EPI_ISL_2549645, EPI_ISL_2549647, EPI_ISL_2549648, EPI_ISL_2549652, EPI_ISL_2549655, EPI_ISL_2549659, EPI_ISL_2549661, EPI_ISL_2549662, EPI_ISL_2549671, EPI_ISL_2549673, EPI_ISL_2549689, EPI_ISL_2549701, EPI_ISL_2549710 | | | |
| see above | Colorado Department of Public Health and Environment | Colorado Department of Public Health and Environment | Alexandria Rossheim; Diana Ir; Emily A. Travanty; Laura Bankers; Molly C. Hetherington-Rauth; Sarah Elizabeth Totten; Shannon R. Matzinger |
| EPI_ISL_677246, EPI_ISL_677253, EPI_ISL_677269 | Colorado Department of Public Health and Environment | Colorado Department of Puplic Health and Environment | Emily A. Travanty; Laura Bankers; Molly Hetherington-Rauth; Sarah Elizabeth Totten; Shannon Ely; Shannon R. Matzinger |
| EPI_ISL_455328, EPI_ISL_455329, EPI_ISL_455330, EPI_ISL_455331, EPI_ISL_539563, EPI_ISL_539564, EPI_ISL_578194 | | | |
| see above | Complejo Hospitalario Universitario La Coruna | Instituto de Salud Carlos III | A. Monzón; F. Casas; I; I. Jiménez; Iglesias-Caballero; J. López; M. Camarero; M. Cuesta; M. González-Esguevillas; M. Molinero Calamita; M. Zaballos; M.A Canizares; M.I Paz; P. Jiménez; S. Juliá; S. Pozo; S. Varona |
| EPI_ISL_539570, EPI_ISL_539571, EPI_ISL_862645, EPI_ISL_862646, EPI_ISL_862647, EPI_ISL_862648, EPI_ISL_862649, EPI_ISL_862650, EPI_ISL_862651 | | | |
| see above | Complejo Hospitalario de Navarra | Instituto de Salud Carlos III | A. Monzón; F. Casas; I; I. Ezpeleta, C.; I. Jiménez; Iglesias-Caballero; J. López; M. Pozo; M. Camarero; M. Cuesta; M. González-Esguevillas; M. Molinero Calamita; M. Zaballos; M.Camarero; P. Jiménez; S. Juliá; S. Molinero Calamita; S. Pozo; S. Varona |
| EPI_ISL_416417, EPI_ISL_416418, EPI_ISL_416419, EPI_ISL_435712, EPI_ISL_435718 | Connecticut State Department of Public Health | Grubaugh Lab - Yale School of Public Health | Adam Moore; Akiko Iwasaki; Albert Ko; Alice Lu; Allison Nelson; Anderson Brito; Anne Wyllie; Anthony Muyombwe; Arnau Casanovas; Catherine Muenker; Chaney Kalinich; Chantal Vogels; Charlese Dela Cruz; Cole Jensen; Ellen Foxman; Isabel Ott; Jafar Razeq; Joseph Fauver; Maria Tokuyama; Marie-Louise Landry; Mary Petrone; Nagarjuna Cheemarla; Nathan Grubaugh; Patrick Wong; Peiwen Lu; Richard Martinello; Saad Omer; Shelli Farhadian; Tara Alpert |
| EPI_ISL_536222 | Conseil Cri de la SSS de la Baie-James | Laboratoire de santé publique du Québec | Guillaume Bourque; Ioannis Ragoussis; Jesse Shapiro; Mark Lathrop and Michel Roger on behalf of the CoVSeQ research group; Sandrine Moreira |
| EPI_ISL_691681 | Consejeria de Sanidad y Asuntos Sociales | Instituto de Salud Carlos III | A. Monzón; F. Casas; I. Gutiérrez, G.; I. Jiménez; Iglesias-Caballero; M. Camarero; M. Cuesta; M. González-Esguevillas; M. Pozo; M. Zaballos; P. Jiménez; S. Juliá; S. Molinero Calamita; S. Varona |
| EPI_ISL_862641, EPI_ISL_862642, EPI_ISL_862643, EPI_ISL_862644 | Consejería de Sanidad y Asuntos Sociales | Instituto de Salud Carlos III | A. Monzón; F. Casas; I. Gutiérrez, G.; I. Jiménez; Iglesias-Caballero; M. Camarero; M. Cuesta; M. González-Esguevillas; M. Pozo; M. Zaballos; P. Jiménez; S. Juliá; S. Molinero Calamita; S. Varona |
| EPI_ISL_468615 | Contra Costa Public Health Lab | Chan-Zuckerberg Biohub | CZB Cliahub Consortium |
| EPI_ISL_468400 | County of San Luis Obispo Public Health Laboratory | Chan-Zuckerberg Biohub | CZB Cliahub Consortium |
| EPI_ISL_476177, EPI_ISL_476224 | DB Diagnósticos do Brasil | Instituto de Medicina Tropical da Univesidade de São Paulo | Camila Alves Maia da Silva; Darlan da Silva Candido; Erika Regina Manuli; Ester Sabino; Flavia Cristina da Silva Sales; Giula Magalhaes Ferreira; Jaqueline Goes de Jesus; Julien Theze; Mariana Severo Ramundo; Nuno Faria; Samples: Nelson Gaburo Jr; Sequencing: Ingra Morales Claro; Thais de Moura Coletti |
| EPI_ISL_804850, EPI_ISL_804884, EPI_ISL_804908 | DC Public Health Lab/ Dept. of Forensic Sciences | DC Public Health Lab/ Dept. of Forensic Sciences | Brittany Hamilton; Connie Maza; David Payne; Elizabeth Zelaya; Jocelyn Hauser; Monica Mann; Scott Nguyen |
| EPI_ISL_424852 | DC Public Health Lab/ Dept. of Forensic Sciences | Pathogen Discovery, Respiratory Viruses Branch, Division of Viral Diseases, Centers for Disease Control and Prevention | Alison S. Laufer Halpin; Anna Uehara; Christopher A. Elkins; Clinton R. Paden; Haibin Wang; Jing Zhang; Krista Queen; Mary S. Keckler; Rachel Marine; Suxiang Tong; Yan Li; Ying Tao |
| EPI_ISL_560567, EPI_ISL_561345, EPI_ISL_561346, EPI_ISL_561348, EPI_ISL_561359, EPI_ISL_565832, EPI_ISL_574586, EPI_ISL_574599, EPI_ISL_575335, EPI_ISL_576151, EPI_ISL_576156, EPI_ISL_576158, EPI_ISL_576160, EPI_ISL_576163, EPI_ISL_576223, EPI_ISL_576224, EPI_ISL_576225, EPI_ISL_576227, EPI_ISL_584013, EPI_ISL_593945, EPI_ISL_593961, EPI_ISL_593963, EPI_ISL_593964, EPI_ISL_593966 | | | |
| see above | Delaware Public Health Lab | Delaware Public Health Lab | Gregory Hovan |
| EPI_ISL_444493 | Departamento de Laboratorios de Salud Publica (DLSP, Division Epidemiologia, Ministerio de Salud Publica) | Facultad de Ciencias (Sección Genética Evolutiva, Sección Virología). | Arbiza; Calleros, L.; Chiparelli, H.; Coppola, L.; Delfraro, A.; Frabasile, S.; Fuques, E.; Goni, N.; Grecco, S.; J. and Perez, R.; Panzera, Y.; Ramos, N.; Ramos, V.; Techera, C. |
| EPI_ISL_508637, EPI_ISL_508644, EPI_ISL_508645 | Departamento de Microbiología, CDB, Hospital Clínic, Barcelona | SeqCOVID-SPAIN consortium/IBV(CSIC) | Aida Peiró and SeqCOVID-SPAIN consortium; Andrea Vergara; Elisa Rubio; Jéssica Navero; Mikel Martínez |
| EPI_ISL_515082 | Department of Biochemistry, Cell and Molecular Biology | WACCBIP, University of Ghana | A.K.; Adu, B.; Amenga-Etego; Ampofo, W.; Amuzu; Anang; Arjarquah, A.; Asante, I.; Awandare; Bediako, Y.; Boatemaa, L.; Bonney, E.; Bonney, K.; C.M.; D.S.; Eshun, M.; G.A.; G.B.; J.K.; J.M.; Kotey, E.; Kumordjie, S.; Kyei; L.N.; Magnussen, V.; Morang'a; Mutungi; Ngoi; Quashie, P.; Tei-Maya, F. |
| EPI_ISL_481251 | Department of Emerging Infectious Diseases, Institute of Tropical Medicine, Nagasaki University | Department of Emerging Infectious Diseases, Institute of Tropical Medicine, Nagasaki University | Haruka Abe; Jiro Yasuda; Rokusuke Yoshikawa; Yuichiro Furusato |
| EPI_ISL_610166 | Department of Health Technology and Informatics, The Hong Kong Polytechnic University | Department of Health Technology and Informatics, The Hong Kong Polytechnic University | A.K.-L.; A.Y.-M.; B.K.-C.; C.T.-M.; Chan; Chau; D.H.-K.; Fung; G.K.-H.; H.-Y.; Ho; J.S.-L.; K.-T.; K.K.-G.; K.S.-C.; K.S.-S.; L.-K.; Lai; Lao; Lee; Leung; Luk, K.; M.C.-Y.; Ng; Que; S.K.-Y.; S.P.; Shum; Siu; T.-L.; T.T.-L.; Tam; To; W.-K.; W.C.; Wong; Wu; Y.W.-M.; Yam; Yau; Yip |
| EPI_ISL_2710829, EPI_ISL_2710873, EPI_ISL_2710879 | Department of Health Technology and Informatics, The Hong Kong Polytechnic University | Department of Health Technology and Informatics, The Hong Kong Polytechnic University | Alan Ka-Lun Wu; Alex Yat-Man Ho; Barry Kin-Chung Wong; Chloe Toi-Mei Chan; David Ho-Keung Shum; Denise Sze-Hang Wong; Gilman Kit-Hang Siu; Hiu-Yin Lao; Hoi-Ching Jim; Ivan Tak-Fai Wong; Jake Siu-Lun Leung; Kam-Tong Yip; Kenneth Siu-Sing Leung; Kingsley King-Gee Tam; Kitty Sau-Chun Fung; Kristine Luk; Lam-Kwong Lee; Miranda Chong-Yee Yau; Sandy Ka-Yee Chau; Shea Ping Yip; Tak-Lun Que; Timothy Ting-Leung Ng; Wing Cheong Yam; Wing-Hei Lo; Wing-Kin To; Yvette Wai-Man Lai |
| EPI_ISL_568556, EPI_ISL_568557, EPI_ISL_636989 | Department of Infectious Diseases and Immunology, National Hospital Organization Nagoya Medical Center | Clinical Research Center, National Hospital Organization Nagoya Medical Center | Hirotaka Ode; Kazuhiro Matsuoka; Mai Kubota; Masakazu Matsuda; Mayumi Imahashi; Miho Nakasuji; Mikiko Mori; Nakasuji Miho; Yasumasa Iwatani; Yoshihiro Nakata; Yoshiyuki Yokomaku |
| EPI_ISL_479821, EPI_ISL_479822, EPI_ISL_479855, EPI_ISL_479856, EPI_ISL_479857, EPI_ISL_479858, EPI_ISL_479859, EPI_ISL_479860, EPI_ISL_479868, EPI_ISL_479986 | | | |
| see above | Department of Infectious Diseases, Kobe Institute of Health | Pathogen Genomics Center, National Institute of Infectious Diseases | Hajime Kamiya; Kentaro Itokawa; Makoto Kuroda; Masanori Hashino; Motoi Suzuki; Rina Tanaka; Ryohei Nomoto; Tsuyoshi Sekizuka |
| EPI_ISL_516805 | Department of Laboratory Medicine, Tan Tock Seng Hospital | Department of Laboratory Medicine, Tan Tock Seng Hospital | Barkham TMS; Chen YYC; Li C; Maurer-Stroh S; Nagarajan N; Sessions OM; Tang WY; Zair X |
| EPI_ISL_1509058, EPI_ISL_1675177 | Department of Laboratory Medicine, Clinical Center, National Institutes of Health | Laboratory of Parasitic Diseases, Systems Genomics Section, National Institute of Allergy and Infectious Diseases, National Institutes of Health | Allison Roder; Elodie Ghedin; Heike Bailin; Jennifer Kwan; Jessica McCormick-Ell; Jung-ho Youn; Matthew Chung; Michael Bell; Rachel Mercado; Sanchita Das; Stephanie Banakis; Tara Palmore; Wei Wang |
| EPI_ISL_408489, EPI_ISL_422415, EPI_ISL_447615 | Department of Laboratory Medicine, National Taiwan University Hospital | Microbial Genomics Core Lab, National Taiwan University Centers of Genomic and Precision Medicine | Chiao-Ling Li; Pei-Jer Chen; Shan-Chwen Chang; Shiou-Hwei Yeh; Sui-Yuan Chang; Ya-Yun Lai; You-Yu Lin |
| EPI_ISL_538452, EPI_ISL_538456, EPI_ISL_538458, EPI_ISL_648064 | Department of Laboratory Medicine, Tan Tock Seng Hospital | Department of Laboratory Medicine, Tan Tock Seng Hospital | Barkham TMS; Chen YYC; Li C; Lim JX; Maurer-Stroh S; Nagarajan N; Sessions OM; Tang WY; Zair X |
| EPI_ISL_3065955, EPI_ISL_3065956, EPI_ISL_3065968, EPI_ISL_3065974, EPI_ISL_3065992, EPI_ISL_3065994 | Department of Microbiology | Department of Microbiology | Paul K.S. Chan; Zigui Chen |
| EPI_ISL_413514, EPI_ISL_413516 | Department of Microbiology, Institute for Viral Diseases, College of Medicine, Korea University | Department of Microbiology, Institute for Viral Diseases, College of Medicine, Korea University | Changmin Kang; Cui Chunguang; Gee Eun Lee; Hee Jin Cheong; Heedo Park; Jeonghun Kim; Ji Yun Noh; Jin Gu Yoon; Jin Il Kim; Joon Young Song; Joon-Yong Bae; Jungmin Lee; Juyoung Cho; Kyeong-ryeol Shin; Man-Seong Park; Woo Joo Kim |
| EPI_ISL_420538 | Department of Microbiology, PathWest QEII Medical Centre | Department of Microbiology, PathWest QEII Medical Centre | Avram Levy; Chisha Sikazwe; David Speers and David Smith; Jurissa Lang |
| EPI_ISL_497832, EPI_ISL_497833 | Department of Microbiology, The University of Hong Kong | Department of Microbiology, The University of Hong Kong | Kelvin K.W. To; Kwok-Yung Yuen |
| EPI_ISL_417176 | Department of Pathology, Princess Margaret Hospital | Department of Health Technology and Informatics, Faculty of Health | Alan Ka-Lun WU; Alex Yat-Man HO; Barry Kin-Chung WONG; David Ho-Keung SHUM; Eugene Yuk-Keung TSO; Gilman Kit-Hang SIU; Hiu-Yin LAO; Kam-Tong Yip; Kenneth Siu-Sing LEUNG; Kingsley King-Gee TAM; Kit-Man SIN; Kitty Sau-Chun |

| | | | |
|---|---|---|---|
| | | and Social Science, The Hong Kong Polytechnic University | FUNG; Kwok-Cheung LUNG; Lam-Kwong LEE; Man-Chun CHAN; Ming-Pan CHOI; Miranda Chong-Yee YAU; Raymond Wai-To LIU; Sandy Ka-Yee CHAU; Shea Ping YIP; Tak-Lun Que; Timothy Ting-Leung NG; Wai-Shing LEUNG; Wing Cheong YAM; Wing-Kin TO; Yuk-Yung NG |
| EPI_ISL_438393, EPI_ISL_438503, EPI_ISL_439486, EPI_ISL_439651, EPI_ISL_440055, EPI_ISL_440209, EPI_ISL_440404 | | | |
| see above | Department of Pathology, University of Cambridge | Wellcome Sanger Institute for the COVID-19 Genomics UK (COG-UK) consortium | Alex Alderton; Aminu S. Jahun; Anna Yakovleva; Charlotte J. Houldcroft; Cordelia Langford; David K. Jackson; Dominic Kwiatkowski; Ewan Harrison; Fahad A Khokhar; Grant Hall; Ian Goodfellow; Ian Johnston; John Sillitoe on behalf of the Wellcome Sanger Institute COVID-19 Surveillance Team (http://www.sanger.ac.uk/covid-team); Laura G Caller; Luke W Meredith; M. Estée Török; Martin D. Curran; Myra Hosmillo; Roberto Amato; Sarah L. Caddy; Sonia Goncalves; Theresa Feltwell; William L. Hamilton |
| EPI_ISL_1240098 | Department of Virology and Immunology, University of Helsinki and Helsinki University Hospital, Huslab Finland | Department of Virology, Faculty of Medicine, University of Helsinki, Helsinki, Finland | Essi Korhonen; Fathiah Zakham; Hanna Jarva; Hanna Liimatainen; Hannimari Kallio-Kokko; Harri Kangas; Hussein Alburkat; Jenni Virtanen; Maija Lappalainen; Maija Suvanto; Mert Erdin; Olli Vapalahti; Pekka Ellonen; Phuoc Truong; Ravi Kant; Sari Hannula; Satu Kurkela; Teemu Smura |
| EPI_ISL_444925, EPI_ISL_444932 | Department of Virus and Microbiological Special Diagnostics, Statens Serum Institut, Copenhagen, Denmark, Artillerivej 5, 2300 Copenahgen S | Albertsen lab, Department of Chemistry and Bioscience, Aalborg University, Denmark | Rasmus Kirkegaard |
| EPI_ISL_614476, EPI_ISL_614477 | Department of Virus and Microbiological Special Diagnostics, Statens Serum Institut, Denmark | Albertsen lab, Department of Chemistry and Bioscience, Aalborg University, Denmark | Danish Covid-19 Genome Consortia |
| EPI_ISL_413485 | Department of microbiology laboratory,Anhui Provincial Center for Disease Control and Prevention | Department of microbiology laboratory,Anhui Provincial Center for Disease Control and Prevention | Bin Su; Jun He; Junling Yu; Qingqing Chen; Weidong Li; Weiwei Li; Yinglu Ge; Yong Sun; Yonglin Shi; Yuan Yuan; Zhirong Liu; Zhuhui Zhang |
| EPI_ISL_408665, EPI_ISL_408666, EPI_ISL_408667 | Dept. of Virology III, National Institute of Infectious Diseases | Pathogen Genomics Center, National Institute of Infectious Diseases | Kazuya Shirato; Makoto Kuroda; Makoto Takeda; Naganori Nao; Shutoku Matsuyama; Tsuyoshi Sekizuka |
| EPI_ISL_413513, EPI_ISL_413515 | Division of Infectious Diseases, Department of Internal Medicine, Korea University College of Medicine | Department of Microbiology, Institute for Viral Diseases, College of Medicine, Korea University | Changmin Kang; Cui Chunguang; Gee Eun Lee; Hee Jin Cheong; Heedo Park; Jeonghun Kim; Ji Yun Noh; Jin Gu Yoon; Jin Il Kim; Joon Young Song; Joon-Yong Bae; Jungmin Lee; Juyoung Cho; Kyeong-ryeol Shin; Man-Seong Park; Woo Joo Kim |
| EPI_ISL_812956, EPI_ISL_812957, EPI_ISL_812958, EPI_ISL_812959, EPI_ISL_812961 | Division of Pathogen Resource Management, Korea National Institute of Health, Korea Disease Control and Prevention Agency | Division of Pathogen Resource Management, Korea National Institute of Health, Korea Disease Control and Prevention Agency | Choi; Choi, C.; E.-J.; J.-M.; Kim; M.-r.; S.T.; S.Y.; Y.S.; Yun |
| EPI_ISL_412869, EPI_ISL_412870, EPI_ISL_412871, EPI_ISL_412873, EPI_ISL_425118, EPI_ISL_426163, EPI_ISL_426167, EPI_ISL_426168, EPI_ISL_426171, EPI_ISL_426173, EPI_ISL_426180, EPI_ISL_426181, EPI_ISL_426182, EPI_ISL_426183, EPI_ISL_426187, EPI_ISL_427812, EPI_ISL_471453, EPI_ISL_481371, EPI_ISL_497962, EPI_ISL_497963, EPI_ISL_497964, EPI_ISL_497965, EPI_ISL_497966, EPI_ISL_498004, EPI_ISL_498005, EPI_ISL_498037, EPI_ISL_498043, EPI_ISL_506957, EPI_ISL_506958, EPI_ISL_506959, EPI_ISL_506960, EPI_ISL_506961, EPI_ISL_506962, EPI_ISL_506963, EPI_ISL_506964 | | | |
| see above | Division of Viral Diseases, Center for Laboratory Control of Infectious Diseases, Korea Centers for Diseases Control and Prevention | Division of Viral Diseases, Center for Laboratory Control of Infectious Diseases, Korea Centers for Diseases Control and Prevention | Daesang Lee; Dong Hyun Song; Heui Man Kim; Hye-Jun Jo; Jeong-Min Kim; Jun-Sub Kim; Junhyeong Jang; Mi-Seon Kim; Myung Guk Han; Namjoo Lee; Sang Hee Woo; Sehee Park; Seong Tae Jeong; Yoon-Seok Chung |
| EPI_ISL_479636, EPI_ISL_479645, EPI_ISL_481744, EPI_ISL_482471 | Dr. Georges-L.-Dumont University Hospital Centre | National Microbiology Laboratory | Anna Majer; Elsie Grudeski; Gary Van Domselaar; Grace Seo; Guillaume Desnoyers; Jennifer Tanner; Kristyn Burak; Morag Graham; Natalie Knox; Nathalie Bastien; Philip Mabon; Rhiannon Huzarewich; Richard Garceau; Russell Mandes; Shari Tyson; Timothy Booth; Yan Li |
| EPI_ISL_906760, EPI_ISL_906764 | Dr. Leonard A. Miller Centre for Health Services | National Microbiology Laboratory (NML) | Adel Malek; Anna Majer; Anneliese Landgraff; CanCOGeN's metadata curation team; Darian Hole; Elsie Grudeski; Gary Van Domselaar; George Zahariadis; Grace Seo; Jennifer Tanner; Kerri Smith; Laura Gilbert; Madison Chapel; Morag Graham; Natalie Knox; Nathalie Bastien; Philip Mabon; Public Health Agency of Canada CanCOGeN team; Rhiannon Huzarewich; Robert Needle; Russell Mandes; Shari Tyson; Timothy Booth; Yan Li; Yang Yu |
| EPI_ISL_414555, EPI_ISL_415461, EPI_ISL_415503, EPI_ISL_415526, EPI_ISL_422648, EPI_ISL_422803, EPI_ISL_422872, EPI_ISL_455255, EPI_ISL_461108, EPI_ISL_461154, EPI_ISL_461158, EPI_ISL_523504, EPI_ISL_523604 | | | |
| see above | Dutch COVID-19 response team | Erasmus Medical Center | Anne van der Linden; Annemiek van der Eijk; Aura Timen; Bas Oude Munnink; Claudia Schapendonk; Corien Swaan; Corine GeurtsvanKessel; David Nieuwenhuijse; Irina Chestakova; Jeroen van Kampen; Jolanda Voermans; Madelief Mollers; Manon Haverkate; Marion Koopmans; Mark Pronk; Mart Stein; Pascal Lexmond; Reina Sikkema; Richard Molenkamp; Sandra Kengne Kamga Mobou; Stefan van Nieuwkoop; Theo Bestebroer; on behalf of the Dutch national COVID-19 response team. |
| EPI_ISL_454760, EPI_ISL_454764, EPI_ISL_547469, EPI_ISL_547470, EPI_ISL_547524, EPI_ISL_547529, EPI_ISL_804386 | | | |
| see above | Dutch COVID-19 response team | National Institute for Public Health and the Environment (RIVM) | Adam Meijer; AnneMarie van den Brandt; Bas van der Veer; Chantal Reusken; Dennis Schmitz; Dirk Eggink; Florian Zwagemaker; Harry Vennema; Jeroen Cremer; Matthijs Welkers; Pieter Overduin; Sharon van den Brink; on behalf of the national COVID-19 response team |
| EPI_ISL_2790694, EPI_ISL_2790697, EPI_ISL_2790699, EPI_ISL_2790700, EPI_ISL_2790705, EPI_ISL_2790707, EPI_ISL_2790708, EPI_ISL_2790711 | | | |
| see above | Emory Molecular Diagnostics Laboratory, Emory Healthcare | Genomics and Discovery, Respiratory Viruses Branch, Division of Viral Diseases, Centers for Disease Control and Prevention | Adam Retchless; Ahmed Babiker; Anna Kelleher; Anna Uehara; Anne Piantadosi; Brian Lynch; Clinton R. Paden; Dhwani Batra; Haibin Wang; Han Jia Justin Ng; Jasmine Padilla; Jing Zhang; Justin Lee; Krista Queen; Mark Burroughs; Mili Sheth; Morgan Davis; Peter Cook; Rachel Marine; Sarah Nobles; Suxiang Tong; Tara Coalter; Yan Li; Ying Tao |
| EPI_ISL_455355, EPI_ISL_455356, EPI_ISL_455360, EPI_ISL_754749, EPI_ISL_754750, EPI_ISL_754751, EPI_ISL_754754, EPI_ISL_754755, EPI_ISL_754756, EPI_ISL_754758, EPI_ISL_754759, EPI_ISL_754760, EPI_ISL_754762, EPI_ISL_754763, EPI_ISL_754764, EPI_ISL_754766, EPI_ISL_754768, EPI_ISL_754776, EPI_ISL_754777, EPI_ISL_1278099, EPI_ISL_1278100, EPI_ISL_1503972 | | | |
| see above | Emory Molecular Diagnostics Laboratory, Emory Healthcare | Piantadosi Lab, Emory Department of Pathology | Ahmed Babiker; Anne Piantadosi |
| EPI_ISL_424350, EPI_ISL_424351 | Environmental and Global Health | Environmental and Global Health | Alam, M.; C.J.; Elbadry; Gibson; J.A.; J.C.; J.G. Jr. and Lednicky; M.A.; Morris; Stephenson; Subramaniam, K.; T.B.; Waltzek |
| EPI_ISL_2836011 | Epidemiology of Microbial Diseases, Yale School of Public Health | Epidemiology of Microbial Diseases, Yale School of Public Health | A.E.; A.F.; Alpert, T.; Breban; Brito; C.B.; C.C.; Fauver; Grubaugh; I.M.; Iwasaki, A.; J.E.; J.R.; Kalinich; Landry; Lucas, C.; M.E.; M.I.; M.L.; N.D.; Ott; Petrone; Rothman; Vogels; Watkins |
| EPI_ISL_514648 | Essentia Health-St. Joseph's Medical Center | Minnesota Department of Health, Public Health Laboratory | Jacob Garfin; Matt Plumb; and Xiong Wang |
| EPI_ISL_424856 | FL Bur. of Public Health Laboratories-Jacksonville | Pathogen Discovery, Respiratory Viruses Branch, Division of Viral Diseases, Centers for Disease Control and Prevention | Alison S. Laufer Halpin; Anna Uehara; Christopher A. Elkins; Clinton R. Paden; Haibin Wang; Jing Zhang; Krista Queen; Mary S. Keckler; Rachel Marine; Suxiang Tong; Yan Li; Ying Tao |
| EPI_ISL_424854 | FL Bureau of Public Health Laboratories-Miami | Pathogen Discovery, Respiratory Viruses Branch, Division of Viral Diseases, Centers for Disease Control and Prevention | Alison S. Laufer Halpin; Anna Uehara; Christopher A. Elkins; Clinton R. Paden; Haibin Wang; Jing Zhang; Krista Queen; Mary S. Keckler; Rachel Marine; Suxiang Tong; Yan Li; Ying Tao |
| EPI_ISL_419560 | FL Bureau of Public Health Laboratories-Tampa | Pathogen Discovery, Respiratory Viruses Branch, Division of Viral Diseases, Centers for Disease Control and Prevention | Anna Uehara; Clinton R. Paden; Haibin Wang; Jasmine Padilla; Jing Zhang; Justin Lee; Krista Queen; Suxiang Tong; Yan Li; Ying Tao |
| EPI_ISL_418252 | FUNDACION JIMENEZ DIAZ | Instituto de Salud Carlos III | A. Monzón; F. Casas; I. Fernández Roblas, R.; I. Jiménez; Iglesias-Caballero; M. Camarero; M. Cuesta; M. González-Esguevillas; M. Molinero Calamita; M. Zaballos; P. Jiménez; S. Juliá; S. Pozo; S. Varona |
| EPI_ISL_480792, EPI_ISL_480793, EPI_ISL_480795, EPI_ISL_480796, EPI_ISL_480797, EPI_ISL_480803, EPI_ISL_480813, EPI_ISL_489800, EPI_ISL_489801, EPI_ISL_489802, EPI_ISL_495346, EPI_ISL_495347, EPI_ISL_495356, EPI_ISL_495391, EPI_ISL_495393, EPI_ISL_508740, EPI_ISL_508741, EPI_ISL_508769, EPI_ISL_508772, EPI_ISL_508775, EPI_ISL_508779, EPI_ISL_508788, EPI_ISL_508799, EPI_ISL_508801, EPI_ISL_508803, EPI_ISL_508806, EPI_ISL_508807, EPI_ISL_509716, EPI_ISL_509718, EPI_ISL_509720, EPI_ISL_509723, EPI_ISL_509734, EPI_ISL_509740, EPI_ISL_509741, EPI_ISL_509755, EPI_ISL_509758, EPI_ISL_509772, EPI_ISL_514141, EPI_ISL_514142, EPI_ISL_517788, EPI_ISL_653155, EPI_ISL_653156, EPI_ISL_653157, EPI_ISL_653177, EPI_ISL_653178, EPI_ISL_653179, EPI_ISL_653184, EPI_ISL_653253, EPI_ISL_653254, EPI_ISL_653255, EPI_ISL_653258, EPI_ISL_653259, EPI_ISL_653260, EPI_ISL_653261, EPI_ISL_653268, EPI_ISL_653269, EPI_ISL_653270 | | | |
| see above | Florida Bureau of Public Health Laboratories | Florida Bureau of Public Health Laboratories | Jason Blanton; Sarah Schmedes |
| EPI_ISL_541145, EPI_ISL_541237 | Florida Bureau of Public Health Laboratories, Florida Department of Health | Florida Bureau of Public Health Laboratories, Florida Department of Health | Blanton, J.; Schmedes, S. |
| EPI_ISL_632119 | Flushing Hospital Medical Center | New York City Public Health Laboratory | Jade Wang; et al. |
| EPI_ISL_2333084 | Foerde Hospital, Department of Microbiology | Norwegian Institute of Public Health, Department of Virology | Atiya R Ali; Debech Nadia; Engebretsen Serina Beate; Garcia Llorente Ignacio; Hilde Elshaug; Hilde Vollan; Jon Bråte; Kamilla Heddeland Instefjord; Karoline Bragstad; Kathrine Stene-Johansen; Line Victoria Moen; Marie Paulsen Madsen; Olav Hungnes; Pedersen Benedikte Nevjen; Rasmus Riis Kopperud |
| EPI_ISL_2433280, EPI_ISL_2433281, EPI_ISL_2433282, EPI_ISL_2433283, EPI_ISL_2433284 | Fourth affiliated hospital, college of medicine, zhejiang university | Fourth affiliated hospital, college of medicine, zhejiang university | huina tang; ke xing; lele li; shufa zheng; tian liu; wei xu; wenxu yu; xiaoping xia; xueke liu; yingping wu; yutong he |
| EPI_ISL_411060, EPI_ISL_431785 | Fujian Center for Disease Control and Prevention | Fujian Center for Disease Control and Prevention | Chen Wei; He Wenxiang; Huang Zhimiao; Lin Qi; Weng Yuwei; Zhang Yanhua |
| EPI_ISL_862839 | Fujita Health University Bantane Hospital | Fujita health University, School of Medicine, Department of Microbiology | Masahiro suzuki; Yohei Doi |
| EPI_ISL_862837, EPI_ISL_862838, EPI_ISL_862840, EPI_ISL_862841 | Fujita Health University Hospital | Fujita health University, School of Medicine, Department of Microbiology | Masahiro suzuki; Yohei Doi |
| EPI_ISL_684422 | Fukuoka Institute of Health and Environmental Sciences | Pathogen Genomics Center, National Institute of Infectious Diseases | Kentaro Itokawa; Makoto Kuroda; Masanori Hashino; Rina Tanaka; Tsuyoshi Sekizuka |
| EPI_ISL_462478 | Fundación Jiménez Díaz | Instituto de Salud Carlos III | A. Monzón; F. Casas; I; I. Jiménez; Iglesias-Caballero; M. Camarero; M. Cuesta; M. González-Esguevillas; M. Molinero Calamita; M. Zaballos; P. Jiménez; R. Fernández; S. Juliá; S. Pozo; S. Varona |
| EPI_ISL_420786, EPI_ISL_420788 | GA Department of Public Health | Pathogen Discovery, Respiratory Viruses Branch, Division of Viral Diseases, Centers for Disease Control and Prevention | Alison S. Laufer Halpin; Anne Uehara; Christopher A. Elkins; Clinton R. Paden; Haibin Wang; Jasmine Padilla; Jing Zhang; Justin Lee; Krista Queen; Mary S. Keckler; Rachel Marine; Suxiang Tong; Yan Li; Ying Tao |
| EPI_ISL_424859, EPI_ISL_424861, EPI_ISL_424864, EPI_ISL_426417 | GA Department of Public Health Laboratory | Pathogen Discovery, Respiratory Viruses Branch, Division of Viral Diseases, Centers for Disease Control and Prevention | Alison S. Laufer Halpin; Anna Uehara; Christopher A. Elkins; Clinton R. Paden; Haibin Wang; Jing Zhang; Krista Queen; Mary S. Keckler; Rachel Marine; Suxiang Tong; Yan Li; Ying Tao |
| EPI_ISL_583463, EPI_ISL_583464 | Garcia-Sastre Laboratory, Department of Microbiology, Icahn School of Medicine at Mount Sinai | van Bakel Laboratory, Genetics and Genomics Sciences, Icahn School of Medicine at Mount Sinai | Adolfo García-Sastre; Adriana van de Guchte; Ajay Obla; Ana S. Gonzalez-Reiche; Genovefa Papanicolaou; Gunjan Shah; Harm van Bakel; Jayeeta Dutta; Judith Aberg; Kent Sepkowitz; Miguel-Angel Perales; Ngolela Esther Babady; Sadaf Aslam; Teresa Aydillo; Tobias Hohl; Zenab Khan; and Mini Kamboj |

| | | | |
|---|---|---|---|
| EPI_ISL_430064 | Geelong Centre for Emerging Infectious Diseases | Geelong Centre for Emerging Infectious Diseases | Alexandersen S.; Bhatta T.R.; Chamings A. |
| EPI_ISL_406801 | General Hospital of Central Theater Command of People's Liberation Army of China | BGI & Institute of Microbiology, Chinese Academy of Sciences & Shandong First Medical University & Shandong Academy of Medical Sciences & General Hospital of Central Theater Command of People's Liberation Army of China | Weifeng Shi and Zhenhong Hu; Weijun Chen; Yuhai Bi |
| EPI_ISL_594456 | Georgia Public Health Laboratory | Pathogen Discovery, Respiratory Viruses Branch, Division of Viral Diseases, Centers for Disease Control and Prevention | Anna Uehara; Clinton Paden; Haibin Wang; Jing Zhang; Julu Bhatnagar; Krista Queen; Suxiang Tong; Yan Li; Ying Tao |
| EPI_ISL_480083, EPI_ISL_480084, EPI_ISL_480085, EPI_ISL_480086, EPI_ISL_480087, EPI_ISL_480088, EPI_ISL_480089 | | | |
| see above | Gifu Prefectural Institute of Public Health and Environmental Sciences | Pathogen Genomics Center, National Institute of Infectious Diseases | Hajime Kamiya; Kentaro Itokawa; Makoto Kuroda; Masanori Hashino; Motoi Suzuki; Rina Tanaka; Tsuyoshi Sekizuka; Yoshihiko Kameyama |
| EPI_ISL_496603, EPI_ISL_496604, EPI_ISL_496605, EPI_ISL_496606, EPI_ISL_496607, EPI_ISL_496608, EPI_ISL_496609, EPI_ISL_496610, EPI_ISL_496611, EPI_ISL_496612, EPI_ISL_496616, EPI_ISL_496617, EPI_ISL_496618, EPI_ISL_496619, EPI_ISL_496621, EPI_ISL_496627, EPI_ISL_496630, EPI_ISL_496631, EPI_ISL_496632, EPI_ISL_496633, EPI_ISL_496634, EPI_ISL_496636, EPI_ISL_496637, EPI_ISL_496638, EPI_ISL_496640, EPI_ISL_496641, EPI_ISL_496642, EPI_ISL_496643, EPI_ISL_496644, EPI_ISL_496645, EPI_ISL_496646, EPI_ISL_496649, EPI_ISL_496650, EPI_ISL_496652, EPI_ISL_496654, EPI_ISL_496655, EPI_ISL_496657, EPI_ISL_496658, EPI_ISL_496659, EPI_ISL_496660, EPI_ISL_496663, EPI_ISL_496664, EPI_ISL_496665, EPI_ISL_496666, EPI_ISL_496669, EPI_ISL_496670, EPI_ISL_496671, EPI_ISL_496677, EPI_ISL_496678, EPI_ISL_496679, EPI_ISL_496681, EPI_ISL_496682, EPI_ISL_496684, EPI_ISL_496685, EPI_ISL_496692, EPI_ISL_496693, EPI_ISL_496694, EPI_ISL_496699, EPI_ISL_496700, EPI_ISL_496701, EPI_ISL_496703, EPI_ISL_496705, EPI_ISL_496706, EPI_ISL_496708, EPI_ISL_496709, EPI_ISL_496710, EPI_ISL_496711, EPI_ISL_496712, EPI_ISL_496713, EPI_ISL_496714, EPI_ISL_496716, EPI_ISL_496718, EPI_ISL_496719, EPI_ISL_496721, EPI_ISL_496722, EPI_ISL_496723, EPI_ISL_496724, EPI_ISL_496725, EPI_ISL_496726, EPI_ISL_496727, EPI_ISL_496728, EPI_ISL_496730, EPI_ISL_496731, EPI_ISL_496732, EPI_ISL_496733, EPI_ISL_496734, EPI_ISL_496735, EPI_ISL_496736, EPI_ISL_496737, EPI_ISL_496740, EPI_ISL_496741, EPI_ISL_496742, EPI_ISL_496745, EPI_ISL_496749, EPI_ISL_496750, EPI_ISL_496751, EPI_ISL_496753, EPI_ISL_496756, EPI_ISL_496764, EPI_ISL_496767, EPI_ISL_496768, EPI_ISL_496770, EPI_ISL_496771, EPI_ISL_496772, EPI_ISL_496773, EPI_ISL_496774, EPI_ISL_496775, EPI_ISL_496776, EPI_ISL_496777, EPI_ISL_496778, EPI_ISL_496781, EPI_ISL_496782, EPI_ISL_496783, EPI_ISL_496784, EPI_ISL_496785, EPI_ISL_496786, EPI_ISL_496787, EPI_ISL_496788, EPI_ISL_496790, EPI_ISL_496791, EPI_ISL_496792, EPI_ISL_496793, EPI_ISL_496795, EPI_ISL_496796, EPI_ISL_496797, EPI_ISL_496798, EPI_ISL_1225382 | | | |
| see above | Gorgas Memorial Laboratory of Health Studies | Gorgas Memorial Laboratory of Health Studies | Adriana Weeden; Alejandra Valoy; Alexander A Martinez; Alexander Martinez; Ambar Moreno; Anyuri Ortiz; Brechla Moreno; Claudia Gonzalez; Claudia Gonzalez Sandra Lopez-Verges; Daniel Castillo; Danilo Franco; Davis Beltran; Dimelza Arauz; Elimelec Valdespino; Gretel Vasquez; Ilka Guerra; Isela Guerrero; Jessica Gondola; Jim Chang; Juan Miguel Pascale; Layda Abrego; Lisseth Saenz; Mabel Martinez-Montero; Maria Chen-German; Marlene Castillo; Melissa Gaitan; Oris Chavarria; Rita Corrales; Rita Rodriguez; Sandra Lopez-Verges; Yamilka Diaz; Yaneth Pitti; Zumara Chaverra |
| EPI_ISL_447734 | Grupo de Investigaciones Microbiológicas-UR (GIMUR), Departamento de Biología, Facultad de Ciencias Naturales, Universidad del Rosario, Bogotá, Colombia | Grupo de Investigaciones Microbiológicas-UR (GIMUR), Departamento de Biología, Facultad de Ciencias Naturales, Universidad del Rosario, Bogotá, Colombia Instituto Nacional de Salud, Bogotá, Colombia Icahn School of Medicine at Mount Sinai, New York, USA | Adriana Castillo; Alberto Paniz-Mondolfi; Ana S. Gonzalez-Reiche; Angelica Rico; Aníbal A. Teherán; Carolina Florez; Carolina Hernandez; David Martínez; Emilia Mia Sordillo; Esther C. Barros; Harm van Bakel; Jesús E. Jaimes; Juan David Ramírez; Laura Vega; Lisseth Pardo; Marina Muñoz; Martha L. Ospina; Matthew M. Hernandez; Nathalia Ballesteros; Sergio Castañeda; Sergio Gomez; Viviana Simon |
| EPI_ISL_403932, EPI_ISL_403933, EPI_ISL_403935 | Guangdong Provincial Center for Diseases Control and Prevention; Guangdong Provincial Public Health | Department of Microbiology, Guangdong Provincial Center for Diseases Control and Prevention | Baisheng Li; Changwen Ke; Feng Ruan; Guanhao He; Haojie Zhong; Huihong Deng; Jianfeng He; Jianpeng Xiao; Jianxiang Geng; Jianxiong Hu; Jie Wu; Jing Lu; Lifeng Lin; Lijun Liang; Lirong Zou; Min Kang; Qi Zhu; Shujiang Mei; Songjian Xiao; Tao Liu; Tie Song; Weilin Zeng; Wenjun Ma; Xing Li; Xiujuan Tang; Xue Zhuang; Xuguang Chen; Ying Wang; Yingchao Song; Yingtao Zhang; Yuhuang Liao; Zhe Liu |
| EPI_ISL_428441, EPI_ISL_428443, EPI_ISL_428447, EPI_ISL_428448, EPI_ISL_428449, EPI_ISL_428457, EPI_ISL_428458, EPI_ISL_428462, EPI_ISL_428463, EPI_ISL_428470, EPI_ISL_428471, EPI_ISL_428476 | | | |
| see above | Guangdong Provincial Center for Diseases Control and Prevention;Guangdong Provincial Institute of Public Health | School of Public Health, The University of Hong Kong | Bosheng Li; Hanri Zeng; Haogao Gu; Hui-Ling Yen; Jie Wu; Leo L.M. Poon; Lijun Liang; Tie Song; Yao Hu; Yingchao Song; Zhencui Li |
| EPI_ISL_413853, EPI_ISL_413854, EPI_ISL_413855, EPI_ISL_413856, EPI_ISL_413858, EPI_ISL_413859, EPI_ISL_413860, EPI_ISL_413862, EPI_ISL_413865, EPI_ISL_413867, EPI_ISL_413873, EPI_ISL_413876, EPI_ISL_413881, EPI_ISL_413889, EPI_ISL_413890, EPI_ISL_413891, EPI_ISL_413892, EPI_ISL_413897, EPI_ISL_413900 | | | |
| see above | Guangdong Provincial Institution of Public Health, Guangdong Provinical Center for Disease Control and Prevention | Guangdong Provincial Institution of Public Health | Andrew Rambaut; Bo Peng; Changwen Ke; Chuming Liang; Huanying Zheng; Huifang Lin; Jing Lu; Jingju Peng; Jiufeng Sun; Josh Quick; Juan Su; Kang Min; Kuibiao Li; Lilian Zeng; Liu Zhe; Louis du Plessis; Minfeng Liang; Moritz Kraemer; Nick Loman; Nuno Faria; Oliver Pybus; Pingping Zhou; Qianlin Xiong; Ru bai; Ruilin Sun; Runyu Yuan; Sarah François; Shisong Fang; Song Tie; Tao Liu; Verity Hill; Wei Li; Wenjun Ma; Wenzhe Su; Xi Tang |
| EPI_ISL_509701 | Guatemala Ministry of Public Health | Pathogen Discovery, Respiratory Viruses Branch, Division of Viral Diseases, Centers for Disease Control and Prevention | Anna Uehara; Clinton Paden; Haibin Wang; Jing Zhang; Krista Queen; Suxiang Tong; Yan Li; Ying Tao |
| EPI_ISL_418185, EPI_ISL_418186 | Gundersen Molecular Diagnostic Laboratory | Kabara Cancer Research Institute | Craig S. Richmond & Paraic A. Kenny |
| EPI_ISL_418187, EPI_ISL_418189 | Gundersen Molecular Diagnostics Laboratory | Kabara Cancer Research Institute | Craig S. Richmond & Paraic A. Kenny |
| EPI_ISL_511406 | H Braga | Instituto Nacional de Saude (INSA) | Borges et al |
| EPI_ISL_421466, EPI_ISL_421467, EPI_ISL_453840, EPI_ISL_453844, EPI_ISL_453845, EPI_ISL_453846, EPI_ISL_453847, EPI_ISL_453848, EPI_ISL_511716 | | | |
| see above | H Evora | Instituto Nacional de Saude (INSA) | Borges et al; Guiomar et al |
| EPI_ISL_455587 | H.R.H. Maha Chakri Sirindhorn Medical Center | National Institute of Health. Department of medical Sciences, Ministry of Public Health, Thailand | Chittaganpitch; Malinee; Okada; Parnmen; Phuygun; Pilailuk; Siripaporn; Sittiporn; Sunthareeya; Thanadachakul; Thanutsapa; Waicharoen; Warawan; Wongboot |
| EPI_ISL_426421 | HI Dept. of Health, State Laboratories Division | Pathogen Discovery, Respiratory Viruses Branch, Division of Viral Diseases, Centers for Disease Control and Prevention | Alison S. Laufer Halpin; Anna Uehara; Christopher A. Elkins; Clinton R. Paden; Haibin Wang; Jing Zhang; Krista Queen; Mary S. Keckler; Rachel Marine; Suxiang Tong; Yan Li; Ying Tao |
| EPI_ISL_418253, EPI_ISL_419240, EPI_ISL_419709 | HOSPITAL TXAGORRITXU | Instituto de Salud Carlos III | A. Monzón; C; F. Casas; I. Gomez-Gonzalez C.; I. Gómez; I. Gómez, C.; I. Jiménez; Iglesias-Caballero; M. Camarero; M. Camarero S. Pozo F. Casas I. Jiménez; M. Cuesta; M. González-Esguevillas; M. Molinero Calamita; M. Zaballos; P. Jiménez; S. Juliá; S. Pozo; S. Varona |
| EPI_ISL_413487 | Harborview Medical Center | University of Washington Virology Lab | Alexander Greninger; Arun Nalla; Hong Xie; Keith Jerome; Pavitra Roychoudhury |
| EPI_ISL_479903, EPI_ISL_479904, EPI_ISL_479905, EPI_ISL_479906, EPI_ISL_479907, EPI_ISL_479908, EPI_ISL_479909, EPI_ISL_479910, EPI_ISL_479912 | | | |
| see above | Himeji City Institute of Environment and Health | Pathogen Genomics Center, National Institute of Infectious Diseases | Hajime Kamiya; Kentaro Itokawa; Makoto Kuroda; Masanori Hashino; Motoi Suzuki; Rina Tanaka; Tsuyoshi Sekizuka |
| EPI_ISL_479815, EPI_ISL_479819, EPI_ISL_479820 | Hokkaido Institute of Public Health | Pathogen Genomics Center, National Institute of Infectious Diseases | Hajime Kamiya; Kentaro Itokawa; Makoto Kuroda; Masanori Hashino; Motoi Suzuki; Rika Komagome; Rina Tanaka; Tsuyoshi Sekizuka |
| EPI_ISL_412028 | Hong Kong Department of Health | School of Public Health, The University of Hon g Kong | Daniel K.W. Chu; Dominic N.C. Tsang; Leo L.M. Poon; Malik Peiris |
| EPI_ISL_2958706, EPI_ISL_2958717, EPI_ISL_2958718, EPI_ISL_2958731 | Hospital | National Reference Center for Viruses of Respiratory Infections, Institut Pasteur, Paris | Angela Brisebarre; BARBUT Patricia; Camille Capel; Christophe Malabat; Corinne Maufrais; Etienne Simon-Lorière; Frédéric Lemoine; Hub Bioinformatique Biostatistiques; Louise Lefrançois; Marion Barbet; Maud Vanpeene; Méline Bizard; Sylvie Behillil; Sylvie Van der Werf; Vincent Enouf |
| EPI_ISL_500219, EPI_ISL_500221, EPI_ISL_510269, EPI_ISL_510270, EPI_ISL_510283, EPI_ISL_510284, EPI_ISL_510290 | | | |
| see above | Hospital Clínico Universitario de Santiago de Compostela | SeqCOVID-SPAIN consortium/IBV(CSIC) | Amparo Coira Nieto; Antonio Aguilera Guirao; Gema Barbeito Castiñeiras; José Javier Costa Alcalde; Mª Luisa Pérez del Molino Bernal; Rocio Trastoy Pena and SeqCOVID-SPAIN consortium |
| EPI_ISL_462477, EPI_ISL_539499 | Hospital Costa del Sol | Instituto de Salud Carlos III | A. Monzón; F. Casas; F. Fernández; I; I. Jiménez; Iglesias-Caballero; M. Camarero; M. Cuesta; M. González-Esguevillas; M. Molinero Calamita; M. Zaballos; P. Jiménez; S. Juliá; S. Pozo; S. Varona |
| EPI_ISL_512655 | Hospital De Niños Dr. Carlos Saenz Herrera [San Jose/San Jose] | Inciensa, Instituto Costarricense de Investigación y Enseñanza en Nutrición y Salud | Adriana Godinez & Melany Calderon; Claudio Soto-Garita; Estela Cordero; Francisco Duarte; Hebleen Porras |
| EPI_ISL_3190068 | Hospital General Universitario Gregorio Marañón | Hospital General Universitario Gregorio Marañón | Cristina Rodriguez-Grande; Darío García de Viedma; Julia Suárez; Laura Pérez-Lago; Marta Herranz Martin; Patricia Muñoz; Pedro Sola Campoy; Pilar Catalán; Sergio Buenestado Serrano; Victor Manuel de la Cueva |
| EPI_ISL_481044, EPI_ISL_481051, EPI_ISL_481052, EPI_ISL_481054, EPI_ISL_481055, EPI_ISL_481057, EPI_ISL_481058, EPI_ISL_481059, EPI_ISL_481060, EPI_ISL_481066, EPI_ISL_481075, EPI_ISL_481087, EPI_ISL_481089, EPI_ISL_481094, EPI_ISL_481096, EPI_ISL_510121, EPI_ISL_510122, EPI_ISL_510146, EPI_ISL_510150, EPI_ISL_510169, EPI_ISL_510171, EPI_ISL_510177, EPI_ISL_510178, EPI_ISL_510180, EPI_ISL_510181, EPI_ISL_510182, EPI_ISL_510183, EPI_ISL_510184, EPI_ISL_510185, EPI_ISL_510187, EPI_ISL_510190, EPI_ISL_510192, EPI_ISL_510195, EPI_ISL_510196, EPI_ISL_510197, EPI_ISL_510198, EPI_ISL_510206, EPI_ISL_510208, EPI_ISL_510209, EPI_ISL_510210, EPI_ISL_510213, EPI_ISL_510215, EPI_ISL_510219, EPI_ISL_510220, EPI_ISL_510228, EPI_ISL_510230, EPI_ISL_510235, EPI_ISL_510237, EPI_ISL_510240, EPI_ISL_654082, EPI_ISL_654188 | | | |
| see above | Hospital General Universitario Gregorio Marañón | SeqCOVID-SPAIN consortium/IBV(CSIC) | Darío García de Viedma; Darío García de Viedma and SeqCOVID-SPAIN consortium; Jon Sicilia; Julia Suárez; Laura Pérez-Lago; Marta Herranz; Patricia Muñoz; Patricia Muñoz and SeqCOVID-SPAIN consortium; Pilar Catalán |
| EPI_ISL_418245, EPI_ISL_418246 | Hospital General y Universitario de Guadalajara | Instituto de Salud Carlos III | A. Monzón; F. Casas; I. Gonzalez-Praetorius A.; I. Jiménez; Iglesias-Caballero; M. Camarero; M. Cuesta; M. González-Esguevillas; M. Molinero Calamita; M. Zaballos; P. Jiménez; S. Juliá; S. Pozo; S. Varona |
| EPI_ISL_862637 | Hospital Nuestra Señora de Sonsoles | Instituto de Salud Carlos III | A. Monzón; A. San Pedro; F. Casas; I; I. Jiménez; Iglesias-Caballero; M. Camarero; M. Cuesta; M. González-Esguevillas; M. Molinero Calamita; M. Zaballos; P. Jiménez; S. Juliá; S. Pozo; S. Varona |
| EPI_ISL_1502818 | Hospital Regional Anita Moreno Los Santos | Gorgas Memorial Laboratory of Health Studies | Best Dayana; Castillo Jorge; Espino Juliee; Franco Danilo; Gonzalez Claudia; Jessica Gondola; Leyda Abrego; Lopez-Verges Sandra; Marlenne Castillo; Martinez Alexander; Moreno Ambar; Moreno Brechla; Oris Chavarria; Ortiz Alma |
| EPI_ISL_419234, EPI_ISL_455336, EPI_ISL_455337, EPI_ISL_455338, EPI_ISL_455339, EPI_ISL_455340, EPI_ISL_455341, EPI_ISL_455343, EPI_ISL_578196, EPI_ISL_578197, EPI_ISL_578198, EPI_ISL_578199, EPI_ISL_578200, EPI_ISL_578201 | | | |
| see above | Hospital San Pedro | Instituto de Salud Carlos III | A. Monzón; Alonso, C.; C. Alonso; Camarero, S.; Casas, I.; Cuesta, I.; F. Casas; González-Esguevillas, M.; I; I. Jiménez; Iglesias-Caballero; Iglesias-Caballero, M.; J.M Azcona; Jiménez, M.; Jiménez, P.; Juliá, M.; M. Blasco; M. Camarero; M. Cuesta; M. González-Esguevillas; M. Molinero Calamita; M. Zaballos; Molinero Calamita, M.; Monzón, S.; P. Jiménez; Pozo, F.; S. Juliá; S. Pozo; S. Varona; Varona, S.; Zaballos, A. |
| EPI_ISL_510325, EPI_ISL_510327, EPI_ISL_510329, EPI_ISL_613411 | Hospital San Pedro de Alcántara (Cáceres) | SeqCOVID-SPAIN consortium/IBV(CSIC) | Cristina Muñoz Cuevas; Guadalupe Rodriguez Rodriguez and SeqCOVID-SPAIN consortium |
| EPI_ISL_434536 | Hospital San Vicente de Paul | Inciensa, Instituto Costarricense de Investigación y Enseñanza en Nutrición y Salud | Adriana Godinez & Melany Calderon; Claudio Soto-Garita; Estela Cordero; Francisco Duarte; Hebleen Porras |
| EPI_ISL_455350, EPI_ISL_455351 | Hospital Txagorritxu | Instituto de Salud Carlos III | A. Monzón; C. Gómez; F. Casas; I; I. Jiménez; Iglesias-Caballero; M. Camarero; M. Cuesta; M. González-Esguevillas; M. Molinero Calamita; M. Zaballos; P. Jiménez; S. Juliá; S. Pozo; S. Varona |
| EPI_ISL_444990, EPI_ISL_819300 | Hospital Universitari Vall d'Hebron - Vall d'Hebron Institut de Recerca | Hospital Universitari Vall d'Hebron - Vall d'Hebron Institut de Recerca | Andrés Antón; Ariadna Rando; Carla Castillo; Cristina Andrés; Damir Garcia-Cehic; Josep F Abril; Josep Quer; Juliana Esperalba; Maria Carmen Martin; Maria Gema Codina; Maria Piñana; Mercedes Guerrero-Murillo; Tomàs Pumarola |
| EPI_ISL_538007, EPI_ISL_538008, EPI_ISL_538010, EPI_ISL_538011, EPI_ISL_538013, EPI_ISL_538014, EPI_ISL_538015, EPI_ISL_538019, EPI_ISL_538024, EPI_ISL_538032, EPI_ISL_538033, EPI_ISL_538042, EPI_ISL_538053 | | | |
| see above | Hospital Universitari i Politècnic La Fe de València | SeqCOVID-SPAIN consortium/IBV(CSIC) | Ana Gil Brusola; Eva González Barbera; José Luis López Hontangas and SeqCOVID-SPAIN consortium; María Dolores Gómez Ruiz; Salvador Giner Almaraz |

| | | | |
|---|---|---|---|
| EPI_ISL_428683, EPI_ISL_428691, EPI_ISL_428692, EPI_ISL_428698, EPI_ISL_428699, EPI_ISL_428700, EPI_ISL_428704, EPI_ISL_428710, EPI_ISL_529983, EPI_ISL_529992 | | | |
| see above | Hospital Universitario 12 de Octubre | Hospital Universitario 12 de Octubre | Elias Dahdouh; Esther Viedma; Fernando Lázaro; Jesús Mingorance; Juan Carlos Galán; Julio García; Mª Dolores Folgueira; Natalia Stella; Rafael Cantón; Rafael Delgado; Raúl Recio; Sara González |
| EPI_ISL_417961 | Hospital Universitario 12 de Octubre | Hospital Universitario La Paz | Elias Dahdouh; Esther Viedma; Fernando Lázaro; Jesús Mingorance; Juan Carlos Galán; Julio García; Mª Dolores Folgueira; Natalia Stella; Rafael Cantón; Rafael Delgado; Sara González |
| EPI_ISL_467092, EPI_ISL_467093, EPI_ISL_467095, EPI_ISL_467096, EPI_ISL_467097, EPI_ISL_467098, EPI_ISL_467099, EPI_ISL_467100, EPI_ISL_467101, EPI_ISL_467102, EPI_ISL_467103, EPI_ISL_467104, EPI_ISL_467105, EPI_ISL_467106, EPI_ISL_467107, EPI_ISL_467108, EPI_ISL_467109, EPI_ISL_467110, EPI_ISL_467111, EPI_ISL_467112, EPI_ISL_467113, EPI_ISL_467114, EPI_ISL_467115, EPI_ISL_467116, EPI_ISL_467117, EPI_ISL_467118, EPI_ISL_467120, EPI_ISL_467121, EPI_ISL_467122, EPI_ISL_467125, EPI_ISL_467126, EPI_ISL_467127, EPI_ISL_467129, EPI_ISL_467130, EPI_ISL_467131, EPI_ISL_467132, EPI_ISL_467133, EPI_ISL_467134, EPI_ISL_467135, EPI_ISL_467136, EPI_ISL_467137, EPI_ISL_467139, EPI_ISL_467140, EPI_ISL_467141, EPI_ISL_467142, EPI_ISL_467143, EPI_ISL_467144, EPI_ISL_467146, EPI_ISL_467147, EPI_ISL_467149, EPI_ISL_467150, EPI_ISL_467153, EPI_ISL_467154, EPI_ISL_467155, EPI_ISL_467156, EPI_ISL_467158, EPI_ISL_467159, EPI_ISL_467160, EPI_ISL_467161, EPI_ISL_467162, EPI_ISL_467164, EPI_ISL_467165, EPI_ISL_467166, EPI_ISL_467168, EPI_ISL_467169, EPI_ISL_467170, EPI_ISL_467171, EPI_ISL_467172, EPI_ISL_467173, EPI_ISL_467174, EPI_ISL_467175, EPI_ISL_467176, EPI_ISL_467177, EPI_ISL_467178, EPI_ISL_467180, EPI_ISL_467181, EPI_ISL_467182, EPI_ISL_467183, EPI_ISL_500228, EPI_ISL_500229, EPI_ISL_500230, EPI_ISL_500231, EPI_ISL_500232, EPI_ISL_500233, EPI_ISL_500234, EPI_ISL_500235, EPI_ISL_500236, EPI_ISL_500237, EPI_ISL_500238, EPI_ISL_500239, EPI_ISL_500240, EPI_ISL_500241, EPI_ISL_500242, EPI_ISL_500243, EPI_ISL_500244, EPI_ISL_500245, EPI_ISL_500246, EPI_ISL_500247, EPI_ISL_500248, EPI_ISL_500249, EPI_ISL_500250, EPI_ISL_500251, EPI_ISL_500252, EPI_ISL_500253, EPI_ISL_500254, EPI_ISL_500255, EPI_ISL_500256, EPI_ISL_500257, EPI_ISL_500258, EPI_ISL_500259, EPI_ISL_500260, EPI_ISL_500261, EPI_ISL_500262, EPI_ISL_500263, EPI_ISL_500264, EPI_ISL_500265, EPI_ISL_500266, EPI_ISL_500267, EPI_ISL_500268, EPI_ISL_500269, EPI_ISL_500270, EPI_ISL_500271, EPI_ISL_500272, EPI_ISL_500273, EPI_ISL_500274, EPI_ISL_500275, EPI_ISL_500276, EPI_ISL_500277, EPI_ISL_500278, EPI_ISL_500279, EPI_ISL_500280, EPI_ISL_500281, EPI_ISL_500282, EPI_ISL_500283, EPI_ISL_500284, EPI_ISL_500285, EPI_ISL_500286, EPI_ISL_509616, EPI_ISL_509617, EPI_ISL_509618, EPI_ISL_537876, EPI_ISL_537878, EPI_ISL_537879, EPI_ISL_537880, EPI_ISL_537881, EPI_ISL_537882, EPI_ISL_537883, EPI_ISL_537884, EPI_ISL_537885, EPI_ISL_537886, EPI_ISL_537887, EPI_ISL_537888, EPI_ISL_537889, EPI_ISL_537890, EPI_ISL_537892, EPI_ISL_537893, EPI_ISL_537894, EPI_ISL_537896, EPI_ISL_537897, EPI_ISL_537898, EPI_ISL_537900, EPI_ISL_537901, EPI_ISL_537902, EPI_ISL_537903, EPI_ISL_537904, EPI_ISL_537905, EPI_ISL_537907, EPI_ISL_537908, EPI_ISL_537909, EPI_ISL_537910, EPI_ISL_537911, EPI_ISL_537912, EPI_ISL_537913, EPI_ISL_537914, EPI_ISL_537916, EPI_ISL_537917, EPI_ISL_537918, EPI_ISL_537919, EPI_ISL_537920, EPI_ISL_537921, EPI_ISL_537922, EPI_ISL_537923, EPI_ISL_537924, EPI_ISL_537925, EPI_ISL_537926, EPI_ISL_537927, EPI_ISL_537928, EPI_ISL_537929, EPI_ISL_537930, EPI_ISL_537931, EPI_ISL_537932, EPI_ISL_537933, EPI_ISL_537934, EPI_ISL_537935, EPI_ISL_537936, EPI_ISL_537937, EPI_ISL_537938, EPI_ISL_537939, EPI_ISL_537940, EPI_ISL_537941, EPI_ISL_537942, EPI_ISL_537943, EPI_ISL_537944, EPI_ISL_538118, EPI_ISL_538231, EPI_ISL_538232 | | | |
| see above | Hospital Universitario Araba. Vitoria-Gasteiz | SeqCOVID-SPAIN consortium/IBV(CSIC) | Amaia Aguirre Quiñonero; Andrés Canut Blasco and SeqCOVID-SPAIN consortium; Andrés Canut Blasco. and SeqCOVID-SPAIN consortium; Carmen Gómez González; Ma Concepción Lecaroz Agara; Ma Rosario Almela Ferrer; Marina Fernández Torres; Mª Concepción Lecaroz Agara; Mª Rosario Almela Ferrer; Silvia Hernáez Crespo |
| EPI_ISL_452692, EPI_ISL_452693, EPI_ISL_452694, EPI_ISL_452695, EPI_ISL_452696, EPI_ISL_452697, EPI_ISL_452698, EPI_ISL_452699, EPI_ISL_452700, EPI_ISL_452701, EPI_ISL_452702, EPI_ISL_452703, EPI_ISL_452704, EPI_ISL_452705, EPI_ISL_452706, EPI_ISL_452707, EPI_ISL_452708, EPI_ISL_452709, EPI_ISL_452710, EPI_ISL_452711, EPI_ISL_452712, EPI_ISL_452713, EPI_ISL_452714, EPI_ISL_452715, EPI_ISL_452716, EPI_ISL_452717, EPI_ISL_452718, EPI_ISL_452719, EPI_ISL_452720, EPI_ISL_452721, EPI_ISL_452722, EPI_ISL_452723, EPI_ISL_452724, EPI_ISL_452725, EPI_ISL_452726, EPI_ISL_452727, EPI_ISL_452728, EPI_ISL_452729, EPI_ISL_452730, EPI_ISL_452731, EPI_ISL_452732, EPI_ISL_452733, EPI_ISL_452734, EPI_ISL_452735, EPI_ISL_452736, EPI_ISL_452737, EPI_ISL_452738, EPI_ISL_452739, EPI_ISL_452740, EPI_ISL_452741, EPI_ISL_452742, EPI_ISL_452743, EPI_ISL_452744, EPI_ISL_452745, EPI_ISL_452746, EPI_ISL_452747, EPI_ISL_452748, EPI_ISL_452749, EPI_ISL_452750, EPI_ISL_452751, EPI_ISL_452752, EPI_ISL_452753, EPI_ISL_452754, EPI_ISL_452755, EPI_ISL_452756, EPI_ISL_452757, EPI_ISL_452758, EPI_ISL_452759, EPI_ISL_452760, EPI_ISL_452761, EPI_ISL_452762, EPI_ISL_452763, EPI_ISL_452764, EPI_ISL_452765, EPI_ISL_452766, EPI_ISL_452767, EPI_ISL_452768, EPI_ISL_452769, EPI_ISL_452770, EPI_ISL_452771, EPI_ISL_452772, EPI_ISL_452773, EPI_ISL_452774, EPI_ISL_452775, EPI_ISL_452776, EPI_ISL_452777, EPI_ISL_452778, EPI_ISL_452779, EPI_ISL_452780, EPI_ISL_452781, EPI_ISL_452782, EPI_ISL_452783, EPI_ISL_452784, EPI_ISL_452785, EPI_ISL_452786 | | | |
| see above | Hospital Universitario Araba. Vitoria-Gasteiz, | SeqCOVID-SPAIN consortium/IBV(CSIC) | Amaia Aguirre Quiñonero; Andrés Canut Blasco and SeqCOVID-SPAIN consortium; Carmen Gómez González; Maria Concepción Lecaroz Agara; Maria Rosario Almela Ferrer; Marina Fernández Torres; Silvia Hernáez Crespo |
| EPI_ISL_428675, EPI_ISL_428676, EPI_ISL_428677, EPI_ISL_428678 | Hospital Universitario La Paz | Hospital Universitario 12 de Octubre | Elias Dahdouh; Esther Viedma; Fernando Lázaro; Jesús Mingorance; Juan Carlos Galán; Julio García; Mª Dolores Folgueira; Natalia Stella; Rafael Cantón; Rafael Delgado; Raúl Recio; Sara González |
| EPI_ISL_538691, EPI_ISL_538705, EPI_ISL_538706, EPI_ISL_538711, EPI_ISL_538715, EPI_ISL_538717, EPI_ISL_538718, EPI_ISL_538720, EPI_ISL_538724, EPI_ISL_538727, EPI_ISL_538728, EPI_ISL_538732, EPI_ISL_538733, EPI_ISL_538734, EPI_ISL_538735, EPI_ISL_538736, EPI_ISL_538740, EPI_ISL_538745, EPI_ISL_538747 | | | |
| see above | Hospital Universitario Marqués de Valdecilla - IDIVAL (Santander, Cantabria) | SeqCOVID-SPAIN consortium/IBV(CSIC) | Daniel Pablo Marcos; Jesús Rodríguez Rodríguez; Jose Manuel Méndez Legaza; María Eliecer Cano García; María Siller Ruiz and SeqCOVID-SPAIN consortium; Mónica Gozalo Margüello |
| EPI_ISL_452455, EPI_ISL_452457, EPI_ISL_452458, EPI_ISL_452459, EPI_ISL_452463, EPI_ISL_452467, EPI_ISL_467068, EPI_ISL_467071, EPI_ISL_467076 | | | |
| see above | Hospital Universitario Puerta del Mar de Cádiz - INIBICA | SeqCOVID-SPAIN consortium/IBV(CSIC) | Fátima-Galán-Sánchez; Manuel Rodríguez-Iglesias and SeqCOVID-SPAIN consortium; Manuel Rodríguez-Iglesias and SeqCOVID-SPAIN consortium; Salud Rodríguez-Pallares |
| EPI_ISL_530094, EPI_ISL_530095, EPI_ISL_530096, EPI_ISL_530097, EPI_ISL_530098, EPI_ISL_530099 | Hospital Universitario Ramón y Cajal | Hospital Universitario 12 de Octubre | Elias Dahdouh; Esther Viedma; Fernando Lázaro; Jesús Mingorance; Juan Carlos Galán; Julio García; Mª Dolores Folgueira; Natalia Stella; Rafael Cantón; Rafael Delgado; Raúl Recio; Sara González |
| EPI_ISL_417979, EPI_ISL_417980, EPI_ISL_417981 | Hospital Universitario Ramón y Cajal | Hospital Universitario La Paz | Elias Dahdouh; Esther Viedma; Fernando Lázaro; Jesús Mingorance; Juan Carlos Galán; Julio García; Mª Dolores Folgueira; Natalia Stella; Rafael Cantón; Rafael Delgado; Sara González |
| EPI_ISL_871973 | Hospital Universitario Severo Ochoa | Instituto de Salud Carlos III | A. Monzón; F. Casas; I. García; I. Jiménez; Iglesias-Caballero; M. Camarero; M. Cuesta; M. González-Esguevillas; M. Pozo; M. Zaballos; M.L.; P. Jiménez; S. Juliá; S. Molinero Calamita; S. Varona |
| EPI_ISL_419230 | Hospital Universitario Virgen de las Nieves | Instituto de Salud Carlos III | Camarero, S.; Casas, I.; Cuesta, I.; González-Esguevillas, M.; Iglesias-Caballero, M.; Jiménez, M.; Jiménez, P.; Juliá, M.; Molinero Calamita, M.; Monzón, S.; Pozo, F.; Sanbonmatsu, S.; Varona, S.; Zaballos, A. |
| EPI_ISL_467060, EPI_ISL_474833, EPI_ISL_474834, EPI_ISL_474841, EPI_ISL_474842, EPI_ISL_474845, EPI_ISL_474854, EPI_ISL_474855, EPI_ISL_474858, EPI_ISL_474860, EPI_ISL_474861, EPI_ISL_474865, EPI_ISL_474866, EPI_ISL_474870, EPI_ISL_474871, EPI_ISL_474877, EPI_ISL_474878, EPI_ISL_474880, EPI_ISL_474883, EPI_ISL_474884, EPI_ISL_474885, EPI_ISL_474897, EPI_ISL_474907, EPI_ISL_474909, EPI_ISL_474923, EPI_ISL_474924, EPI_ISL_474925, EPI_ISL_474926, EPI_ISL_474929, EPI_ISL_474931, EPI_ISL_474942, EPI_ISL_474957, EPI_ISL_510444, EPI_ISL_538676, EPI_ISL_538680, EPI_ISL_538682, EPI_ISL_538683, EPI_ISL_538685, EPI_ISL_538688 | | | |
| see above | Hospital Universitario Virgen de las Nieves de Granada-SAS | SeqCOVID-SPAIN consortium/IBV(CSIC) | Irene Pedrosa Corral; José M. Navarro-Marí and SeqCOVID-SPAIN consortium; Mercedes Pérez Ruiz; Sara Sanbonmatsu Gámez |
| EPI_ISL_419233 | Hospital Universitario de Canarias | Instituto de Salud Carlos III | Camarero, S.; Casas, I.; Castro, B.; Cuesta, I.; González-Esguevillas, M.; Iglesias-Caballero, M.; Jiménez, M.; Jiménez, P.; Juliá, M.; Molinero Calamita, M.; Monzón, S.; Pozo, F.; Varona, S.; Zaballos, A. |
| EPI_ISL_474918, EPI_ISL_537692, EPI_ISL_537693, EPI_ISL_537699, EPI_ISL_537701, EPI_ISL_537704, EPI_ISL_537710, EPI_ISL_537797, EPI_ISL_537803, EPI_ISL_537806 | | | |
| see above | Hospital Universitario de Gran Canaria Dr. Negrín | SeqCOVID-SPAIN consortium/IBV(CSIC) | Ana Bordes Benítez and SeqCOVID-SPAIN consortium; Francisco J. Chamizo López; M. Carmen Pérez González |
| EPI_ISL_455316, EPI_ISL_455317, EPI_ISL_455320, EPI_ISL_455321, EPI_ISL_455322 | Hospital Virgen de las Nieves | Instituto de Salud Carlos III | A. Monzón; F. Casas; I; I. Jiménez; Iglesias-Caballero; M. Camarero; M. Cuesta; M. González-Esguevillas; M. Molinero Calamita; M. Zaballos; P. Jiménez; S. Juliá; S. Pozo; S. Sanbonmatsu; S. Varona |
| EPI_ISL_455314 | Hospital Virgen del Rocio | Instituto de Salud Carlos III | A. Monzón; F. Casas; I; I. Jiménez; Iglesias-Caballero; J. Lepe; M. Camarero; M. Cuesta; M. González-Esguevillas; M. Molinero Calamita; M. Zaballos; P. Jiménez; S. Juliá; S. Pozo; S. Varona |
| EPI_ISL_414577, EPI_ISL_414578 | Hospital de Talca, Chile | Instituto de Salud Publica de Chile | Alejandra Acevedo; Andrés E. Castillo; Bárbara Parra; Carolina Tambley; Gabriel Leal; Gisselle Barra; Jaime Lagos; Javier Tognarelli; Jorge Fernández.; Loredana Arata; Patricia Bustos; Paz Tapia; Rodrigo Fasce; Soledad Ulloa; Winston Andrade |
| EPI_ISL_510253 | Hospital de la Santa Creu i Sant Pau. Servicio de Microbiología | SeqCOVID-SPAIN consortium/IBV(CSIC) | Elisenda Miró and SeqCOVID-SPAIN consortium; Ferran Navarro; Núria Rabella |
| EPI_ISL_498255, EPI_ISL_498258, EPI_ISL_498261 | Hospital for Tropical Diseases | COVID-19 Network Investigations (CONI) Alliance | Akanitt Jittmittraphap; Angkana Huang; Anthony R. Jones; Arporn Wangwiwatsin; Bhakbhoom Panthan; Chonticha Klungtong; Ekawat Pasomsub; Elizabeth Batty; Insee Sensorn; Janjira Thaipadungpanit; Khajohn Joonsalak; Kingkan Rakmanee; Krittikorn Kumpornsin; Namfon Kotanan; Nantarat Chantawat; Pornsawan Leaungwutiwong; Stefan Fernandez; Thanat Chookajorn; Theerarat Kochakarn; Treewat Watthanachockchai; Wasun Chantratita; Weena Janwitthayanan; Wudtichai Manasatienkij |
| EPI_ISL_434716, EPI_ISL_434717, EPI_ISL_434723, EPI_ISL_434724, EPI_ISL_434731, EPI_ISL_434732, EPI_ISL_434738, EPI_ISL_434740, EPI_ISL_434741, EPI_ISL_434758, EPI_ISL_434760, EPI_ISL_434766, EPI_ISL_434775, EPI_ISL_434777, EPI_ISL_434788, EPI_ISL_434791, EPI_ISL_434797, EPI_ISL_434801, EPI_ISL_434809, EPI_ISL_434810, EPI_ISL_434812, EPI_ISL_434813, EPI_ISL_434814, EPI_ISL_434815, EPI_ISL_434816, EPI_ISL_434818, EPI_ISL_434825, EPI_ISL_434826, EPI_ISL_434827, EPI_ISL_434828, EPI_ISL_434829, EPI_ISL_434831, EPI_ISL_434835, EPI_ISL_434836, EPI_ISL_434838, EPI_ISL_434840, EPI_ISL_434843, EPI_ISL_434844, EPI_ISL_434849, EPI_ISL_434854, EPI_ISL_434855, EPI_ISL_434862, EPI_ISL_434863, EPI_ISL_434867, EPI_ISL_434872, EPI_ISL_434873, EPI_ISL_434882, EPI_ISL_434883, EPI_ISL_434884, EPI_ISL_434886, EPI_ISL_434887, EPI_ISL_434891, EPI_ISL_434892, EPI_ISL_434896, EPI_ISL_434897, EPI_ISL_434902, EPI_ISL_434904, EPI_ISL_434905, EPI_ISL_434907, EPI_ISL_434913, EPI_ISL_434916, EPI_ISL_434923, EPI_ISL_434924, EPI_ISL_434925, EPI_ISL_434932, EPI_ISL_434934, EPI_ISL_434953, EPI_ISL_434961, EPI_ISL_434964, EPI_ISL_434967, EPI_ISL_434969, EPI_ISL_434971, EPI_ISL_434974, EPI_ISL_434978, EPI_ISL_434981, EPI_ISL_434983, EPI_ISL_434986, EPI_ISL_542587, EPI_ISL_542803 | | | |
| see above | Houston Methodist Hospital | Houston Methodist Hospital | Chia-Wei Chou; Concepcion C. Cantu; Daniel Boutz; David W. Bernard; Ghazaleh Eskandari; Hakon Jonsson; Heather Hendrickson; Hoang A. T. Nguyen; Hung-Che Kuo; Ilya J. Finkelstein; J. Hunter Long; James J. Davis; Jason S. McLellan; Jimmy Gollihar; Jule Goike; Kamyab Javanmardi; Kari Stefansson; Layne Pruitt; Marcus Nguyen; Matthew Ojeda Saavedra; Maulik Shukla; Muthiah Kumaraswami; Paul A. Christensen; Prasanti Yerramilli; Randall J. Olsen; S. Wesley Long; Sishir Subedi; and James M. Musser |
| EPI_ISL_454636, EPI_ISL_454638, EPI_ISL_468446 | Humboldt County Public Health Laboratory | Chan-Zuckerberg Biohub | CZB Cliahub Consortium |
| EPI_ISL_535983, EPI_ISL_536203, EPI_ISL_536225, EPI_ISL_1241283 | Hôpital Charles-LeMoyne | Laboratoire de santé publique du Québec | Guillaume Bourque; Ioannis Ragoussis; Jesse Shapiro; Mark Lathrop and Michel Roger on behalf of the CoVSeQ research group; Sandrine Moreira |
| EPI_ISL_535755 | Hôpital Maisonneuve-Rosemont | Laboratoire de santé publique du Québec | Guillaume Bourque; Ioannis Ragoussis; Jesse Shapiro; Mark Lathrop and Michel Roger; Sandrine Moreira |
| EPI_ISL_450317, EPI_ISL_535718, EPI_ISL_535786, EPI_ISL_536276 | Hôpital Pierre-Boucher | Laboratoire de santé publique du Québec | Guillaume Bourque; Ioannis Ragoussis; Jesse Shapiro; Mark Lathrop and Michel Roger; Mark Lathrop and Michel Roger on behalf of the CoVSeQ research group; Sandrine Moreira |
| EPI_ISL_536061, EPI_ISL_536083, EPI_ISL_536137, EPI_ISL_536139 | Hôpital Pierre-Le Gardeur | Laboratoire de santé publique du Québec | Guillaume Bourque; Ioannis Ragoussis; Jesse Shapiro; Mark Lathrop and Michel Roger on behalf of the CoVSeQ research group; Sandrine Moreira |
| EPI_ISL_536010 | Hôpital Sainte-Croix | Laboratoire de santé publique du Québec | Guillaume Bourque; Ioannis Ragoussis; Jesse Shapiro; Mark Lathrop and Michel Roger on behalf of the CoVSeQ research group; Sandrine Moreira |
| EPI_ISL_465686, EPI_ISL_535911, EPI_ISL_535993, EPI_ISL_536117, EPI_ISL_536363, EPI_ISL_536387 | Hôpital de Hull | Laboratoire de santé publique du Québec | Guillaume Bourque; Ioannis Ragoussis; Jesse Shapiro; Mark Lathrop and Michel Roger; Mark Lathrop and Michel Roger on behalf of the CoVSeQ research group; Sandrine Moreira |
| EPI_ISL_535885, EPI_ISL_536111, EPI_ISL_536112 | Hôpital de Maria | Laboratoire de santé publique du Québec | Guillaume Bourque; Ioannis Ragoussis; Jesse Shapiro; Mark Lathrop and Michel Roger on behalf of the CoVSeQ research group; Sandrine Moreira |
| EPI_ISL_535889, EPI_ISL_535941, | Hôpital de Saint-Eustache | Laboratoire de santé publique du Québec | Guillaume Bourque; Ioannis Ragoussis; Jesse Shapiro; Mark Lathrop and Michel Roger; Mark Lathrop and Michel Roger on behalf of the CoVSeQ research group; Sandrine Moreira |

| | | | |
|---|---|---|---|
| EPI_ISL_536049, EPI_ISL_536058 | | | |
| EPI_ISL_536026, EPI_ISL_536200, EPI_ISL_536259, EPI_ISL_536274 | Hôpital du Suroît | Laboratoire de santé publique du Québec | Guillaume Bourque; Ioannis Ragoussis; Jesse Shapiro; Mark Lathrop and Michel Roger on behalf of the CoVSeQ research group; Sandrine Moreira |
| EPI_ISL_535725, EPI_ISL_535758, EPI_ISL_535759 | Hôtel-Dieu de Lévis | Laboratoire de santé publique du Québec | Guillaume Bourque; Ioannis Ragoussis; Jesse Shapiro; Mark Lathrop and Michel Roger; Mark Lathrop and Michel Roger on behalf of the CoVSeQ research group; Sandrine Moreira |
| EPI_ISL_410045 | IL Department of Public Health Chicago Laboratory | Pathogen Discovery, Respiratory Viruses Branch, Division of Viral Diseases, Centers for Dieases Control and Prevention | Anna Uehara; Brett L. Whitaker; Brian Lynch; Clinton R. Paden; Janna' R. Murray; Jing Zhang; Krista Queen; Lijuan Wang; Senthil Kumar K. Sakthivel; Shifaq Kamili; Stephen Lindstrom; Susan I. Gerber; Suxiang Tong; Xiaoyan Lu; Yan Li; Ying Tao |
| EPI_ISL_424851 | IL Department of Public Health Chicago Laboratory | Pathogen Discovery, Respiratory Viruses Branch, Division of Viral Diseases, Centers for Disease Control and Prevention | Alison S. Laufer Halpin; Anna Uehara; Christopher A. Elkins; Clinton R. Paden; Haibin Wang; Jing Zhang; Krista Queen; Mary S. Keckler; Rachel Marine; Suxiang Tong; Yan Li; Ying Tao |
| EPI_ISL_1577817 | INHRR | Laboratorio de Virología Molecular | Aguilar M; Alarcon V; D Angelo P; Delgado M; Garzaro D; Jaspe RC; Loureiro CL; Pujol FH; Rangel HR; Rodriguez L; Zambrano JL |
| EPI_ISL_453835, EPI_ISL_453836 | INSA | Instituto Nacional de Saude (INSA) | Borges et al |
| EPI_ISL_479939, EPI_ISL_479940 | Ibaraki Prefectural Institute of Public Health | Pathogen Genomics Center, National Institute of Infectious Diseases | Hajime Kamiya; Keiko Goto; Kentaro Itokawa; Makoto Kuroda; Masanori Hashino; Motoi Suzuki; Rina Tanaka; Tsuyoshi Sekizuka |
| EPI_ISL_683662, EPI_ISL_683665, EPI_ISL_683666, EPI_ISL_683667, EPI_ISL_779720, EPI_ISL_779722, EPI_ISL_779739, EPI_ISL_779741, EPI_ISL_779742, EPI_ISL_779745, EPI_ISL_779746, EPI_ISL_779747, EPI_ISL_779759, EPI_ISL_779760, EPI_ISL_779772, EPI_ISL_855407 | | | |
| see above | IdISSC/Hospital Clínico San Carlos de Madrid | SeqCOVID-SPAIN consortium/IBV(CSIC) | Alberto Delgado-Iribarren; Esther Culebras Lopez; Esther Culebras López; Jorge Matias-Guiu; Luis Ortega Medina; Silvia Sánchez Ramón; Ulises Gómez-Pinedo and SeqCOVID-SPAIN consortium; Vicente Estrada Pérez |
| EPI_ISL_632849, EPI_ISL_632851, EPI_ISL_632874, EPI_ISL_632882, EPI_ISL_632886 | Idaho Bureau of Laboratories | Center for Global Health, University of New Mexico Health Sciences Center | Christopher Ball; Darrell Dinwiddie; Daryl Domman; Kurt Schwalm; Matthew Burns; Robert Voermans |
| EPI_ISL_2976249 | Idaho Bureau of Laboratories | Idaho Bureau of Laboratories | "R. Beukelman; Aimee Ceniseros; Christian Loera; Christopher Ball"; Matthew Charles Burns; Robert L. Voermans |
| EPI_ISL_848243, EPI_ISL_848244, EPI_ISL_848251, EPI_ISL_848274, EPI_ISL_848275, EPI_ISL_848277, EPI_ISL_848280, EPI_ISL_848443 | | | |
| see above | Illinois Department of Public Health | Gagnon Lab, Southern Illinois University | Keith Gagnon |
| EPI_ISL_413523 | Indian Council of Medical Research-National Institute of Virology | National Influenza Center, Indian Council of Medical Research-National Institute of Virology | Choudhary ML; Potdar V; Shete-Aich A; Yadav PD |
| EPI_ISL_413518, EPI_ISL_413519, EPI_ISL_413520, EPI_ISL_413521 | Infectious Disease Control Center, Center for Disease Control and Prevention of PLA | Infectious Disease Control Center, Center for Disease Control and Prevention of PLA | Li; Li, J.; Li, L.; Li, Z.; P. and Li, P.; Qiu, S.; Song, H. |
| EPI_ISL_962529, EPI_ISL_962531, EPI_ISL_962533, EPI_ISL_962534, EPI_ISL_962540, EPI_ISL_962541, EPI_ISL_962544, EPI_ISL_962545, EPI_ISL_962547, EPI_ISL_962548, EPI_ISL_962556, EPI_ISL_962560, EPI_ISL_962561, EPI_ISL_962562, EPI_ISL_962565, EPI_ISL_962566, EPI_ISL_962571, EPI_ISL_962573, EPI_ISL_962574, EPI_ISL_962576, EPI_ISL_962579, EPI_ISL_962581, EPI_ISL_962582, EPI_ISL_962869, EPI_ISL_962870, EPI_ISL_962871, EPI_ISL_962874, EPI_ISL_962875 | | | |
| see above | Infectious Disease Control and Prevention Institute | Infectious Disease Control and Prevention Institute | Bo Pang; Jianxing Wang; Julong Wu; Mingxiao Yao; Ti Liu; Xiaolin Jiang; Yan Li; Yujie He; Yuwei Zhang; Zengqiang Kou |
| EPI_ISL_2896270 | Infectious Disease Core Research, Abbott Diagnostics Division | Infectious Disease Core Research, Abbott Diagnostics Division | Berg; Cloherty; Faron, M.; Forberg, K.; Fox, A.; G.A.; G.S.; Goldstein, Y.; Jennings, C.; M.G.; Meyer; Mohaimani, A.; Mowermann, I.; Orf; T.V. |
| EPI_ISL_1440879 | Institut National d'Hygiène | Unité Mixte Internationale TransVIHMI (UMI 233 IRD – U1175 INSERM - Université de Montpellier) IRD (Institut de recherche pour le développement) | Abla A. KONOU; Adodo SADJI; Ahidjo AYOUBA; Akoélé SILIADIN; Alassane OURO-MEDELI; Amivi EHLAN; Améyo DORKENOO; Anoumou DAGNRA; Christelle BUTEL; Déléma MABA; Eric DELAPORTE; Issaka Maman; Kokou TEGUENI; Laetitia SERRANO; Martine PEETERS; Messanh DOUFFAN; Mireille PRINCE-DAVID; Mounerou SALOU; Sidonie A.M.KAGNISSODE; Sika DOSSIM; Wembo A. HALATOKO |
| EPI_ISL_1442993 | Institut National d'hygiène | "Unité Mixte Internationale TransVIHMI (UMI 233 IRD – U1175 INSERM - Université de Montpellier) IRD (Institut de recherche pour le développement)" | Abla A. KONOU; Adodo SADJI; Ahidjo AYOUBA; Akoélé SILIADIN; Alassane OURO-MEDELI; Amivi EHLAN; Améyo DORKENOO; Anoumou DAGNRA; Christelle BUTEL; Déléma MABA; Eric DELAPORTE; Issaka Maman; Kokou TEGUENI; Laetitia SERRANO; Martine PEETERS; Messanh DOUFFAN; Mireille PRINCE-DAVID; Mounerou SALOU; Sidonie A.M.KAGNISSODE; Sika DOSSIM; Wembo A. HALATOKO |
| EPI_ISL_1293351, EPI_ISL_1303031, EPI_ISL_1321604 | Institut National d'Hygiène (INH) | Unité Mixte Internationale TransVIHMI (UMI 233 IRD – U1175 INSERM - Université de Montpellier)IRD (Institut de recherche pour le développement) | Abla A. KONOU; Adodo SADJI; Ahidjo AYOUBA; Akoélé SILIADIN; Alassane OURO-MEDELI; Amivi EHLAN; Améyo DORKENOO; Anoumou DAGNRA; Christelle BUTEL; Déléma MABA; Eric DELAPORTE; Issaka Maman; Kokou TEGUENI; Laetitia SERRANO; Martine PEETERS; Messanh DOUFFAN; Mireille PRINCE-DAVID; Mounerou SALOU; Sidonie A.M.KAGNISSODE; Sika DOSSIM; Wembo A. HALATOKO |
| EPI_ISL_418216, EPI_ISL_420077, EPI_ISL_420078 | Institut Pasteur Dakar | Institut Pasteur de Dakar | Amadou Alpha Sall; Mamadou Diop; Moussa Moise Diagne; Ndongo Dia; Ousmane Faye |
| EPI_ISL_855187, EPI_ISL_855188, EPI_ISL_855193, EPI_ISL_855195, EPI_ISL_855200, EPI_ISL_855233, EPI_ISL_855237, EPI_ISL_855243, EPI_ISL_855282 | | | |
| see above | Institute for Computational Biomedicine, Weill Cornell Medicine | Institute for Computational Biomedicine, Weill Cornell Medicine | A.J.; A.M.; Afshinnekoo, E.; B.W.; Bezdan, D.; Bhattacharya, C.; Butler; C.D.; C.E.; Cong, L.; Craney, A.; Cushing, M.; D.J.; Danko, D.; F.J.; Foox, J.; Gawrys, J.; Gisladottir, U.; Hajirasouliha, I.; Horner; Iftner, A.; Iftner, T.; Imielinski, M.; Ivanov; L.F.; Langhorst; Levy, S.; Loda, M.; MacKay, M.; Mason; Meleshko, D.; Melnick; Meydan, C.; Mozsary, C.; N.A.; P.A.D.; Pohle, D.; R.E.; Ramlall, V.; Rennert, H.; Rosiene, J.; Ruggiero, P.; Ryon, K.; S.M.; Salvatore, M.; Schwartz; Sedlazeck; Shaiber, A.; Shemesh; Sierra, M.; Sipley, J.; Steel; Tanner, N.; Tatonetti, N.; Thierry-Mieg, D.; Thierry-Mieg, J.; Velu, P.; Westblade; Westover; Wu, S.; Xiang, J.; Xu, D.; Young, B.; Zietz, M. |
| EPI_ISL_455790 | Institute for Medical Research, Infectious Disease Research Centre, National Institutes of Health, Ministry of Health Malaysia | Malaysia Genome Institute | Azrin Ahmad; Enizza Kasim; Irni Suhayu Sapian; Jeyanthi Suppiah; Mohd Faizal Abu Bakar; Mohd Noor Mat Isa; Norazah Ahmad; Nurhezreen Md Iqbal; Ravindran Thayan; Shahrul Hisham Zainal Ariffin; Shamsidar Sopie; Siti Noraini Othman; Tahir Aris; Yusuf Muhammad Noor; Zarina Mohd Zawawi |
| EPI_ISL_527871 | Institute of Medical Biology, Chinese Academy of Medical Sciences and Peking Union Medical College | Institute of Medical Biology, Chinese Academy of Medical Sciences and Peking Union Medical College | Chen, H.; Fan, S.; G. and Li, Q.; Jiang; Liao, Y.; Liu, L.; Wang, L.; Xie, Z.; Xu, X.; Zheng, H.; Zhou, X. |
| EPI_ISL_2643912 | Institute of Microbiology and Immunology, Faculty of Medicine, University of Ljubljana | Institute of Microbiology and Immunology, Faculty of Medicine, University of Ljubljana | Alen Suljič; Andraž Celar; Dominika Šturm; Doroteja Vlaj; Mario Poljak; Matic Brvar; Miša Korva; Patricija Pozvek; Samo Zakotnik; Tatjana Avšič – Županc; Tomaž Mark Zorec; Špela Pleh |
| EPI_ISL_983519 | Institute of Virology, University Hospital, University of Bonn and German Center for Infection Research (DZIF), Bonn-Cologne, Bonn, Germany | Institute of Virology, University Hospital, University of Bonn and German Center for Infection Research (DZIF), Bonn-Cologne, Bonn, Germany | Marek Korencak et al |
| EPI_ISL_426364 | Instituto Nacional de Ciencias Medicas y Nutricion Salvador Zubiran | Instituto Nacional de Ciencias Medicas y Nutricion | Adnan Araiza Rodríguez; Alejandro Sánchez; Alfredo Ponce de León Garduño; Blanca Taboada; Carlos F. Arias; Carolina González Torres; Celia Boukadida; Cesar Raúl González Bonilla; Concepción Grajales Muñiz; Edgar Mendieta Condado; Eduardo Becerril Vargas; Fabiola Garcés Ayala; Fernando Ledesma Barrientos; Francisco Javier Gaytán Cervantes; Francisco Pulido; Gisela Barrera Badillo; Gloria Vázquez; Guillermo M. Ruiz-Palacios; Irma López Martínez; Joel Armando Vázquez Pérez; José Arturo Martínez Orozco; José Ernesto Ramírez González; José Esteban Muñoz Medina; Lucia Hernández Rivas; Luis Alberto García Andrade; Mario Mújica Sánchez; Pavel Isa; Pilar Ramos Cervantes; Ricardo Grande; Santiago Avila Ríos; Victor Hugo Borja Aburto; Violeta Ibarra Gonzalez |
| EPI_ISL_426361, EPI_ISL_426362, EPI_ISL_426363 | Instituto Nacional de Ciencias Medicas y Nutricion Salvador Zubiran | Instituto Nacional de Ciencias Medicas y Nutricion Salvador Zubiran | Adnan Araiza Rodríguez; Alejandro Sánchez; Alfredo Ponce de León Garduño; Blanca Taboada; Carlos F. Arias; Carolina González Torres; Celia Boukadida; Cesar Raúl González Bonilla; Concepción Grajales Muñiz; Edgar Mendieta Condado; Eduardo Becerril Vargas; Fabiola Garcés Ayala; Fernando Ledesma Barrientos; Francisco Javier Gaytán Cervantes; Francisco Pulido; Gisela Barrera Badillo; Gloria Vázquez; Guillermo M. Ruiz-Palacios; Irma López Martínez; Joel Armando Vázquez Pérez; José Arturo Martínez Orozco; José Ernesto Ramírez González; José Esteban Muñoz Medina; Lucia Hernández Rivas; Luis Alberto García Andrade; Mario Mújica Sánchez; Pavel Isa; Pilar Ramos Cervantes; Ricardo Grande; Santiago Avila Ríos; Victor Hugo Borja Aburto; Violeta Ibarra Gonzalez |
| EPI_ISL_424627 | Instituto Nacional de Enfermedades Respiratorias | Instituto Nacional de Enfermedades Respiratorias | Adnan Araiza Rodríguez; Alejandro Sánchez; Alfredo Ponce de León Garduño; Blanca Taboada; Carlos F. Arias.; Carolina González Torres; Celia Boukadida; Cesar Raúl González Bonilla; Concepción Grajales Muñiz; Edgar Mendieta Condado; Eduardo Becerril Vargas; Fabiola Garcés Ayala; Fernando Ledesma Barrientos; Francisco Javier Gaytán Cervantes; Francisco Pulido; Gisela Barrera Badillo; Gloria Vázquez; Guillermo M. Ruiz-Palacios; Irma López Martínez; Joel Armando Vázquez Pérez; Jorge Salas Hernández; José Arturo Martínez Orozco; José Ernesto Ramírez González; José Esteban Muñoz Medina; Lucia Hernández Rivas; Luis Alberto García Andrade; Mario Mújica Sánchez; Pavel Isa; Pilar Ramos Cervantes; Ricardo Grande; Santiago Avila Ríos; Victor Hugo Borja Aburto; Violeta Ibarra Gonzalez |
| EPI_ISL_837616 | Instituto Nacional de Enfermedades Respiratorias (INER) | Instituto Nacional de Enfermedades Respiratorias (INER) | Alejandra Hernández-Terán; Alma Rincón-Rubio; Celia Boukadida; Edgar Sevilla-Reyes; Eduardo Becerril-Vargas; Fidencio Mejía-Nepomuceno; Hector Esteban Paz-Juárez; Joel Armando Vázquez-Pérez; Jorge Salas-Hernández; José Arturo Martínez-Orozco; Margarita Matías-Florentino; Mario Mújica-Sánchez; Olivia Briceño; Santiago Ávila-Ríos |
| EPI_ISL_1301520, EPI_ISL_1301718 | Instituto Nacional de Enfermedades Respiratorias (INER) | Instituto de Biotecnología de la UNAM | Alejandra Hernández-Terán; Alejandro Sanchez-Flores; Alma Rincón-Rubio; Andrea Santos Coy-Arechavaleta; Authors from IBT; Blanca Taboada; Celia Boukadida; Clara Esperanza Santacruz-Tinoco; Edgar Mendieta-Condado; Eduardo Becerril-Vargas; Fidencio Mejía-Nepomuceno; Francisco Pulido; Gisela Barrera-Badillo; Gloria Vazquez; Hector Esteban Paz-Juárez; IMSS; InDRE and INER (in alphabetical order): Carlos F. Arias; Irma Lopez-Martinez; Jerome Jean Verleyen; Joel Armando Vázquez-Pérez; Jorge Salas-Hernández; José Arturo Martínez-Orozco; José Ernesto Ramírez-González; José Esteban Muñoz-Medina; Larissa Fernandes-Matano; Lucia Hernandez-Rivas; Luis Alberto Ochoa-Carrera; Margarita Matías-Florentino; Mario Mújica-Sánchez; Natividad Cruz-Ortiz; Pavel Isa; Ricardo Grande; Santiago Ávila-Ríos; Tatiana Nunez-Garcia; Teresita Rojas-Mendoza |
| EPI_ISL_536537, EPI_ISL_536539, EPI_ISL_536540, EPI_ISL_536541, EPI_ISL_536556, EPI_ISL_536561 | Instituto Nacional de Salud | Laboratorio de Infecciones Respiratorias Agudas | David Tarazona; Dennis Carhuaricra; Eduardo Juscamayta Lopez; Faviola Valdivia Guerrero; Lenin Maturrano Hernandez; Nancy Rojas Serrano; Ronnie Gavilan Chavez |
| EPI_ISL_456117, EPI_ISL_456119, EPI_ISL_456123 | Instituto Nacional de Salud - Unidad de Secuenciación y Análisis Genómico | Instituto Nacional de Salud, Universidad Cooperativa de Colombia, Instituto Alexander von Humboldt, Imperial College-London, London School of Hygiene & Tropical Medicine | Astrid C. Flórez; Carlos Franco-Muñoz; Christian Julian Villabona-Arenas; Diana Marcela Walteros-Acero; Diego A. Álvarez-Díaz; Erika Ospitia; Gloria Puerto; Jose A. Usme-Ciro; Juliana Barbosa; Katherine Laiton-Donato; Liz Villabona-Arenas; Luz Dary Rodriguez; Mailyn A.Gonzalez; Marcela Mercado-Reyes.; Martha Lucia Ospina Martinez; Nicolas D. Franco-Sierra; Sergio Gomez-Rangel; Sussy Echeverria; Zulma M. Cucunubá |
| EPI_ISL_447757 | Instituto Nacional de Salud, Bogotá, Colombia | Grupo de Investigaciones Microbiológicas-UR (GIMUR), Departamento de Biología, Facultad de Ciencias Naturales, Universidad del Rosario, | Adriana Castillo; Alberto Paniz-Mondolfi; Ana S. Gonzalez-Reiche; Angelica Rico; Aníbal A. Teherán; Carolina Florez; Carolina Hernandez; David Martínez; Emilia Mia Sordillo; Esther C. Barros; Harm van Bakel; Jesús E. Jaimes; Juan David Ramírez; Laura Vega; Lisseth Pardo; Marina Muñoz; Martha L. Ospina; Matthew M. Hernandez; Nathalia Ballesteros; Sergio Castañeda; Sergio Gomez; Viviana Simon |

| | | | |
|---|---|---|---|
| | | Bogotá, Colombia Instituto Nacional de Salud, Bogotá, Colombia Icahn School of Medicine at Mount Sinai, New York, USA | |
| EPI_ISL_498162, EPI_ISL_498169, EPI_ISL_498170 | Instituto Nacional de Salud, Bogotá, Colombia | Instituto Nacional de Salud, Bogotá, Colombia | Astrid C. Flórez; Carlos Andrés Durán; Carlos Franco-Muñoz; Carolina Ferro; Christian Julian Villabona-Arenas; Diana Marcela Walteros-Acero; Diego A. Álvarez-Díaz; Diego Andrés Prada; Franklin Prieto; Jonathan Reales; Jose A. Usme-Ciro; Katherine Laiton-Donato; Liz Villabona-Arenas; Marcela Mercado-Reyes; Martha Lucia Ospina Martinez; Nicolas D. Franco-Sierra; Sussy Echeverria; Zulma M. Cucunubá |
| EPI_ISL_424673 | Instituto de Diagnostico y Referencia Epidemiologicos | Instituto de Diagnostico y Referencia Epidemiologicos | Adnan Araiza Rodríguez; Alejandro Sánchez; Alfredo Ponce de León Garduño; Blanca Taboada; Carlos F. Arias; Carolina González Torres; Celia Boukadida; Cesar Raúl González Bonilla; Concepción Grajales Muñiz; Edgar Mendieta Condado; Eduardo Becerril Vargas; Fabiola Garcés Ayala; Fernando Ledesma Barrientos; Francisco Javier Gaytán Cervantes; Francisco Pulido; Gisela Barrera Badillo; Gloria Vázquez; Guillermo M. Ruiz-Palacios; Irma López Martínez; Joel Armando Vázquez Pérez; José Arturo Martínez Orozco; José Ernesto Ramírez González; José Esteban Muñoz Medina; Lucia Hernández Rivas; Luis Alberto García Andrade; Mario Mújica Sánchez; Pavel Isa; Pilar Ramos Cervantes; Ricardo Grande; Santiago Avila Ríos; Victor Hugo Borja Aburto; Violeta Ibarra Gonzalez |
| EPI_ISL_455432, EPI_ISL_455439, EPI_ISL_455456 | Instituto de Diagnostico y Referencia Epidemiologicos (INDRE) | Instituto de Diagnostico y Referencia Epidemiologicos (INDRE) | ; Araiza-Rodriguez Adnan; Barrera-Badillo Gisela; Garces-Ayala Fabiola; Garces-Ayala Fabiola. Taboada Ramírez Blanca. Ramirez-Gonzalez Ernesto; Hernandez-Rivas Lucia; Lopez-Martinez Irma; Lopez-Martinez Irma.; Mendieta-Condado Edgar; Ramirez-Gonzalez Ernesto.; Rodriguez-Maldonado Abril; Taboada Ramirez Blanca; Taboada Ramírez Blanca. Ramirez-Gonzalez Ernesto; Wong-Arambula Claudia; Wong-Arambula Claudia. |
| EPI_ISL_956391 | Isolation - Virology Unit, Institut Pasteur du Cambodge; Sequencing - US National Institute of Allergy and Infectious Diseases Cambodia, US Naval Medical Research Unit -2, Cambodia National Institute for Public Health | Virology Unit, Institut Pasteur du Cambodge | Chau Darapheak; Chin Savuth; Erik A Karlsson; Jennifer Bohl; Jessica Manning; Jose A Garcia-Rivera; Kraing Sidonn; Ly Sovann; Sophana Chea; Sreyngim Lay; Veasna Duong; Vireak Heang; Yi Sengdoeurn |
| EPI_ISL_483708, EPI_ISL_514308, EPI_ISL_649060, EPI_ISL_649068, EPI_ISL_649076, EPI_ISL_649087, EPI_ISL_649088, EPI_ISL_649091, EPI_ISL_649092, EPI_ISL_649095, EPI_ISL_649096, EPI_ISL_649107 | | | |
| see above | Israel Central Virology laboratory | Israel Central Virology laboratory | Efrat Dahan Bucris; Ella Mendelson; Michal Mandelboim; Neta Zuckerman; Oran Erster |
| EPI_ISL_631790, EPI_ISL_631791, EPI_ISL_631970, EPI_ISL_632008, EPI_ISL_632009, EPI_ISL_632172 | Jamaica Hospital Medical Center | New York City Public Health Laboratory | Jade Wang; et al. |
| EPI_ISL_421237, EPI_ISL_421238, EPI_ISL_421239, EPI_ISL_421240, EPI_ISL_421241, EPI_ISL_421242, EPI_ISL_421243, EPI_ISL_421244, EPI_ISL_421245, EPI_ISL_421246, EPI_ISL_421247, EPI_ISL_421248, EPI_ISL_421249, EPI_ISL_455460, EPI_ISL_455461, EPI_ISL_455462, EPI_ISL_455464, EPI_ISL_455467 | | | |
| see above | Jiangxi Province Center for Disease Control and Prevention | Jiangxi Province Center for Disease Control and Prevention | DaJin Xiao; Fang Xiao; Gang Xu; JianXiong Li; Jun Zhou; ShiWen Liu; Tian Gong; XiaoQing Liu; Xin Ran; YanNi Zhang; Ying Xiong; Yong Shi |
| EPI_ISL_434686, EPI_ISL_438234, EPI_ISL_438241, EPI_ISL_438246, EPI_ISL_457756, EPI_ISL_457762, EPI_ISL_457767, EPI_ISL_457768, EPI_ISL_457773, EPI_ISL_457779, EPI_ISL_457782, EPI_ISL_457783, EPI_ISL_457789, EPI_ISL_457800, EPI_ISL_457807, EPI_ISL_457813, EPI_ISL_511866, EPI_ISL_511867, EPI_ISL_2884922, EPI_ISL_2884925, EPI_ISL_2884926, EPI_ISL_2884927, EPI_ISL_2884938, EPI_ISL_2884949, EPI_ISL_2884998 | | | |
| see above | Johns Hopkins Hospital Department of Pathology | Johns Hopkins Hospital Department of Pathology | Adannaya Amadi; Amanda Ernlund; C. Paul Morris; Chun Huai Luo; Craig Howser; Heba H. Mostafa; Heba Mostafa; Kristina Zudock; Mark Hopkins; Matthew Schwartz; Melanie Kirsche; Michael C. Schatz; Michael Schatz; N_di_ Trovao; Nicholas Gallagher; Nidia Trovao; Norah Sadowski; Nídiá Trovao; Oluwaseun Falade-Nwulia; Paul Morris; Peter M. Thielen; Shirlee Wohl; Srividya Ramakrishnan; Stuart C. Ray; Thomas Mehoke; Timothy Gilpatrick; Victoria Gniazdowski; Winston Timp; Yunfan Fan; _Amanda Ernlund; _Oluwaseun Falade-Nwulia |
| EPI_ISL_407976 | KU Leuven, Clinical and Epidemiological Virology | KU Leuven, Clinical and Epidemiological Virology | Annabel Rector; Bert Vanmechelen; Elke Wollants; Els Keyaerts; Lies Laenen; Marc Van Ranst; and Piet Maes |
| EPI_ISL_462264 | KU Leuven, Rega Institute, Clinical and Epidemiological Virology | KU Leuven, Rega Institute, Clinical and Epidemiological Virology | Bert Vanmechelen; Joan Marti-Carerras; Piet Maes; Tony Wawina-Bokalanga |
| EPI_ISL_690824 | Kanagawa Prefectural Institute of Public Health | Pathogen Genomics Center, National Institute of Infectious Diseases | Kentaro Itokawa; Makoto Kuroda; Masanori Hashino; Rina Tanaka; Tsuyoshi Sekizuka |
| EPI_ISL_875679 | Kansas Health and Environmental Lab | Kansas Health and Environmental Lab | Ben Olsen; Carissa Robertson; Mike Grose; and Phil Adam |
| EPI_ISL_965817, EPI_ISL_965818, EPI_ISL_965821 | Kentucky State Public Health Lab | Kentucky State Public Health Lab | Joshua Tobias; Karim George; Matthew Johnson; Rachel Zinner; Rhonda Lucas; Stephanie Lunn; Vaneet Arora; William Grooms |
| EPI_ISL_483553 | Kingdom of Bahrein Ministry of Health | Erasmus Medical Center | Amjad Ghanem Mohamed; Anne van der Linden; Bas Oude Munnink; Claudia Schapendonk; David Nieuwenhuijse; Ebrahim Shehad; Fatema; Hashmeya Al Wasti; Irina Chestakova; Marion Koopmans; Mark Pronk; Pascal Lexmond; Reina Sikkema; Richard Molenkamp; Stefan van Nieuwkoop; Theo Bestebroer; on behalf of the Dutch national COVID-19 response team. |
| EPI_ISL_459878, EPI_ISL_459879 | Kingston Health Sciences Center | Queen's Genomics Lab at Ongwanada (Q-GLO) | Ayub M; Colautti R; Evans GA; Guan H; Huang D; Hudson ML; Perez-Patrigeon S; Rustom N; Sheth P; Sjaarda CP; Soares CN; Wong H |
| EPI_ISL_479823 | Kochi Prefectural Institute of Public Health | Pathogen Genomics Center, National Institute of Infectious Diseases | Akihiko Tokaji; Hajime Kamiya; Kentaro Itokawa; Makoto Kuroda; Masanori Hashino; Motoi Suzuki; Rina Tanaka; Tsuyoshi Sekizuka |
| EPI_ISL_407193 | Korea Centers for Disease Control & Prevention (KCDC) Center for Laboratory Control of Infectious Diseases Division of Viral Diseases | Korea Centers for Disease Control & Prevention (KCDC) Center for Laboratory Control of Infectious Diseases Division of Viral Diseases | Heui Man Kim; Hye-Joon Jo; Jeong-Min Kim; Mi-Seon Kim; Myung Guk Han; Namjoo Lee; SangHee Woo; Sehee Park; Yoon-Seok Chung |
| EPI_ISL_480103, EPI_ISL_480104, EPI_ISL_480105 | Koshigaya City Public Health Center | Pathogen Genomics Center, National Institute of Infectious Diseases | Aya Tamura; Hajime Kamiya; Kentaro Itokawa; Kyohei Sakata; Makoto Kuroda; Masanori Hashino; Motoi Suzuki; Rina Tanaka; Takumi Daimon; Tsuyoshi Sekizuka; Yoko Togawa; Yoshiko Hamada; Yuka Furui |
| EPI_ISL_515181, EPI_ISL_515182, EPI_ISL_515183, EPI_ISL_515184, EPI_ISL_515248 | Kumasi Centre for Collaborative Research in Tropical Medicine, Kumasi. | Institute of Virology, Charité – Universitätsmedizin Berlin | Augustina Sylverken; Christian Drosten; Eric Adu; Jesse Addo Asamoah; Julia Schneider; Jörn Beheim-Schwarzbach; Michael Owusu; Philip El-Duah; Richard Phillips.; Richmond Gorman; Richmond Yeboah; Sherihane Aryeetey; Victor Max Corman |
| EPI_ISL_528539 | LVM/UFRJ | LNCC | Amilcar Tanuri; Ana Teresa R. Vasconcelos; Bruno B. Bezerra; Diana Marianni; Elena Cobos; Fabio Limonte; Gustavo M. Romário M. de Souza; Isadora A. Correa; Luciana B. Arruda; Luciana J. Costa.; Lucio A. Caldas; Luiz Almeida; Luiza Higga; Marcelo Bozza; Orlando Ferreira; Sharton V. A. Coelho; Terezinha M. Castineiras; Wanderley de Souza |
| EPI_ISL_434364 | Lab voor klinische biologie | Onderzoeksgroep Virologie | Bruno Verhasselt; Hans Nauwynck; Jozefien De Clercq; Laurens Lambrechts; Linos Vandekerckhove; Marthe Pauwels; Nick Vereecke; Sebastiaan Theuns |
| EPI_ISL_456172, EPI_ISL_456174, EPI_ISL_456180, EPI_ISL_456200, EPI_ISL_456212, EPI_ISL_579120 | LabPLUS | Institute of Environmental Science and Research (ESR) | Anja Werno; Antje van der Linden; Arlo Upton; Chris Mansell; David Hammer; Dragana Drinkovic; Erasmus Smit; Gary McAuliffe; Hana Sofia Andersson; Hermes Perez; James Ussher; Jill Sherwood; Jing Wang; Joep de Ligt; Josh Freeman; Julia Howard; Juliet Elvy; Lauren Jelly; Mary DeAlmeida; Matt Blakiston; Matt Storey; Matthew Rogers; Max Bloomfield; Michael Addidle; Michelle Balm; Muhammad Faisal; Nikki Freed; Olin Silander; Sally Roberts; Sarah Jefferies; Sharmini Muttaiyah; Susan Morpeth; Susan Taylor; Timothy Blackmore; Vani Sathyendran; Veronica Playle; Virginia Hope; Xiaoyun Ren |
| EPI_ISL_421735, EPI_ISL_429738, EPI_ISL_429743, EPI_ISL_429774 | Laboratoire National de Sante, Microbiology, Virology | Laboratoire National de Sante, Microbiology, Epidemiology and Microbial Genomics | Anke Wienecke-Baldacchino; Ardashel Latsuzbaia; Catherine Ragimbeau; Guillaume Fournier; Jessica Tapp; Joel Mossong; Tamir Abdelrahman; Trung Nguyen Nguyen |
| EPI_ISL_414600, EPI_ISL_414623 | Laboratoire de Virologie Institut de Virologie - INSERM U 1109 Hôpitaux Universitaires de Strasbourg | National Reference Center for Viruses of Respiratory Infections, Institut Pasteur, Paris | Angela Brisebarre; Flora Donati Vincent Enouf; Marion Barbet; Maud Vanpeene; Méline Bizard; Mélnie Albert; Samira Fafi-Kremer; Sylvie Behillil; Sylvie van der Werf |
| EPI_ISL_889814, EPI_ISL_889817, EPI_ISL_889841, EPI_ISL_889867, EPI_ISL_889911, EPI_ISL_889963, EPI_ISL_889970, EPI_ISL_890000, EPI_ISL_890001, EPI_ISL_890006, EPI_ISL_890023, EPI_ISL_890391, EPI_ISL_890400, EPI_ISL_890401, EPI_ISL_890406, EPI_ISL_890407, EPI_ISL_890426, EPI_ISL_890436, EPI_ISL_890445, EPI_ISL_890447, EPI_ISL_890448, EPI_ISL_890452, EPI_ISL_890472, EPI_ISL_890475, EPI_ISL_890485, EPI_ISL_890486, EPI_ISL_890487, EPI_ISL_890488, EPI_ISL_890495, EPI_ISL_890523, EPI_ISL_890535, EPI_ISL_890545, EPI_ISL_890588, EPI_ISL_890603, EPI_ISL_890621, EPI_ISL_890624, EPI_ISL_890661, EPI_ISL_890671, EPI_ISL_890680, EPI_ISL_890704, EPI_ISL_890721, EPI_ISL_890740, EPI_ISL_890747, EPI_ISL_890758, EPI_ISL_890766, EPI_ISL_890778, EPI_ISL_890798, EPI_ISL_890799, EPI_ISL_890815, EPI_ISL_890822, EPI_ISL_890830, EPI_ISL_890875, EPI_ISL_890880, EPI_ISL_890886, EPI_ISL_1241112, EPI_ISL_1241114, EPI_ISL_1241115, EPI_ISL_1241120, EPI_ISL_1241130, EPI_ISL_1241150, EPI_ISL_1241165, EPI_ISL_1241169, EPI_ISL_1241188, EPI_ISL_1241189, EPI_ISL_1241193, EPI_ISL_1241196, EPI_ISL_1365839, EPI_ISL_1378971, EPI_ISL_1390778, EPI_ISL_1390789 | | | |
| see above | Laboratoire de santé publique du Québec | Laboratoire de santé publique du Québec | Guillaume Bourque; Ioannis Ragoussis; Jesse Shapiro; Mark Lathrop and Michel Roger on behalf of the CoVSeQ research group; Mark Lathrop and Michel Roger on behalf of the CoVSeQ research group (http://covseq.ca/researchgroup); Sandrine Moreira |
| EPI_ISL_476833 | Laboratoire des Fièvres Hémorragiques Virales du Benin | Charité-Universitätsmedizin Berlin | Anges; Drexler; Jan Felix; Moreira-Soto Andres; Sander Anna-Lena; Yadouleton |
| EPI_ISL_792476 | Laboratorio Central Mg. Luis Alfredo Pianciola | Hospital Regional Ushuaia - Centro Austral De Investigaciones Científicas - Universidad Nacional De Tierra Del Fuego on behalf of 'Proyecto Argentino Interinstitucional de genomica de SARS-CoV-2' (PAIS Consortium) | C; CF; Ceballos; F; Gallego; Gramundi; ID; M; MC; Mazzeo; Nardi; Pianciola, L.; Pinto; SG; Ziehm |
| EPI_ISL_3048752 | Laboratorio Central de Saude Publica do Estado do Rio Grande do Sul (LACEN-RS) | Laboratório de Biologia Molecular da Universidade Federal de Ciências da Saúde de Porto Alegre | Adriana Seixas; Ana B. G. Veiga; Ana Paula Muterle Varela; Fabiana Quoos Mayer; Fernando Hayashi Sant'Anna; Janira Prichula; Letícia Garay Martins; Richard Steiner Salvato; Tatiana Schäffer Gregianini |
| EPI_ISL_424667 | Laboratorio Estatal de Salud Publica del Estado de México | Instituto de Diagnóstico y Referencia Epidemiológicos | Adnan Araiza Rodríguez; Alejandro Sánchez; Alfredo Ponce de León Garduño; Blanca Taboada; Carlos F. Arias.; Carolina González Torres; Celia Boukadida; Cesar Raúl González Bonilla; Concepción Grajales Muñiz; Edgar Mendieta Condado; Eduardo Becerril Vargas; Fabiola Garcés Ayala; Fernando Ledesma Barrientos; Francisco Javier Gaytán Cervantes; Francisco Pulido; Gisela Barrera Badillo; Gloria Vázquez; Guillermo M. Ruiz-Palacios; Irma López Martínez; Joel Armando Vázquez Pérez; José Arturo Martínez Orozco; José Ernesto Ramírez González; José Esteban Muñoz Medina; Lucia Hernández Rivas; Luis Alberto García Andrade; Mario Mújica Sánchez; Pavel Isa; Pilar Ramos Cervantes; Ricardo Grande; Santiago Avila Ríos; Victor Hugo Borja Aburto; Violeta Ibarra Gonzalez |
| EPI_ISL_424670 | Laboratorio Estatal de Salud Publica del Estado de Queretaro | Instituto de Diagnóstico y Referencia Epidemiologicos | Adnan Araiza Rodríguez; Alejandro Sánchez; Alfredo Ponce de León Garduño; Blanca Taboada; Carlos F. Arias; Carolina González Torres; Celia Boukadida; Cesar Raúl González Bonilla; Concepción Grajales Muñiz; Edgar Mendieta Condado; Eduardo Becerril Vargas; Fabiola Garcés Ayala; Fernando Ledesma Barrientos; Francisco Javier Gaytán Cervantes; Francisco Pulido; Gisela Barrera Badillo; Gloria Vázquez; Guillermo M. Ruiz-Palacios; Irma López Martínez; Joel Armando Vázquez Pérez; José Arturo Martínez Orozco; José Ernesto Ramírez González; José Esteban Muñoz Medina; Lucia Hernández Rivas; Luis Alberto García Andrade; Mario Mújica Sánchez; Pavel Isa; Pilar Ramos Cervantes; Ricardo Grande; Santiago Avila Ríos; Victor Hugo Borja Aburto; Violeta Ibarra Gonzalez |
| EPI_ISL_457940, EPI_ISL_457941, EPI_ISL_457942, EPI_ISL_457943, EPI_ISL_457945, EPI_ISL_457946, EPI_ISL_457947, EPI_ISL_457948, EPI_ISL_457950, EPI_ISL_457951, EPI_ISL_457952, EPI_ISL_457956, EPI_ISL_457958, EPI_ISL_457960, EPI_ISL_480428 | | | |
| see above | Laboratorio de Biología Molecular Asociación Española Primera en Salud | Departments of Pathology and Medicine, New York University School of Medicine | Adriana Heguy; Christian Marier; Gael Westby; Gonzalo Manrique; Maria Noel Zubillaga; Maria Victoria Elizondo; Matthew T Maurano; Paul Zappile |
| EPI_ISL_417034 | Laboratorio de Ecologia de Doencas Transmissiveis na | Laboratorio de Ecologia de Doencas Transmissiveis na Amazonia, | André Corado; Debora Duarte; Felipe Naveca on behalf of the Fiocruz COVID-19 Genomic Surveillance Network; Fernanda Nascimento; Luciana Gonçalves; Michele Jesus; Sérgio Luz; Valdinete Nascimento; Ágatha Costa |

| | | | |
|---|---|---|---|
| | Amazonia, Instituto Leonidas e Maria Deane - Fiocruz Amazonia | Instituto Leonidas e Maria Deane - Fiocruz Amazonia | |
| EPI_ISL_648380 | Laboratorio de Infectología Servicio de Infectología Hospital Universitario Dr. José Eleuterio González - Universidad Autónoma de Nuevo León | Laboratorio de Infectología Molecular Departamento de Bioquímica y Medicina Molecular Facultad de Medicina - Universidad Autónoma de Nuevo León | Adrian Camacho-Ortiz; Ana M. Rivas-Estilla; Consuelo Treviño-Garza; Daniel Arellanos-Soto; Eduardo Perez-Alba; Elvira Garza-González; Kame A. Galán-Huerta; Laura Nuzzolo-Shihadeh; Manuel E. de-la-O-Cavazos; María F. Herrera-Saldivar; Natalia Martínez-Acuña; Paola Bocanegra-Ibarias; Roberto Montes-de-Oca; Samantha M. Flores-Treviño; Sonia A. Lozano-Sepúlveda |
| EPI_ISL_779169 | Laboratorio de Infectología, Servicio de Infectología, Hospital Universitario Dr. José Eleuterio González - Universidad Autónoma de Nuevo León | Laboratorio de Infectología Molecular, Departamento de Bioquímica y Medicina Molecular,Facultad de Medicina - Universidad Autónoma de Nuevo León | Adrian Camacho-Ortiz; Ana M. Rivas-Estilla; Daniel Arellanos-Soto; Eduardo Perez-Alba; Elvira Garza-González; Kame A. Galán-Huerta; Laura Nuzzolo-Shihadeh; María F. Herrera-Saldivar; Natalia Martínez-Acuña; Paola Bocanegra-Ibarias; Samantha M. Flores-Treviño; Sonia A. Lozano-Sepúlveda |
| EPI_ISL_648318 | Laboratorio de Investigaciones de Baney | University Hospital Basel, Clinical Bacteriology | Adrian Egli; Alfredo Mari; Bonifacio Manguire Nlavo; Carlos Cortes; Claudia Daubenberger; Diosdado Odjama Nseng Ada; Elizabeth Nyakarungu; Guillermo Garcia; Helena Seth-Smith; Madlen Stange; Maximilian Mpina; Mitoha Ondo O Ayekaba; Philip Wonder Phiri; Salome Hosch; Tim Roloff; Tobias Schindler |
| EPI_ISL_523810, EPI_ISL_525206 | Laboratorio de Referencia Nacional de Virus Respiratorio. Centro Nacional de Salud Publica. Instituto Nacional de Salud Peru. | Laboratorio de Referencia Nacional de Biotecnología y Biología Molecular. Centro Nacional de Salud Publica. Instituto Nacional de Salud Peru. | Carlos Padilla Rojas; Henri Bailon Calderon; Johanna Balbuena Torres; Karolyn Vega Chozo; Marco Galarza Perez; Maribel Huaringa Nuñez; Nancy Rojas Serrano.; Omar Caceres Rey; Priscila Lope Pari |
| EPI_ISL_489988, EPI_ISL_491427, EPI_ISL_491429, EPI_ISL_491430, EPI_ISL_491464 | Laboratorio de Referencia Nacional de Virus Respiratorio. Instituto Nacional de Salud Perú | Laboratorio de Referencia Nacional de Biotecnología y Biología Molecular. Instituto Nacional de Salud Perú | Carlos Padilla Rojas; Henri Bailon Calderon; Johanna Balbuena Torres; Johanna Balbuena Torrez; Karolyn Chozo Vega; Karolyn Vega Chozo; Marco Galarza Perez; Maribel Huaringa Nuñez; Nancy Rojas Serrano; Nancy Rojas Serrano.; Omar Caceres Rey; Priscila Lope Pari |
| EPI_ISL_1111164, EPI_ISL_1111214, EPI_ISL_1111218, EPI_ISL_1111223, EPI_ISL_1111225, EPI_ISL_1111231, EPI_ISL_1111232 | | | |
| see above | Laboratorio de Referencia Nacional de Virus Respiratorio. Instituto Nacional de Salud Perú | Laboratorio de Referencia Nacional de Enteropatógenos. Instituto Nacional de Salud del Perú | Fiorella Orellana Peralta; Iris Silva Molina; Junior Caro Castro; Ronnie Gavilan Chavez; Veronica Hurtado Vela; Willi Quino Sifuentes |
| EPI_ISL_417518 | Laboratory Medicine | Department of Laboratory Medicine, Lin-Kou Chang Gung Memorial Hospital, Taoyuan, Taiwan | Cheng-Hsun Chiu; Cheng-Ta Yang; Chung-Guei Huang; Guang-Wu Chen; Kuo-Chien Tsao; Kuo-Ming Lee; Mei-Jen Hsiao; Peng-Nien Huang; Po-Wei Huang; Shin-Ru Shih; Shu-Li Yang; Yi-Chun Liu; Yu-Nong Gong |
| EPI_ISL_529208, EPI_ISL_529209 | Laboratory Medicine, University of Washington | University of Washington, Laboratory Medicine | Greninger, A.; Jerome, K.; Roychoudhury, P. |
| EPI_ISL_437625 | Laboratory of Genomics & Bioinformatics, Institute of Immunology and Experimental Therapy, Polish Academy of Sciences Oddzial Mikrobiologii Wojewodzkiej Stacji Sanitarno-Epidemiologicznej. | Laboratory of Genomics & Bioinformatics, Institute of Immunology and Experimental Therapy, Polish Academy of Sciences | Aleksandra Herud; Dariusz Martynowski; Dorota Kujawa; Grazyna Zalewska; Joanna Sikorska; Krzysztof Jakub Pawlik; Oskar Karpinski and Lukasz Laczmanski; Paulina Zebrowska |
| EPI_ISL_418265 | Laboratory of Microbiology, Department of Medicine, National and Kapodistrian University of Athens, Greece | Laboratory of Biology, Department of Medicine, Democritus University of Thrace, Greece | Elisavet Gatzidou; Ioannis Karakasiliotis; Maria Bampali; Nikolaos Dovrolis; Nikolaos Spanakis; Stavroula Veletza |
| EPI_ISL_434456, EPI_ISL_434464, EPI_ISL_437907 | Laboratory of Microbiology, Medical School, National and Kapodistrian University of Athens | Laboratory of Biology, Department of Medicine, Democritus University of Thrace | Bampali, M.; Dovrolis, N.; Froukala, E.; Gatzidou, E.; Kassela K.; N. and Karakasiliotis, I.; Spanakis; Stavropoulou, A.; Tsakris, A.; Veletza, S. |
| EPI_ISL_415658, EPI_ISL_415660, EPI_ISL_415661, EPI_ISL_801607 | Laboratory of Molecular Virology, Pontificia Universidad Católica de Chile | MSHS Pathogen Surveillance Program | Adolfo Garcia-Sastre; Adriana van De Guchte; Ajay Obla; Ana Maria Contreras; Ana S. Gonzalez-Reiche; Ana Silvia Gonzalez-Reiche; Bremy Alburquerque; Carlos Palma; Constanza Maldonado; Edward C. Holmes; Eileen Serrano; Erick Salinas; Hala Alshammary; Harm van Bakel; Jayeeta Dutta; Jorge Levican; Juan Soto; Leonardo I. Almonacid; M. Belen Leyton; Marcela Ferres; Matthew Hernandez; Matthew M. Hernandez; Melissa Smith; Mitchell Sullivan; Pablo Vial; Rafael A. Medina; Rafael A. Medina.; Robert Sebra; Shwetha Hara Sridhar; Shwetha Sridhar Hara; Tamara Garcia; Tamara García-Salum; Viviana Simon; Ying-Chih Wang; Zenab Khan |
| EPI_ISL_427299 | Laboratory of Respiratory Viruses and Measles, Oswaldo Cruz Institute, FIOCRUZ | Laboratory of Respiratory Viruses and Measles, Oswaldo Cruz Institute, FIOCRUZ | Aline Mattos; Braulia Caetano; Cristiana Garcia; Fernando Motta; Jonathan Lopes; Luciana Appolinario; Maria Ogrzewalska; Marilda Siqueira on behalf of the Fiocruz COVID-19 Genomic Surveillance Network; Milene Miranda; Paola Resende; Priscila Born; Sunando Roy |
| EPI_ISL_2158598 | Laboratório de Biologia Molecular Jean Piaget | MRCG at LSHTM, Genomics lab | Abdoulie Kanteh; Abdul Karim Sesay; Adul Candé; Aicha Balde; Aladje Balde; Bakary Sanyang; Bubacar Delgado Pinto Embaló; Dabiri Damilari; Edmira Maria da Costa; Erica Luis Maria Magalhães; Faatu Cassama; Mariama Kujabi; Milanca Agostinho Cá; Paulina Joãozinho da Costa Jarra Manneh; Rei José Pereira; Rui Inndi; Sainabou Laye Ndure; Simão Tchuda Bióté |
| EPI_ISL_451489, EPI_ISL_451611, EPI_ISL_455041 | Laverty Pathology | NSW Health Pathology - Institute of Clinical Pathology and Medical Research; Westmead Hospital; University of Sydney | CIDM-PH et al. |
| EPI_ISL_434636 | Lednicky Laboratory, Emerging Pathogens Institute, University of Florida | Lednicky Laboratory at Emerging Pathogens Institute, University of Florida | Alam, M.; C.J.; Elbadry; Gibson; J.A.; J.C.; J.G. Jr. and Lednicky; M.A.; Morris; Stephenson; Subramaniam, K.; T.B.; Waltzek |
| EPI_ISL_434637 | Lednicky Laboratory, Emerging Pathogens Institute, University of Florida. | Lednicky Laboratory, Emerging Pathogens Institute, University of Florida. | C.J.; Elbadry; Gibson; J.A.; J.C.; J.G. Jr. and Lednicky; M.A.; Morris; Stephenson; Subramaniam, K.; T.B.; Waltzek |
| EPI_ISL_538972, EPI_ISL_539009 | Leeds Teaching Hospitals NHS Trust and Public Health England, National Infection Service (Leeds laboratory) | Wellcome Sanger Institute for the COVID-19 Genomics UK (COG-UK) consortium | Antony Hale and Alex Alderton; Cordelia Langford; David K. Jackson; Dominic Kwiatkowski; Ewan Harrison; Holli Carden; Ian Johnston; John Sillitoe on behalf of the Wellcome Sanger Institute COVID-19 Surveillance Team; Katherine L. Harper; Louissa Macfarlane-Smith; Roberto Amato; Sonia Goncalves |
| EPI_ISL_449336, EPI_ISL_449343, EPI_ISL_449350, EPI_ISL_449413, EPI_ISL_449437, EPI_ISL_449547, EPI_ISL_449554, EPI_ISL_499663, EPI_ISL_499693, EPI_ISL_499836, EPI_ISL_499926, EPI_ISL_517094, EPI_ISL_517180, EPI_ISL_517221, EPI_ISL_517328, EPI_ISL_517436, EPI_ISL_611821, EPI_ISL_612070 | | | |
| see above | Liverpool Clinical Laboratories | COVID-19 Genomics UK (COG-UK) Consortium | A Alrezaihi; Alessandro Gerada; Alistair Darby; Angela Cowell; Anita Lucaci; Anu Chawla; Cassie Olateju; Catherine Hartley; Charlotte Nelson; Ecaterina Vamos; Elaine O'Toole; Eleanor G Bentley; Ghada T Shawli; Isabel García-Dorival; James Johnson; James P Stewart; Jenifer Manson; Joanne Watts; Jones Benjamin; Jordan J Clark; Julian Hiscox; L Luu; Lucille Rainbow; M Almsaud; Margaret Hughes; Mark Whitehead; Matthew Gemmell; Miren Iturriza-Gomara; Muhannad Alruwaili; N.P Randle; Neil Swainston; PKF Gilmore; Parul Sharma; Rebekah Penrice-Randal; Richard Eccles; Richard Gregory; Sam Haldenby; Steve Paterson; Stuart D Armstrong; Trevor Ian Robinson; Ximeng Han |
| EPI_ISL_1501697, EPI_ISL_1501698, EPI_ISL_1501699, EPI_ISL_1501700, EPI_ISL_1501701, EPI_ISL_1501702, EPI_ISL_1501703, EPI_ISL_1501704 | | | |
| see above | Lurie Children's Hospital of Chicago | Northwestern University - Ozer Lab | Egon A. Ozer; Judd F. Hultquist; Lacy M. Simons; Larry K. Kociolek; Michael G. Ison; Ramon Lorenzo-Redondo; Taylor J. Dean; William J. Muller; Xiaotian; Zheng |
| EPI_ISL_683684 | M Health Fairview | Minnesota Department of Health, Public Health Laboratory | Alexandra Lorentz; Jacob Garfin; Matt Plumb; and Xiong Wang |
| EPI_ISL_514637, EPI_ISL_514650, EPI_ISL_514657 | M Health Fairview St. Joseph's Hospital | Minnesota Department of Health, Public Health Laboratory | Jacob Garfin; Matt Plumb; and Xiong Wang |
| EPI_ISL_1257923, EPI_ISL_1257934, EPI_ISL_1257938, EPI_ISL_1257983, EPI_ISL_1257992 | MB-Cadham Provincial laboratory | National Microbiology Laboratory (NML) | Anna Majer; Anneliese Landgraff; CanCOGeN's metadata curation team; Darian Hole; David Alexander; Elsie Grudeski; Gary Van Domselaar; Grace Seo; Jared Bullard; Jennifer Tanner; Kerry Dust; Kirsten Biggar; Madison Chapel; Morag Graham; Natalie Knox; Nathalie Bastien; Paul Van Caeseele; Philip Mabon; Public Health Agency of Canada CanCOGeN team; Rhiannon Huzarewich; Russell Mandes; Shari Tyson; Timothy Booth; Yan Li |
| EPI_ISL_445358 | MEGASALUD SPA. | Instituto de Salud Publica de Chile | Alejandra Acevedo; Andrés E Castillo; Bárbara Parra; Carolina Tambley; Gabriel Leal; Jaime Lagos; Jorge Fernandez; Loredana Arata; Patricia Bustos; Paz Tapia; Rodrigo Fasce; Winston Andrade |
| EPI_ISL_568989, EPI_ISL_569257, EPI_ISL_569470 | MEPHI, Aix Marseille University | MEPHI, Aix Marseille University | Anthony LEVASSEUR |
| EPI_ISL_452133 | MN Department of Health | Pathogen Discovery, Respiratory Viruses Branch, Division of Viral Diseases, Centers for Disease Control and Prevention | Alison S. Laufer Halpin; Anna Montmayeur; Anna Uehara; Christopher A. Elkins; Clinton R. Paden; Haibin Wang; Jing Zhang; Krista Queen; Mary S. Keckler; Rachel Marine; Suxiang Tong; Yan Li; Ying Tao; Zachary Weiner |
| EPI_ISL_426426 | MN PHL Division, Minnesota Department of Health | Pathogen Discovery, Respiratory Viruses Branch, Division of Viral Diseases, Centers for Disease Control and Prevention | Alison S. Laufer Halpin; Anna Uehara; Christopher A. Elkins; Clinton R. Paden; Haibin Wang; Jing Zhang; Krista Queen; Mary S. Keckler; Rachel Marine; Suxiang Tong; Yan Li; Ying Tao |
| EPI_ISL_648018, EPI_ISL_648020, EPI_ISL_648026, EPI_ISL_648028 | MS Public Health Laboratory | Pathogen Discovery, Respiratory Viruses Branch, Division of Viral Diseases, Centers for Disease Control and Prevention | Anna Montmayeur; Anna Uehara; Brian Lynch; Clinton R. Paden; Haibin Wang; Jing Zhang; Krista Queen; Rachel Marine; Suxiang Tong; Yan Li; Ying Tao |
| EPI_ISL_415151, EPI_ISL_421364, EPI_ISL_421365, EPI_ISL_421393, EPI_ISL_421426, EPI_ISL_422517, EPI_ISL_422523, EPI_ISL_422545, EPI_ISL_450016, EPI_ISL_450091, EPI_ISL_450102, EPI_ISL_450110, EPI_ISL_450115, EPI_ISL_450126, EPI_ISL_450139, EPI_ISL_450144, EPI_ISL_450151 | | | |
| see above | MSHS Clinical Microbiology Laboratories | MSHS Pathogen Surveillance Program | Adolfo Garcia-Sarstre; Adolfo Garcia-Sastre; Ajay Obla; Alberto Paniz-mondolfi; Ana S. Gonzalez-Reiche; Ana Silvia Gonzalez-Reiche; Bremy Albuquerque; Bremy Alburquerque; Emilia Sordillo; Florian Krammer; Gopi Patel; Harm van Bakel; Jayeeta Dutta; Jose Polanco; Juan Soto; Judith Aberg; Lisa Miorin; Matthew Hernandez; Melissa Gitman; Melissa Smith; Mitchell Sullivan; Nancy Francoeur; Randy Albrecht; Robert Sebra; Shelcie Fabre; Shwetha Sridhar Hara; Viviana Simon; Wen-chun Liu; Ying-Chih Wang; Zenab Khan |
| EPI_ISL_513429, EPI_ISL_513442, EPI_ISL_513462, EPI_ISL_513463, EPI_ISL_513464, EPI_ISL_513467, EPI_ISL_513483, EPI_ISL_513489, EPI_ISL_513505 | | | |
| see above | Maine HETL | Tewhey Lab, The Jackson Laboratory | Barter, M.; Dewey, H.; H. and Tewhey, R.; Lynch, R.; Matluk, N.; Munger |
| EPI_ISL_454429, EPI_ISL_454430, EPI_ISL_454431 | Maryland Department of Health Laboratories Administration | Maryland Department of Health Laboratories Administration | MDH Laboratories Administration |
| EPI_ISL_460102, EPI_ISL_460114, EPI_ISL_460119, EPI_ISL_460170, EPI_ISL_460178, EPI_ISL_460179, EPI_ISL_460186, EPI_ISL_460205, EPI_ISL_460214, EPI_ISL_460232, EPI_ISL_460264, EPI_ISL_460285, EPI_ISL_460320, EPI_ISL_460338, EPI_ISL_460355, EPI_ISL_460375, EPI_ISL_460382, EPI_ISL_460388, EPI_ISL_460393, EPI_ISL_460411, EPI_ISL_460415, EPI_ISL_460433, EPI_ISL_460446, EPI_ISL_460467, EPI_ISL_460470 | | | |
| see above | Massachusetts General Hospital | Infectious Disease Program, Broad Institute of Harvard and MIT | A.E.; Adams, G.; Anahtar, M.; B.L.; B.W.; Bauer, M.; Birren; Branda, J.; Carter, A.; Cerrato, F.; Chaluvadi, S.; Chapman; Cusick, C.; D.J.; DeRuff, K.; Flowers, K.; Gallagher, G.; Gladden-Young, A.; Gnirke, A.; Harris, J.; J.E.; K.J.; LaRocque, R.; Lagerborg, K.; Lemieux; Lin; Loreth, C.; MacInnis; Neumann, A.; Normandin, E.; P.C.; Park; Pierce, V.; Reilly, S.; Rosenberg, E.; Rudy, M.; Ryan, E.; S.B.; Sabeti; Shaw, B.; Siddle; Slater, D.; Smole, S.; Tomkins-Tinch, C.; Turbett, S. |
| EPI_ISL_791516, | Massachusetts State Public Health Laboratory | Infectious Disease Program, Broad Institute of Harvard and MIT | A.E.; Adams, G.; Anahtar, M.; B.L.; B.W.; Bauer, M.; Birren; Branda, J.; Carter, A.; Cerrato, F.; Chaluvadi, S.; Chapman; Cusick, C.; D.J.; DeRuff, K.; E. and Sabeti; Flowers, K.; Gallagher, G.; Gladden-Young, A.; Gnirke, A.; Harris, J.; J.E.; K.J.; |

| | | | |
|---|---|---|---|
| EPI_ISL_791518 | | | LaRocque, R.; Lagerborg, K.; Lemieux; Lin; Loreth, C.; MacInnis; Neumann, A.; Normandin, E.; P.C.; Park; Pierce, V.; Reilly, S.; Rosenberg; Rudy, M.; Ryan, E.; S.B.; Shaw, B.; Siddle; Slater, D.; Smole, S.; Tomkins-Tinch, C.; Turbett, S. |
| EPI_ISL_420879 | Mater Pathology | Public Health Virology Laboratory | Alyssa Pyke; Amanda De Jong; Andrew Van Den Hurk; Bixing Huang; Carmel Taylor; David Warrilow; Doris Genge; Elisabeth Gamez; Glen Hewitson; Ian Maxwell Mackay; Inga Sultana; Jamie McMahon; Jean Barcelon; Judy Northill; Mitchell Finger; Natalie Simpson; Neelima Nair; Peter Burtonclay; Peter Moore; Sarah Wheatley; Sean Moody; Sonja Hall-Mendelin; Timothy Gardam; and Frederick Moore |
| EPI_ISL_477291, EPI_ISL_495597, EPI_ISL_514638, EPI_ISL_514639, EPI_ISL_514655, EPI_ISL_518860, EPI_ISL_518863, EPI_ISL_518866, EPI_ISL_518868, EPI_ISL_518869, EPI_ISL_518871, EPI_ISL_518875, EPI_ISL_518879, EPI_ISL_518880, EPI_ISL_518884, EPI_ISL_518885, EPI_ISL_518894 | | | |
| see above | Mayo Clinic & Mayo Clinic Laboratories | Minnesota Department of Health, Public Health Laboratory | Jacob Garfin; Kelly Pung; Matt Plumb; and Xiong Wang |
| EPI_ISL_500513, EPI_ISL_500515, EPI_ISL_500516, EPI_ISL_500534, EPI_ISL_500535, EPI_ISL_501074, EPI_ISL_501075 | | | |
| see above | Mayo Clinic Laboratories | University of Washington Virology Lab | Alexander Greninger; Amin Addetia; Hong Xie; Keith R Jerome; Lasata Shrestha; Meei-Li Huang; Pavitra Roychoudhury; Truong Nguyen; Victoria M Rachleff |
| EPI_ISL_583461, EPI_ISL_583462 | Memorial Sloan Kettering Cancer Center | van Bakel Laboratory, Genetics and Genomics Sciences, Icahn School of Medicine at Mount Sinai | Adolfo García-Sastre; Adriana van de Guchte; Ajay Obla; Ana S. Gonzalez-Reiche; Genovefa Papanicolaou; Gunjan Shah; Harm van Bakel; Jayeeta Dutta; Judith Aberg; Kent Sepkowitz; Miguel-Angel Perales; Ngolela Esther Babady; Sadaf Aslam; Teresa Aydillo; Tobias Hohl; Zenab Khan; and Mini Kamboj |
| EPI_ISL_2099222, EPI_ISL_2099225 | Michael E. DeBakey VA Medical Center | Pathogen Discovery, Respiratory Viruses Branch, Division of Viral Diseases, Centers for Disease Control and Prevention | Adam Retchless; Anna Kelleher; Anna Uehara; Brian Lynch; Clinton R. Paden; Dhwani Batra; Haibin Wang; Han Jia Justin Ng; Jasmine Padilla; Jing Zhang; Justin Lee; Krista Queen; Mark Burroughs; Mili Sheth; Morgan Davis; Peter Cook; Rachel Marine; Sarah Nobles; Suxiang Tong; Tara Coalter; Yan Li; Ying Tao |
| EPI_ISL_436809, EPI_ISL_436820, EPI_ISL_436823, EPI_ISL_436828, EPI_ISL_436833, EPI_ISL_436840, EPI_ISL_436847, EPI_ISL_436850, EPI_ISL_436865, EPI_ISL_437120, EPI_ISL_437122, EPI_ISL_437129, EPI_ISL_437137, EPI_ISL_437139, EPI_ISL_437145, EPI_ISL_437155, EPI_ISL_437159, EPI_ISL_447173, EPI_ISL_447187, EPI_ISL_447192, EPI_ISL_447201, EPI_ISL_447212, EPI_ISL_450583, EPI_ISL_452268, EPI_ISL_452325, EPI_ISL_460008, EPI_ISL_471678, EPI_ISL_507637, EPI_ISL_507647, EPI_ISL_507672, EPI_ISL_507688, EPI_ISL_507870, EPI_ISL_507912, EPI_ISL_507929, EPI_ISL_516303, EPI_ISL_516317, EPI_ISL_529810, EPI_ISL_565961, EPI_ISL_594199, EPI_ISL_1905543, EPI_ISL_1905626, EPI_ISL_2246326 | | | |
| see above | Michigan Department of Health and Human Services, Bureau of Laboratories | Michigan Department of Health and Human Services, Bureau of Laboratories | Blankenship HM; Riner D; Soehnlen MK |
| EPI_ISL_1069173 | Micrbiological Laboratory,Lu`an Center for Disease Control and Prevention | Micrbiological Laboratory,Lu`an Center for Disease Control and Prevention | Chang Hongwei; Chen Beilei; Chen Zhichao; Gao Dawei; Li Zhaoyang; Yang Wei; Zhang Feng; Zhang Limei; Zhang Qin; Zhu Rui |
| EPI_ISL_426583 | Microbial Genomics Laboratory, Institut Pasteur Monteivdeo | Microbial Genomics Laboratory, Institut Pasteur Montevideo, Uruguay | Cecilia Salazar; Florencia Díaz-Viraqué; Gonzalo Moratorio; Gregorio Iraola; Marianoel Pereira; Pilar Moreno |
| EPI_ISL_426479, EPI_ISL_426480 | Microbial Genomics Laboratory, Institut Pasteur Montevideo | Microbial Genomics Laboratory, Institut Pasteur Montevideo | Cecilia Salazar; Florencia Díaz-Viraqué; Gonzalo Moratorio; Gregorio Iraola; Marianoel Pereira; Pilar Moreno |
| EPI_ISL_426476, EPI_ISL_426477, EPI_ISL_426478 | Microbial Genomics Laboratory, Institut Pasteur Montevideo | Microbial Genomics Laboratory, Institut Pasteur Montevideo, Uruguay | Cecilia Salazar; Florencia Díaz-Viraqué; Gonzalo Moratorio; Gregorio Iraola; Marianoel Pereira; Pilar Moreno |
| EPI_ISL_426481, EPI_ISL_426482 | Microbial Genomics Laboratory, Institut Pasteur Montevideo, Uruguay | Microbial Genomics Laboratory, Institut Pasteur Montevideo | Cecilia Salazar; Florencia Díaz-Viraqué; Gonzalo Moratorio; Gregorio Iraola; Marianoel Pereira; Pilar Moreno |
| EPI_ISL_426584 | Microbial Genomics Laboratory, Institut Pasteur Montevideo, Uruguay | Microbial Genomics Laboratory, Institut Pasteur Montevideo, Uruguay | Cecilia Salazar; Florencia Díaz-Viraqué; Gonzalo Moratorio; Gregorio Iraola; Marianoel Pereira; Pilar Moreno |
| EPI_ISL_480561 | Microbiological Diagnostic Unit - Public Health Laboratory (MDU-PHL) | MDU-PHL | Sait, M.; Schultz M.; Seemann T.; Sherry, N. |
| EPI_ISL_419714, EPI_ISL_419715, EPI_ISL_419716, EPI_ISL_419717, EPI_ISL_419719, EPI_ISL_419727, EPI_ISL_419728, EPI_ISL_419823, EPI_ISL_420007, EPI_ISL_420008, EPI_ISL_427059, EPI_ISL_427066, EPI_ISL_427067, EPI_ISL_430645 | | | |
| see above | Microbiological Diagnostic Unit Public Health Laboratory | Microbiological Diagnostic Unit Public Health Laboratory | Sait, M.; Schultz M.; Seemann T.; Sherry, N. |
| EPI_ISL_1069198, EPI_ISL_1069200, EPI_ISL_1069213 | Microbiology Laboratory,Lu`an Center for Disease Control and Prevention | Microbiology Laboratory,Lu`an Center for Disease Control and Prevention | Chang Hongwei; Chen Beilei; Chen Zhichao; Gao Dawei; Li Zhaoyang; Yang Wei; Zhang Feng; Zhang Limei; Zhang Qin; Zhu Rui |
| EPI_ISL_468067 | Microbiology and Immunology, The Peter Doherty Institute for Infection and Immunity | Microbiology and Immunology, The Peter Doherty Institute for Infection and Immunity | Antonia da Costa, E.; Barreto, I.; Caly, L.; Canisia, D.; Dakh, F.; Dolores de Jesus da Costa, M.; Douglas; Druce, J.; Francis, J.; Freeman, K.; Jayanti Pereira Tilman, A.; M.B.; Marr, I.; Meumann, E.; N. and Baird, R.; Sait, M.; Salles de Sousa, A.; Schultz; Seemann, T.; Sherry, N.; Soares da Silva, E.; Wapling, J.; Ximenes, J. |
| EPI_ISL_434571 | Microbiology, The University of Hong Kong | Microbiology, The University of Hong Kong | Chan; J.F.W. and Yuen; K.-Y. |
| EPI_ISL_475720, EPI_ISL_475721 | Microbiology, University Hospital Donostia | Microbiology, University Hospital Donostia | Cilla, G.; J.M.; Marimon; Montes, M.; Pineiro, L. |
| EPI_ISL_456213, EPI_ISL_579132, EPI_ISL_579303, EPI_ISL_579304, EPI_ISL_579311 | Middlemore Hospital | Institute of Environmental Science and Research (ESR) | Anja Werno; Antje van der Linden; Arlo Upton; Chris Mansell; David Hammer; Dragana Drinkovic; Erasmus Smit; Gary McAuliffe; Hana Sofia Andersson; Hermes Perez; James Ussher; Jill Sherwood; Jing Wang; Joep de Ligt; Josh Freeman; Julia Howard; Juliet Elvy; Lauren Jelly; Mary DeAlmeida; Matt Blakiston; Matt Storey; Matthew Rogers; Max Bloomfield; Michael Addidle; Michelle Balm; Muhammad Faisal; Nikki Freed; Olin Silander; Sally Roberts; Sarah Jefferies; Sharmini Muttaiyah; Susan Morpeth; Susan Taylor; Timothy Blackmore; Vani Sathyendran; Veronica Playle; Virginia Hope; Xiaoyun Ren |
| EPI_ISL_437317 | Ministry of Health Turkey | Ministry of Health Turkey | Fatma Bayrakdar; Selçuk Kılıç; Süleyman Yalçın; Tülin Demir |
| EPI_ISL_414588, EPI_ISL_414590, EPI_ISL_417194, EPI_ISL_417477, EPI_ISL_417480, EPI_ISL_417500, EPI_ISL_417503, EPI_ISL_419391, EPI_ISL_419392, EPI_ISL_421687, EPI_ISL_421690, EPI_ISL_427280, EPI_ISL_437361, EPI_ISL_437362, EPI_ISL_437363, EPI_ISL_450748, EPI_ISL_450752, EPI_ISL_450759, EPI_ISL_450761, EPI_ISL_450771, EPI_ISL_450773, EPI_ISL_450774, EPI_ISL_460045, EPI_ISL_460046, EPI_ISL_460052, EPI_ISL_460053, EPI_ISL_460058, EPI_ISL_460063, EPI_ISL_460064, EPI_ISL_460068, EPI_ISL_460072, EPI_ISL_482988, EPI_ISL_482991, EPI_ISL_482992, EPI_ISL_482994, EPI_ISL_482996, EPI_ISL_483001, EPI_ISL_483005, EPI_ISL_483012, EPI_ISL_483017, EPI_ISL_495613, EPI_ISL_495619, EPI_ISL_495626, EPI_ISL_496919, EPI_ISL_496920, EPI_ISL_507979, EPI_ISL_514663, EPI_ISL_514665, EPI_ISL_514671, EPI_ISL_530224 | | | |
| see above | Minnesota Department of Health, Public Health Laboratory | Minnesota Department of Health, Public Health Laboratory | Jacob Garfin; Jacob Garfin and Xiong Wang; Jake Garfin and Xiong Wang; Matt Plumb; Xiong Wang; and Xiong Wang |
| EPI_ISL_1362553, EPI_ISL_1362561, EPI_ISL_1362565, EPI_ISL_1362572, EPI_ISL_1362576, EPI_ISL_1362593, EPI_ISL_1392807, EPI_ISL_1392816 | | | |
| see above | Missouri State Public Health Laboratory | Missouri State Public Health Laboratory | Ashley New; Joshua Barry; Matthew Sinn |
| EPI_ISL_435137, EPI_ISL_463740 | Mohammed Bin Rashid University of Medicine and Health Sciences | Al Jalila Genomics Center | Abdulmajeed Alkhaja; Abiola Catherine Senok; Ahmad Abou Tayoun; Alawi Alsheikh-Ali; Divinlal Harilal; Hamda Khansaheb; Hanan Al Suwaidi; Mohammed Uddin; Norbert Nowotny; Qutayba Hamid; Rabih Halwani; Rifat Hamoudi; Rupa Murthy Varghese; Sathishkumar Ramaswamy; Tom Loney; Zulfa Omar Deesi |
| EPI_ISL_444455 | Molecular Infectious Disease | Molecular Infectious Disease | Anderson; B.P.; Gerasimova, A.; Kagan; R.M. and Owen, R.; Rosenthal; S.H. |
| EPI_ISL_884533, EPI_ISL_884541, EPI_ISL_884542, EPI_ISL_884545, EPI_ISL_884549, EPI_ISL_884552, EPI_ISL_884573 | | | |
| see above | Molecular Microbiology & Immunology, University of Missouri | Molecular Microbiology & Immunology, University of Missouri | C.Y.; G.M.; Hang, J.; Li, T.; Lidl; Tang; Wan; X.-F. |
| EPI_ISL_826646 | Montefiore Medical Center | Albert Einstein College of Medicine, Dept. of Microbiology & Immunology, Chandran lab | Amy S. Fox; Ariel S. Wirchnianski; Aviv Bergman; Catalina Florez; D. Yitzchak Goldstein; David L. Goldman; Denise Haslwanter; Ethan Laudermilch; Hnin Khine; J. Maximilian Fels; Jason Barnhill; Johanna P. Daily; Karin A. Skalina; Kartik Chandran; Libusha Kelly; Lucia R. Wolgast; M. Eugenia Dieterle; Robert H. Bortz III; Rohit K. Jangra; Ryan Forster; Saad Khan; Surksha Sirichand; Wendy Szymczak; William B. Mitchell |
| EPI_ISL_631913 | Mount Sinai Hospital | New York City Public Health Laboratory | Jade Wang; et al. |
| EPI_ISL_631507, EPI_ISL_631880 | Mount Sinai West | New York City Public Health Laboratory | Jade Wang; et al. |
| EPI_ISL_424871, EPI_ISL_452124 | NC State Laboratory of Public Health | Pathogen Discovery, Respiratory Viruses Branch, Division of Viral Diseases, Centers for Disease Control and Prevention | Alison S. Laufer Halpin; Anna Montmayeur; Anna Uehara; Christopher A. Elkins; Clinton R. Paden; Haibin Wang; Jing Zhang; Krista Queen; Mary S. Keckler; Rachel Marine; Suxiang Tong; Yan Li; Ying Tao; Zachary Weiner |
| EPI_ISL_632243, EPI_ISL_632246 | NHSGGC West of Scotland Specialist Virology Centre | MRC-University of Glasgow Centre for Virus Research | Alasdair MacLean; Alex Alderton; Ana da Silva Filipe; Antonia Ho; Cordelia Langford; Daniel Mair; David K. Jackson; David L Robertson; Dominic Kwiatkowski; Emma Thomson; Ewan Harrison; Ian Johnston; James Shepherd; Jenna Nichols; John Sillitoe on behalf of the Wellcome Sanger Institute COVID-19 Surveillance Team (http://www.sanger.ac.uk/covid-team); Joseph Hughes; Kathy Li; Kathy Smollett; Kyriaki Nomikou; Lily Tong; Natasha Jesudason; Natasha Johnson; Rajiv Shah; Richard Orton; Roberto Amato; Rory Gunson; Sonia Goncalves; Sreenu Vattipally; Stephen Carmichael |
| EPI_ISL_459520, EPI_ISL_459544, EPI_ISL_459550, EPI_ISL_459551, EPI_ISL_459553, EPI_ISL_459561, EPI_ISL_459570, EPI_ISL_459573, EPI_ISL_459592, EPI_ISL_459618, EPI_ISL_459625, EPI_ISL_459640, EPI_ISL_459650, EPI_ISL_459652, EPI_ISL_459653, EPI_ISL_459667, EPI_ISL_459671, EPI_ISL_459693, EPI_ISL_469944, EPI_ISL_469972, EPI_ISL_469973, EPI_ISL_469975, EPI_ISL_469997, EPI_ISL_470000, EPI_ISL_470003, EPI_ISL_470010, EPI_ISL_489393, EPI_ISL_489588, EPI_ISL_489589, EPI_ISL_489594, EPI_ISL_489611, EPI_ISL_489636, EPI_ISL_489647, EPI_ISL_489661, EPI_ISL_489662, EPI_ISL_489664, EPI_ISL_489667, EPI_ISL_489681, EPI_ISL_489684, EPI_ISL_489698, EPI_ISL_524563 | | | |
| see above | NHSGGC West of Scotland Specialist Virology Centre / MRC-University of Glasgow Centre for Virus Research | Wellcome Sanger Institute for the COVID-19 Genomics UK (COG-UK) consortium | Alasdair MacLean; Alice Broos; Ana da Silva Filipe; Antonia Ho; Cordelia Langford; Daniel Mair; David K. Jackson; David L Robertson; Dominic Kwiatkowski; Elihu Aranday-Cortes; Emma Thomson and Alex Alderton; Ewan Harrison; Ian Johnston; James Shepherd; Jenna Nichols; John Sillitoe on behalf of the Wellcome Sanger Institute COVID-19 Surveillance Team (http://www.sanger.ac.uk/covid-team); Joseph Hughes; Kathy Li; Kathy Smollett; Kirstyn Brunker; Kyriaki Nomikou; Lily Tong; Marc Niebel; Natasha Jesudason; Natasha Johnson; Patawee Asamaphan; Rajiv Shah; Richard Orton; Roberto Amato; Rory Gunson; Sarah McDonald; Sonia Goncalves; Sreenu Vattipally; Stephen Carmichael; Yasmin Parr |
| EPI_ISL_452202, EPI_ISL_452203, EPI_ISL_452204, EPI_ISL_452205, EPI_ISL_454531, EPI_ISL_454534, EPI_ISL_454535, EPI_ISL_454536, EPI_ISL_454537, EPI_ISL_454538 | | | |
| see above | NIV Influenza | NIV Influenza | Potdar V |
| EPI_ISL_450324 | NIV Pune | CSIR-Centre for Cellular and Molecular Biology | A. Awhale; A. Gondhalikar; A. Jagtap; Archana Bharadwaj Siva; B. Minhas S. Bharadwaj; Dhiviya Vedagiri; Disha Nanda; Divya Das; Divya Gupta; Dr ML Choudhary; Dr Priya Abraham; Dr V A Potdar; Gangumala Srinivas Reddy; H Kaushal Y. Gurav; H. Kengle; Jotin Gogoi; K. Patel; Karthik Bharadwaj Tallapaka; Krishnan Harinivas Harshan; Lamuk Zaveri; M. Kakade; Manish Bhattacharjee; N Srivastava; Namami Gaur; P Shinde; P. Malsane; Payel Mukherjee; Priya Singh; Purushotham Vodnala; R. Gunjikar; Rakesh K Mishra; Ravi Prasad Mukku; Renu Sudhakar; S. Bhorekar; S. Digraskar; S. Hundekar; S. Jadhav; S. Salve; S. Tomar; Sakshi Shambhavi; Santosh Kumar Kuncha; Shagufta Khan; Sofia Banu; Somesh Gorde; Sujoy Deb; Swati Bayyana; Tulasi Nagabandi; U. Saha; V Malik; V. Awtade; V. Vipat; Vishal Sah; Yogesh Balakartik; Zeba Rizvi |
| EPI_ISL_422387, EPI_ISL_422397, EPI_ISL_422402, EPI_ISL_422403, EPI_ISL_422406 | NMIMR, Department of Virology | WACCBIP, University of Ghana | Abraham Kwabena Anang; Augustina Arjarquah; Bright Adu; Collins M. Morang'a; Dominic S. Y. Amuzu; Erasmus Kotey; Evelyn Bonney; Fred Tei-Maya; George B. Kyei; Gordon A. Awandare; Ivy Asante; Joyce M. Ngoi; Kofi Bonney; Linda Boatemaa; Miriam Eshun; Peter Quashie; Selassie Kumordjie; Vanessa Magnussen; William Ampofo |
| EPI_ISL_1666667 | NS-QEII Health Sciences Centre | National Microbiology Laboratory (NML) | Anna Majer; Anneliese Landgraff; CanCOGeN's metadata curation team; Dan Gaston; Darian Hole; Elsie Grudeski; Gary Van Domselaar; Grace Seo; Janice Pettipas; Jason LeBlanc; Jennifer Tanner; Kirsten Biggar; Madison Chapel; Morag Graham; Natalie Knox; Nathalie Bastien; Philip Mabon; Public Health Agency of Canada CanCOGeN team; Rhiannon Huzarewich; Russell Mandes; Shari Tyson; Timothy Booth; Todd Hatchette; Yan Li |
| EPI_ISL_458630, EPI_ISL_488600, EPI_ISL_488602, EPI_ISL_488680, EPI_ISL_488682, EPI_ISL_488732, EPI_ISL_488783 | | | |
| see above | NU-OMICS DNA Sequencing research facility, Northumbria University | Wellcome Sanger Institute for the COVID-19 Genomics UK (COG-UK) consortium | Andrew Nelson; Brendan Payne; Chris Duncan; Clive Graham; Cordelia Langford; Darren Smith and Alex Alderton; David K. Jackson; Debra Padgett; Dominic Kwiatkowski; Edward Barton; Emma Swindells; Ewan Harrison; Garren Scott; Gary Black; Gary Eltringham; Greg Young; Ian Johnston; Jane Greenaway; Jennifer Collins; John Allan; John Sillitoe on behalf of the Wellcome Sanger Institute COVID-19 Surveillance Team (http://www.sanger.ac.uk/covid-team); Joshua Loh; Lynn Dover; Matthew Bashton; Paul Baker; Roberto Amato; Sarah Essex; Sheia Waugh; Shirelle Burton-Fanning; Sonia Goncalves; Steve Ligget; Wen Yew; Yusri Taha |
| EPI_ISL_436047, | NYC Department of Health and Mental Hygiene | Pathogen Discovery, Respiratory Viruses Branch, Division of Viral | Anna Uehara; Bettina Bankamp; Christy Harrison; Clinton R. Paden; Haibin Wang; Jasmine Padilla; Jennifer Rakeman; Jing Zhang; Justin Lee; Krista Queen; Suxiang Tong; Yan Li; Ying Tao; Zachary Weiner |

| | | | |
|---|---|---|---|
| EPI_ISL_436068 | | Diseases, Centers for Disease Control and Prevention | |
| EPI_ISL_631557, EPI_ISL_632072, EPI_ISL_632186 | NYC HH Lincoln Medical And Mental Health Center | New York City Public Health Laboratory | Jade Wang; et al. |
| EPI_ISL_418970 | NYU Langone Health | Department of Pathology and Medicine, New York University School of Medicine | Adriana Heguy; Amy Rapkiewicz; Antonio Serrano; Christian Marier; Dacia Dimartino; Emily Huang; Gael Westby; George Jour; Guomiao Shen; Jared Pinnell; John Cadley; John Chen; Margaret Black; Maria Aguero-Rosenfeld; Marie Samanovic-Golden; Mark J. Mulligan; Matija Snuderl; Matthew T. Maurano; Megan Hogan; Nick Vulpescu; Paolo Cotzia; Paul Zappile; Peter Meyn; Sitharam Ramaswami; Xiaojun Feng; Yutong Zhang |
| EPI_ISL_419696, EPI_ISL_419698, EPI_ISL_421707, EPI_ISL_421714, EPI_ISL_421723, EPI_ISL_436738, EPI_ISL_444632, EPI_ISL_444637, EPI_ISL_444652, EPI_ISL_444682, EPI_ISL_451443, EPI_ISL_451468 | | | |
| see above | NYU Langone Health | Departments of Pathology and Medicine, New York University School of Medicine | Adriana Heguy; Alireza Khodadadi-Jamayran; Amy Rapkiewicz; Andre M. Ribeiro-dos-Santos; Andrew Lytle; Antonio Serrano; Brendan Belovarac; Christian Marier; Dacia Dimartino; Emily Guzman; Emily Huang; Gael Westby; George Jour; Guomiao Shen; Iman Osman; Jared Pinnell; John Cadley; John Chen; Lawrence H. Lin; Ludovic Boytard; Margaret Black; Maria Aguero-Rosenfeld; Marie Samanovic-Golden; Mark J. Mulligan; Matija Snuderl; Matthew T. Maurano; Megan Hogan; Nick Vulpescu; Paolo Cotzia; Paul Zappile; Peter Meyn; Raquel Ordonez Ciriza; Raven Luther; Sitharam Ramaswami; Tatyana Gindin; Theodore Vougiouklakis; Vanessa Raabe; Xiaojun Feng; Yutong Zhang |
| EPI_ISL_480002 | Nagasaki Prefectural Institute for Environmental Research and Public Health | Pathogen Genomics Center, National Institute of Infectious Diseases | Fumiaki Matsumoto; Hajime Kamiya; Kentaro Itokawa; Makoto Kuroda; Masanori Hashino; Motoi Suzuki; Rina Tanaka; Tsuyoshi Sekizuka |
| EPI_ISL_480042, EPI_ISL_480043, EPI_ISL_480044, EPI_ISL_480045, EPI_ISL_480046, EPI_ISL_480047, EPI_ISL_480048, EPI_ISL_480049, EPI_ISL_480050, EPI_ISL_480051, EPI_ISL_480052, EPI_ISL_480053, EPI_ISL_480054, EPI_ISL_480055, EPI_ISL_480056, EPI_ISL_480057, EPI_ISL_480058, EPI_ISL_480059, EPI_ISL_480060, EPI_ISL_480061, EPI_ISL_480062, EPI_ISL_480063, EPI_ISL_480064 | | | |
| see above | Nagoya City Public Health Research Institute | Pathogen Genomics Center, National Institute of Infectious Diseases | Hajime Kamiya; Kentaro Itokawa; Makoto Kuroda; Masanori Hashino; Motoi Suzuki; Rina Tanaka; Shinichiro Shibata; Takuya Miki; Tsuyoshi Sekizuka |
| EPI_ISL_454572 | National Center of Expertise | National Center for Expertise, Kazakhstan National Center for Biotechnology, Kazakhstan | ; Abdaliyev Askar; Akhmetollayev Ilyas; Amirgazin Asylulan; Aushakhmetova Zabira; Kalendar Ruslan; Lutsay Viktoriya; Rakhmetova Akbota; Ramankulov Yerlan; Shevtsov Alexandr |
| EPI_ISL_454571 | National Center of Expertise | National Center for Expertise, National Center for Biotechnology, Kazakhstan | ; Abdaliyev Askar; Akhmetollayev Ilyas; Amirgazin Asylulan; Aushakhmetova Zabira; Kalendar Ruslan; Lutsay Viktoriya; Rakhmetova Akbota; Ramankulov Yerlan; Shevtsov Alexandr |
| EPI_ISL_435110, EPI_ISL_435111 | National Centre for Disease control (NCDC), CSIR-Institute of Genomics and Integrative Biology (CSIR-IGIB) | NCDC/CSIR-IGIB | Aarti Tewari; Anurag Agrawal; Bharathram Uppili; Bibhash Nandi; Debasis Dash; Dhirendra Kumar; Hema Gogia; Hemlata Lall; Himanshu Vashisht; Mahesh Dhar; Manju Bala; Meena Datta; Mitali Mukerji; Mohammed Faruq; Nidhi Saini; Nishu Tyagi; Partha Rakshit; Pooja Sharma; Poonam Gupta; Pramod Kumar; Prateek Singh; Preeti Madan; Priyanka Singh; Rajesh Pandey; Sandhya Kabra; Saruchi Wadhwa; Satyabrata Bag; Simrita Singh; Sujeet Singh; Uma Sharma; Varun Jaiswal; Vivekanand A |
| EPI_ISL_410535, EPI_ISL_414378, EPI_ISL_414379, EPI_ISL_414380, EPI_ISL_420099, EPI_ISL_420100, EPI_ISL_420107 | | | |
| see above | National Centre for Infectious Diseases | Programme in Emerging Infectious Diseases, Duke-NUS Medical School | Barnaby E Young; Danielle E Anderson; David CB Lye; Gavin JD Smith; Jayanthi Jayakumar; Louisa Sun; Martin Linster; Yan Zhuang; Yee Sin Leo; Yvonne CF Su |
| EPI_ISL_408668 | National Influenza Center - National Institute of Hygiene and Epidemiology (NIHE) | National Influenza Center - National Institute of Hygiene and Epidemiology (NIHE) | ; Hoang Vu Mai Phuong; Le Thi Quynh Mai; Le Thi Thanh; Nguyen Le Khanh Hang; Nguyen Phuong Anh; Nguyen Vu Son; Pham Thi Hien; Tran Thu Huong; Ung Thi Hong Trang; Vuong Duc Cuong |
| EPI_ISL_416427, EPI_ISL_416429, EPI_ISL_660541 | National Influenza Center, National Institute of Hygiene and Epidemiology (NIHE) | National Influenza Center, National Institute of Hygiene and Epidemiology (NIHE) | Dang Duc Anh; Futoshi Hasebe; Hoang Vu Mai Phuong; Kouichi Morita; Le Quynh Mai; Le Thi Thanh; Meng Ling Moi; Nguyen Le Khanh Hang; Nguyen Phuong Anh; Nguyen Vu Son; Pham Hong Quynh Anh; Pham Thi Hien; Taichiro Takemura; Takeshi Nabeshima; Tran Thu Huong; Ung Thi Hong Trang; Vuong Duc Cuong |
| EPI_ISL_476838 | National Influenza Centre for Northern Greece | National Influenza Centre for Northern Greece | Maria Christoforidi |
| EPI_ISL_596386 | National Institute for Allergy and Infectious Diseases Integrated Research Facility - Frederick (NIAID IRF-Frederick), National Institutes of Health (NIH) | National Institute for Allergy and Infectious Diseases Integrated Research Facility - Frederick (NIAID IRF- Frederick), National Institutes of Health (NIH) | B. and Palacios, G.; Beitzel; J.R.; Kocher, G.; Kugelman |
| EPI_ISL_408480, EPI_ISL_408483, EPI_ISL_408484, EPI_ISL_408485 | National Institute for Viral Disease Control and Prevention, China CDC | National Institute for Viral Disease Control & Prevention, CCDC | Baoying Huang; Baoying Huang , Li Zhao , Fei Ye , Wenbo Xu , George F. Gao , Guizhen Wu; Chongshan Li; Fei Ye; George F. Gao; Guizhen Wu; Huiping Yang; Jianan Xu; Li Zhao; Peihua Niu; Quanyi Wang; Roujian Lu; Wenbo Xu; Wenjie Tan; Wenjie Tan , Xiaoqing Fu , Xiang Zhao , Wenling Wang , Peihua Niu , Roujian Lu; Wenjie Tan , Zhen Teng; Wenling Wang; Xiang Zhao; Xiang Zhao , Wenling Wang , Peihua Niu , Roujian Lu; Yang Pan; Yanhong Sun , Baoying Huang , Li Zhao , Fei Ye , Wenbo Xu , George F. Gao , Guizhen Wu |
| EPI_ISL_408487 | National Institute for Viral Disease Control and Prevention, China CDC | National Institute for Viral Disease Control & Prevention, China CDC | Baoying Huang; Fei Ye; George F. Gao; Guizhen Wu; Jin Xu; Li Zhao; Peihua Niu; Roujian Lu; Wenbo Xu; Wenjie Tan; Wenling Wang; Xiang Zhao; Xueyong Huang |
| EPI_ISL_455584, EPI_ISL_455589, EPI_ISL_455590, EPI_ISL_455591, EPI_ISL_455592, EPI_ISL_455624 | National Institute of Health. Department of medical Sciences, Ministry of Public Health, Thailand | National Institute of Health. Department of medical Sciences, Ministry of Public Health, Thailand | Chittaganpitch; Malinee; Okada; Parnmen; Phuygun; Pilailuk; Siripaporn; Sittiporn; Sunthareeya; Thanadachakul; Thanutsapa; Waicharoen; Warawan; Wongboot |
| EPI_ISL_644743, EPI_ISL_644750, EPI_ISL_644764, EPI_ISL_647976 | National Microbiology Reference Laboratory | Quadram Institute Bioscience | Agnes Juru; Alexander Goredema; Ana-Victoria Gutierrez; Andrew J. Page; Andrew Tarupiwa; Barbra Murwira; Beuty Makamure; Charles Nyagupe; David Baker; Gaetan Thilliez; Gemma Kay; Gibson Mhlanga; Hlanai Gumbo; Isaac Phiri; Justin O'Grady; Leonardo de Oliveira Martins; Muchaneta Mugabe; Portia Manangazira; Robert Kingsley; Sekesai Zinyowera; Tapfumanei Mashe; Tatenda Takawira; Thanh Le Viet |
| EPI_ISL_416885, EPI_ISL_416886 | National Public Health Laboratory | Malaysia Genome Institute | Azrin Ahmad; Enizza Kasim; Hani Mat Hussin; Irni Suhayu Sapian; Mohd Faizal Abu Bakar; Mohd Noor Mat Isa; Noorliza Mohamad Noordin; Nor Azfa Johari; Norazimah Tajudin; Nurhezreen Md Iqbal; Rehan Shuhada Abu Bakar; Selvanesan Sengol; Shahrul Hisham Zainal Ariffin; Shamsidar Sopie; Siti Noraini Othman; W Afiza W Mohd Arifin; Yu Kie Chem; Yusuf Muhammad Noor |
| EPI_ISL_418993, EPI_ISL_418995, EPI_ISL_418996, EPI_ISL_418997, EPI_ISL_418998, EPI_ISL_418999, EPI_ISL_419000, EPI_ISL_419001, EPI_ISL_428822, EPI_ISL_428827, EPI_ISL_428830, EPI_ISL_443231, EPI_ISL_462292, EPI_ISL_462350, EPI_ISL_462415, EPI_ISL_462420, EPI_ISL_469132, EPI_ISL_475963, EPI_ISL_475997, EPI_ISL_475998, EPI_ISL_536435, EPI_ISL_536447, EPI_ISL_574531, EPI_ISL_1315626 | | | |
| see above | National Public Health Laboratory, National Centre for Infectious Diseases | National Public Health Laboratory, National Centre for Infectious Diseases | Chavatte JM; Chavatte Jean-Marc; Cui L; Cui Lin; Lin Cui; Lin RTP; Lin Raymond Tzer Pin; Mak TM; Mak Tze Minn; Octavia S; Octavia Sophie; Raymond Tzer Pin Lin; Sophie Octavia; Tze Minn Mak; Zhenyang Zhou; Zhou Z |
| EPI_ISL_671375, EPI_ISL_1620750 | National Virus Reference Laboratory | Irish Coronavirus Sequencing Consortium - Teagasc Moorepark | Calum Walsh; Fiona Crispie; John Kenny; Jose Sanchez-Morgado; Paul Cotter |
| EPI_ISL_493165, EPI_ISL_493168, EPI_ISL_493169, EPI_ISL_493171, EPI_ISL_493173, EPI_ISL_493174, EPI_ISL_493175, EPI_ISL_493178, EPI_ISL_493181, EPI_ISL_493183, EPI_ISL_493184, EPI_ISL_493185, EPI_ISL_493186, EPI_ISL_493187, EPI_ISL_493188, EPI_ISL_493189, EPI_ISL_493190 | | | |
| see above | National Virus Resource Center, Chinese Academy of Sciences, Wuhan 430071, China | Computational Virology Group, Center for Bacteria and Viruses Resources and Bioinformation, Wuhan Institute of Virology, Chinese Academy of Sciences , Wuhan 430071, China | Di Liu; Hongping Wei; Jianjun Chen; Jin Xiong; Yi Huang; Yi Yan |
| EPI_ISL_444994 | Naval Health Research Center | Naval Medical Research Center Biological Defense Research Directorate | Adrian Paskey; Chris Myers; Dessiree Pena-Gomez; Ewell Hollis; Kimberly Bishop-Lilly; Kyle Long; Logan Voegtly; Melinda Balansay-Ames; Nathaniel Christy; Regina Cer; Roger Pan |
| EPI_ISL_491913 | Naval Infectious Diseases Diagnostic Laboratory | Naval Medical Research Center Biological Defense Research Directorate | Adrian Paskey; Dessiree Pena-Gomez; Francisco Malgon Bautista; Hua Wei Chen; Kimberly Bishop-Lilly; Kyle Long; Lindsay Glang; Logan Voegtly; Mark Simons; Megan Schilling; Regina Cer; Victor Sugiharto |
| EPI_ISL_428203 | Nebraska Public Health Laboratory | UNMC COVID-19 Response Team | UNMC COVID-19 Response Team |
| EPI_ISL_515294, EPI_ISL_515295, EPI_ISL_515298, EPI_ISL_515299, EPI_ISL_515303, EPI_ISL_515304, EPI_ISL_515308, EPI_ISL_515317, EPI_ISL_515322, EPI_ISL_515327, EPI_ISL_515333, EPI_ISL_515343 | | | |
| see above | Nevada State Public Health Laboratory | Nevada State Public Health Laboratory | Andrew Gorzalski; Chris Laverdure; Cyprian Rossetto; David Jackson; Heather Kerwin; Joel R. Sevinsky; Natalie Crawford; Paul Hartley; Richard Tillett; Stephanie Van Hooser; Subhash C. Verma; and Mark Pandori |
| EPI_ISL_467596, EPI_ISL_467603, EPI_ISL_467622, EPI_ISL_467623, EPI_ISL_467624, EPI_ISL_467625, EPI_ISL_467627 | | | |
| see above | New Mexico Department of Health Scientific Laboratory Division | Center for Global Health, University of New Mexico Health Sciences Center | Darrell Dinwiddie; Daryl Domman; Joseph Hicks; Kurt Schwalm; Michael Edwards; Twila Kunde |
| EPI_ISL_427527, EPI_ISL_427531, EPI_ISL_427536, EPI_ISL_427544, EPI_ISL_427583 | NewYork-Presbyterian & Mason Lab | Mason Lab | Alon Shaiber; Arryn Craney; Benjamin Young; Cem Meydan; Chandrima Bhattacharya; Christopher E. Mason; Christopher Mozsary; Craig D. Westover; Daniel J. Butler; David Danko; Dmitry Meleshko; Dong Xu; Ebrahim Afshinnekoo; Fritz J. Sedlazeck; Hanna Rennert; Iman Hajirasouliha; Jenny Xiang; Joel Rosiene; John Sipley; Jonathan Foox; Justyna Gawrys; Krista Ryon; Lars F. Westblade; Lin Cong; Marcin Imielinski; Maria Sierra; Massimo Loda; Matthew MacKay; Melissa Cushing; Mirella Salvatore; Nikolay A. Ivanov; Phyllis Ruggiero; Priya Velu; Shawn Levy |
| EPI_ISL_455422, EPI_ISL_455426 | Nigeria Centre for Disease Control | African Centre of Excellence for Genomics of Infectious Diseases (ACEGID), Redeemer's University, Ede, Osun State, Nigeria | Ajogbasile F.V.; Folarin O.A.; Happi C.T.; Ihekweazu C.; Kayode A.; Oguzie J.; Olawoye I.; Olumade T.; Oluniyi P.E.; Uwanibe J. |
| EPI_ISL_455414, EPI_ISL_455415, EPI_ISL_455418, EPI_ISL_455423, EPI_ISL_455424, EPI_ISL_455425, EPI_ISL_455427, EPI_ISL_527876, EPI_ISL_527877 | | | |
| see above | Nigeria Centre for Disease Control (NCDC) | African Centre of Excellence for Genomics of Infectious Diseases (ACEGID), Redeemer's University, Ede, Osun State, Nigeria | Ajogbasile F.V.; Folarin O.A.; Happi C.T.; Ihekweazu C.; Kayode A.; Oguzie J.; Olawoye I.; Olumade T.; Oluniyi P.E.; Oluniyi P.E. et al; Uwanibe J. |
| EPI_ISL_479913, EPI_ISL_479914, EPI_ISL_479915, EPI_ISL_479916, EPI_ISL_479917, EPI_ISL_479918, EPI_ISL_479919, EPI_ISL_479920, EPI_ISL_479921, EPI_ISL_479922, EPI_ISL_479923, EPI_ISL_479924 | | | |
| see above | Niigata City Public Health Research Institute | Pathogen Genomics Center, National Institute of Infectious Diseases | Hajime Kamiya; Kentaro Itokawa; Makoto Kuroda; Masanori Hashino; Motoi Suzuki; Rina Tanaka; Tsuyoshi Sekizuka; Yurie Takahashi |
| EPI_ISL_479869, EPI_ISL_685297 | Niigata Prefectural Institute of Public Health and Environmental Sciences | Pathogen Genomics Center, National Institute of Infectious Diseases | Hajime Kamiya; Kentaro Itokawa; Makoto Kuroda; Masanori Hashino; Motoi Suzuki; Reiko Arai; Rina Tanaka; Tsuyoshi Sekizuka |
| EPI_ISL_848177, EPI_ISL_848178, EPI_ISL_848546 | North Dakota Department of Health, Public Health Laboratory | North Dakota Department of Health, Public Health Laboratory | Lisa Wingerter |
| EPI_ISL_516851, EPI_ISL_524665 | North West London Pathology, Imperial College Healthcare NHS Trust | Wellcome Sanger Institute for the COVID-19 Genomics UK (COG-UK) consortium | Aileen Rowan; Alison Holmes; Anjna Badhan; Carolina Herrera and Alex Alderton; Cordelia Langford; David K. Jackson; David Muir; Dominic Kwiatkowski; Ewan Harrison; Frankie Bolt; Graham Taylor; Ian Johnston; James Price; John Sillitoe on behalf of the Wellcome Sanger Institute COVID-19 Surveillance Team (http://www.sanger.ac.uk/covid-team); Ling Li; Paul Randell; Roberto Amato; Sonia Goncalves |
| EPI_ISL_478535 | Northumbria University / South Tees Hospitals NHS Foundation Trust / North Cumbria Integrated Care NHS | COVID-19 Genomics UK (COG-UK) Consortium | Andrew Nelson; Brendan Payne; Clive Graham; Darren L Smith; Debra Padgett; Edward Barton; Emma Swindells; Garren Scott; Gary Black; Gary Eltringham; Giles S Holt; Greg R Young; Jane Greenaway; Jennifer Collins; John Allan; Joshua Loh; Lynn Dover; Matthew Bashton; Mohammad A Tariq; Paul Baker; Sarah Essex; Steve Liggett; Wen C Yew; Yusri Taha |

| | | | |
|---|---|---|---|
| | Foundation Trust / North Tees and Hartlepool NHS Foundation Trust / Newcastle Hospitals NHS Foundation Trust | | |
| EPI_ISL_444522, EPI_ISL_444526, EPI_ISL_444530, EPI_ISL_444532, EPI_ISL_444539, EPI_ISL_444543, EPI_ISL_444549, EPI_ISL_444551, EPI_ISL_444554, EPI_ISL_444555, EPI_ISL_444557, EPI_ISL_444559, EPI_ISL_444561, EPI_ISL_444562, EPI_ISL_444563, EPI_ISL_444564, EPI_ISL_444565, EPI_ISL_444566, EPI_ISL_444567, EPI_ISL_444568, EPI_ISL_444569, EPI_ISL_444572, EPI_ISL_444573, EPI_ISL_444577, EPI_ISL_444583, EPI_ISL_444590, EPI_ISL_444593, EPI_ISL_444594, EPI_ISL_444596, EPI_ISL_444597, EPI_ISL_444601, EPI_ISL_444602, EPI_ISL_444603, EPI_ISL_444604, EPI_ISL_444606, EPI_ISL_626357, EPI_ISL_626360, EPI_ISL_626361, EPI_ISL_626362, EPI_ISL_626363, EPI_ISL_626368, EPI_ISL_626369, EPI_ISL_626370, EPI_ISL_626374, EPI_ISL_626375, EPI_ISL_626382, EPI_ISL_626464, EPI_ISL_626467, EPI_ISL_626468, EPI_ISL_936497 | | | |
| see above | Northwestern Memorial Hospital | Ozer Lab | Alan R. Hauser; Chad J. Achenbach; Chao Qi; Egon A. Ozer; Hannah H. Nam; Judd F. Hultquist; Lacy M. Simons; Lawrence J. Jennings; Michael G. Ison; Ramon Lorenzo-Redondo; Scott C. Roberts |
| EPI_ISL_509071 | OHSU Lab Services Molecular Microbiology Lab | Oregon SARS-CoV-2 Genome Sequencing Center | Alec J. Hirsch; Andrew C. Adey; Benjamin N. Bimber; Brendan L. O'Connell; Brian J. O'Roak; Daniel N. Streblow; Guang Fan; Ruth V. Nichols; Sally B. Grindstaff; William B. Messer |
| EPI_ISL_498171 | OUCRU | OUCRU | Dinh Nguyen Huy Man; Guy Thwaites; Lam Anh Nguyet; Lam Minh Yen; Le Manh Hung; Le Nguyen Truc Nhu; Nghiem My Ngoc; Ngo Ngoc Quang Minh; Nguyen Tan Binh; Nguyen Thanh Dung; Nguyen Thanh Phong; Nguyen Thanh Truong; Nguyen Thi Han Ny; Nguyen Thi Thu Hong; Nguyen Tri Dung; Nguyen Van Vinh Chau; Tang Chi Thuong; Tran Chanh Xuan; Tran Nguyen Hoang Tu; Tran Tan Thanh; Tran Tinh Hien; Vo Thanh Lam; and Le Van Tan; for OUCRU COVID-19 research group* |
| EPI_ISL_450741, EPI_ISL_450743 | OUCRU/HTD | OUCRU/HTD | Dinh Nguyen Huy Man; Guy Thwaites; Lam Anh Nguyet; Lam Minh Yen; Le Manh Hung; Le Nguyen Truc Nhu; Nghiem My Ngoc; Ngo Ngoc Quang Minh; Nguyen Tan Binh; Nguyen Thanh Dung; Nguyen Thanh Phong; Nguyen Thanh Truong; Nguyen Thi Han Ny; Nguyen Thi Thu Hong; Nguyen Tri Dung; Nguyen Van Vinh Chau; Tang Chi Thuong; Tran Chanh Xuan; Tran Nguyen Hoang Tu; Tran Tan Thanh; Tran Tinh Hien; Vo Thanh Lam; and Le Van Tan; for OUCRU COVID-19 research group* |
| EPI_ISL_935520, EPI_ISL_940158, EPI_ISL_940553, EPI_ISL_2555539, EPI_ISL_2555540, EPI_ISL_2598025, EPI_ISL_2625640, EPI_ISL_2625732 | | | |
| see above | Ohio Department of Health Laboratory | Ohio Department of Health Laboratory | Allison Black; Brent Lee; Caitlin McDonnell; Eric Brandt; Erica Leasure; Glen McGillivary; Heather Blankenship; Holmes; Jade Mowery; Jennifer; Kelsey Florek; Keoni Omura; Kirtana Ramadugu; Quanta Brown; Stephanie Mccracken; Tyler Payne; and Tammy Bannerman |
| EPI_ISL_479979, EPI_ISL_479980, EPI_ISL_479981, EPI_ISL_479982, EPI_ISL_479983, EPI_ISL_479984, EPI_ISL_480109, EPI_ISL_480110, EPI_ISL_480111, EPI_ISL_480112, EPI_ISL_480113, EPI_ISL_480114, EPI_ISL_480115, EPI_ISL_480116, EPI_ISL_480117 | | | |
| see above | Oita Prefectural Institute of Public Health and Environmental Science | Pathogen Genomics Center, National Institute of Infectious Diseases | Hajime Kamiya; Kentaro Itokawa; Makoto Kuroda; Mari Sasaki; Masanori Hashino; Motoi Suzuki; Rina Tanaka; Tsuyoshi Sekizuka |
| EPI_ISL_457938, EPI_ISL_457976, EPI_ISL_457991, EPI_ISL_457993 | Oman-NIC | Oman-NIC | Abdulla Balkhair; Ahlam Al-Amri; Aisha Al-Amri; Aisha Al-Busaidi; Amina Al Jardani; Fahad Zadjali; Fatma BaAlawi; Hamida AL Barwani; Hanan Al-kindi; Intisar Al-Shukri; Khulood Al-Mammary; Mohammed Al-Tobi; Samiha Al Kharusi; Samira Al-Maruqi; Zeyana AL-Dahmani |
| EPI_ISL_1739546 | Ontario's COVID-19 Genomics Rapid Response Coalition | McMaster University | Ahmed Draia; Allison McGeer; Andrew G. McArthur; Angel Li; Emily Panousis; Hooman Derakhshani; Jalees Nasir; Kuganya Nirmalarajah; Michael Surette; Patryk Aftanas; Samira Mubareka |
| EPI_ISL_2443714 | Orange County Public Health Laboratory | University of California, Irvine | Amanda N. Pinski; Ilhem Messaoudi |
| EPI_ISL_448824 | Oxford Viromics, NDM, University of Oxford; Oxford University Hospitals; Basingstoke and North Hampshire Hospital | COVID-19 Genomics UK (COG-UK) Consortium | Alex Mobbs; Amy Trebes; Anita Justice; Catrin Moore; Christophe Fraser; David Bonsall; David Buck; Emma Wise; George Macintyre; Jessica Lynch; John Todd; Mariateresa de Cesare; Monique Andersson; Nathan Moore; Nick Cortes; Robert Shaw; Stephen Kidd; Tanya Golubchik |
| EPI_ISL_1040031, EPI_ISL_1040034, EPI_ISL_1040036, EPI_ISL_1040037, EPI_ISL_1040039, EPI_ISL_1040040, EPI_ISL_1040044, EPI_ISL_1040046, EPI_ISL_1040048 | | | |
| see above | PCR Laboratory, The First Affiliated Hospital of Zhengzhou University, Zhengzhou, Henan, China | PCR Laboratory, The First Affiliated Hospital of Zhengzhou University, Zhengzhou, Henan, China | Guanglin Cui; Hao Guo; Junhu Wan; Lei Zheng; Li Sun; Lijuan Rong; Meijing Long; Shijie Zhang; Ting Sun; Xianchun Meng; Xiangbo Huang |
| EPI_ISL_440043, EPI_ISL_440094, EPI_ISL_440140, EPI_ISL_443688, EPI_ISL_443746, EPI_ISL_443749, EPI_ISL_443810, EPI_ISL_443999, EPI_ISL_444012 | | | |
| see above | PHE South West Regional Laboratory, National Infection Service | Wellcome Sanger Institute for the COVID-19 Genomics UK (COG-UK) consortium | Alex Alderton; Barry Vipond; Cordelia Langford; David K. Jackson; Dominic Kwiatkowski; Dr Peter Muir; Ewan Harrison; Hannah Pymont; Ian Johnston; John Sillitoe on behalf of the Wellcome Sanger Institute COVID-19 Surveillance Team (http://www.sanger.ac.uk/covid-team); Rich Hopes; Roberto Amato; Sonia Goncalves; Stephanie Hutchings; and Alex Alderton |
| EPI_ISL_455605 | Panyananthaphikkhu Chonprathan Medical Center | National Institute of Health. Department of medical Sciences, Ministry of Public Health, Thailand | Chittaganpitch; Malinee; Okada; Parnmen; Phuygun; Pilailuk; Siripaporn; Sittiporn; Sunthareeya; Thanadachakul; Thanutsapa; Waicharoen; Warawan; Wongboot |
| EPI_ISL_434701 | Panyanunthaphikkhu Chonprathan Medical Center (PCMC) | National Institute of Health. Department of medical Sciences, Ministry of Public Health, Thailand | Chittaganpitch; Malinee; Okada; Parnmen; Phuygun; Pilailuk; Siripaporn; Sittiporn; Sunthareeya; Thanadachakul; Thanutsapa; Waicharoen; Warawan; Wongboot |
| EPI_ISL_654866, EPI_ISL_654874, EPI_ISL_654878, EPI_ISL_654884 | Pasteur Institute in Ho Chi Minh city | Department of Microbiology and Immunology - Pasteur Institute in Ho Chi Minh city | Cao Minh Thắng; Hoàng Minh; Hoàng Như Đào; Hoàng Quốc Cường; Huỳnh Phương Thảo; Huỳnh Thị Kim Loan; Lương Chấn Quang; Nguyễn Hoàng Anh; Nguyễn Hoàng Quân; Nguyễn Thanh Long; Nguyễn Thu Ngọc; Nguyễn Thị Ngọc Thảo; Nguyễn Thị Thanh Thương; Nguyễn Trung Hiếu; Nguyễn Trọng Toàn; Nguyễn Vũ Thượng; Phan Trọng Lân; Phạm Duy Quang; Phạm Thị Nhung; Phạm Thị Thu Hằng; Trần Thị Hồng Kim; Trần Tôn; Viên Đặng Khánh Linh; Vũ Phạm Hồng Nhung; Đào Huy Mạnh; Đặng Thanh Giang |
| EPI_ISL_456215 | PathLab Bay of Plenty | Institute of Environmental Science and Research (ESR) | Anja Werno; Antje van der Linden; Arlo Upton; Chris Mansell; David Hammer; Dragana Drinkovic; Erasmus Smit; Gary McAuliffe; Hana Sofia Andersson; James Ussher; Jill Sherwood; Joep de Ligt; Josh Freeman; Julia Howard; Juliet Elvy; Lauren Jelly; Mary DeAlmeida; Matt Blakiston; Matt Storey; Matthew Rogers; Max Bloomfield; Michael Addidle; Michelle Balm; Sally Roberts; Sarah Jefferies; Sharmini Muttaiyah; Susan Morpeth; Susan Taylor; Timothy Blackmore; Vani Sathyendran; Veronica Playle; Virginia Hope; Xiaoyun Ren |
| EPI_ISL_470840, EPI_ISL_470845, EPI_ISL_470846, EPI_ISL_470849, EPI_ISL_470856, EPI_ISL_470863, EPI_ISL_470864, EPI_ISL_470865, EPI_ISL_470866, EPI_ISL_470867, EPI_ISL_470868, EPI_ISL_470869, EPI_ISL_470872 | | | |
| see above | PathWest Laboratory Medicine WA | PathWest Laboratory Medicine WA | Avram Levy; Chisha Sikazwe; David Smith and David Speers; Jurissa Lang |
| EPI_ISL_512741, EPI_ISL_512743, EPI_ISL_512755, EPI_ISL_596693, EPI_ISL_596697, EPI_ISL_596706, EPI_ISL_596723, EPI_ISL_596724, EPI_ISL_596744, EPI_ISL_596745, EPI_ISL_596749, EPI_ISL_596754, EPI_ISL_596778, EPI_ISL_596830, EPI_ISL_596835, EPI_ISL_596845, EPI_ISL_596848, EPI_ISL_596853, EPI_ISL_596855, EPI_ISL_596868, EPI_ISL_596879, EPI_ISL_672631, EPI_ISL_672632, EPI_ISL_672633, EPI_ISL_672634, EPI_ISL_672635, EPI_ISL_672654, EPI_ISL_708748 | | | |
| see above | PathWest Laboratory Medicine WA | PathWest Laboratory Medicine WA Microbial Surveillance Unit | PathWest Laboratory Medicine WA Microbial Surveillance Unit |
| EPI_ISL_684128, EPI_ISL_684132, EPI_ISL_684140, EPI_ISL_684141, EPI_ISL_684142, EPI_ISL_684143, EPI_ISL_684144, EPI_ISL_684145, EPI_ISL_684146, EPI_ISL_684156, EPI_ISL_684158, EPI_ISL_684164, EPI_ISL_684165, EPI_ISL_684168, EPI_ISL_684169, EPI_ISL_684170, EPI_ISL_684171, EPI_ISL_684172, EPI_ISL_684693, EPI_ISL_684736, EPI_ISL_685203, EPI_ISL_685238, EPI_ISL_685647, EPI_ISL_685695, EPI_ISL_685696, EPI_ISL_686021, EPI_ISL_686038, EPI_ISL_688757, EPI_ISL_688758, EPI_ISL_688759, EPI_ISL_690845, EPI_ISL_690847, EPI_ISL_690848, EPI_ISL_690850, EPI_ISL_690987, EPI_ISL_690988, EPI_ISL_898565, EPI_ISL_901221, EPI_ISL_901222, EPI_ISL_901224 | | | |
| see above | Pathogen Genomics Center, National Institute of Infectious Diseases | Pathogen Genomics Center, National Institute of Infectious Diseases | Kentaro Itokawa; Makoto Kuroda; Masanori Hashino; Rina Tanaka; Tsuyoshi Sekizuka |
| EPI_ISL_437459, EPI_ISL_437460, EPI_ISL_437461, EPI_ISL_437462, EPI_ISL_437465, EPI_ISL_437466, EPI_ISL_437467, EPI_ISL_437484, EPI_ISL_437495, EPI_ISL_437757, EPI_ISL_437759, EPI_ISL_437760, EPI_ISL_437761, EPI_ISL_678042, EPI_ISL_678080, EPI_ISL_678141 | | | |
| see above | Pathogen Genomics Lab King Abdullah University of Science and Technology(KAUST) | Pathogen Genomics Lab King Abdullah University of Science and Technology(KAUST) | Abdulaziz Alahmadi; Afrah Alsomali; Amit Kumar Subudhi; Amit Subudhi; Anwar Hashem; Arnab Pain; Asim Khogeer; Fadwa Alofi; Fathia Ben Rached; Fathia Ben-Rached; Fathia Rached; Jumana Taha; Kahled Algithami; Khaled Algithami; Naif Almontashiri; Raeece Naeem; Rahul P Salunke; Rahul Salunke; Sara Mfarrej; Sharif Hala |
| EPI_ISL_2976233 | Pathologists' Regional Laboratory | Idaho Bureau of Laboratories | "R. Beukelman; Aimee Ceniseros; Christian Loera; Christopher Ball"; Matthew Charles Burns; Robert L. Voermans |
| EPI_ISL_451552 | Pathology North - NSW Health Pathology | NSW Health Pathology - Institute of Clinical Pathology and Medical Research; Westmead Hospital; University of Sydney | CIDM-PH et al. |
| EPI_ISL_407894, EPI_ISL_407896, EPI_ISL_410717, EPI_ISL_410718, EPI_ISL_417031 | Pathology Queensland | Public Health Virology Laboratory | Alyssa Pyke; Amanda De Jong; Andrew Van Den Hurk; Ben Huang; Bixing Huang; Carmel Taylor; David Warrilow; Doris Genge; Elisabeth Gamez; Fred Hewitson; Ian Maxwell Mackay; Inga Sultana; Jamie McMahon; Jean Barcelon; Judy Northill; Mitchell Finger; Natalie Simpson; Neelima Nair; Peter Burtonclay; Peter Moore; Sarah Wheatley; Sean Moody; Sonja Hall-Mendelin; Timothy Gardam; and Frederick Moore; and Frederick Moore. |
| EPI_ISL_451608 | Pathology Sydney South West - NSW Health Pathology | NSW Health Pathology - Institute of Clinical Pathology and Medical Research; Westmead Hospital; University of Sydney | CIDM-PH et al. |
| EPI_ISL_451494, EPI_ISL_451496, EPI_ISL_451506, EPI_ISL_451513, EPI_ISL_451515, EPI_ISL_451516, EPI_ISL_451522, EPI_ISL_451557, EPI_ISL_451580, EPI_ISL_455044, EPI_ISL_455045, EPI_ISL_498748, EPI_ISL_498749, EPI_ISL_593662 | | | |
| see above | Pathology West - NSW Health Pathology | NSW Health Pathology - Institute of Clinical Pathology and Medical Research; Westmead Hospital; University of Sydney | CIDM-PH et al. |
| EPI_ISL_455585, EPI_ISL_455608 | Phramongkutklao Hospital | National Institute of Health. Department of medical Sciences, Ministry of Public Health, Thailand | Chittaganpitch; Malinee; Okada; Parnmen; Phuygun; Pilailuk; Siripaporn; Sittiporn; Sunthareeya; Thanadachakul; Thanutsapa; Waicharoen; Warawan; Wongboot |
| EPI_ISL_430841, EPI_ISL_434699, EPI_ISL_434702, EPI_ISL_434703, EPI_ISL_434704, EPI_ISL_434705, EPI_ISL_434706, EPI_ISL_455606, EPI_ISL_455607 | | | |
| see above | Praram 9 Hospital | National Institute of Health. Department of medical Sciences, Ministry of Public Health, Thailand | Chittaganpitch; Malinee; Okada; Parnmen; Phuygun; Pilailuk; Siripaporn; Sittiporn; Sunthareeya; Thanadachakul; Thanutsapa; Waicharoen; Warawan; Wongboot |
| EPI_ISL_404895 | Providence Regional Medical Center | Division of Viral Diseases, Centers for Disease Control and Prevention | C.R.; Gerber; Li, Y.; Lindstrom, S.; Lu, X.; Paden; Queen, K.; S.I.; Tao, Y.; Tong, S.; Zhang, J. |
| EPI_ISL_482299, EPI_ISL_482306, EPI_ISL_482318, EPI_ISL_482321, EPI_ISL_482324, EPI_ISL_482327, EPI_ISL_482449, EPI_ISL_482463 | | | |
| see above | Providence St. Joseph Health Molecular Genomics Laboratory | Providence St. Joseph Health Molecular Genomics Laboratory | Alexa K Dowdell; Brian D Piening; Carlo B Bifulco; Fred L Robinson; Mary Campbell |
| EPI_ISL_429815, EPI_ISL_482479 | Public Health Laboratory | National Microbiology Laboratory | Adel Malek; Anna Majer; Elsie Grudeski; Gary Van Domselaar; George Zahariadis; Grace Seo; Jennifer Tanner; Kristyn Burak; Laura Gilbert; Matthew Gilmour; Morag Graham; Natalie Knox; Nathalie Bastien; Philip Mabon; Rhiannon Huzarewich; Robert Needle; Russell Mandes; Shari Tyson; Timothy Booth; Yan Li; Yang Yu |
| EPI_ISL_593597, EPI_ISL_593598, EPI_ISL_593599, EPI_ISL_593600, EPI_ISL_593603, EPI_ISL_593611, EPI_ISL_593622, EPI_ISL_593624, EPI_ISL_593626, EPI_ISL_593627, EPI_ISL_639744, EPI_ISL_639747, EPI_ISL_639748, EPI_ISL_639749, EPI_ISL_639753, EPI_ISL_639754, EPI_ISL_639756, EPI_ISL_639766, EPI_ISL_639771, EPI_ISL_639774, EPI_ISL_639776, EPI_ISL_639779, EPI_ISL_639781, EPI_ISL_639782, EPI_ISL_639783, EPI_ISL_639794, EPI_ISL_639799, EPI_ISL_639800, EPI_ISL_639801, EPI_ISL_639803, EPI_ISL_639807, EPI_ISL_639815, EPI_ISL_693251, EPI_ISL_693258, EPI_ISL_693261, EPI_ISL_693266, EPI_ISL_693273, EPI_ISL_693281, EPI_ISL_693288, EPI_ISL_693290, EPI_ISL_693293, EPI_ISL_693295, EPI_ISL_849664, EPI_ISL_849666, EPI_ISL_849668, EPI_ISL_849688, EPI_ISL_849708, EPI_ISL_849711, EPI_ISL_849712, EPI_ISL_849714, EPI_ISL_849715, EPI_ISL_849716 | | | |
| see above | Public Health Virology Laboratory, Forensic and Scientific Services (PHV-FSS) | Public Health Virology Laboratory, Forensic and Scientific Services (PHV-FSS) | Son Nguyen et al |
| EPI_ISL_962181 | Public Health Virology-Forensic and Scientific Services (PHV-FSS) | Public Health Virology-Forensic and Scientific Services (PHV-FSS) | Son Nguyen et al |
| EPI_ISL_512785, EPI_ISL_512786 | Public Health, United States Air Force School of Aerospace Medicine | Public Health, United States Air Force School of Aerospace Medicine | A.C.; A.K.; A.W.; B.C.; C.R.; Chapleau; Connors; E.A.; Fries; J.R.; Javorina; Lambert; Macias; Meyer; Purves; R.R.; S.M.; Starr |

| | | | |
|---|---|---|---|
| EPI_ISL_434541, EPI_ISL_434543, EPI_ISL_434547 | Puerto Rico Department of Health | Centers for Disease Control and Prevention, Dengue Branch | Betzabel Flores; Chaney Kalinich; Fabiola Cruz; Gilberto A. Santiago; Glenda Gonzalez; Jessica I. Falcon; Jorge L. Munoz-Jordan; Joseph Fauver; Keyla Charriez; Nathan Grubaugh |
| EPI_ISL_579516, EPI_ISL_579570, EPI_ISL_579572 | QEII Health Sciences Centre | National Microbiology Laboratory (NML) | Anna Majer; Anneliese Landgraff; CanCOGeN's metadata curation team; Darian Hole; Elsie Grudeski; Gary Van Domselaar; Grace Seo; Jason LeBlanc; Jennifer Tanner; Morag Graham; Natalie Knox; Nathalie Bastien; Philip Mabon; Public Health Agency of Canada's CanCOGeN team; Rhiannon Huzarewich; Russell Mandes; Shari Tyson; Timothy Booth; Todd Hatchette; Yan Li |
| EPI_ISL_425564, EPI_ISL_425636 | Queens Medical Centre, Clinical Microbiology Department / DeepSeq Nottingham | COVID-19 Genomics UK (COG-UK) Consortium | Christopher Moore; Fei Sang; Gemma Clark; Hannah Howson-Wells; Johnny Debebe; Jonathan Ball; Joseph Chappell; Manjinder Khakh; Matthew Carlisle; Matthew Loose; Nadine Holmes; Patrick McClure; Theocharis Tsoleridis; Victoria Wright; Wendy Smith |
| EPI_ISL_530261, EPI_ISL_530262, EPI_ISL_530268, EPI_ISL_530273 | Queensland Health Forensic and Scientific Services, Public Health Virology | Public Health Virology Laboratory, Forensic and Scientific Services, Queensland Health | Son Nguyen et al |
| EPI_ISL_454692, EPI_ISL_454693, EPI_ISL_467814, EPI_ISL_467816, EPI_ISL_467817, EPI_ISL_467829, EPI_ISL_467831, EPI_ISL_467834, EPI_ISL_467836, EPI_ISL_467840, EPI_ISL_467842, EPI_ISL_467843, EPI_ISL_467845, EPI_ISL_467846, EPI_ISL_467847, EPI_ISL_467848, EPI_ISL_467849, EPI_ISL_467850, EPI_ISL_467855, EPI_ISL_467862, EPI_ISL_467871, EPI_ISL_467890, EPI_ISL_467891, EPI_ISL_467897, EPI_ISL_467903, EPI_ISL_467904, EPI_ISL_467909, EPI_ISL_467910, EPI_ISL_467911, EPI_ISL_467917, EPI_ISL_467919, EPI_ISL_467920, EPI_ISL_467921, EPI_ISL_467927, EPI_ISL_468563, EPI_ISL_468565, EPI_ISL_468567, EPI_ISL_468570, EPI_ISL_468590, EPI_ISL_571019, EPI_ISL_571031, EPI_ISL_571034, EPI_ISL_571040, EPI_ISL_571053, EPI_ISL_571056, EPI_ISL_571107, EPI_ISL_571153, EPI_ISL_571162, EPI_ISL_571171, EPI_ISL_571186, EPI_ISL_571270, EPI_ISL_571277, EPI_ISL_571286, EPI_ISL_571287, EPI_ISL_571288, EPI_ISL_571289, EPI_ISL_571290, EPI_ISL_571291, EPI_ISL_571293, EPI_ISL_571294, EPI_ISL_571295, EPI_ISL_571296, EPI_ISL_571297, EPI_ISL_571331, EPI_ISL_571336, EPI_ISL_571359, EPI_ISL_571361, EPI_ISL_571390, EPI_ISL_571391, EPI_ISL_571399, EPI_ISL_571401, EPI_ISL_571402, EPI_ISL_571403, EPI_ISL_571405, EPI_ISL_571406, EPI_ISL_571412, EPI_ISL_571417, EPI_ISL_571421, EPI_ISL_571422, EPI_ISL_571426, EPI_ISL_571427, EPI_ISL_571428, EPI_ISL_571433, EPI_ISL_571434, EPI_ISL_571491, EPI_ISL_571495, EPI_ISL_571504, EPI_ISL_571515, EPI_ISL_571520, EPI_ISL_571523, EPI_ISL_571524, EPI_ISL_571528, EPI_ISL_571530, EPI_ISL_571543, EPI_ISL_571548, EPI_ISL_571549, EPI_ISL_571561, EPI_ISL_571583, EPI_ISL_571590, EPI_ISL_571598, EPI_ISL_571604, EPI_ISL_571609, EPI_ISL_571655, EPI_ISL_571661, EPI_ISL_571685, EPI_ISL_571704, EPI_ISL_571708, EPI_ISL_571710, EPI_ISL_571712, EPI_ISL_571715, EPI_ISL_571733, EPI_ISL_571797, EPI_ISL_571801, EPI_ISL_571802, EPI_ISL_571806, EPI_ISL_571810, EPI_ISL_571812, EPI_ISL_571813, EPI_ISL_571823, EPI_ISL_571847, EPI_ISL_571902, EPI_ISL_571903, EPI_ISL_571905, EPI_ISL_571906, EPI_ISL_571918, EPI_ISL_571922, EPI_ISL_571927, EPI_ISL_571939, EPI_ISL_571958, EPI_ISL_571961, EPI_ISL_571992, EPI_ISL_572009, EPI_ISL_572021, EPI_ISL_572036, EPI_ISL_572038, EPI_ISL_572052, EPI_ISL_572055, EPI_ISL_572056, EPI_ISL_572062, EPI_ISL_572063, EPI_ISL_572071, EPI_ISL_572074, EPI_ISL_572083, EPI_ISL_572086, EPI_ISL_572093, EPI_ISL_572096, EPI_ISL_572131, EPI_ISL_572175, EPI_ISL_572181, EPI_ISL_603764, EPI_ISL_603766, EPI_ISL_603767, EPI_ISL_603773, EPI_ISL_603780, EPI_ISL_603781, EPI_ISL_603784, EPI_ISL_603796, EPI_ISL_603798, EPI_ISL_603811, EPI_ISL_603813, EPI_ISL_603816, EPI_ISL_603818, EPI_ISL_603823, EPI_ISL_603826, EPI_ISL_603832, EPI_ISL_603834, EPI_ISL_603836, EPI_ISL_603837, EPI_ISL_603842, EPI_ISL_603847, EPI_ISL_603848, EPI_ISL_603851, EPI_ISL_603855, EPI_ISL_603857, EPI_ISL_603860, EPI_ISL_603861, EPI_ISL_603868, EPI_ISL_603872, EPI_ISL_603883, EPI_ISL_603884, EPI_ISL_603889, EPI_ISL_603890, EPI_ISL_603896, EPI_ISL_603908, EPI_ISL_603913, EPI_ISL_603921, EPI_ISL_603924, EPI_ISL_603929, EPI_ISL_603937, EPI_ISL_603940, EPI_ISL_603969, EPI_ISL_604012, EPI_ISL_604014, EPI_ISL_604015, EPI_ISL_604024, EPI_ISL_604025, EPI_ISL_604027, EPI_ISL_604032, EPI_ISL_604053, EPI_ISL_604056, EPI_ISL_604093, EPI_ISL_604105, EPI_ISL_604118, EPI_ISL_604119, EPI_ISL_604137, EPI_ISL_604143, EPI_ISL_604144, EPI_ISL_604146, EPI_ISL_604148, EPI_ISL_604151, EPI_ISL_604158, EPI_ISL_604159, EPI_ISL_604163, EPI_ISL_604168, EPI_ISL_604189, EPI_ISL_604196, EPI_ISL_604197, EPI_ISL_604199, EPI_ISL_604201, EPI_ISL_604202, EPI_ISL_604205, EPI_ISL_604210, EPI_ISL_604211, EPI_ISL_604214, EPI_ISL_604217, EPI_ISL_604222, EPI_ISL_604225, EPI_ISL_604226, EPI_ISL_604263, EPI_ISL_604279, EPI_ISL_604286, EPI_ISL_604290, EPI_ISL_604291, EPI_ISL_604295, EPI_ISL_604304, EPI_ISL_604306, EPI_ISL_604308, EPI_ISL_604310, EPI_ISL_604311, EPI_ISL_604319, EPI_ISL_604326, EPI_ISL_604328, EPI_ISL_604335, EPI_ISL_604361, EPI_ISL_604376, EPI_ISL_604378, EPI_ISL_604390, EPI_ISL_604394, EPI_ISL_604395, EPI_ISL_604398, EPI_ISL_604399, EPI_ISL_604400, EPI_ISL_604415, EPI_ISL_604421, EPI_ISL_604423, EPI_ISL_604429, EPI_ISL_604445, EPI_ISL_604446, EPI_ISL_604447, EPI_ISL_604449, EPI_ISL_604450, EPI_ISL_604451, EPI_ISL_604454, EPI_ISL_604455, EPI_ISL_604465, EPI_ISL_604486, EPI_ISL_604492, EPI_ISL_604499, EPI_ISL_604509, EPI_ISL_604511, EPI_ISL_604514, EPI_ISL_604527, EPI_ISL_604529, EPI_ISL_604530, EPI_ISL_604531, EPI_ISL_604537, EPI_ISL_604538, EPI_ISL_604543, EPI_ISL_604551, EPI_ISL_604552, EPI_ISL_604557, EPI_ISL_604578, EPI_ISL_604583, EPI_ISL_604584, EPI_ISL_604587, EPI_ISL_604589, EPI_ISL_604592, EPI_ISL_604593, EPI_ISL_604603, EPI_ISL_604606, EPI_ISL_604607, EPI_ISL_604613, EPI_ISL_604614, EPI_ISL_604617, EPI_ISL_604618, EPI_ISL_604620, EPI_ISL_604622, EPI_ISL_604623, EPI_ISL_604624, EPI_ISL_604625, EPI_ISL_604626, EPI_ISL_604627, EPI_ISL_604629, EPI_ISL_604630, EPI_ISL_604641, EPI_ISL_604649, EPI_ISL_604652, EPI_ISL_604653, EPI_ISL_604673, EPI_ISL_604680, EPI_ISL_604683, EPI_ISL_604684, EPI_ISL_604685, EPI_ISL_604687, EPI_ISL_604689, EPI_ISL_604690, EPI_ISL_604691, EPI_ISL_604692, EPI_ISL_604693, EPI_ISL_604694, EPI_ISL_604695, EPI_ISL_604696, EPI_ISL_604697, EPI_ISL_604716, EPI_ISL_604733, EPI_ISL_604734, EPI_ISL_604744, EPI_ISL_604745, EPI_ISL_604748, EPI_ISL_604749, EPI_ISL_604751, EPI_ISL_604752, EPI_ISL_604757, EPI_ISL_604775, EPI_ISL_604776, EPI_ISL_604777, EPI_ISL_604778, EPI_ISL_604779, EPI_ISL_604780, EPI_ISL_604781, EPI_ISL_604785, EPI_ISL_604837, EPI_ISL_604839, EPI_ISL_604848, EPI_ISL_604849, EPI_ISL_604850, EPI_ISL_604852, EPI_ISL_604853, EPI_ISL_604854, EPI_ISL_604857, EPI_ISL_604860, EPI_ISL_604865, EPI_ISL_604869, EPI_ISL_604871, EPI_ISL_604876, EPI_ISL_604877, EPI_ISL_604881, EPI_ISL_604890, EPI_ISL_604898, EPI_ISL_604902, EPI_ISL_604904, EPI_ISL_604909, EPI_ISL_604931, EPI_ISL_604939, EPI_ISL_604953, EPI_ISL_604954, EPI_ISL_604969, EPI_ISL_604972, EPI_ISL_604973, EPI_ISL_604976, EPI_ISL_604978, EPI_ISL_604979, EPI_ISL_604988, EPI_ISL_604993, EPI_ISL_604998, EPI_ISL_604999, EPI_ISL_605003, EPI_ISL_605005, EPI_ISL_605012, EPI_ISL_605015, EPI_ISL_605016, EPI_ISL_605020, EPI_ISL_605021, EPI_ISL_605025, EPI_ISL_605033, EPI_ISL_605037, EPI_ISL_605047, EPI_ISL_605057, EPI_ISL_661253 | | | |
| see above | Quest Diagnostics | Quest Diagnostics | Anderson; Anderson, B.; B.P.; D.F.; Gerasimova, A.; Grover, D.; Hua, M.; K.E.; Kagan; Lacbawan, F.; Liu Y.; Livingston; Owen, R.; R.M.; R.M. and Owen, R.; Rosenthal; S.H.; Shalhout |
| EPI_ISL_416477 | R. G. Lugar Center for Public Health Research, National Center for Disease Control and Public Health (NCDC) of Georgia. | R. G. Lugar Center for Public Health Research, National Center for Disease Control and Public Health (NCDC) of Georgia. | Adam Kotorashvili; Amiran Gamkrelidze.; Ana Papkiauri; Ann Machablishvili; Anna Kasradze; Davit Tsaguria; Ekaterine Khmaladze; Ekaterine Zangaladze; Ekaterine Zhghenti; Giorgi Tomashvili; Gvantsa Brachveli; Gvantsa Chanturia; Irma Burjanadze; Ketevan Sidamonidze; Khatuna Zakhashvili; Lela Sabadze; Lela Urushadze; Magda Dgebuadze; Maia Alkhazashvili; Mari Gavashelidze; Mariam Zakalashvili; Marine Murtskhvaladze; Meri Pantsulaia; Nato Kotaria; Nino Berishvili; Paata Imnadze; Roena Sukhiashvili; Tamar Jashiashvili; Tata Imnadze; Tea Tevdoradze |
| EPI_ISL_423040, EPI_ISL_429164, EPI_ISL_429165, EPI_ISL_429166, EPI_ISL_429167, EPI_ISL_429168, EPI_ISL_429170, EPI_ISL_429171, EPI_ISL_429173, EPI_ISL_429176, EPI_ISL_429177, EPI_ISL_447002, EPI_ISL_447005, EPI_ISL_447007, EPI_ISL_450723, EPI_ISL_455908, EPI_ISL_455909, EPI_ISL_455911, EPI_ISL_455912, EPI_ISL_455914, EPI_ISL_455916, EPI_ISL_455924, EPI_ISL_455925, EPI_ISL_455926, EPI_ISL_455928, EPI_ISL_455929, EPI_ISL_455930, EPI_ISL_455931, EPI_ISL_455932, EPI_ISL_455933, EPI_ISL_455935, EPI_ISL_455936, EPI_ISL_455937, EPI_ISL_455938, EPI_ISL_455939, EPI_ISL_455940, EPI_ISL_455941, EPI_ISL_455942, EPI_ISL_455944, EPI_ISL_455945, EPI_ISL_455946, EPI_ISL_455947, EPI_ISL_455948, EPI_ISL_498263, EPI_ISL_498265, EPI_ISL_512848, EPI_ISL_512851, EPI_ISL_512852, EPI_ISL_512853, EPI_ISL_512854, EPI_ISL_512856, EPI_ISL_512857, EPI_ISL_512858, EPI_ISL_512859, EPI_ISL_512860, EPI_ISL_512863, EPI_ISL_512865, EPI_ISL_512868, EPI_ISL_512870, EPI_ISL_512871, EPI_ISL_568581, EPI_ISL_568582, EPI_ISL_568583, EPI_ISL_568586, EPI_ISL_693334, EPI_ISL_693335, EPI_ISL_693337, EPI_ISL_693338, EPI_ISL_693339, EPI_ISL_693340, EPI_ISL_693341, EPI_ISL_693342, EPI_ISL_693344, EPI_ISL_693346, EPI_ISL_693347, EPI_ISL_693348, EPI_ISL_693349, EPI_ISL_693350, EPI_ISL_693351, EPI_ISL_693352, EPI_ISL_693353, EPI_ISL_693354, EPI_ISL_693355, EPI_ISL_693356, EPI_ISL_693358, EPI_ISL_693359, EPI_ISL_693360, EPI_ISL_693362, EPI_ISL_693363, EPI_ISL_693365, EPI_ISL_693366, EPI_ISL_693367, EPI_ISL_693368, EPI_ISL_693369, EPI_ISL_693370, EPI_ISL_693372, EPI_ISL_693373, EPI_ISL_693374, EPI_ISL_693375, EPI_ISL_693376, EPI_ISL_693377, EPI_ISL_693378, EPI_ISL_693379, EPI_ISL_693380, EPI_ISL_693385 | | | |
| see above | Ramathibodi Hospital | COVID-19 Network Investigations (CONI) Alliance | Angkana Huang; Anthony R. Jones; Arporn Wangwiwatsin; Bhakbhoom Panthan; Chonticha Klungtong; Ekawat Pasomsub; Elizabeth Batty; Insee Sensorn; Janjira Thaipadungpanit; Khajohn Joonsalak; Kingkan Rakmanee; Krittikorn Kumpornsin; Namfon Kotanan; Poramate Jiaranai; Stefan Fernandez; Thanat Chookajorn; Theerarat Kochakarn; Treewat Watthanachockchai; Wasun Chantratita; Wudtichai Manasatienkij |
| EPI_ISL_434700, EPI_ISL_455604 | Ramkhamhaeng Hospital | National Institute of Health. Department of medical Sciences, Ministry of Public Health, Thailand | Chittaganpitch; Malinee; Okada; Parnmen; Phuygun; Pilailuk; Siripaporn; Sittiporn; Sunthareeya; Thanadachakul; Thanutsapa; Waicharoen; Warawan; Wongboot |
| EPI_ISL_454417, EPI_ISL_454418 | Research and Experiment Center, Meizhou People Hospital | Research and Experiment Center, Meizhou People Hospital | Guo, X.; L. and Yu, Z.; Zeng |
| EPI_ISL_2735785 | Respiratory Virus Unit, Microbiology Services Colindale, Public Health England | COVID-19 Genomics UK (COG-UK) Consortium | PHE Covid Sequencing Team |
| EPI_ISL_407071, EPI_ISL_407073, EPI_ISL_420524, EPI_ISL_423169, EPI_ISL_423179, EPI_ISL_423180, EPI_ISL_423352, EPI_ISL_423492, EPI_ISL_423506, EPI_ISL_423523, EPI_ISL_423859, EPI_ISL_423918, EPI_ISL_423976, EPI_ISL_424127, EPI_ISL_464744, EPI_ISL_465052, EPI_ISL_465897, EPI_ISL_466032 | | | |
| see above | Respiratory Virus Unit, Microbiology Services Colindale, Public Health England | Respiratory Virus Unit, Microbiology Services Colindale, Public Health England | Angie Lackenby; Joanna Ellis; Jonathan Hubb; Kirstin Edwards; Leena Bhaw; Maria Zambon; Maria Zambon.; Monica Galiano; Omolola Akinbami; PHE Covid Sequencing Team; Richard Myers; Shahjahan Miah; Steven Platt; Tiina Talts |
| EPI_ISL_682298 | Respiratory virus Laboratory, Chinese Academy of Medical Science | Respiratory virus Laboratory, Chinese Academy of Medical Science | Chen; Li, J.; Y. and Liu, L.; Zhen, H. |
| EPI_ISL_631816 | Richmond University Medical Center | New York City Public Health Laboratory | Jade Wang; et al. |
| EPI_ISL_436098 | Royal Brisbane and Women's Hospital | Public Health Virology Laboratory, Forensic and Scientific Services, Queensland Health | Alyssa Pyke; Amanda De Jong; Andrew Van Den Hurk; Bixing Huang; Carmel Taylor; David Warrilow; Doris Genge; Elisabeth Gamez; Glen Hewitson; Ian Maxwell Mackay; Inga Sultana and Frederick Moore; Jamie McMahon; Jean Barcelon; Judy Northill; Lisa Leckie; Mitchell Finger; Natalie Simpson; Neelima Nair; Peter Burtonclay; Peter Moore; Sarah Wheatley; Sean Moody; Sonja Hall-Mendelin |
| EPI_ISL_426634, EPI_ISL_426635 | Royal Darwin Hospital Pathology | Microbiological Diagnostic Unit Public Health Laboratory and Victorian Infectious Diseases Reference Laboratory, Doherty Institute | Caly L.; Druce J.; Meumann, E.; Sait, M.; Schultz M.; Seemann T.; Sherry, N. |
| EPI_ISL_430497 | Royal Darwin Hospital Pathology | Microbiological Diagnostic Unit Public Health Laboratory and Victorian Infectious Diseases Reference Laboratory, The Peter Doherty Institute for Infection and Immunity | Caly L.; Druce J.; Meumann, E.; Sait, M.; Schultz M.; Seemann T.; Sherry, N. |
| EPI_ISL_522570, EPI_ISL_522572, EPI_ISL_522579, EPI_ISL_522582, EPI_ISL_522583, EPI_ISL_522584, EPI_ISL_522588, EPI_ISL_522590, EPI_ISL_522606, EPI_ISL_522611, EPI_ISL_522620, EPI_ISL_522636, EPI_ISL_522644, EPI_ISL_522652, EPI_ISL_522653, EPI_ISL_522721, EPI_ISL_522727, EPI_ISL_522730, EPI_ISL_522735, EPI_ISL_522736, EPI_ISL_522737, EPI_ISL_522738, EPI_ISL_522744, EPI_ISL_522745, EPI_ISL_522746, EPI_ISL_522753, EPI_ISL_522758, EPI_ISL_522763, EPI_ISL_522764, EPI_ISL_522769 | | | |
| see above | Royal Hobart Hospital Microbiology Department | MDU-PHL | Cooley L.; M.B.; Sait M.; Schultz; Seemann T.; Sherry N.; van Haeften R. |
| EPI_ISL_3024688, EPI_ISL_3024689, EPI_ISL_3024690, EPI_ISL_3024692, EPI_ISL_3024694, EPI_ISL_3024695, EPI_ISL_3024697 | | | |
| see above | Rush University Medical Center | RIPHL at Rush University Medical Center | Felix Araujo Perez; Jinal Patel; Kevin Kunstman; Laura Furtado; Marieta Hyde; Mary Hayden; Max Kolton; Stefan Green |
| EPI_ISL_451080, EPI_ISL_451081, EPI_ISL_451083, EPI_ISL_451086, EPI_ISL_451096, EPI_ISL_451104, EPI_ISL_451134, EPI_ISL_451137, EPI_ISL_451140, EPI_ISL_451141, EPI_ISL_451148, EPI_ISL_467985, EPI_ISL_467992, EPI_ISL_468006, EPI_ISL_468008, EPI_ISL_468015, EPI_ISL_468018, EPI_ISL_468022, EPI_ISL_468030, EPI_ISL_468033, EPI_ISL_468034, EPI_ISL_468042, EPI_ISL_483077, EPI_ISL_483079, EPI_ISL_483080, EPI_ISL_483082, EPI_ISL_483083, EPI_ISL_483085, EPI_ISL_483088, EPI_ISL_483090, EPI_ISL_483091, EPI_ISL_483094, EPI_ISL_483100, EPI_ISL_483101, EPI_ISL_483103, EPI_ISL_483107, EPI_ISL_483108, EPI_ISL_483110, EPI_ISL_483116, EPI_ISL_483117, EPI_ISL_483118, EPI_ISL_483120, EPI_ISL_483122, EPI_ISL_483127, EPI_ISL_483129, EPI_ISL_483130, EPI_ISL_483132, EPI_ISL_483133, EPI_ISL_483135, EPI_ISL_483137, EPI_ISL_483138, EPI_ISL_492109, EPI_ISL_492118, EPI_ISL_492119, EPI_ISL_492120, EPI_ISL_492134, EPI_ISL_492136, EPI_ISL_492138, EPI_ISL_492140, EPI_ISL_492141, EPI_ISL_492143, EPI_ISL_492144, EPI_ISL_492149, EPI_ISL_492150, EPI_ISL_492156, EPI_ISL_492157, EPI_ISL_492158, EPI_ISL_492159, EPI_ISL_492163, EPI_ISL_492165, EPI_ISL_492166, EPI_ISL_492169, EPI_ISL_492172, EPI_ISL_508123, EPI_ISL_508131, EPI_ISL_508138, EPI_ISL_508143, EPI_ISL_508144 | | | |
| see above | SA Pathology | SA Pathology | Chuan Kok Lim; Geoff Higgins; Ivan Bastian; Lex Leong; Mark Turra |
| EPI_ISL_491097, EPI_ISL_491098, EPI_ISL_491099, EPI_ISL_491100, EPI_ISL_491101, EPI_ISL_491102, EPI_ISL_491103, EPI_ISL_491104 | | | |
| see above | SC Department of Health and Environmental Control | SC Department of Health and Environmental Control | Flores, H. |
| EPI_ISL_424841, EPI_ISL_424842, EPI_ISL_424888, EPI_ISL_424902, EPI_ISL_424903, EPI_ISL_424904 | SC Dept of Health and Env. Control-Bureau of Laboratories | Pathogen Discovery, Respiratory Viruses Branch, Division of Viral Diseases, Centers for Disease Control and Prevention | Alison S. Laufer Halpin; Anna Uehara; Christopher A. Elkins; Clinton R. Paden; Haibin Wang; Jing Zhang; Krista Queen; Mary S. Keckler; Rachel Marine; Suxiang Tong; Yan Li; Ying Tao |
| EPI_ISL_511357, EPI_ISL_511359 | SYNLAB | Instituto Nacional de Saude (INSA) | Borges et al |
| EPI_ISL_511221, EPI_ISL_511240, EPI_ISL_511251 | SYNLAB | Instituto Nacional de Saude (INSA) and Instituto Gulbenkian de Ciencia (IGC) | Borges et al |
| EPI_ISL_479805, EPI_ISL_479806, EPI_ISL_479807, EPI_ISL_479931, EPI_ISL_479932, EPI_ISL_684148 | Saitama Prefectural Institute of Public Health | Pathogen Genomics Center, National Institute of Infectious Diseases | Hajime Kamiya; Hayato Ehara; Kentaro Itokawa; Makoto Kuroda; Masanori Hashino; Motoi Suzuki; Rina Tanaka; Tsuyoshi Sekizuka |
| EPI_ISL_479925, EPI_ISL_479926, EPI_ISL_479927, EPI_ISL_480065, EPI_ISL_684147 | Sakai City Institute of Public Health | Pathogen Genomics Center, National Institute of Infectious Diseases | Hajime Kamiya; Kentaro Itokawa; Makoto Kuroda; Masanori Hashino; Motoi Suzuki; Rina Tanaka; Tatsuya Miyoshi; Tsuyoshi Sekizuka |
| EPI_ISL_467950, EPI_ISL_467951, EPI_ISL_467954, EPI_ISL_467956, EPI_ISL_467958, EPI_ISL_467964, EPI_ISL_467968, EPI_ISL_467970, EPI_ISL_467973, EPI_ISL_467978, EPI_ISL_467982, EPI_ISL_494624 | | | |
| see above | San Diego County Public Health Laboratory | Andersen lab at Scripps Research | Brett Austin; Jovan Shephard; SEARCH Alliance San Diego with Tracy Basler |

| | | | |
|---|---|---|---|
| EPI_ISL_513876 | San Francisco Public Health Laboratory | Chan-Zuckerberg Biohub | CZB Cliahub Consortium |
| EPI_ISL_468506, EPI_ISL_468507, EPI_ISL_468508, EPI_ISL_468509, EPI_ISL_468511, EPI_ISL_468512, EPI_ISL_468514, EPI_ISL_468515, EPI_ISL_468516, EPI_ISL_468517, EPI_ISL_468518, EPI_ISL_468524, EPI_ISL_468528, EPI_ISL_468559 | | | |
| see above | San Joaquin County Public Health Lab | Chan-Zuckerberg Biohub | CZB Cliahub Consortium |
| EPI_ISL_542346 | San Matteo Hospital Pavia | Dep. Of Oncology and Hemato-Oncology University of Milan | Antonio Piralla; Carlo Federico Perno; Chiara Vismara; Claudia Alteri; Elisa Matarazzo; Fausto Baldanti; Federica Giardina; Federica Novazzi; Luna Colagrossi; Maria Antonello; Massimo Puoti; Monica Tallarita; Oscar Massimiliano Epis; Roberto Fumagalli; Silvia Renica; Stefano Gaiarsa; Valentino Costabile; Valeria Cento |
| EPI_ISL_435603, EPI_ISL_435611, EPI_ISL_435619, EPI_ISL_435625, EPI_ISL_435631, EPI_ISL_435640, EPI_ISL_671765, EPI_ISL_671768 | | | |
| see above | Santa Clara County Public Health Department | Chiu Laboratory, University of California, San Francisco | Brandon Bonin; Debra A. Wadford; Elsa Villarino; Scot Federman; Wei Gu; Xianding Deng; and Charles Y. Chiu |
| EPI_ISL_660108 | Santa Clara County Public Health Laboratory | Chan-Zuckerberg Biohub | CZB Cliahub Consortium |
| EPI_ISL_479833, EPI_ISL_479835 | Sapporo City Institute of Public Health | Pathogen Genomics Center, National Institute of Infectious Diseases | Asami Ohnishi; Hajime Kamiya; Kentaro Itokawa; Makoto Kuroda; Masanori Hashino; Motoi Suzuki; Rina Tanaka; Tsuyoshi Sekizuka |
| EPI_ISL_437563, EPI_ISL_445165 | Scripps Medical Laboratory | Andersen lab at Scripps Research | Ellen Stefanski; Ian Mchardy; SEARCH Alliance San Diego with Michael Quigley |
| EPI_ISL_413560, EPI_ISL_416460, EPI_ISL_416461, EPI_ISL_416462, EPI_ISL_416463, EPI_ISL_416464, EPI_ISL_416465, EPI_ISL_416466, EPI_ISL_417163, EPI_ISL_417164, EPI_ISL_417165, EPI_ISL_430113, EPI_ISL_430114, EPI_ISL_430116, EPI_ISL_430117, EPI_ISL_430118, EPI_ISL_430119, EPI_ISL_430122, EPI_ISL_430123, EPI_ISL_430124, EPI_ISL_430125, EPI_ISL_430126, EPI_ISL_430127, EPI_ISL_430129, EPI_ISL_430131, EPI_ISL_430132, EPI_ISL_430133, EPI_ISL_430134, EPI_ISL_430135, EPI_ISL_430137, EPI_ISL_430140, EPI_ISL_430141, EPI_ISL_430142, EPI_ISL_430144, EPI_ISL_430145, EPI_ISL_430146, EPI_ISL_430150, EPI_ISL_430151, EPI_ISL_430152, EPI_ISL_430153, EPI_ISL_430155, EPI_ISL_430156, EPI_ISL_495657, EPI_ISL_495663, EPI_ISL_575033 | | | |
| see above | Seattle Flu Study | Seattle Flu Study | Amanda Adler; Barry R. Lutz; Benjamin Pelle; Brian Hiatt; Caitlin R. Wolf; Chris D. Frazar; Chu et al; Chu etl al; Deborah A. Nickerson; Elisabeth Brandstetter; Geoff Melly; Helen Y. Chu; Janet A. Englund; Jay Shendure; Jeff Duchin; Jover Lee; Kairsten Fay; Karen Cowgill; Kirsten Lacombe; Lea M. Starita; Mark J. Rieder; Matthew Richardson; Matthew Thompson; Melissa Truong; Michael Boeckh; Michael Famulare; Misja Ilcisin; Peter D. Han; Philip Dykema; Romesh Gautom; Scott Lindquist; Stephanie Schrag; Thomas R. Sibley; Trevor Bedford |
| EPI_ISL_413456 | Seattle Flu Study, University of Washington Medical Center | Seattle Flu Study, University of Washington Medical Center | Chu et al |
| EPI_ISL_417924 | Secretaría de Salud Medellín | Instituto Nacional de Salud, Universidad Cooperativa de Colombia, Instituto Alexander von Humboldt, Imperial College-London, London School of Hygiene & Tropical Medicine | Astrid C. Flórez; Carlos Franco-Muñoz; Christian Julian Villabona-Arenas; Diana Marcela Walteros-Acero; Diego A. Álvarez-Díaz; Erika Ospitia; Gloria Puerto; Jose A. Usme-Ciro; Juliana Barbosa; Katherine Laiton-Donato; Liz Villabona-Arenas; Luz Dary Rodriguez; Mailyn A. Gonzalez; Marcela Mercado-Reyes; Martha Lucia Ospina Martinez; Nicolás D. Franco-Sierra; Sergio Gomez Rangel; Sussy Echeverria-Londoño; Zulma M. Cucunubá |
| EPI_ISL_414598 | Servicio Microbiologia, Hospital Clinico Universitario, Valencia | Sequencing and Bioinformatics Service and Molecular Epidemiology Research Group. FISABIO-Public Health. | David Navarro; Fernando Gonzalez-Candelas; Giuseppe D'Auria; Griselda De Marco; Maria Alma Bracho; Neris Garcia-Gonzalez |
| EPI_ISL_414496 | Servicio Microbiologia. Hospital Clinico Universitario. Valencia. | Sequencing and Bioinformatics Service. Molecular Epidemiology Laboratory. FISABIO-Public Health | David Navarro; Fernando Gonzalez-Candelas; Giuseppe D'Auria; Griselda De Marco; María Alma Bracho; Neris Garcia-Gonzalez |
| EPI_ISL_416485, EPI_ISL_417206 | Servicio de Microbiologia. Consorcio Hospital General Universitario de Valencia | Sequencing and Bioinformatics Service and Molecular Epidemiology Research Group. FISABIO-Public Health | Concepcion Gimeno; Fernando Gonzalez-Candelas; Giuseppe D'Auria; Griselda De Marco; Maria Alma Bracho; Maria Dolores Ocete; Neris Garcia-Gonzalez |
| EPI_ISL_537996, EPI_ISL_538000 | Servicio de Microbiologia. Hospital Arnau de Vilanova | SeqCOVID-SPAIN consortium/IBV(CSIC) | Fernando Gonzalez-Candelas and SeqCOVID-SPAIN consortium; Giuseppe D'Auria; Griselda De Marco; Inma Galán Vendrell; Ivan Ansari; Lidia Ruiz Roldan; Llúcia Martínez-Priego; Loreto Ferrús Abad; Maria Alma Bracho; Mariana Reyes-Prieto; Neris Garcia-Gonzalez; Paula Ruiz-Hueso; Sandra Carbo; Vicente Soriano Chirona; Victoria Dominguez |
| EPI_ISL_539223, EPI_ISL_539224, EPI_ISL_539232, EPI_ISL_539233, EPI_ISL_539236, EPI_ISL_539237, EPI_ISL_539238, EPI_ISL_539239 | | | |
| see above | Servicio de Microbiologia. Hospital Clinico Universitario de Valencia | SeqCOVID-SPAIN consortium/IBV(CSIC) | Beatriz Beamud; Concepcion Gimeno; Fernando Gonzalez-Candelas and SeqCOVID-SPAIN consortium; Giuseppe D'Auria; Griselda De Marco; Inma Galán Vendrell; Lidia Ruiz Roldan; Llúcia Martínez-Priego; Loreto Ferrús Abad; Maria Alma Bracho; Maria Dolores Ocete; Mariana Reyes-Prieto; Marta Pla Diaz; Neris Garcia-Gonzalez; Paula Ruiz-Hueso; Vicente Soriano Chirona |
| EPI_ISL_421520, EPI_ISL_436280, EPI_ISL_436284, EPI_ISL_436288, EPI_ISL_436296, EPI_ISL_436298, EPI_ISL_436303, EPI_ISL_436304, EPI_ISL_436306, EPI_ISL_436307, EPI_ISL_436311, EPI_ISL_436320, EPI_ISL_436332, EPI_ISL_436333, EPI_ISL_436334, EPI_ISL_436335, EPI_ISL_436338, EPI_ISL_436341, EPI_ISL_436346, EPI_ISL_436355, EPI_ISL_436364, EPI_ISL_436399, EPI_ISL_436401, EPI_ISL_436403, EPI_ISL_436404, EPI_ISL_436406, EPI_ISL_436410, EPI_ISL_447470, EPI_ISL_447481, EPI_ISL_447482, EPI_ISL_447484, EPI_ISL_447486, EPI_ISL_447490, EPI_ISL_447491, EPI_ISL_447495 | | | |
| see above | Servicio de Microbiologia. Hospital Clinico Universitario de Valencia | Sequencing and Bioinformatics Service and Molecular Epidemiology Research Group. FISABIO-Public Health | Beatriz Beamud; David Navarro; Eliseo Albert; Fernando Gonzalez-Candelas; Giuseppe 'Auria; Giuseppe D'Auria; Griselda De Marco; Inma Galán Vendrell; Ivan Ansari; Lidia Ruiz Roldan; Llúcia Martínez-Priego; Loreto Ferrús Abad; Maria Alma Bracho; Mariana Reyes-Prieto; Marta Pla Diaz; Neris Garcia-Gonzalez; Paula Ruiz-Hueso; Sandra Carbo; Vicente Soriano Chirona |
| EPI_ISL_455722, EPI_ISL_455741 | Servicio de Microbiologia. Hospital Clinico Universitario de Valencia | Sequencing and Bioinformatics Service and Molecular Epidemiology Research Group. FISABIO-Public Health, and SeqCOVID-Spain Consortium | David Navarro; Eliseo Albert; Fernando Gonzalez-Candelas; Giuseppe 'Auria; Griselda De Marco; Inma Galán Vendrell; Ivan Ansari; Lidia Ruiz Roldan; Llúcia Martínez-Priego; Loreto Ferrús Abad; Maria Alma Bracho; Mariana Reyes-Prieto; Neris Garcia-Gonzalez; Paula Ruiz-Hueso; Sandra Carbo; Vicente Soriano Chirona |
| EPI_ISL_436236, EPI_ISL_436241, EPI_ISL_436248, EPI_ISL_436253, EPI_ISL_436254, EPI_ISL_436256, EPI_ISL_436257, EPI_ISL_436263, EPI_ISL_436264, EPI_ISL_436373, EPI_ISL_436374, EPI_ISL_436377 | | | |
| see above | Servicio de Microbiologia. Hospital Universitario Doctor Peset | Sequencing and Bioinformatics Service and Molecular Epidemiology Research Group. FISABIO-Public Health | Beatriz Beamud; Fernando Gonzalez-Candelas; Giuseppe D'Auria; Griselda De Marco; Inma Galán Vendrell; Ivan Ansari; Jose Miguel Nogueira Coito; Juan Alberola Enguídanos; Juan Jose Camarena Miñana; Lidia Ruiz Roldan; Llúcia Martínez-Priego; Loreto Ferrús Abad; Maria Alma Bracho; Mariana Reyes-Prieto; Marta Pla Diaz; Neris Garcia-Gonzalez; Paula Ruiz-Hueso; Rosa González Pellicer; Sandra Carbo; Vicente Soriano Chirona |
| EPI_ISL_455735, EPI_ISL_455736 | Servicio de Microbiologia. Hospital_Arnau_de_Vilanova | Sequencing and Bioinformatics Service and Molecular Epidemiology Research Group. FISABIO-Public Health, and SeqCOVID-Spain Consortium | Fernando Gonzalez-Candelas; Giuseppe 'Auria; Griselda De Marco; Inma Galán Vendrell; Ivan Ansari; Lidia Ruiz Roldan; Llúcia Martínez-Priego; Loreto Ferrús Abad; Maria Alma Bracho; Mariana Reyes-Prieto; Neris Garcia-Gonzalez; Paula Ruiz-Hueso; Sandra Carbo; Vicente Soriano Chirona; Victoria Dominguez |
| EPI_ISL_1382903, EPI_ISL_1382916, EPI_ISL_1382918, EPI_ISL_1382919, EPI_ISL_1382920, EPI_ISL_1382921, EPI_ISL_1382922, EPI_ISL_1382923, EPI_ISL_1382924, EPI_ISL_1382948, EPI_ISL_1382954, EPI_ISL_1382955, EPI_ISL_1382957, EPI_ISL_1382960, EPI_ISL_1382974, EPI_ISL_1382995, EPI_ISL_1383002, EPI_ISL_1383009, EPI_ISL_1383018, EPI_ISL_1383022, EPI_ISL_1383025, EPI_ISL_1383034, EPI_ISL_1383045, EPI_ISL_1383070, EPI_ISL_1383082, EPI_ISL_1383087, EPI_ISL_1383098, EPI_ISL_1383099, EPI_ISL_1383100, EPI_ISL_1383101, EPI_ISL_1383113, EPI_ISL_1383115, EPI_ISL_1383121, EPI_ISL_1383123, EPI_ISL_1383125, EPI_ISL_1383132, EPI_ISL_1383169, EPI_ISL_1383173, EPI_ISL_1383178, EPI_ISL_1383185, EPI_ISL_1383202, EPI_ISL_1383209, EPI_ISL_1383216, EPI_ISL_1383219, EPI_ISL_1383229, EPI_ISL_1383232 | | | |
| see above | Servicio de Microbiología Clínica (Complejo Hospitalario de Navarra, Pamplona) | Centro de Secuenciación NASERTIC | Ana Miqueleiz; Ana Navascués; Carmen Ezpeleta Baquedano |
| EPI_ISL_538056, EPI_ISL_538061, EPI_ISL_538179, EPI_ISL_538182, EPI_ISL_538186, EPI_ISL_538193, EPI_ISL_538199, EPI_ISL_538200, EPI_ISL_538205, EPI_ISL_538215, EPI_ISL_538217, EPI_ISL_538230 | | | |
| see above | Servicio de Microbiología y Parasitología clínica. UCEIMP. Hospital Universitario Virgen del Rocío/IBIS/CSIC/US | SeqCOVID-SPAIN consortium/IBV(CSIC) | Guillermo Martín Gutiérrez; Javier Aznar Martín and SeqCOVID-SPAIN consortium; Lidia Gálvez Benitez; Verónica González Galán; Ángel Rodríguez Villodres |
| EPI_ISL_452545, EPI_ISL_452547, EPI_ISL_452548, EPI_ISL_452553, EPI_ISL_452557, EPI_ISL_452559, EPI_ISL_452561, EPI_ISL_452563, EPI_ISL_452565, EPI_ISL_452566, EPI_ISL_452568, EPI_ISL_452570, EPI_ISL_452581, EPI_ISL_452584, EPI_ISL_452588, EPI_ISL_452591, EPI_ISL_452594, EPI_ISL_452599, EPI_ISL_452600, EPI_ISL_452608 | | | |
| see above | Servicio de Microbiología y Parasitología clínica. UCEIMP. Hospital Universitario Virgen del Rocío/IBIS/CSIC/US. | SeqCOVID-SPAIN consortium/IBV(CSIC) | Guillermo Martín Gutiérrez; Javier Aznar Martín and SeqCOVID-SPAIN consortium; Lidia Gálvez Benitez; Verónica González Galán; Ángel Rodríguez Villodres |
| EPI_ISL_468766, EPI_ISL_468776, EPI_ISL_468779, EPI_ISL_468789, EPI_ISL_468790, EPI_ISL_468792, EPI_ISL_468796, EPI_ISL_468798, EPI_ISL_468802, EPI_ISL_468804, EPI_ISL_468817, EPI_ISL_468818, EPI_ISL_468821, EPI_ISL_468822, EPI_ISL_468825, EPI_ISL_468827, EPI_ISL_468834, EPI_ISL_468837, EPI_ISL_468845, EPI_ISL_468848, EPI_ISL_468855, EPI_ISL_468857 | | | |
| see above | Servicio de Microbiología, Hospital Miguel Servet, Zaragoza | SeqCOVID-SPAIN consortium/IBV(CSIC) | Alexander Tristancho Baró; Ana Milagro; Antonio Rezusta López; Nieves Martínez Cameo and SeqCOVID-SPAIN consortium; Yolanda Gracia Grataloup |
| EPI_ISL_654621, EPI_ISL_654623, EPI_ISL_654624, EPI_ISL_654637, EPI_ISL_654641, EPI_ISL_654654 | Servicio de Microbiología, Hospital Universitario Central de Asturias | SeqCOVID-SPAIN consortium/IBV(CSIC) | Cristián Castelló Abietar; Jose A. Boga; Marta Elena Álvarez-Argüelles; Santiago Melón and SeqCOVID-SPAIN consortium; Susana Rojo-Alba |
| EPI_ISL_582699, EPI_ISL_582709, EPI_ISL_582710, EPI_ISL_582712, EPI_ISL_582717, EPI_ISL_582720, EPI_ISL_582723, EPI_ISL_582725, EPI_ISL_582731, EPI_ISL_582736, EPI_ISL_582737, EPI_ISL_582745, EPI_ISL_582747, EPI_ISL_582761, EPI_ISL_582763, EPI_ISL_582764 | | | |
| see above | Servicio de Microbiología, Hospital Universitario Cruces, Bilbao | SeqCOVID-SPAIN consortium/IBV(CSIC) | Maitane Aranzamendi Zaldumbide and SeqCOVID-SPAIN consortium |
| EPI_ISL_467057, EPI_ISL_468954, EPI_ISL_468955, EPI_ISL_468960, EPI_ISL_468961, EPI_ISL_468965, EPI_ISL_468969, EPI_ISL_468973, EPI_ISL_468978, EPI_ISL_468979, EPI_ISL_468982, EPI_ISL_468985, EPI_ISL_468991, EPI_ISL_468994, EPI_ISL_468996, EPI_ISL_469001, EPI_ISL_469007, EPI_ISL_469011, EPI_ISL_541963, EPI_ISL_541966 | | | |
| see above | Servicio de Microbiología, Hospital Universitario Son Espases | SeqCOVID-SPAIN consortium/IBV(CSIC) | Antonio Oliver and SeqCOVID-SPAIN consortium; Carla López-Causapé; Jordi Reina; Jordi Reina y Antonio Oliver and SeqCOVID-SPAIN consortium |
| EPI_ISL_537608, EPI_ISL_539242, EPI_ISL_539289, EPI_ISL_539290, EPI_ISL_539293, EPI_ISL_539295, EPI_ISL_539296, EPI_ISL_539298 | | | |
| see above | Servicio de Microbiología. Consorcio Hospital General Universitario de Valencia | SeqCOVID-SPAIN consortium/IBV(CSIC) | Beatriz Beamud; David Navarro; Fernando Gonzalez-Candelas and SeqCOVID-SPAIN consortium; Giuseppe D'Auria; Griselda De Marco; Inma Galán Vendrell; Ivan Ansari; Lidia Ruiz Roldan; Llúcia Martínez-Priego; Loreto Ferrús Abad; Maria Alma Bracho; Mariana Reyes-Prieto; Marta Pla Diaz; Neris Garcia-Gonzalez; Paula Ruiz-Hueso; Sandra Carbo; Vicente Soriano Chirona |
| EPI_ISL_416483, EPI_ISL_416484, EPI_ISL_416487, EPI_ISL_417205, EPI_ISL_419676, EPI_ISL_419678, EPI_ISL_419679, EPI_ISL_419682, EPI_ISL_419683, EPI_ISL_419684, EPI_ISL_419685, EPI_ISL_419686, EPI_ISL_419687, EPI_ISL_419688, EPI_ISL_419690, EPI_ISL_420112, EPI_ISL_420113, EPI_ISL_420114, EPI_ISL_420115, EPI_ISL_420116, EPI_ISL_420117, EPI_ISL_420118, EPI_ISL_420119, EPI_ISL_420120, EPI_ISL_420121, EPI_ISL_420122, EPI_ISL_420123, EPI_ISL_420124, EPI_ISL_425178, EPI_ISL_425179, EPI_ISL_425180, EPI_ISL_425181, EPI_ISL_425182, EPI_ISL_425183, EPI_ISL_425184, EPI_ISL_425185, EPI_ISL_425186, EPI_ISL_425191, EPI_ISL_425192, EPI_ISL_425194, EPI_ISL_425195, EPI_ISL_425196, EPI_ISL_425197, EPI_ISL_425198, EPI_ISL_425213, EPI_ISL_425216, EPI_ISL_425217, EPI_ISL_425218, EPI_ISL_425219, EPI_ISL_436196, EPI_ISL_436199, EPI_ISL_436202, EPI_ISL_436208, EPI_ISL_436211, EPI_ISL_436213, EPI_ISL_436214, EPI_ISL_436220, EPI_ISL_436221, EPI_ISL_436222, EPI_ISL_436223, EPI_ISL_436225, EPI_ISL_436226, EPI_ISL_436234, EPI_ISL_436361, EPI_ISL_436382, EPI_ISL_436383, EPI_ISL_436385, EPI_ISL_436395 | | | |
| see above | Servicio de Microbiología. Consorcio Hospital General Universitario de Valencia | Sequencing and Bioinformatics Service and Molecular Epidemiology Research Group. FISABIO-Public Health | ; Beatriz Beamud; Concepcion Gimeno; David Navarro; Fernando Gonzalez-Candelas; Giuseppe D'Auria; Griselda De Marco; Inma Galán Vendrell; Ivan Ansari; Lidia Ruiz Roldan; Llúcia Martínez-Priego; Llúcia Martínez-PriegoGriselda De Marco; Loreto Ferrús Abad; Maria Alma Bracho; Maria Dolores Ocete; Mariana Reyes-Prieto; Marta Pla Diaz; Neris Garcia-Gonzalez; Paula Ruiz-Hueso; Sandra Carbo; Vicente Soriano Chirona |
| EPI_ISL_452368, EPI_ISL_452372, EPI_ISL_452373, EPI_ISL_452374, EPI_ISL_452375, EPI_ISL_452376, EPI_ISL_452377, EPI_ISL_452378, EPI_ISL_452379, EPI_ISL_452380, EPI_ISL_452381, EPI_ISL_452382, EPI_ISL_452384, EPI_ISL_452386, EPI_ISL_452387, EPI_ISL_452388, EPI_ISL_452389, EPI_ISL_452391, EPI_ISL_452392, EPI_ISL_452393, EPI_ISL_452394, EPI_ISL_452395, EPI_ISL_452397, EPI_ISL_452398, EPI_ISL_452399, EPI_ISL_452400, EPI_ISL_452401, EPI_ISL_452402, EPI_ISL_452403, EPI_ISL_452404, EPI_ISL_452405, EPI_ISL_452406, EPI_ISL_452407, EPI_ISL_452408, EPI_ISL_452409, EPI_ISL_452411, EPI_ISL_452412, EPI_ISL_452413, EPI_ISL_452414, EPI_ISL_452415, EPI_ISL_452416, EPI_ISL_452417, EPI_ISL_452418, EPI_ISL_452419, EPI_ISL_452420, EPI_ISL_452421, EPI_ISL_452422, EPI_ISL_452423, EPI_ISL_452424, EPI_ISL_452425, EPI_ISL_452426, EPI_ISL_452427, EPI_ISL_452428, EPI_ISL_452429, EPI_ISL_452430, EPI_ISL_452432, EPI_ISL_452433, EPI_ISL_452434, EPI_ISL_452435, EPI_ISL_452436, EPI_ISL_452437, EPI_ISL_452438, EPI_ISL_452440, EPI_ISL_452441, EPI_ISL_452442, EPI_ISL_452443, EPI_ISL_452444, EPI_ISL_452445, EPI_ISL_452446, EPI_ISL_452448, EPI_ISL_452450, EPI_ISL_452451, EPI_ISL_452452, EPI_ISL_509602, EPI_ISL_509603, EPI_ISL_509604, EPI_ISL_509605, EPI_ISL_509606, EPI_ISL_509607, EPI_ISL_509608, EPI_ISL_509609, EPI_ISL_509611, EPI_ISL_509620, EPI_ISL_509622, EPI_ISL_509623, EPI_ISL_509625, EPI_ISL_509627, EPI_ISL_509629, EPI_ISL_509631, EPI_ISL_509632, EPI_ISL_565908, EPI_ISL_565909, EPI_ISL_565910, EPI_ISL_565911, EPI_ISL_565914, EPI_ISL_565915, EPI_ISL_565916, EPI_ISL_565917, EPI_ISL_565918, EPI_ISL_565919 | | | |
| see above | Servicio de Microbiología. HRU de Málaga. Servicio Andaluz de Salud | SeqCOVID-SPAIN consortium/IBV(CSIC) | Begoña Palop Borrás and SeqCOVID-SPAIN consortium; Inmaculada de Toro Peinado; Inmaculada de Toro Peinado. MªConcepción Mediavilla Gradolph. Begoña Palop Borrás and SeqCOVID-SPAIN consortium; Ma Concepción Mediavilla Gradolph; Maria Concepción Mediavilla Gradolph |
| EPI_ISL_537968, EPI_ISL_537978, EPI_ISL_537984, EPI_ISL_537985, EPI_ISL_537989, EPI_ISL_538624, EPI_ISL_538630, EPI_ISL_538638, EPI_ISL_538648, EPI_ISL_583933, EPI_ISL_583941, EPI_ISL_583943, EPI_ISL_583947, EPI_ISL_583948, EPI_ISL_654481, EPI_ISL_657450, EPI_ISL_722239, EPI_ISL_796068, EPI_ISL_796106, EPI_ISL_796151 | | | |
| see above | Servicio de Microbiología. Hospital General Universitario de Castellón | SeqCOVID-SPAIN consortium/IBV(CSIC) | María Dolores Tirado Balaguer and SeqCOVID-SPAIN consortium; María Dolores Tirado and SeqCOVID-SPAIN consortium; Rosario Moreno; Rosario Moreno Muñoz |
| EPI_ISL_480956, EPI_ISL_480959, EPI_ISL_480964, EPI_ISL_480966, EPI_ISL_480968, EPI_ISL_480973, EPI_ISL_480974, EPI_ISL_480976, EPI_ISL_480979, EPI_ISL_480980, EPI_ISL_480982, EPI_ISL_480985, EPI_ISL_480986, EPI_ISL_480987, EPI_ISL_480990, EPI_ISL_480991, EPI_ISL_480992, EPI_ISL_480993, EPI_ISL_480997, EPI_ISL_481001, EPI_ISL_481005, EPI_ISL_481009, EPI_ISL_481010, EPI_ISL_481012, EPI_ISL_481016, EPI_ISL_481019, EPI_ISL_481020, EPI_ISL_481024, EPI_ISL_481026, EPI_ISL_481027, EPI_ISL_481028, EPI_ISL_481030, EPI_ISL_481036, EPI_ISL_481039, EPI_ISL_481040, EPI_ISL_538066, EPI_ISL_538069, EPI_ISL_538070 | | | |

| | | | |
|---|---|---|---|
| see above | Servicio de Microbiología. Hospital Universitario Donostia. OSI Donostialdea. Área de Enfermedades Infecciosas, Grupo de Infección Respiratoria y Resistencia Antimicrobiana. Instituto de Investigación Sanitaria Biodonostia | SeqCOVID-SPAIN consortium/IBV(CSIC) | Gustavo Cilla; Jose Maria Marimón and SeqCOVID-SPAIN consortium; Luis Piñeiro; Milagrosa Montes |
| EPI_ISL_452618, EPI_ISL_452624, EPI_ISL_452625, EPI_ISL_452628, EPI_ISL_452632, EPI_ISL_452637, EPI_ISL_452638, EPI_ISL_452648, EPI_ISL_452649, EPI_ISL_452660, EPI_ISL_452662, EPI_ISL_452667, EPI_ISL_452673, EPI_ISL_452675, EPI_ISL_452676, EPI_ISL_452677, EPI_ISL_452678, EPI_ISL_452685, EPI_ISL_452690 | | | |
| see above | Servicio de Microbiología. Hospital Universitario Donostia. OSI Donostialdea. Área de Enfermedades Infecciosas, Grupo de Infección Respiratoria y Resistencia Antimicrobiana. Instituto de Investigación Sanitaria Biodonostia. | SeqCOVID-SPAIN consortium/IBV(CSIC) | Gustavo Cilla; Jose Maria Marimón and SeqCOVID-SPAIN consortium; Luis Piñeiro; Milagrosa Montes |
| EPI_ISL_414934, EPI_ISL_414935, EPI_ISL_414936, EPI_ISL_414937, EPI_ISL_414938, EPI_ISL_414939, EPI_ISL_414940, EPI_ISL_414941 | | | |
| see above | Shandong Provincial Center for Disease Control and Prevention | Beijing Institute of Microbiology and Epidemiology | Bo Pang; Can-Fang Lin; Cun-Bao Li; Dian-Ming Kang; Feng Gao; Guo-Lin Wang; Jie Lei; Li-Jun Duan; Lin Yao; Mai-Juan Ma; Ming-Xiao Yao; Sheng-Xiang Ji; Wen-Kui Sun; Xiang-Na Zhao; Xiao Wei; Xiao-Li Zhang; Xiao-Lin Jiang; Yang Hang; Zeng-Qiang Kou |
| EPI_ISL_416316, EPI_ISL_416317, EPI_ISL_416318, EPI_ISL_416319, EPI_ISL_416322, EPI_ISL_416323, EPI_ISL_416326, EPI_ISL_416328, EPI_ISL_416330, EPI_ISL_416333, EPI_ISL_416335, EPI_ISL_416338, EPI_ISL_416339, EPI_ISL_416341, EPI_ISL_416343, EPI_ISL_416344, EPI_ISL_416350, EPI_ISL_416352, EPI_ISL_416357, EPI_ISL_416366, EPI_ISL_416368, EPI_ISL_416372, EPI_ISL_416377, EPI_ISL_416378, EPI_ISL_416381, EPI_ISL_416392, EPI_ISL_416402, EPI_ISL_416403, EPI_ISL_416409 | | | |
| see above | Shanghai Public Health Clinical Center, Shanghai Medical College, Fudan University | National Research Center for Translational Medicine (Shanghai), Ruijin Hospital affiliated to Shanghai Jiao Tong University School of Medicine & Shanghai Public Health Clinical Center | Gang Lu; Hongzhou Lu; Saijuan Chen; Shengyue Wang; Xiaonan Zhang; Yun Ling; Yun Tan |
| EPI_ISL_463889, EPI_ISL_463894, EPI_ISL_463895, EPI_ISL_463896, EPI_ISL_463898, EPI_ISL_463901, EPI_ISL_463902 | | | |
| see above | Shaoxing Center for Disease Control and Prevention | Department of Pathology and Laboratory Medicine, University of California Los Angeles | Evann E. Hilt; Fan Li; Huan Wu; JiLing Wang; Jialiang Tang; Jinkun Chen; QinChao Zhang; Shangxin Yang; Yifang Wang; Zhuojing Jiang; Ziqin Li |
| EPI_ISL_582125, EPI_ISL_582126 | Sheikh Khalifa Medical City | Molecular Surveillance lab Sheikh Khalifa Medical City | Amirtharaj Francis; Hala Imambaccus; Hiba Saud; Sahar Almarzooqi; Sajeed Abdul; Stefan Weber |
| EPI_ISL_582608 | Sheikh Khalifa Medical City | Molecular/Surveillance lab Sheikh Khalifa Medical City | Amirtharaj Francis; Hala Imambaccus; Hiba Saud; Sahar Almarzooqi; Sajeed Abdul; Stefan Weber |
| EPI_ISL_406593 | Shenzhen Key Laboratory of Pathogen and Immunity, National Clinical Research Center for Infectious Disease, Shenzhen Third People's Hospital | Shenzhen Key Laboratory of Pathogen and Immunity, National Clinical Research Center for Infectious Disease, Shenzhen Third People's Hospital | Chenguang Shen; Haixia Zheng; Li Xing; Yang Yang; Yingxia Liu; Zhixiang Xu |
| EPI_ISL_462086 | Singapore General Hospital | Department of Microbiology | Chenhao Li; Karrie Ko; Kern Rei Chng; Kian Sing Chan; Kun Lee Lim; Lynette Oon; Niranjan Nagarajan; Nurdyana Abdul Rahman |
| EPI_ISL_1113236, EPI_ISL_1113262 | Sonora Quest Laboratories | TGen North | "Jolene Bowers; Ashlyn Pfeiffer; Chris French; Darrin Lemmer; Dave Engelthaler; Hayley Yaglom; Megan Folkerts; The Arizona COVID Genomics Union (ACGU)" |
| EPI_ISL_696419, EPI_ISL_696446 | Sonora Quest Laboratories, Laboratory Sciences of Arizona | TGen North | Ashlyn Pfeiffer; Chris French; Darrin Lemmer; Dave Engelthaler; Hayley Yaglom; Jolene Bowers; Megan Folkerts; The Arizona COVID Genomics Union (ACGU) |
| EPI_ISL_529171, EPI_ISL_529172, EPI_ISL_529173, EPI_ISL_529174, EPI_ISL_529175 | South Carolina Department of Health and Environmental Control | South Carolina Department of Health and Environmental Control | Haley V. Flores |
| EPI_ISL_451634, EPI_ISL_455078, EPI_ISL_455079, EPI_ISL_455080, EPI_ISL_455082, EPI_ISL_455088, EPI_ISL_455089, EPI_ISL_455096, EPI_ISL_455098 | | | |
| see above | South Eastern Area Laboratory Services | NSW Health Pathology - Institute of Clinical Pathology and Medical Research; Westmead Hospital; University of Sydney | CIDM-PH et al. |
| EPI_ISL_478683, EPI_ISL_478684, EPI_ISL_478685, EPI_ISL_478686, EPI_ISL_478687, EPI_ISL_478688, EPI_ISL_478689, EPI_ISL_478690, EPI_ISL_478691, EPI_ISL_478692, EPI_ISL_478708, EPI_ISL_490032, EPI_ISL_490034, EPI_ISL_490035, EPI_ISL_498751 | | | |
| see above | South Eastern Area Laboratory Services (SEALS) | NSW Health Pathology - Institute of Clinical Pathology and Medical Research; Westmead Hospital; University of Sydney | CIDM-PH et al. |
| EPI_ISL_456276, EPI_ISL_456277, EPI_ISL_456286 | Southern Community Labs Dunedin | Institute of Environmental Science and Research (ESR) | Anja Werno; Antje van der Linden; Arlo Upton; Chris Mansell; David Hammer; Dragana Drinkovic; Erasmus Smit; Gary McAuliffe; Hana Sofia Andersson; James Ussher; Jill Sherwood; Joep de Ligt; Josh Freeman; Julia Howard; Juliet Elvy; Lauren Jelly; Mary DeAlmeida; Matt Blakiston; Matt Storey; Matthew Rogers; Max Bloomfield; Michael Addidle; Michelle Balm; Sally Roberts; Sarah Jefferies; Sharmini Muttaiyah; Susan Morpeth; Susan Taylor; Timothy Blackmore; Vani Sathyendran; Veronica Playle; Virginia Hope; Xiaoyun Ren |
| EPI_ISL_469243 | Special Infectious Agents Unit | Special Infectious Agents Unit | A.M.; Al-Sobahy; Azhar; E.I.; El-Kafrawy; Farraj; Hassan; N.A.; S.A.; T.L.; Tolah; Uthman |
| EPI_ISL_2976228, EPI_ISL_2976237 | St. Luke's Health Systems Reference Lab | Idaho Bureau of Laboratories | "R. Beukelman; Aimee Ceniseros; Christian Loera; Christopher Ball"; Matthew Charles Burns; Robert L. Voermans |
| EPI_ISL_514646 | St. Luke's Hospital | Minnesota Department of Health, Public Health Laboratory | Jacob Garfin; Matt Plumb; and Xiong Wang |
| EPI_ISL_450457, EPI_ISL_450473, EPI_ISL_450477, EPI_ISL_476768, EPI_ISL_476776, EPI_ISL_476779, EPI_ISL_476781, EPI_ISL_476783 | | | |
| see above | Stanford clinical virology lab | Chan-Zuckerberg Biohub | Benjamin Pinksy; Carlos Bustamante; Euan Ashley; Hannah N. DeJong; Jason Andrews; John Gorzynski; Katharine Walter; Manuel Rivas; Matthew T. Wheeler; Victoria N. Parikh; with CZB Cliahub Consortium |
| EPI_ISL_416473 | State Key Laboratory for Diagnosis and Treatment of Infectious Diseases, National Clinical Research Center for Infectious Diseases, First Affiliated Hospital, Zhejiang University School of Medicine, Hangzhou, China 310003 | State Key Laboratory for Diagnosis and Treatment of Infectious Diseases, National Clinical Research Center for Infectious Diseases, First Affiliated Hospital, Zhejiang University School of Medicine, Hangzhou, China 310003 | Changzhong Jin; Chao Jiang; Fumin Liu; Haibo Wu; Hangping Yao; Lanjuan Li; Linfang Cheng; Min Zheng; Nanping Wu; Xiangyun Lu; Zhigang Wu |
| EPI_ISL_416042 | State Key Laboratory for Diagnosis and Treatment of Infectious Diseases, National Clinical Research Center for Infectious Diseases, First Affiliated Hospital, Zhejiang University School of Medicine, Hangzhou, China. 310003 | State Key Laboratory for Diagnosis and Treatment of Infectious Diseases, National Clinical Research Center for Infectious Diseases, First Affiliated Hospital, Zhejiang University School of Medicine, Hangzhou, China. 310003 | Changzhong Jin; Chao Jiang; Fumin Liu; Haibo Wu; Hangping Yao; Lanjuan Li; Linfang Cheng; Min Zheng; Nanping Wu; Xiangyun Lu; Zhigang Wu |
| EPI_ISL_417443 | State Key Laboratory for Emerging Infectious Diseases Department of Microbiology Li Ka Shing Faculty of Medicine The University of Hong Kong | State Key Laboratory for Emerging Infectious Diseases Department of Microbiology Li Ka Shing Faculty of Medicine The University of Hong Kong | Bobo Wing-Yee Mok; Honglin Chen; Kwok-Yung Yuen; Pui Wang; Shaofeng Deng; Siu-Ying Lau; Wenjun Song |
| EPI_ISL_1655937 | State Key Laboratory of Emerging Infectious Diseases, The University of Hong Kong | State Key Laboratory of Emerging Infectious Diseases, The University of Hong Kong | Huachen Zhu; Jia Wang; Lifeng Li; Yi Guan; Yiu Man Cheung |
| EPI_ISL_414663, EPI_ISL_414688, EPI_ISL_414689, EPI_ISL_414690, EPI_ISL_414691 | State Key Laboratory of Respiratory Disease, National Clinical Research Center for Respiratory Disease, Guangzhou Institute of Respiratory Health, the First Affiliated Hospital of Guangzhou Medical University | The First Affiliated Hospital of Guangzhou Medical University & BGI-Shenzhen | Zhao et al |
| EPI_ISL_752609, EPI_ISL_752618 | State Laboratories Division, Hawaii State Department of Health | State Laboratories Division, Hawaii State Department of Health | Drew Kuwazaki; Edward Desmond; Pamela O'Brien; Razvan Sultana; Sabrina Diemert |
| EPI_ISL_428893, EPI_ISL_428898 | State Research Center of Virology and Biotechnology VECTOR, Department of Collection of Microorganisms | State Research Center of Virology and Biotechnology VECTOR, Department of Collection of Microorganisms | Alexander N. Shvalov; Elena V. Gavrilova; Oleg V. Pyankov; Rinat A. Maksyutov; Sergey A. Bodnev; Tatyana V. Tregubchak |
| EPI_ISL_478676, EPI_ISL_478677, EPI_ISL_478678, EPI_ISL_478680, EPI_ISL_478682, EPI_ISL_593756 | Sydney South West Pathology Service (SSWPS) - Liverpool Hospital - NSW Health Pathology | NSW Health Pathology - Institute of Clinical Pathology and Medical Research; Westmead Hospital; University of Sydney | CIDM-PH et al. |
| EPI_ISL_426504, EPI_ISL_426534, EPI_ISL_694047, EPI_ISL_694052 | TGen North | TGen North | Ashlyn Pfeiffer; Chris French; Darrin Lemmer; Dave Engelthaler; Hayley Yaglom; Jolene Bowers; Megan Folkerts; The Arizona COVID Genomics Union (ACGU) |
| EPI_ISL_436101 | TSGH-CP molecular lab | TSGH-CP molecular lab | Cherng-Lih Perng; Chien-Wen Chen; Chih-Kai Chang; Feng-Yee Chang; Hsing-Yi Chung; Hung-Sheng Shang; Jung-Chung Lin; Kuo-Ming Yeh; Kuo-Sheng Hung; Ming-Jr JIAN; Sheng-Kang Chiu; Shih-Hung Tsai; Tien-Yao Chang |
| EPI_ISL_411926 | Taiwan Centers for Disease Control | Taiwan Centers for Disease Control | Ji-Rong Yang; Jung-Jung Mu; Ming-Tsan-Liu; Yu-Chi-Lin |
| EPI_ISL_1379445, EPI_ISL_1379447, EPI_ISL_1379452, EPI_ISL_1379464, EPI_ISL_1379468, EPI_ISL_1379469, EPI_ISL_1379471, EPI_ISL_1379472, EPI_ISL_1379487, EPI_ISL_1379489, EPI_ISL_1379490, EPI_ISL_1379500, EPI_ISL_1492444, EPI_ISL_1492445 | | | |
| see above | Tampa General Hospital Esoteric Lab | Tampa General Hospital Esoteric Research & Development Lab | Amorce Lima; Deanna Becker; Dominic Uy; Grant Vestal; Suzane Silbert; Vicki Healer |
| EPI_ISL_529150 | Technology Centre, Guangzhou Customs | Technology Centre, Guangzhou Customs | Dai, J.; Huang, J.; Huang, S.; Li, X.; Shi, Y.; Sun, F.; Sun, J.; Wang, Y.; Zhang, Z.; Zheng, K.; Zhu, .; Zhuang, Z. |
| EPI_ISL_514438, EPI_ISL_514439 | Tewhey Lab, The Jackson Laboratory | The Jackson Laboratory | Barter, M.; Dewey, H.; Lynch, R.; Matluk, N.; Munger, H.; Tewhey, R. |
| EPI_ISL_452115 | Texas DSHS Lab Services | Pathogen Discovery, Respiratory Viruses Branch, Division of Viral | Alison S. Laufer Halpin; Anna Montmayeur; Anna Uehara; Christopher A. Elkins; Clinton R. Paden; Haibin Wang; Jing Zhang; Krista Queen; Mary S. Keckler; Rachel Marine; Suxiang Tong; Yan Li; Ying Tao; Zachary Weiner |

| | | | |
|---|---|---|---|
| | | Diseases, Centers for Disease Control and Prevention | |
| EPI_ISL_411956 | Texas Department of State Health Services | Pathogen Discovery, Respiratory Viruses Branch, Division of Viral Diseases, Centers for Dieases Control and Prevention | Anna Uehara; Brett L. Whitaker; Brian Lynch; Clinton R. Paden; Haibin Wang; Janna' R. Murray; Jing Zhang; Krista Queen; Lijuan Wang; Senthil Kumar K. Sakthivel; Shifaq Kamili; Stephen Lindstrom; Susan I. Gerber; Suxiang Tong; Xiaoyan Lu; Yan Li; Ying Tao |
| EPI_ISL_524053, EPI_ISL_524054, EPI_ISL_524055, EPI_ISL_524056, EPI_ISL_524057, EPI_ISL_524061, EPI_ISL_524062, EPI_ISL_525763, EPI_ISL_525765, EPI_ISL_631652, EPI_ISL_631668, EPI_ISL_631670, EPI_ISL_631673, EPI_ISL_631675, EPI_ISL_631681, EPI_ISL_644171, EPI_ISL_644177 | | | |
| see above | Texas Department of State Health Services | Texas Department of State Health Services | Anita Pokharel; Bonnie Oh; Cara Akrout; Chun Wang; Chung Wang; Grace Kubin; Jenny Zhang; Maliha Rahman; Mayela Pedrueza; Myong Koag; Rachel Lee; Rashmi Tuladhar |
| EPI_ISL_594455 | Texas Department of State Health Services Public Health Laboratory | Pathogen Discovery, Respiratory Viruses Branch, Division of Viral Diseases, Centers for Disease Control and Prevention | Anna Uehara; Clinton Paden; Haibin Wang; Jing Zhang; Julu Bhatnagar; Krista Queen; Suxiang Tong; Yan Li; Ying Tao |
| EPI_ISL_918160, EPI_ISL_918162, EPI_ISL_937542, EPI_ISL_976893, EPI_ISL_976895, EPI_ISL_976899, EPI_ISL_1296447, EPI_ISL_1296453, EPI_ISL_1296457, EPI_ISL_1296459, EPI_ISL_1296461 | | | |
| see above | Thai Red Cross Emerging Infectious Diseases Health Science Centre, Chulalongkorn Hospital, Faculty of Medicine, Chulalongkorn University | Thai Red Cross Emerging Infectious Diseases Center and Faculty of Medicine, Chulalongkorn University | Apaporn Rodpan; Gompol Suwanpimolkul; Leilani Paitoonpong; Opass Putcharoen; Pattama Torvorapanit; Sininat Petcharat; Sopon Iamsirithaworn; Supaporn Wacharapluesadee; Thiravat Hemachudha; Watsamon Jantarabenjakul; Weenassarin Ampoot; Yutthana Joyjinda |
| EPI_ISL_434707 | Thammasat University Hospital | National Institute of Health. Department of medical Sciences, Ministry of Public Health, Thailand | Chittaganpitch; Malinee; Okada; Parnmen; Phuygun; Pilailuk; Siripaporn; Sittiporn; Sunthareeya; Thanadachakul; Thanutsapa; Waicharoen; Warawan; Wongboot |
| EPI_ISL_672161 | The Ashley Laboratory, Stanford University | Chan-Zuckerberg Biohub | CZB Cliahub Consortium |
| EPI_ISL_412978 | The Central Hospital Of Wuhan | Hubei Provincial Center for Disease Control and Prevention | Bin Fang; Bo Yang; Bo Yu; Faxian Zhan; Guojun Ye; Jing Li; Junqiang Xu; Kun Cai; Linlin Liu; Xiang Li; Xiao Yu; Xixiang Huo; Yongzhong Jiang. |
| EPI_ISL_450442 | The Department of Infectious Disease Prevention and Control, Henan Provincial Center for Disease Control and Prevention | The Department of Infectious Disease Prevention and Control, Henan Provincial Center for Disease Control and Prevention | Hu, X.; Huang; Li, D.; Li, X.; Lu, S.; Wu, B.; X. and Guo, W. |
| EPI_ISL_429075, EPI_ISL_429082, EPI_ISL_429083, EPI_ISL_429084, EPI_ISL_429090, EPI_ISL_429091, EPI_ISL_429092, EPI_ISL_429093, EPI_ISL_429094, EPI_ISL_429095, EPI_ISL_429099 | | | |
| see above | The First Affiliated Hospital of Guangzhou Medical University | BGI-shenzhen & The First Affiliated Hospital of Guangzhou Medical University | ; Daxi Wang; Lu Zhang; Wanying Sun; Yanqun Wang; Zhaoyong Zhang et al. |
| EPI_ISL_457687, EPI_ISL_457688, EPI_ISL_457689, EPI_ISL_457690, EPI_ISL_457692, EPI_ISL_457693, EPI_ISL_457695, EPI_ISL_457696 | | | |
| see above | The First Affiliated Hospital of Guangzhou Medical University, Guangzhou, China | BGI-shenzhen & The First Affiliated Hospital of Guangzhou Medical University | Daxi Wang; Lu Zhang; Wanying Sun; Yanqun Wang; Zhaoyong Zhang et al. |
| EPI_ISL_489708 | The National Institute of Public Health | The National Institute of Public Health and State Veterinary Institute Prague | A; D; H; J; Jirincova; L; Nagy; Novakova; Trnka; Vecerova |
| EPI_ISL_417577, EPI_ISL_417587, EPI_ISL_417615, EPI_ISL_417717, EPI_ISL_417720, EPI_ISL_417724, EPI_ISL_417762, EPI_ISL_417789, EPI_ISL_417791, EPI_ISL_417818, EPI_ISL_417871, EPI_ISL_424429, EPI_ISL_424436, EPI_ISL_424462, EPI_ISL_424469, EPI_ISL_1586632 | | | |
| see above | The National University Hospital of Iceland | deCODE genetics | Agnar Helgason; Alma Moller; Arna B Agustsdottir; Arnaldur Gylfason; Asgeir Sigurdsson; Aslaug Jonasdottir; Berglind Eiriksdottir; Bjarni Thorbjornsson; Brynjar O Jensson; Daniel F Gudbjartsson; Droplaug N Magnusdottir; Elisabet E Gardarsdottir; Emil A Thorarensen; Gardar Sveinbjornsson; Gisli Masson; Gudmundur Georgsson; Gudmundur L Norddahl; Gudrun Sigmundsdottir; Hakon Jonsson; Hannes Eggertsson; Hilma Holm; Ingileif Jonsdottir; Jona Saemundsdottir; Kamilla S Josefsdottir; Kari Stefansson; Karl G Kristinsson; Kjartan R Gudmundsson; Kristin E Sveinsdottir; Kristin E Sveinsdottir; Louise le Roux; Maney Sveinsdottir; Olafia S Gretarsdottir; Olafia S Gretarsdottir; Olafur T Magnusson; Pall Melsted; Patrick Sulem; Run Fridriksdottir; Solvi Rognvaldsson; Thora R Gunnarsdottir; Thordur Kristjansson; Thorolfur Gudnason; Unnur Thorsteinsdottir |
| EPI_ISL_427288 | The Ohio State University | The Ohio State University-James Molecular Lab at Polaris | Dan Jones; Erica Vincent; Huolin Tu; Jason Garee; Joan-Miquel Balada-Llasat; Matthew Hunt; Preeti Panchioli; Weiqiang Zhao |
| EPI_ISL_405839, EPI_ISL_406030 | The University of Hong Kong - Shenzhen Hospital | Li Ka Shing Faculty of Medicine, The University of Hong Kong | C.C.-Y.; C.K.-M. and Yuen; Cai; Chan; Chen, H.; Cheng; Chu, H.; H.W.; Hui; Ip; J.D.; J.F.-W.; J.P.; K.H.; K.K.-W.; K.Y.; Kok; Liu, J.; Lo; Poon; R.W.-S.; S.K.-F.; To; Tsai; V.C.-C.; V.K.-M.; W.M.; Xing, F.; Yang, J.; Yip; Yuan, S. |
| EPI_ISL_412983 | Tianmen Center for Disease Control and Prevention | Hubei Provincial Center for Disease Control and Prevention | Bin Fang; Bo Yang; Bo Yu; Faxian Zhan; Guojun Ye; Jing Li; Junqiang Xu; Kun Cai; Linlin Liu; Xiang Li; Xiao Yu; Xierong Li; Xixiang Huo; Yangyang Tao; YiFa Zhu; Yongzhong Jiang. |
| EPI_ISL_480034 | Tochigi Prefectural Institute of Public Health and Environmental Science | Pathogen Genomics Center, National Institute of Infectious Diseases | Ako Nakajima; Hajime Kamiya; Kentaro Itokawa; Makoto Kuroda; Masanori Hashino; Motoi Suzuki; Rina Tanaka; Tsuyoshi Sekizuka |
| EPI_ISL_479888, EPI_ISL_479890, EPI_ISL_479892, EPI_ISL_479959, EPI_ISL_479960, EPI_ISL_480227, EPI_ISL_684137, EPI_ISL_684154, EPI_ISL_684512, EPI_ISL_685606 | | | |
| see above | Tokyo Metropolitan Institute of Public Health | Pathogen Genomics Center, National Institute of Infectious Diseases | Hajime Kamiya; Kenji Sadamasu; Kentaro Itokawa; Makoto Kuroda; Mami Nagashima; Masanori Hashino; Motoi Suzuki; Rina Tanaka; Takashi Chiba; Tsuyoshi Sekizuka |
| EPI_ISL_463993, EPI_ISL_586278, EPI_ISL_586356, EPI_ISL_586390, EPI_ISL_755815, EPI_ISL_755854, EPI_ISL_770828, EPI_ISL_933603, EPI_ISL_933606 | | | |
| see above | Toronto Invasive Bacterial Diseases Network | McMaster University | Ahmed Draia; Allison McGeer; Andrew G. McArthur; Angel Li; Emily Panousis; Hooman Derakhshani; Jalees Nasir; Kuganya Nirmalarajah; Michael Surette; Patryk Aftanas; Samira Mubareka |
| EPI_ISL_445096, EPI_ISL_445099, EPI_ISL_445100, EPI_ISL_445102, EPI_ISL_445103, EPI_ISL_483309, EPI_ISL_483391 | | | |
| see above | UC San Diego Center for Advanced Laboratory Medicine | Andersen lab at Scripps Research | Ji H Shin; SEARCH Alliance San Diego with David Pride |
| EPI_ISL_537493, EPI_ISL_537495, EPI_ISL_537539, EPI_ISL_537572, EPI_ISL_537576, EPI_ISL_537580, EPI_ISL_537582, EPI_ISL_537587, EPI_ISL_537588, EPI_ISL_537603 | | | |
| see above | UCLA Pathology Clinical Microbiology Lab | Kruglyak Lab | Guo et al. |
| EPI_ISL_428997, EPI_ISL_429000, EPI_ISL_429005, EPI_ISL_429008, EPI_ISL_429010, EPI_ISL_429019, EPI_ISL_429020, EPI_ISL_429021, EPI_ISL_429025, EPI_ISL_429032, EPI_ISL_429039, EPI_ISL_429040, EPI_ISL_429041, EPI_ISL_429045, EPI_ISL_429048, EPI_ISL_429049, EPI_ISL_429055, EPI_ISL_429056, EPI_ISL_429057, EPI_ISL_429058, EPI_ISL_429059, EPI_ISL_429060, EPI_ISL_429063, EPI_ISL_429064, EPI_ISL_429065, EPI_ISL_429066, EPI_ISL_429068, EPI_ISL_429069, EPI_ISL_429070, EPI_ISL_429071, EPI_ISL_429072, EPI_ISL_429073, EPI_ISL_445170, EPI_ISL_445172, EPI_ISL_445173, EPI_ISL_445174 | | | |
| see above | UCSF Clinical Microbiology Laboratory | Chan-Zuckerberg Biohub | CZB Cliahub Consortium |
| EPI_ISL_450234 | UCSF Clinical Microbiology Laboratory | Chiu Laboratory, University of California, San Francisco | Scot Federman; Wei Gu; Xianding Deng; and Charles Y. Chiu |
| EPI_ISL_511007, EPI_ISL_511020, EPI_ISL_511118 | ULS Guarda | Instituto Nacional de Saude (INSA) | Borges et al |
| EPI_ISL_511177 | UNILABS | Instituto Nacional de Saude (INSA) | Borges et al |
| EPI_ISL_445322 | UNIV.DE CHILE HOSP.CLINICO | Instituto de Salud Publica de Chile | Alejandra Acevedo; Andrés E Castillo; Bárbara Parra; Carolina Tambley; Gabriel Leal; Jaime Lagos; Jorge Fernandez; Loredana Arata; Patricia Bustos; Paz Tapia; Rodrigo Fasce; Winston Andrade |
| EPI_ISL_445247 | UNIVERSIDAD DE LOS ANDES | Instituto de Salud Publica de Chile | Alejandra Acevedo; Andrés E Castillo; Bárbara Parra; Carolina Tambley; Gabriel Leal; Jaime Lagos; Jorge Fernandez; Loredana Arata; Patricia Bustos; Paz Tapia; Rodrigo Fasce; Winston Andrade |
| EPI_ISL_436487, EPI_ISL_436503 | UPMC Clinical Laboratory | Microbial Genome Sequencing Center, Microbial Genomic Epidemiological Laboratory | Chinelo Ezeonwuku; Dan Snyder; Jane W. Marsh; Kady D Waggle; Lee H. Harrison; Marissa P Griffith; Mustapha M Mustapha; Stephanie L Mitchell; Vatsala R Srinivasa; Vaughn S. Cooper |
| EPI_ISL_454361, EPI_ISL_454366, EPI_ISL_454384 | UPMC Clinical Microbiology Laboratory | Microbial Genome Sequencing Center, Microbial Genomic Epidemiological Laboratory | Chinelo Ezeonwuku; Dan Snyder; Jane W. Marsh; Kady D. Waggle; Lee H. Harrison; Marissa P. Griffith; Mustapha M. Mustapha; Stephanie L. Mitchell; Vatsala R. Srinivasa; Vaughn S. Cooper |
| EPI_ISL_424889 | UT-Unified State Labs: Public Health Utah DOH | Pathogen Discovery, Respiratory Viruses Branch, Division of Viral Diseases, Centers for Disease Control and Prevention | Alison S. Laufer Halpin; Anna Uehara; Christopher A. Elkins; Clinton R. Paden; Haibin Wang; Jing Zhang; Krista Queen; Mary S. Keckler; Rachel Marine; Suxiang Tong; Yan Li; Ying Tao |
| EPI_ISL_413562, EPI_ISL_413563, EPI_ISL_413601, EPI_ISL_413649, EPI_ISL_413650, EPI_ISL_413651, EPI_ISL_413652, EPI_ISL_413653, EPI_ISL_414363, EPI_ISL_414364, EPI_ISL_414365, EPI_ISL_414366, EPI_ISL_414367, EPI_ISL_414368, EPI_ISL_414369, EPI_ISL_414592, EPI_ISL_414593, EPI_ISL_414595, EPI_ISL_414596, EPI_ISL_414597, EPI_ISL_414617, EPI_ISL_414618, EPI_ISL_414619, EPI_ISL_414620, EPI_ISL_414621, EPI_ISL_414622, EPI_ISL_415591, EPI_ISL_415592, EPI_ISL_415593, EPI_ISL_415594, EPI_ISL_415595, EPI_ISL_415596, EPI_ISL_415598, EPI_ISL_415599, EPI_ISL_415600, EPI_ISL_415602, EPI_ISL_415603, EPI_ISL_415604, EPI_ISL_415605, EPI_ISL_415606, EPI_ISL_415607, EPI_ISL_415608, EPI_ISL_415609, EPI_ISL_415610, EPI_ISL_415611, EPI_ISL_415612, EPI_ISL_415613, EPI_ISL_415614, EPI_ISL_415615, EPI_ISL_415616, EPI_ISL_415617, EPI_ISL_415618, EPI_ISL_415619, EPI_ISL_415620, EPI_ISL_415621, EPI_ISL_415622, EPI_ISL_415623, EPI_ISL_415624, EPI_ISL_415626, EPI_ISL_415627, EPI_ISL_416433, EPI_ISL_416435, EPI_ISL_416436, EPI_ISL_416437, EPI_ISL_416439, EPI_ISL_416440, EPI_ISL_416441, EPI_ISL_416442, EPI_ISL_416443, EPI_ISL_416444, EPI_ISL_416445, EPI_ISL_416446, EPI_ISL_416447, EPI_ISL_416448, EPI_ISL_416450, EPI_ISL_416451, EPI_ISL_416453, EPI_ISL_416454, EPI_ISL_416455, EPI_ISL_416456, EPI_ISL_416635, EPI_ISL_416636, EPI_ISL_416637, EPI_ISL_416638, EPI_ISL_416639, EPI_ISL_416640, EPI_ISL_416641, EPI_ISL_416643, EPI_ISL_416644, EPI_ISL_416645, EPI_ISL_416649, EPI_ISL_416650, EPI_ISL_416651, EPI_ISL_416652, EPI_ISL_416654, EPI_ISL_416656, EPI_ISL_416657, EPI_ISL_416659, EPI_ISL_416662, EPI_ISL_416663, EPI_ISL_416664, EPI_ISL_416665, EPI_ISL_416666, EPI_ISL_416667, EPI_ISL_416668, EPI_ISL_416669, EPI_ISL_416670, EPI_ISL_416677, EPI_ISL_416679, EPI_ISL_416680, EPI_ISL_416681, EPI_ISL_416682, EPI_ISL_416683, EPI_ISL_416684, EPI_ISL_416685, EPI_ISL_416686, EPI_ISL_416687, EPI_ISL_416688, EPI_ISL_416689, EPI_ISL_416690, EPI_ISL_416692, EPI_ISL_416694, EPI_ISL_416695, EPI_ISL_416696, EPI_ISL_416700, EPI_ISL_416701, EPI_ISL_416702, EPI_ISL_416703, EPI_ISL_416705, EPI_ISL_416709, EPI_ISL_416710, EPI_ISL_416711, EPI_ISL_416712, EPI_ISL_416713, EPI_ISL_416714, EPI_ISL_416715, EPI_ISL_416717, EPI_ISL_416718, EPI_ISL_416720, EPI_ISL_416721, EPI_ISL_416722, EPI_ISL_416726, EPI_ISL_416727, EPI_ISL_416728, EPI_ISL_417344, EPI_ISL_417347, EPI_ISL_417348, EPI_ISL_417349, EPI_ISL_417350, EPI_ISL_417351, EPI_ISL_417353, EPI_ISL_417354, EPI_ISL_417355, EPI_ISL_417356, EPI_ISL_417357, EPI_ISL_417358, EPI_ISL_417360, EPI_ISL_417361, EPI_ISL_417363, EPI_ISL_417364, EPI_ISL_417365, EPI_ISL_417366, EPI_ISL_417367, EPI_ISL_417368, EPI_ISL_417369, EPI_ISL_417370, EPI_ISL_417371, EPI_ISL_417374, EPI_ISL_417375, EPI_ISL_417377, EPI_ISL_417378, EPI_ISL_417380, EPI_ISL_417381, EPI_ISL_417448, EPI_ISL_417449, EPI_ISL_417450, EPI_ISL_417453, EPI_ISL_417456, EPI_ISL_418029, EPI_ISL_418030, EPI_ISL_418031, EPI_ISL_418034, EPI_ISL_418038, EPI_ISL_418040, EPI_ISL_418045, EPI_ISL_418046, EPI_ISL_418047, EPI_ISL_418048, EPI_ISL_418050, EPI_ISL_418052, EPI_ISL_418055, EPI_ISL_418059, EPI_ISL_418061, EPI_ISL_418062, EPI_ISL_418063, EPI_ISL_418064, EPI_ISL_418065, EPI_ISL_418066, EPI_ISL_418069, EPI_ISL_418070, EPI_ISL_418071, EPI_ISL_418074, EPI_ISL_418075, EPI_ISL_418077, EPI_ISL_418078, EPI_ISL_418079, EPI_ISL_418082, EPI_ISL_418866, EPI_ISL_418867, EPI_ISL_418868, EPI_ISL_418869, EPI_ISL_418870, EPI_ISL_418871, EPI_ISL_418872, EPI_ISL_418873, EPI_ISL_418874, EPI_ISL_418877, EPI_ISL_418881, EPI_ISL_418882, EPI_ISL_418883, EPI_ISL_418884, EPI_ISL_418885, EPI_ISL_418886, EPI_ISL_418887, EPI_ISL_418888, EPI_ISL_418890, EPI_ISL_418892, EPI_ISL_418897, EPI_ISL_418899, EPI_ISL_418901, EPI_ISL_418903, EPI_ISL_418905, EPI_ISL_418906, EPI_ISL_418907, EPI_ISL_418908, EPI_ISL_418909, EPI_ISL_418910, EPI_ISL_418911, EPI_ISL_418912, EPI_ISL_418914, EPI_ISL_418917, EPI_ISL_418921, EPI_ISL_418922, EPI_ISL_418927, EPI_ISL_418928, EPI_ISL_418930, EPI_ISL_418931, EPI_ISL_418934, EPI_ISL_418935, EPI_ISL_418936, EPI_ISL_418937, EPI_ISL_418938, EPI_ISL_418939, EPI_ISL_418940, EPI_ISL_418942, EPI_ISL_418943, EPI_ISL_418944, EPI_ISL_418945, EPI_ISL_418946, EPI_ISL_418947, EPI_ISL_418948, EPI_ISL_418949, EPI_ISL_418950, EPI_ISL_418952, EPI_ISL_418955, EPI_ISL_422961, EPI_ISL_422962, EPI_ISL_422963, EPI_ISL_422964, EPI_ISL_422965, EPI_ISL_422971, EPI_ISL_422973, EPI_ISL_422974, EPI_ISL_422975, EPI_ISL_422982, EPI_ISL_422984, EPI_ISL_422986, EPI_ISL_422988, EPI_ISL_422989, EPI_ISL_422990, EPI_ISL_422991, EPI_ISL_422992, EPI_ISL_422993, EPI_ISL_422995, EPI_ISL_422997, EPI_ISL_422999, EPI_ISL_423003, EPI_ISL_423004, EPI_ISL_423005, EPI_ISL_423007, EPI_ISL_423008, EPI_ISL_423009, EPI_ISL_423012, EPI_ISL_423014, EPI_ISL_423016, EPI_ISL_423017, EPI_ISL_423022, EPI_ISL_423025, EPI_ISL_423027, EPI_ISL_423033, EPI_ISL_424168, EPI_ISL_424169, EPI_ISL_424173, EPI_ISL_424175, EPI_ISL_424177, EPI_ISL_424179, EPI_ISL_424180, EPI_ISL_424181, EPI_ISL_424185, EPI_ISL_424191, EPI_ISL_424196, EPI_ISL_424198, EPI_ISL_424199, EPI_ISL_424206, EPI_ISL_424209, EPI_ISL_424210, EPI_ISL_424213, EPI_ISL_424217, EPI_ISL_424218, EPI_ISL_424219, EPI_ISL_424220, EPI_ISL_424222, EPI_ISL_424223, EPI_ISL_424226, EPI_ISL_424228, EPI_ISL_424230, EPI_ISL_424231, EPI_ISL_424232, EPI_ISL_424240, EPI_ISL_424241, EPI_ISL_424243, EPI_ISL_424247, EPI_ISL_424249, EPI_ISL_424250, EPI_ISL_424252, EPI_ISL_424258, EPI_ISL_424261, EPI_ISL_424262, EPI_ISL_424264, EPI_ISL_424265, EPI_ISL_424266, EPI_ISL_424267, EPI_ISL_424268, EPI_ISL_424269, EPI_ISL_424272, EPI_ISL_424275, EPI_ISL_424276, EPI_ISL_424279, EPI_ISL_424282, EPI_ISL_424283, EPI_ISL_424284, EPI_ISL_424285, EPI_ISL_424286, EPI_ISL_424287, EPI_ISL_424288, EPI_ISL_424289, EPI_ISL_424292, EPI_ISL_424295, EPI_ISL_424296, EPI_ISL_424297, EPI_ISL_424298, EPI_ISL_424303, EPI_ISL_424308, EPI_ISL_424323, EPI_ISL_424326, EPI_ISL_424329, EPI_ISL_424336, EPI_ISL_424338, EPI_ISL_426082, EPI_ISL_426096, EPI_ISL_426102, EPI_ISL_426110, EPI_ISL_426111, EPI_ISL_426119, EPI_ISL_426120, EPI_ISL_426129, EPI_ISL_426130, EPI_ISL_426131, EPI_ISL_426133, EPI_ISL_427174, EPI_ISL_427175, EPI_ISL_427177, EPI_ISL_427178, EPI_ISL_427179, EPI_ISL_427182, EPI_ISL_427186, EPI_ISL_427187, EPI_ISL_427190, EPI_ISL_427193, EPI_ISL_427195, EPI_ISL_427203, EPI_ISL_427204, EPI_ISL_427206, EPI_ISL_427208, EPI_ISL_427214, EPI_ISL_427218, EPI_ISL_427220, EPI_ISL_427226, EPI_ISL_427231, EPI_ISL_427237, EPI_ISL_427243, EPI_ISL_427249, EPI_ISL_427252, EPI_ISL_427265, EPI_ISL_427266, EPI_ISL_429598, EPI_ISL_429612, EPI_ISL_429623, EPI_ISL_430867, EPI_ISL_430870, EPI_ISL_430877, EPI_ISL_430878, EPI_ISL_430881, EPI_ISL_430883, EPI_ISL_430886, EPI_ISL_430887, EPI_ISL_430888, EPI_ISL_430889, EPI_ISL_430890, EPI_ISL_430892, EPI_ISL_430894, EPI_ISL_430895, EPI_ISL_430938, EPI_ISL_437804, EPI_ISL_437805, EPI_ISL_437809, EPI_ISL_437810, EPI_ISL_437812, EPI_ISL_437822, EPI_ISL_437824, EPI_ISL_437829, EPI_ISL_460621, EPI_ISL_460623, EPI_ISL_460624, EPI_ISL_460625, EPI_ISL_460627, EPI_ISL_460631, EPI_ISL_460632, EPI_ISL_461422, EPI_ISL_461445, EPI_ISL_476901, EPI_ISL_476902, EPI_ISL_476903, EPI_ISL_476904, EPI_ISL_476905, EPI_ISL_476906, EPI_ISL_476909, EPI_ISL_476910, EPI_ISL_476920, EPI_ISL_476922, EPI_ISL_476923, EPI_ISL_476924, EPI_ISL_476925, EPI_ISL_476926, EPI_ISL_476929, EPI_ISL_476930, EPI_ISL_476932, EPI_ISL_476933, EPI_ISL_476934, EPI_ISL_476935, EPI_ISL_476936, EPI_ISL_476938, EPI_ISL_476939, EPI_ISL_476940, EPI_ISL_477693, EPI_ISL_477694, EPI_ISL_477697, EPI_ISL_477698, EPI_ISL_477699, EPI_ISL_477700, EPI_ISL_477703, EPI_ISL_477704, EPI_ISL_477708, EPI_ISL_477713, EPI_ISL_477717, EPI_ISL_477719, EPI_ISL_477721, EPI_ISL_477723, EPI_ISL_485945, EPI_ISL_485971, EPI_ISL_485974, EPI_ISL_485976, EPI_ISL_485978, EPI_ISL_485981, EPI_ISL_485982, EPI_ISL_485987, EPI_ISL_485988, EPI_ISL_485994, EPI_ISL_486003, EPI_ISL_486004, EPI_ISL_486017, EPI_ISL_486034, EPI_ISL_486040, EPI_ISL_486045, EPI_ISL_486046, EPI_ISL_486047, EPI_ISL_486052, EPI_ISL_486056, EPI_ISL_486059, EPI_ISL_486061, EPI_ISL_486070, EPI_ISL_486074, EPI_ISL_486078, EPI_ISL_486083, EPI_ISL_486084, EPI_ISL_486085, EPI_ISL_486086, EPI_ISL_486088, EPI_ISL_486090, EPI_ISL_486093, EPI_ISL_486095, EPI_ISL_486099, EPI_ISL_486103, EPI_ISL_486105, EPI_ISL_486113, EPI_ISL_490982, EPI_ISL_490996, EPI_ISL_491016, EPI_ISL_491019, EPI_ISL_491022, EPI_ISL_491029, EPI_ISL_491030, EPI_ISL_491031, EPI_ISL_491032, EPI_ISL_497872, EPI_ISL_570028, EPI_ISL_570033, EPI_ISL_570034, EPI_ISL_570040, EPI_ISL_570044, EPI_ISL_570046, EPI_ISL_570047, EPI_ISL_570051, EPI_ISL_570052, EPI_ISL_570058, EPI_ISL_570060, EPI_ISL_570064, EPI_ISL_570069, EPI_ISL_570156, EPI_ISL_570201, EPI_ISL_570204, EPI_ISL_570209, EPI_ISL_570536, EPI_ISL_570537, EPI_ISL_570547, EPI_ISL_570765, EPI_ISL_570798, EPI_ISL_570802, EPI_ISL_570806, EPI_ISL_570895, EPI_ISL_570901, EPI_ISL_570972, EPI_ISL_570973, EPI_ISL_570974, EPI_ISL_570976, EPI_ISL_570977, EPI_ISL_570979, EPI_ISL_570983, EPI_ISL_570986, EPI_ISL_570987, EPI_ISL_576492, EPI_ISL_576494, EPI_ISL_576496, EPI_ISL_576497, EPI_ISL_603257, EPI_ISL_603260, EPI_ISL_603264, EPI_ISL_776695, EPI_ISL_792092, EPI_ISL_792093, EPI_ISL_792095, EPI_ISL_792097, EPI_ISL_792099, EPI_ISL_792100 | | | |
| see above | UW Virology Lab | UW Virology Lab | Alexander Greninger; Amin Addetia; Hong Xie; Keith Jerome; Keith R Jerome; Lasata Shrestha; Meei-Li Huang; Pavitra Roychoudhury; Truong Nguyen; Victoria M Rachleff |
| EPI_ISL_737954 | Uganda Central Public Health Lab and Uganda Virus Research Institute | MRC/UVRI & LSHTM Uganda Research Unit | Dan Lule Bugembe; Matthew Cotten; My V.T. Phan; Pontiano Kaleebu et al. |
| EPI_ISL_451183, | Uganda Virus Research Institute | MRC/UVRI & LSHTM Uganda Research Unit | Beatrice Dhaala; Dan Lule Bugembe; Deogratius Ssemwanga; Henry Kyobe; Henry Mwebesa; Jane Aceng; John Kayiwa; Jonas Lexow; Julius Lutwama; Matthew Cotten; My V.T Phan; Phionah Tushabe; Pontiano Kaleebu; Stephen Balinandi |

| | | | |
|---|---|---|---|
| EPI_ISL_451184, EPI_ISL_451194 | | | |
| EPI_ISL_412979, EPI_ISL_412980 | Union Hospital of Tongji Medical College, Huazhong University of Science and Technology | Hubei Provincial Center for Disease Control and Prevention | Bin Fang; Bo Yang; Bo Yu; Faxian Zhan; Guojun Ye; Jing Li; Junqiang Xu; Kun Cai; Linlin Liu; Xiang Li; Xiao Yu; Xixiang Huo; Yongzhong Jiang. |
| EPI_ISL_812308, EPI_ISL_812320, EPI_ISL_812343 | United States Air Force School of Aerospace Medicine | United States Air Force School of Aerospace Medicine | Amanda Javorina; Anthony Fries; Clarise Starr; Elizabeth Macias; Jennifer Meyer; Sarah Purves; William Gruner |
| EPI_ISL_464002, EPI_ISL_464006 | Unity Health Toronto | Ontario Institute for Cancer Research | Annette Gower; Bernard Lam; Ilinca Lungu; Jared Simpson; Jeremy Johns; Karel Boissinot; Larissa M. Matukas; Mark Downing; Paul Krzyzanowski; Philip Zuzarte; Ramzi Fattouh; Richard de Borja; Samira Mubareka; TIBDN |
| EPI_ISL_660733 | Unité des Virus Émergents | CNR Virus des Infections Respiratoires - France SUD | Antoine Nougairède; Antonin Bal; Bruno Lina; Gregory Destras; Gwendolyne Burfin; Hadrien Règue; Laetitia Ninove; Laurence Josset; Léa Luciani; Martine Valette; Quentin Semanas |
| EPI_ISL_418959 | Universidade Federal do Rio de Janeiro - UFRJ | Bioinformatics Laboratory / LNCC | Alexandra Gerber; Amilcar Tanuri; Ana Paula Guimarães; Cristiano Xavier Lima; Cássia Cristina Alves Gonçalves; Diana Mariani; Ester Cerdeira Sabino; Filipe Romero; Ingra Morales Claro; Isabela de Carvalho Leitão; Jaqueline Goes de Jesus; Laboratorio Hermes Pardini; Laboratorio Simile; Luciana Jesus Costa e André Felipe Santos; Luiz Gonzaga Paula de Almeida; Lídia Boullosa; Mariane Talon; Mauro Teixeira; Nuno Rodrigues Faria; Orlando Ferreira Jr.; Pinto Castiñeiras; Rafael de Mello Galliez; Renato Santana Aguiar; Richard Araújo Maia; Rodrigo Brindeiro; Ronaldo da Silva Francisco Junior; Terezinha Marta Pereira; e Ana Tereza Vasconcelos; Érica Ramos dos Santos Nascimento |
| EPI_ISL_812967 | University Clinical Research Center, University of Sciences | University Clinical Research Center, University of Sciences | A.A.; Bane, S.; Dao, S.; Diakite, M.; Diarra, B.; Doumbia, S.; Guindo, I.; Iknane; Kone, A. |
| EPI_ISL_528239 | University Hospital Basel, Clinical Virology | University Hospital Basel, Clinical Bacteriology | Adrian Egli; Alexander Gensch; Alfredo Mari; Christian Nickel; Hans Hirsch; Hans Pargger; Helena MB Seth-Smith; Julia Bielicki; Karoline Leuzinger; Kirstine K. Soegaard; Madlen Stange; Manuel Battegay; Martin Siegemund; Michael Osthoff; Michael Schweitzer; Myrta Brunner; Rita Schneider-Sliwa; Roland Bingisser; Sarah Tschudin-Sutter; Simon Fuchs; Stefano Bassetti; Tim Roloff |
| EPI_ISL_775610, EPI_ISL_775611, EPI_ISL_775623, EPI_ISL_775624, EPI_ISL_775626, EPI_ISL_775645, EPI_ISL_775660, EPI_ISL_775666, EPI_ISL_775695, EPI_ISL_775699, EPI_ISL_775708, EPI_ISL_775725, EPI_ISL_775755, EPI_ISL_775761, EPI_ISL_775765, EPI_ISL_775778, EPI_ISL_775783, EPI_ISL_775785, EPI_ISL_775786, EPI_ISL_775797, EPI_ISL_775807, EPI_ISL_775809, EPI_ISL_775825, EPI_ISL_775832, EPI_ISL_775845, EPI_ISL_775863, EPI_ISL_775873, EPI_ISL_775890, EPI_ISL_775897, EPI_ISL_775909, EPI_ISL_775938, EPI_ISL_776067, EPI_ISL_776071, EPI_ISL_776088, EPI_ISL_776169, EPI_ISL_776174, EPI_ISL_776175, EPI_ISL_776176, EPI_ISL_776177, EPI_ISL_776186, EPI_ISL_776187, EPI_ISL_776189, EPI_ISL_776190, EPI_ISL_776211, EPI_ISL_776222, EPI_ISL_776230, EPI_ISL_776231, EPI_ISL_776234, EPI_ISL_776237, EPI_ISL_776242, EPI_ISL_776243, EPI_ISL_776244, EPI_ISL_776245, EPI_ISL_776248, EPI_ISL_776256, EPI_ISL_776257, EPI_ISL_776258, EPI_ISL_776259, EPI_ISL_776260, EPI_ISL_776261, EPI_ISL_776263, EPI_ISL_776264, EPI_ISL_776266, EPI_ISL_776267, EPI_ISL_776269, EPI_ISL_776270, EPI_ISL_776271, EPI_ISL_776272, EPI_ISL_776273, EPI_ISL_776274, EPI_ISL_776275, EPI_ISL_776283, EPI_ISL_776287, EPI_ISL_776298, EPI_ISL_776306, EPI_ISL_776309, EPI_ISL_776310, EPI_ISL_776314, EPI_ISL_776316, EPI_ISL_776317, EPI_ISL_776319, EPI_ISL_776322, EPI_ISL_776335, EPI_ISL_776340, EPI_ISL_776363, EPI_ISL_776366, EPI_ISL_776378, EPI_ISL_776379, EPI_ISL_776380, EPI_ISL_776382, EPI_ISL_776386, EPI_ISL_776389, EPI_ISL_776390, EPI_ISL_776391, EPI_ISL_776395, EPI_ISL_776396, EPI_ISL_776397, EPI_ISL_776400, EPI_ISL_776403, EPI_ISL_776405, EPI_ISL_776406, EPI_ISL_776410, EPI_ISL_776414, EPI_ISL_776419, EPI_ISL_776424, EPI_ISL_776444, EPI_ISL_776445, EPI_ISL_776446, EPI_ISL_776450, EPI_ISL_776550, EPI_ISL_776560, EPI_ISL_776562, EPI_ISL_776565 | | | |
| see above | University Medical Center Hamburg Eppendorf | Heinrich Pette Institute, Leibniz Institute for Experimental Virology | Adam Grundhoff; Alexis Robitaille; Johannes Knobloch; Martin Aepfelbacher; Nicole Fischer; Thomas Günther |
| EPI_ISL_444298, EPI_ISL_444300, EPI_ISL_529615 | University of Birmingham | COVID-19 Genomics UK (COG-UK) Consortium | Alex Richter; Andrew Bosworth; Andrew Bosworth. Queen Elizabeth Hospital: Anna Casey; Andrew D Beggs // PHE Heartlands Lab: Husam Osman; Andrew D Beggs PHE Heartlands Lab: Husam Osman; Charlotte Poxon; Institute of Microbiology; Joanne Stockton; Josh Quick; Kasun Wanigasooriya; Loman Lab: Claire McMurray; Mike Kidd; Nicholas Loman // UHB Lab: Celina M Whalley; Nicholas Loman. University of Birmingham Testing Laboratory: Celina M Whalley; Oliver Pickles; Radoslaw Poplawski; Samuel Nicholls; University of Birmingham: Claire McMurray; Will Rowe |
| EPI_ISL_457124, EPI_ISL_457163, EPI_ISL_457171, EPI_ISL_471968, EPI_ISL_471970 | University of Exeter | COVID-19 Genomics UK (COG-UK) Consortium | Aaron Jeffries; Audrey Farbos; Ben Temperton; Jane Masoli; Joanna Warwick-Dugdale; Michelle Michelsen; Robyn Manley; Stephen Michell |
| EPI_ISL_484710, EPI_ISL_484731, EPI_ISL_484735, EPI_ISL_484737, EPI_ISL_484751, EPI_ISL_484752 | University of Michigan Clinical Microbiology Laboratory | Lauring Lab, University of Michigan, Department of Microbiology and Immunology | Valesano et al. |
| EPI_ISL_485002 | University of Ulsan College of Medicine and Asan Medical Center | University of Ulsan College of Medicine and Asan Medical Center | Heungsup Sung; Jaewoong Lee; Jina Lee; Jiwon Jung; Jongsik Chun; Kihyun Lee; Kuenyoul Park; Kyu-Hwa Hur; Mauricio Chalita; Mi-Na Kim; Seok-Hwan Yoon; Sung-Han Kim; and Hae Kyung Lee |
| EPI_ISL_524444 | University of Washington Virology Lab | Fred Hutchinson Cancer Research Center | Greninger, A.; Jerome, K.; Roychoudhury, P. |
| EPI_ISL_524438, EPI_ISL_524439, EPI_ISL_524442 | University of Washington Virology Lab | Laboratory Medicine, University of Washington | Greninger, A.; Jerome, K.; Roychoudhury, P. |
| EPI_ISL_515274, EPI_ISL_515277, EPI_ISL_515280, EPI_ISL_515286 | University of Washington Virology Lab | University of Washington Virology Lab | Alexander Greninger; Amin Addetia; Hong Xie; Keith R Jerome; Lasata Shrestha; Meei-Li Huang; Pavitra Roychoudhury; Truong Nguyen; Victoria M Rachleff |
| EPI_ISL_529155, EPI_ISL_529156, EPI_ISL_529157, EPI_ISL_529200 | University of Washington, Laboratory Medicine | University of Washington, Laboratory Medicine | Greninger, A.; Jerome, K.; Roychoudhury, P. |
| EPI_ISL_418809 | University of Wisconsin - Madison: Influenza Research Institute | University of Wisconsin Madison, AIDS Vaccine Research Laboratories | Gage Moreno; Katarina Braun; Peter Halfmann; et al. |
| EPI_ISL_417200, EPI_ISL_417201, EPI_ISL_417203, EPI_ISL_417504, EPI_ISL_417507, EPI_ISL_417514, EPI_ISL_421286, EPI_ISL_421291, EPI_ISL_421292, EPI_ISL_421296, EPI_ISL_421297, EPI_ISL_421308, EPI_ISL_421309, EPI_ISL_421317, EPI_ISL_421319, EPI_ISL_421324, EPI_ISL_421325, EPI_ISL_421327, EPI_ISL_421328, EPI_ISL_421329, EPI_ISL_421331, EPI_ISL_421334, EPI_ISL_421335, EPI_ISL_425173, EPI_ISL_425174, EPI_ISL_428262, EPI_ISL_428263, EPI_ISL_428280, EPI_ISL_428285, EPI_ISL_428314, EPI_ISL_428331, EPI_ISL_428341, EPI_ISL_428726, EPI_ISL_480381, EPI_ISL_480387 | | | |
| see above | University of Wisconsin-Madison AIDS Vaccine Research Laboratories | University of Wisconsin-Madison AIDS Vaccine Research Laboratories | Gage Moreno; Katarina Braun; et al. AIDS Vaccine Research Laboratories |
| EPI_ISL_977366 | University of Zambia, School of Veterinary Medicine | UNZAVET and PATH | Daniel Bridges; Mulenga Mwenda-Chimfwembe; Ngonda Saasa |
| EPI_ISL_415541, EPI_ISL_415542, EPI_ISL_417028, EPI_ISL_417960, EPI_ISL_417964, EPI_ISL_417966, EPI_ISL_420806, EPI_ISL_420808, EPI_ISL_421561, EPI_ISL_435349, EPI_ISL_435351, EPI_ISL_435352, EPI_ISL_450539, EPI_ISL_575034, EPI_ISL_575035, EPI_ISL_575036 | | | |
| see above | Utah Public Health Laboratory | Utah Public Health Laboratory | Erin Young; Kelly Oakeson |
| EPI_ISL_603049 | Utah Public Health Laboratory, Utah Public Health Laboratory Infectious Disease submission group | Utah Public Health Laboratory, Utah Public Health Laboratory Infectious Disease submission group | E.L.; Oakeson, K.; Young |
| EPI_ISL_424893, EPI_ISL_424907 | VA-Division of Consolidated Laboratory Services | Pathogen Discovery, Respiratory Viruses Branch, Division of Viral Diseases, Centers for Disease Control and Prevention | Alison S. Laufer Halpin; Anna Uehara; Christopher A. Elkins; Clinton R. Paden; Haibin Wang; Jing Zhang; Krista Queen; Mary S. Keckler; Rachel Marine; Suxiang Tong; Yan Li; Ying Tao |
| EPI_ISL_464167 | VI-US Virgin Islands Department of Health | Pathogen Discovery, Respiratory Viruses Branch, Division of Viral Diseases, Centers for Disease Control and Prevention | Alison S. Laufer Halpin; Anna Uehara; Christopher A. Elkins; Clinton R. Paden; Haibin Wang; Jasmine Padilla; Jing Zhang; Justin Lee; Krista Queen; Mary S. Keckler; Suxiang Tong; Yan Li; Ying Tao |
| EPI_ISL_2367400 | VIOLLIER MORGES SA | Laboratory of genomics and metagenomics | Claire Bertelli; Damien Jacot; Gilbert Greub; Sébastien Aeby; Trestan Pillonel |
| EPI_ISL_447846 | VT Dept. of Health Laboratory | Pathogen Discovery, Respiratory Viruses Branch, Division of Viral Diseases, Centers for Disease Control and Prevention | Alison S. Laufer Halpin; Anna Uehara; Christopher A. Elkins; Clinton R. Paden; Haibin Wang; Jasmine Padilla; Jing Zhang; Justin Lee; Krista Queen; Mary S. Keckler; Suxiang Tong; Yan Li; Ying Tao |
| EPI_ISL_413486 | Valley Medical Center | University of Washington Virology Lab | Alexander Greninger; Arun Nalla; Hong Xie; Keith Jerome; Pavitra Roychoudhury |
| EPI_ISL_468495, EPI_ISL_548349 | Ventura County Public Health Lab | Chan-Zuckerberg Biohub | CZB Cliahub Consortium |
| EPI_ISL_471471 | Victorian Infectious Diseases Reference Laboratory | Victorian Infectious Diseases Reference Laboratory | Antonia da Costa, E.; Barreto, I.; Caly, L.; Canisia, D.; Dakh, F.; Dolores de Jesus da Costa, M.; Douglas; Druce, J.; Francis, J.; Freeman, K.; Jayanti Pereira Tilman, A.; M.B.; Marr, I.; N. Meumann, E.; N. and Baird, R.; Sait, M.; Salles de Sousa, A.; Schultz; Seemann, T.; Sherry; Soares da Silva, E.; Wapling, J.; Ximenes, J. |
| EPI_ISL_426640, EPI_ISL_426644, EPI_ISL_426645, EPI_ISL_426647, EPI_ISL_426658, EPI_ISL_426659, EPI_ISL_426663, EPI_ISL_426666, EPI_ISL_426667, EPI_ISL_426672, EPI_ISL_426673, EPI_ISL_426675, EPI_ISL_426686, EPI_ISL_426688, EPI_ISL_426693, EPI_ISL_426715, EPI_ISL_426723, EPI_ISL_426730, EPI_ISL_426732, EPI_ISL_426733, EPI_ISL_426762, EPI_ISL_426763, EPI_ISL_426773, EPI_ISL_426794, EPI_ISL_426798, EPI_ISL_426825, EPI_ISL_426828, EPI_ISL_426841, EPI_ISL_426842, EPI_ISL_426845, EPI_ISL_426852, EPI_ISL_426853, EPI_ISL_426855, EPI_ISL_426856, EPI_ISL_426857, EPI_ISL_426869, EPI_ISL_426877, EPI_ISL_426878, EPI_ISL_426927, EPI_ISL_426942, EPI_ISL_426943, EPI_ISL_426945, EPI_ISL_426952, EPI_ISL_426958, EPI_ISL_426965, EPI_ISL_426967, EPI_ISL_426971, EPI_ISL_426981, EPI_ISL_426998, EPI_ISL_427012, EPI_ISL_427013, EPI_ISL_427015, EPI_ISL_427019, EPI_ISL_427024, EPI_ISL_427039, EPI_ISL_427046, EPI_ISL_427052 | | | |
| see above | Victorian Infectious Diseases Reference Laboratory (VIDRL) | Microbiological Diagnostic Unit Public Health Laboratory and Victorian Infectious Diseases Reference Laboratory, Doherty Institute | Caly L.; Druce J.; Sait, M.; Schultz M.; Seemann T.; Sherry, N. |
| EPI_ISL_430477, EPI_ISL_430479, EPI_ISL_430483, EPI_ISL_430527, EPI_ISL_430542, EPI_ISL_430543 | Victorian Infectious Diseases Reference Laboratory (VIDRL) | Microbiological Diagnostic Unit Public Health Laboratory and Victorian Infectious Diseases Reference Laboratory, The Peter Doherty Institute for Infection and Immunity | Caly L.; Druce J.; Sait, M.; Schultz M.; Seemann T.; Sherry, N. |
| EPI_ISL_521898, EPI_ISL_521899, EPI_ISL_563817 | Victorian Infectious Diseases Reference Laboratory (VIDRL) | VIDRL and MDU-PHL | Caly L.; Caly, L.; Druce J.; M. B.; Sait, M.; Schultz; Schultz M.; Seemann T.; Seemann, T.; Sherry, N. |
| EPI_ISL_416413, EPI_ISL_416415, EPI_ISL_416514, EPI_ISL_419733, EPI_ISL_419736, EPI_ISL_419739, EPI_ISL_419741, EPI_ISL_419746, EPI_ISL_419747, EPI_ISL_419751, EPI_ISL_419752, EPI_ISL_419755, EPI_ISL_419758, EPI_ISL_419760, EPI_ISL_419769, EPI_ISL_419772, EPI_ISL_419777, EPI_ISL_419779, EPI_ISL_419781, EPI_ISL_419782, EPI_ISL_419783, EPI_ISL_419785, EPI_ISL_419788, EPI_ISL_419789, EPI_ISL_419795, EPI_ISL_419796, EPI_ISL_419805, EPI_ISL_419812, EPI_ISL_419814, EPI_ISL_419816, EPI_ISL_419821, EPI_ISL_419826, EPI_ISL_419834, EPI_ISL_419836, EPI_ISL_419837, EPI_ISL_419847, EPI_ISL_419856, EPI_ISL_419857, EPI_ISL_419859, EPI_ISL_419862, EPI_ISL_419870, EPI_ISL_419874, EPI_ISL_419885, EPI_ISL_419899, EPI_ISL_419900, EPI_ISL_419904, EPI_ISL_419908, EPI_ISL_419910, EPI_ISL_419917, EPI_ISL_419920, EPI_ISL_419924, EPI_ISL_419927, EPI_ISL_419936, EPI_ISL_419939, EPI_ISL_419958, EPI_ISL_419970, EPI_ISL_419979, EPI_ISL_419984, EPI_ISL_419986, EPI_ISL_419995, EPI_ISL_420036 | | | |
| see above | Victorian Infectious Diseases Reference Laboratory (VIDRL) | Victorian Infectious Diseases Reference Laboratory and Microbiological Diagnostic Unit Public Health Laboratory, Doherty Institute | Caly L.; Druce J.; Sait, M.; Schultz M.; Seemann T.; Sherry, N.; Taiaroa, G. |

| | | | |
|---|---|---|---|
| EPI_ISL_451727 | Viollier AG | Department of Biosystems Science and Engineering, ETH Zürich | Christian Beisel; Christiane Beckmann; Christoph Noppen; Elodie Burcklen; Ina Nissen; Ivan Topolsky; Maurice Redondo; Natascha Santacroce; Niko Beerenwinkel; Noemie Santamaria de Souza; Olivier Kobel; Pedro Ferreira; Philipp Jablonski; Sarah Nadeau; Sophie Seidel; Susana Posada-Céspedes; Tanja Stadler; Tobias Schär |
| EPI_ISL_420841, EPI_ISL_961010 | Viral Respiratory Lab, National Institute for Biomedical Research (INRB) | Pathogen Sequencing Lab, National Institute for Biomedical Research (INRB) | Allison Black; Amuri Aziza; Andrew Rambaut; Catherine Pratt; Eddy Kinganda-Lusamaki; Edith Nkwembe; Emmanuel Lokilo Lofiko; Francisca Muyembe Mawete; Ian Goodfellow; James Hadfield; Jean-Claude Makangara Cigolo; Jean-Jacques Muyembe Tamfum; Josh Quick; Kristian Andersen; Matthias Pauthner; Michael Wiley; Nick Loman; Placide Mbala-Kingebeni; Steve Ahuka-Mundeke; Trevor Bedford |
| EPI_ISL_418864, EPI_ISL_418956, EPI_ISL_418957, EPI_ISL_418958, EPI_ISL_419262, EPI_ISL_419263, EPI_ISL_419711, EPI_ISL_419713, EPI_ISL_420028, EPI_ISL_420029, EPI_ISL_420628, EPI_ISL_429970, EPI_ISL_429971, EPI_ISL_485838, EPI_ISL_485839, EPI_ISL_485842, EPI_ISL_485844, EPI_ISL_485845, EPI_ISL_526804, EPI_ISL_526811, EPI_ISL_526815, EPI_ISL_526819 | | | |
| see above | Virginia DCLS | Virginia DCLS | Virginia DCLS |
| EPI_ISL_710487, EPI_ISL_710488 | Virology (microbiology), Hospital Universitario Central de Asturias | Virology (microbiology), Hospital Universitario Central de Asturias | Abreu, F.; Alvarez-Argueelles; Boga; Castello, C.; Costales, I.; Coto, E.; Gomez-deOna, J.; J.A.; M.E.; Martin-Rodriguez, G.; Melon, S.; Perez-Martinez, Z.; Rojo, S.; Sandoval, M. |
| EPI_ISL_487822, EPI_ISL_491545, EPI_ISL_491689 | Virology Department, Royal Infirmary of Edinburgh, NHS Lothian / School of Biological Sciences, University of Edinburgh | Wellcome Sanger Institute for the COVID-19 Genomics UK (COG-UK) consortium | Colquhoun R; Cordelia Langford; David K. Jackson; Dewar R; Dominic Kwiatkowski; Ewan Harrison; Hill V; Ian Johnston; Jackson B; John Sillitoe on behalf of the Wellcome Sanger Institute COVID-19 Surveillance Team (http://www.sanger.ac.uk/covid-team); McCrone JT; McHugh M; O'Toole Á; Rambaut A; Roberto Amato; Rooke S; Scher E; Sonia Goncalves; Templeton K and Alex Alderton; Yu X |
| EPI_ISL_425948, EPI_ISL_425972, EPI_ISL_425977, EPI_ISL_426007, EPI_ISL_426008, EPI_ISL_426015 | Virology Department, Royal Infirmary of Edinburgh, NHS Lothian / School of Biological Sciences, University of Edinburgh / Institute of Genetics and Molecular Medicine, University of Edinburgh | COVID-19 Genomics UK (COG-UK) Consortium | Balcaza C; Colquhoun R; Dewar R; Gallagher M; Hill V; Jackson B; McCrone JT; McHugh M; O'Toole A; Rambaut A; Rooke S; Scher E; Templeton K; Williams TC; Yu X |
| EPI_ISL_493203, EPI_ISL_493204, EPI_ISL_493206 | Virology Lab,Department of Pathology, National Cheng Kung University Hospital | Virology Lab,Department of Pathology, National Cheng Kung University Hospital | Huey-Pin Tsai; et al |
| EPI_ISL_610245 | Virology, National Institute for Biological Standards and Control | Virology, National Institute for Biological Standards and Control | Bentley, E.; E.T.; Fritzsche; M. and Mattiuzzo, G.; Mate, R.; Mee; Routley, S. |
| EPI_ISL_418771, EPI_ISL_418772, EPI_ISL_418773, EPI_ISL_418774, EPI_ISL_418775, EPI_ISL_418776, EPI_ISL_418777, EPI_ISL_418778, EPI_ISL_418779, EPI_ISL_418780, EPI_ISL_418781, EPI_ISL_418782, EPI_ISL_418783, EPI_ISL_418784, EPI_ISL_418785, EPI_ISL_418786, EPI_ISL_418787, EPI_ISL_418788, EPI_ISL_418789, EPI_ISL_418790, EPI_ISL_418791, EPI_ISL_426436, EPI_ISL_426437, EPI_ISL_426438, EPI_ISL_426439, EPI_ISL_426440, EPI_ISL_426441, EPI_ISL_426442, EPI_ISL_426443, EPI_ISL_426444, EPI_ISL_426445, EPI_ISL_426446, EPI_ISL_426447, EPI_ISL_426448, EPI_ISL_426449, EPI_ISL_426450, EPI_ISL_426451, EPI_ISL_426452, EPI_ISL_426453 | | | |
| see above | WA State Department of Health | Pathogen Discovery, Respiratory Viruses Branch, Division of Viral Diseases, Centers for Disease Control and Prevention | Anna Uehara; Brian Hiatt; Clinton R. Paden; Denny Russell; Haibin Wang; Jesica Jacobs; Jessica Gant; Jing Zhang; Krista Queen; Suxiang Tong; Yan Li; Ying Tao |
| EPI_ISL_525607, EPI_ISL_525609, EPI_ISL_525611, EPI_ISL_525613, EPI_ISL_525621, EPI_ISL_525625, EPI_ISL_525626, EPI_ISL_525630, EPI_ISL_534731 | | | |
| see above | Wadsworth Center, New York State Department of Health | Wadsworth Center, New York State Department of Health | Daryl M. Lamson; Erica Lasek-Nesselquist; Jonathan Plitnick; Kirsten St. George; Matthew D. Shudt; Navjot Singh; Sara Griesemer |
| EPI_ISL_426302, EPI_ISL_676669, EPI_ISL_676689, EPI_ISL_676699, EPI_ISL_676704, EPI_ISL_676734, EPI_ISL_676740, EPI_ISL_676743, EPI_ISL_676756, EPI_ISL_676762 | | | |
| see above | Wadsworth Center, New York State Department.of Health | Wadsworth Center, New York State Department.of Health | Alexis Russel; Daryl M. Lamson; Erasmus Schneider; Erica Lasek-Nesselquist; John Kelly; Jonathan Plitnick; Kirsten St. George; Matthew D. Shudt; Navjot Singh; Sara Griesemer |
| EPI_ISL_418113, EPI_ISL_418146, EPI_ISL_422299, EPI_ISL_445804, EPI_ISL_445922 | Wales Specialist Virology Centre | Public Health Wales Microbiology Cardiff | Alec Birchley; Alexander Adams; Amy Gaskin; Bree Gatica-Wilcox; Catherine Moore; Jason Coombes; Joanne Watkins; Johnathan Evans; Laura Gifford; Lauren Gilbert; Lee Graham; Malorie Perry; Matt Bull; Matthew Bull; Nicole Pacchiarini; Sally Corden; Sara Kumziene-Summerhayes; Sara Rey; Sarah Taylor; Simon Cottrell; Sophie Jones; Tom Connor |
| EPI_ISL_407214, EPI_ISL_407215 | Washington State Department of Health | Pathogen Discovery, Respiratory Viruses Branch, Division of Viral Diseases, Centers for Dieases Control and Prevention | Anna Uehara; Azaibi Tamin; Clinton R. Paden; Haibin Wang; Janna' R. Murray; Jennifer Harcourt; Jing Zhang; Krista Queen; Natalie Thornburg; Rashi Gautam; Shifaq Kamili; Stephen Lindstrom; Susan I. Gerber; Suxiang Tong; Xiaoyan Lu; Yan Li; Ying Tao |
| EPI_ISL_412970, EPI_ISL_417065, EPI_ISL_417066, EPI_ISL_417067, EPI_ISL_417068, EPI_ISL_417069, EPI_ISL_417070, EPI_ISL_417071, EPI_ISL_417072, EPI_ISL_417073, EPI_ISL_417074, EPI_ISL_417075, EPI_ISL_417076, EPI_ISL_417077, EPI_ISL_417078, EPI_ISL_417079, EPI_ISL_417080, EPI_ISL_417081, EPI_ISL_417082, EPI_ISL_417083, EPI_ISL_417084, EPI_ISL_417085, EPI_ISL_417086, EPI_ISL_417087, EPI_ISL_417088, EPI_ISL_417089, EPI_ISL_417090, EPI_ISL_417091, EPI_ISL_417092, EPI_ISL_417093, EPI_ISL_417094, EPI_ISL_417095, EPI_ISL_417096, EPI_ISL_417097, EPI_ISL_417098, EPI_ISL_417099, EPI_ISL_417100, EPI_ISL_417101, EPI_ISL_417102, EPI_ISL_417103, EPI_ISL_417104, EPI_ISL_417105, EPI_ISL_417106, EPI_ISL_417107, EPI_ISL_417108, EPI_ISL_417109, EPI_ISL_417110, EPI_ISL_417111, EPI_ISL_417112, EPI_ISL_417114, EPI_ISL_417115, EPI_ISL_417116, EPI_ISL_417117, EPI_ISL_417118, EPI_ISL_417119, EPI_ISL_417120, EPI_ISL_417121, EPI_ISL_417122, EPI_ISL_417123, EPI_ISL_417124, EPI_ISL_417125, EPI_ISL_417126, EPI_ISL_417127, EPI_ISL_417128, EPI_ISL_417129, EPI_ISL_417130, EPI_ISL_417131, EPI_ISL_417132, EPI_ISL_417133, EPI_ISL_417134, EPI_ISL_417135, EPI_ISL_417136, EPI_ISL_417137, EPI_ISL_417139, EPI_ISL_417140, EPI_ISL_417141, EPI_ISL_417142, EPI_ISL_417143, EPI_ISL_417144, EPI_ISL_417145, EPI_ISL_417146, EPI_ISL_417147, EPI_ISL_417148, EPI_ISL_417149, EPI_ISL_417150, EPI_ISL_417151, EPI_ISL_417152, EPI_ISL_417153, EPI_ISL_417154, EPI_ISL_417155, EPI_ISL_417156, EPI_ISL_417157, EPI_ISL_417158, EPI_ISL_417159, EPI_ISL_417160, EPI_ISL_417161, EPI_ISL_417162, EPI_ISL_417166, EPI_ISL_417167, EPI_ISL_417168, EPI_ISL_417169, EPI_ISL_417170, EPI_ISL_417171, EPI_ISL_417172, EPI_ISL_417173, EPI_ISL_417174, EPI_ISL_417175, EPI_ISL_430160, EPI_ISL_430161, EPI_ISL_430162, EPI_ISL_430163, EPI_ISL_430164, EPI_ISL_430165, EPI_ISL_430166, EPI_ISL_430167, EPI_ISL_430168, EPI_ISL_430169, EPI_ISL_430170, EPI_ISL_430171, EPI_ISL_430172, EPI_ISL_430173, EPI_ISL_430174, EPI_ISL_430175, EPI_ISL_430176, EPI_ISL_430177, EPI_ISL_430178, EPI_ISL_430179, EPI_ISL_430180, EPI_ISL_430181, EPI_ISL_430182, EPI_ISL_430184, EPI_ISL_430187, EPI_ISL_430188, EPI_ISL_430189, EPI_ISL_430190, EPI_ISL_430191, EPI_ISL_430192, EPI_ISL_430193, EPI_ISL_430194, EPI_ISL_430196, EPI_ISL_430197, EPI_ISL_430198, EPI_ISL_430199, EPI_ISL_430203, EPI_ISL_430204, EPI_ISL_430208, EPI_ISL_430209, EPI_ISL_430210, EPI_ISL_430211, EPI_ISL_430212, EPI_ISL_430213, EPI_ISL_430214, EPI_ISL_430215, EPI_ISL_430216, EPI_ISL_430219, EPI_ISL_430221, EPI_ISL_430222, EPI_ISL_430223, EPI_ISL_430225, EPI_ISL_430228, EPI_ISL_430231, EPI_ISL_430232, EPI_ISL_430234, EPI_ISL_430238, EPI_ISL_430239, EPI_ISL_430245, EPI_ISL_430248, EPI_ISL_430249, EPI_ISL_430250, EPI_ISL_430251, EPI_ISL_430253, EPI_ISL_430257, EPI_ISL_430259, EPI_ISL_430260, EPI_ISL_430262, EPI_ISL_430265, EPI_ISL_430267, EPI_ISL_430268, EPI_ISL_430270, EPI_ISL_430271, EPI_ISL_430272, EPI_ISL_430273, EPI_ISL_430274, EPI_ISL_430275, EPI_ISL_430276, EPI_ISL_430277, EPI_ISL_430278, EPI_ISL_430279, EPI_ISL_430280, EPI_ISL_430281, EPI_ISL_430282, EPI_ISL_430283, EPI_ISL_430284, EPI_ISL_430285, EPI_ISL_430286, EPI_ISL_430287, EPI_ISL_430288, EPI_ISL_430289, EPI_ISL_430290, EPI_ISL_430291, EPI_ISL_430292, EPI_ISL_430293, EPI_ISL_430294, EPI_ISL_430295, EPI_ISL_430296, EPI_ISL_434078, EPI_ISL_434081, EPI_ISL_434082, EPI_ISL_434111, EPI_ISL_434112, EPI_ISL_434113, EPI_ISL_434114, EPI_ISL_434115, EPI_ISL_434116, EPI_ISL_434117, EPI_ISL_434118, EPI_ISL_434119, EPI_ISL_434120, EPI_ISL_434128, EPI_ISL_434129, EPI_ISL_434132, EPI_ISL_434134, EPI_ISL_434135, EPI_ISL_434136, EPI_ISL_434137, EPI_ISL_434140, EPI_ISL_434143, EPI_ISL_434145, EPI_ISL_434146, EPI_ISL_434151, EPI_ISL_434153, EPI_ISL_434154, EPI_ISL_434156, EPI_ISL_434157, EPI_ISL_434159, EPI_ISL_434160, EPI_ISL_434161, EPI_ISL_434162, EPI_ISL_434163, EPI_ISL_434165, EPI_ISL_434167, EPI_ISL_434169, EPI_ISL_434170, EPI_ISL_434171, EPI_ISL_434172, EPI_ISL_434174, EPI_ISL_434175, EPI_ISL_434176, EPI_ISL_434177, EPI_ISL_434178, EPI_ISL_434179, EPI_ISL_434180, EPI_ISL_434182, EPI_ISL_434183, EPI_ISL_434184, EPI_ISL_434185, EPI_ISL_434186, EPI_ISL_434187, EPI_ISL_434188, EPI_ISL_434189, EPI_ISL_434190, EPI_ISL_434192, EPI_ISL_434193, EPI_ISL_434196, EPI_ISL_434197, EPI_ISL_434201, EPI_ISL_434202, EPI_ISL_434210, EPI_ISL_434211, EPI_ISL_434212, EPI_ISL_434214, EPI_ISL_434215, EPI_ISL_434247, EPI_ISL_434250, EPI_ISL_434252, EPI_ISL_434256, EPI_ISL_434257, EPI_ISL_434259, EPI_ISL_434263, EPI_ISL_434265, EPI_ISL_434269, EPI_ISL_434270, EPI_ISL_434279, EPI_ISL_434280, EPI_ISL_434286, EPI_ISL_434287, EPI_ISL_434288, EPI_ISL_434289, EPI_ISL_434298, EPI_ISL_434299, EPI_ISL_434302, EPI_ISL_434305, EPI_ISL_434306, EPI_ISL_434308, EPI_ISL_434313, EPI_ISL_434315, EPI_ISL_434316, EPI_ISL_434323, EPI_ISL_434327, EPI_ISL_434328, EPI_ISL_434332, EPI_ISL_434333, EPI_ISL_434334, EPI_ISL_434335, EPI_ISL_434336, EPI_ISL_434337, EPI_ISL_434338, EPI_ISL_434339, EPI_ISL_434342, EPI_ISL_434343, EPI_ISL_434344, EPI_ISL_434345, EPI_ISL_434346 | | | |
| see above | Washington State Department of Health | Seattle Flu Study | Barry Lutz; Chu et al; Chu etl al; Deborah Nickerson; Helen Chu; Janet Englund; Jay Shendure; Lea Starita; Mark Rieder; Matthew Thompson; Michael Boeckh; Michael Famulare; and Trevor Bedford |
| EPI_ISL_413457, EPI_ISL_413458 | Washington State Public Health Lab | UW Virology Lab | Alexander Greninger; Arun Nalla; Hong Xie; Keith Jerome; Pavitra Roychoudhury |
| EPI_ISL_413455 | Washington State Public Health Lab | University of Washington Virology Lab | Alexander Greninger; Arun Nalla; Hong Xie; Keith Jerome; Pavitra Roychoudhury |
| EPI_ISL_594447, EPI_ISL_594448, EPI_ISL_594449, EPI_ISL_594450, EPI_ISL_594451, EPI_ISL_594452, EPI_ISL_594453, EPI_ISL_594454, EPI_ISL_594459 | | | |
| see above | Washington State Public Health Laboratories | Pathogen Discovery, Respiratory Viruses Branch, Division of Viral Diseases, Centers for Disease Control and Prevention | Anna Uehara; Clinton Paden; Haibin Wang; Jing Zhang; Julu Bhatnagar; Krista Queen; Suxiang Tong; Yan Li; Ying Tao |
| EPI_ISL_416538, EPI_ISL_416539 | Wellington Hospital | Institute of Environmental Science and Research (ESR) | Craig Thornley; Erasmus Smit; Joep de Ligt; Lauren Jelly; Matt Storey; Maxim Bloomfield; New Zealand; Newtown; Riddiford Street; Wellington 6021; Wellington Hospital; Wellington SCL; Xiaoyun Ren |
| EPI_ISL_456182 | Wellington SCL | Institute of Environmental Science and Research (ESR) | Anja Werno; Antje van der Linden; Arlo Upton; Chris Mansell; David Hammer; Dragana Drinkovic; Erasmus Smit; Gary McAuliffe; Hana Sofia Andersson; James Ussher; Jill Sherwood; Joep de Ligt; Josh Freeman; Julia Howard; Juliet Elvy; Lauren Jelly; Mary DeAlmeida; Matt Blakiston; Matt Storey; Matthew Rogers; Max Bloomfield; Michael Addidle; Michelle Balm; Sally Roberts; Sarah Jefferies; Sharmini Muttaiyah; Susan Morpeth; Susan Taylor; Timothy Blackmore; Vani Sathyendran; Veronica Playle; Virginia Hope; Xiaoyun Ren |
| EPI_ISL_451346, EPI_ISL_451348, EPI_ISL_451354, EPI_ISL_451360, EPI_ISL_451365, EPI_ISL_451368, EPI_ISL_451370, EPI_ISL_451371, EPI_ISL_451383, EPI_ISL_451384 | | | |
| see above | West China Hospital of Sichuan University | State Key Laboratory of Biotherapy of Sichuan University | Baowen Du; Binwu Ying; Chao Tang; Chuan Chen; Hancheng Wei; Jia Geng; Jing-wen Lin; Lu Chen; Mingxia Yu; Minjin Wang; Weimin Li; Yongzhao Zhou |
| EPI_ISL_425658, EPI_ISL_425713, EPI_ISL_425718, EPI_ISL_425723, EPI_ISL_425725, EPI_ISL_425735, EPI_ISL_425744, EPI_ISL_425752, EPI_ISL_425781, EPI_ISL_425793, EPI_ISL_433277, EPI_ISL_433298, EPI_ISL_433363, EPI_ISL_433367, EPI_ISL_433506, EPI_ISL_433512, EPI_ISL_433514, EPI_ISL_433515, EPI_ISL_433518, EPI_ISL_433525, EPI_ISL_433541, EPI_ISL_433558, EPI_ISL_438757, EPI_ISL_438766, EPI_ISL_438769, EPI_ISL_438770, EPI_ISL_438777, EPI_ISL_438781, EPI_ISL_438782, EPI_ISL_438783, EPI_ISL_438786, EPI_ISL_438911, EPI_ISL_438930, EPI_ISL_439089, EPI_ISL_439095, EPI_ISL_439099, EPI_ISL_439110, EPI_ISL_439112, EPI_ISL_439114, EPI_ISL_439126, EPI_ISL_439137, EPI_ISL_439139, EPI_ISL_478161, EPI_ISL_493741 | | | |
| see above | West of Scotland Specialist Virology Centre, NHSGGC / MRC-University of Glasgow Centre for Virus Research | COVID-19 Genomics UK (COG-UK) Consortium | Alasdair MacLean; Alice Broos; Ana da Silva Filipe; Antonia Ho; Daniel Mair; David L Robertson; Elihu Aranday-Cortes; Emma Thomson; James Shepherd; Jenna Nichols; Joseph Hughes; Kathy Li; Kathy Smollett; Kirstyn Brunker; Kyriaki Nomikou; Lily Tong; Marc Niebel; Natasha Jesudason; Natasha Johnson; Patawee Asamaphan; Rajiv Shah; Richard Orton; Rory Gunson; Rory Gunson.; Sarah McDonald; Sreenu Vattipally; Stephen Carmichael; Yasmin Parr |
| EPI_ISL_471203, EPI_ISL_471213, EPI_ISL_471222, EPI_ISL_471252, EPI_ISL_471258, EPI_ISL_471266, EPI_ISL_578380, EPI_ISL_578384, EPI_ISL_578419, EPI_ISL_578424, EPI_ISL_578433, EPI_ISL_803306, EPI_ISL_803307, EPI_ISL_803327, EPI_ISL_803638 | | | |
| see above | Wisconsin State Laboratory of Hygiene Communicable Disease Division | Wisconsin State Laboratory of Hygiene Communicable Disease Division | Abigail C. Shockey; Kelsey R. Florek |
| EPI_ISL_2264432, EPI_ISL_2264434 | World Reference Center of Emerging Viruses and Arboviruses (WRCEVA), University of Texas Medical Branch (UTMB), Galveston | ClickSeq Lab, University of Texas Medical Branch, UTMB, Galveston | Dong J.; Jaworski E.; Judy B.; Ksiazek T.; Langsjoen R.M.; Menachery V.D.; Miller A.L.; Newman P.; Plante J.A.; Plante K.S.; Pyles R.B.; Ren P.; Routh A.; Swetnam D.; Weaver S.C; Zhou Y. |
| EPI_ISL_454989, EPI_ISL_454996, EPI_ISL_454998, EPI_ISL_455368, EPI_ISL_455376, EPI_ISL_455406 | Wuhan Chain Medical Labs (CMLabs) | State Key Laboratory of Biotherapy of Sichuan University | Baowen Du; Binwu Ying; Chao Tang; Chuan Chen; Hancheng Wei; Jia Geng; Jing-wen Lin; Lu Chen; Mingxia Yu; Minjin Wang; Weimin Li; Yongzhao Zhou |
| EPI_ISL_412982 | Wuhan Lung Hospital | Hubei Provincial Center for Disease Control and Prevention | Bin Fang; Bo Yang; Bo Yu; Faxian Zhan; Guojun Ye; Jing Li; Junqiang Xu; Kun Cai; Linlin Liu; Xiang Li; Xiao Yu; Xixiang Huo; Yongzhong Jiang. |
| EPI_ISL_631808 | Wyckoff Heights Medical Center | New York City Public Health Laboratory | Jade Wang; et al. |
| EPI_ISL_421272, EPI_ISL_421548, EPI_ISL_421549, EPI_ISL_421552, EPI_ISL_421554, EPI_ISL_462917, EPI_ISL_462922, EPI_ISL_462924, EPI_ISL_462925, EPI_ISL_462933, EPI_ISL_462941 | | | |
| see above | Wyoming Public Health Laboratory | Center for Global Health, University of New Mexico Health Sciences Center | Cari Sloma; Darrell Dinwiddie; Daryl Domman; Kurt Schwalm; Noah Hull; Rob Christensen; Wanda Manley |
| EPI_ISL_539566 | Xerencia de Xestión Integrada de Pontevedra e o Salnés | Instituto de Salud Carlos III | A. Monzón; F. Casas; I; I. Jiménez; I. Pérez; Iglesias-Caballero; M. Camarero; M. Cuesta; M. González-Esguevillas; M. Molinero Calamita; M. Zaballos; P. Jiménez; S. Juliá; S. Pozo; S. Varona |
| EPI_ISL_419513, EPI_ISL_428378 | Yale COVID-19 Biorepository | Grubaugh Lab - Yale School of Public Health | Adam Moore; Akiko Iwasaki; Albert Ko; Alice Lu; Allison Nelson; Anderson Brito; Anne Wyllie; Arnau Casanovas; Catherine Muenker; Chaney Kalinich; Chantal Vogels; Charlese Dela Cruz; Isabel Ott; Joseph Fauver; Maria Tokuyama; Mary Petrone; Nathan Grubaugh; Patrick Wong; Peiwen Lu; Richard Martinello; Saad Omer; Shelli Farhadian; Tara Alpert |
| EPI_ISL_416421, EPI_ISL_416422, EPI_ISL_416423, EPI_ISL_416424, EPI_ISL_419524, EPI_ISL_419526, EPI_ISL_420086, EPI_ISL_420091, EPI_ISL_428370, EPI_ISL_428372, EPI_ISL_428381, EPI_ISL_428382, EPI_ISL_476544 | | | |
| see above | Yale Clinical Virology Laboratory | Grubaugh Lab - Yale School of Public Health | Adam Moore; Akiko Iwasaki; Albert Ko; Alice Lu; Allison Nelson; Anderson Brito; Anne Wyllie; Anthony Muyombwe; Arnau Casanovas; Catherine Muenker; Chaney Kalinich; Chantal Vogels; Charlese Dela Cruz; Cole Jensen; Ellen Foxman; Isabel Ott; Jafar Razeq; Joseph Fauver; Maria Tokuyama; Marie Landry; Marie-Louise Landry; Mary Petrone; Nagarjuna Cheemarla; Nathan Grubaugh; Patrick Wong; Peiwen Lu; Richard Martinello; Saad Omer; Shelli Farhadian; Tara Alpert |

| | | | |
|---|---|---|---|
| EPI_ISL_408478 | Yongchuan District Center for Disease Control and Prevention | Chongqing Municipal Center for Disease Control and Prevention | Chen Shuang; Li Qing; Ling Hua; Rong Rong; Su Kun; Tan ZhangPing; Tang Wenge; Tang Yun; Ye Sheng; Yu zhen |
| EPI_ISL_417618 | deCODE genetics | deCODE genetics | Agnar Helgason; Alma Moller; Arna B Agustsdottir; Arnaldur Gylfason; Asgeir Sigurdsson; Aslaug Jonasdottir; Berglind Eiriksdottir; Bjarni Thorbjornsson; Brynjar O Jensson; Daniel F Gudbjartsson; Droplaug N Magnusdottir; Elisabet E Gardarsdottir; Emil A Thorarensen; Gardar Sveinbjornsson; Gisli Masson; Gudmundur Georgsson; Gudmundur L Norddahl; Gudrun Sigmundsdottir; Hakon Jonsson; Hilma Holm; Ingileif Jonsdottir; Jona Saemundsdottir; Kamilla S Josefsdottir; Kari Stefansson; Karl G Kristinsson; Kjartan R Gudmundsson; Kristin E Sveinsdottir; Louise le Roux; Maney Sveinsdottir; Olafia S Gretarsdottir; Olafur T Magnusson; Pall Melsted; Patrick Sulem; Run Fridriksdottir; Thora R Gunnarsdottir; Thordur Kristjansson; Thorolfur Gudnason; Unnur Thorsteinsdottir |
| EPI_ISL_447911, EPI_ISL_447912, EPI_ISL_447913, EPI_ISL_447918, EPI_ISL_447920 | n/a | National Institute of Health. Department of medical Sciences, Ministry of Public Health, Thailand | Chittaganpitch; Malinee; Okada; Parnmen; Phuygun; Pilailuk; Siripaporn; Sittiporn; Sunthareeya; Thanadachakul; Thanutsapa; Waicharoen; Warawan; Wongboot |
| EPI_ISL_458286 | unknown | Bundeswehr Institute of Microbiology | Antwerpen; Bestehorn-Willmann; Eckstein, S.; Handrick, S.; M.C.; M.H.; M.S.; Naija, H.; R. and Ben Moussa, M.; Rehn, A.; Stoecker, K.; Walter; Woelfel |
| EPI_ISL_463001 | unknown | Clinical virology | Fares, W.; Triki, H. |
| EPI_ISL_468064, EPI_ISL_468065 | unknown | Computer Science and Engineering | A.B.; Adcock; Alejandro, B.; Arnold; Carrico, R.; Chariker; Chung, D.; E.C.; F.W.; Hwang; J.H.; J.W.; J.Y.; K.E.; L.A.; Lasnik; Palmer; Park; R.S.; Ramirez, J.; Rouchka; Waigel, S.; Wolf; Zacharias, W.; Zhang, M. |
| EPI_ISL_447836 | unknown | Department of Medicine | Bampali, M.; Dovrolis, N.; Froukala, E.; Gatzidou, E.; Kassela, K.; N. and Karakasiliotis, I.; Spanakis; Stavropoulou, A.; Tsakris, A.; Veletza, S. |
| EPI_ISL_455683, EPI_ISL_455684, EPI_ISL_455685, EPI_ISL_455687, EPI_ISL_455692, EPI_ISL_468726 | unknown | Department of Microbiology | Bao, L.; Cai, F.; Deng, W.; Gao, H.; Gao, Q.; Ge, X.; Gong, X.; Hu, Y.; Jiang, D.; Jiang, L.; Kan, B.; Li, C.; Li, J.; Li, Y.; Liu, J.; Lou, X.; Lu, J.; Lv, Z.; Mao, H.; Peng, H.; Qi, Z.; Qin, C.; Shi, W.; Sun, Y.; Tang, H.; Wang, L.; Wang, N.; Wang, X.; Wu, D.; Xu, K.; Yang, M.; Yin, W.; Yin, Y.; Zhang, H.; Zhang, Y.; Zhao, P.; Zhu, L. |
| EPI_ISL_449476, EPI_ISL_449477, EPI_ISL_449478, EPI_ISL_449479, EPI_ISL_449480, EPI_ISL_449481, EPI_ISL_449482, EPI_ISL_449483, EPI_ISL_449484, EPI_ISL_449485, EPI_ISL_449486 | | | |
| see above | unknown | Department of Respiratory and Critical Care | Bai, X.; Chang, Y.; Deng, T.; Gao, Y.; He, Y.; Jiang, N.; Liang, L.; Liu, L.; Liu, W.; Ma; Ma, L.; Ma, X.; Pan, T.; Shi, H.; W.L. and Gao, Z.; Wang, G.; Wang, X.; Wei, X.; Xu, Y.; Yang, B.; Yang, D.; Zhai, B.; Zhang, J.; Zhang, Z.; Zheng, Y.; Zhou, Q. |
| EPI_ISL_437602, EPI_ISL_437603, EPI_ISL_437604, EPI_ISL_437605, EPI_ISL_437607, EPI_ISL_437609, EPI_ISL_437612, EPI_ISL_437613, EPI_ISL_437615, EPI_ISL_437616, EPI_ISL_437620 | | | |
| see above | unknown | Faculty of Medicine | Buathong, R.; Bunprakob, S.; Ghai, S.; Joyjinda, Y.; Mungaomklang, A.; Petcharat, S.; Plipat; Prasithsirikul, W.; Rodpan, A.; Sirichan, N.; T. and Hemachudha, T.; Wacharapluesadee, S. |
| EPI_ISL_450212, EPI_ISL_450213 | unknown | Microbiological Diagnostic Unit Public Health Laboratory (MDU-PHL) and Victorian Infectious Disease Reference Laboratory (VIDRL) | Alpren, C.; B.P.; Ballard; C.R.; Caly, L.; Catton, M.; D.A.; Dougal, S.; Druce, J.; Duchene, S.; Easton, M.; Goncalves da Silva, A.; Hoang, T.; Horan, K.; Howden; Lane; M.B.; N.L.; S.A.; Sait, M.; Schultz; Seemann, T.; Sherry; Stinear; Sutton, B.; T.P.; Williamson; van Diemen, A. |
| EPI_ISL_450412 | unknown | Microbiology | K.-Y.; K.K.W.; To; Yuen |
| EPI_ISL_434567, EPI_ISL_434568 | unknown | Microbiology, The University of Hong Kong | K.-Y.; K.K.W. and Yuen; To |
| EPI_ISL_444518 | unknown | Molecular Infectious Disease | Anderson; B.P.; Gerasimova, A.; Kagan; R.M. and Owen, R.; Rosenthal; S.H. |
